# Potato foliar infection with *Phytophthora infestans* drives strong, cultivar-specific shifts in rhizosphere communities

**DOI:** 10.64898/2026.03.06.709792

**Authors:** Vivien Pichon, Mout De Vrieze, Fares Bellameche, Rares Cristea, Floriane L’Haridon, Laurent Falquet, Laure Weisskopf

**Affiliations:** Department of Biology, University of Fribourg, 1700 Fribourg, Switzerland; Food Research and Innovation Centre, University of Fribourg, 1700 Fribourg, Switzerland; Department of Life Sciences, University of Modena and Reggio Emilia, 42122 Reggio Emilia, Italy; Swiss Institute of Bioinformatics, 1015 Lausanne, Switzerland; Agroscope, 1260 Nyon, Switzerland

## Abstract

**Background:** Potato is an important crop worldwide, yet its production is severely threatened by *Phytophthora infestans*, the causal agent of late blight. Alternatives to the current control strategies are needed, as these rely heavily on environmentally harmful treatments. The recruitment of beneficial microbes by plants upon stress (“cry-for-help” mechanism) may represent an opportunity to find new biocontrol agents but this has not yet been reported for potato. The aim of this study was to analyse whether foliar late blight infection induces shifts in the phyllosphere, rhizosphere and soil bacterial communities associated with two potato cultivars of differing sensitivity to late blight. Moreover, we aimed at isolating members of the plant microbiota to test whether bacteria putatively recruited upon infection would be particularly active in protecting the plant against late blight.

**Results:** Controlled foliar infection triggered substantial, cultivar-specific shifts in the rhizosphere communities across two successive generations. Despite the number of differentially abundant ASVs detected being ten times higher in the second generation than in the first one, the same taxonomic groups were concerned by the shifts: *Burkholderiales*, *Flavobacteriales*, and *Bacillales*. Furthermore, the communities linked to the susceptible cultivar consistently shifted more strongly than the communities linked to the resistant cultivar. The obtained ASV sequences were used to identify 163 corresponding isolates. The inhibition potential of these strains against *P. infestans* spores was assessed through biological assays, which revealed the biocontrol potential of strains otherwise not yet known to inhibit phytopathogenic organisms, such as *Advenella*, *Nocardioides* and *Phyllobacterium* strains. Although we found no correlation between the relative abundance shift of the ASVs upon infection and the activity of the corresponding strains, we observed that the overall activity of strains isolated from the resistant cultivar was higher than that of the strains isolated from the susceptible one.

**Conclusion:** Taken together, the higher activity of the strains isolated from the resistant cultivar, along with its comparatively modest microbiome shifts upon infection suggest that the investigated resistant cultivar might harbour specific microbiota enriched in strains with efficient protective abilities against their host plant’s pathogens, which possibly contribute to its higher resistance against *P. infestans*.

## Background

Originally domesticated about 10,000 years ago in the Andes (South America), potato (*Solanum tuberosum L.*) is the 6th most produced crop, accounting for 4% of the global agricultural production [1]. It is, however, an extremely challenging culture due to its sensitivity to many diseases, one of the most devastating being late blight caused by *Phytophthora infestans*. Potato late blight is a major challenge for farmers because it is very hard to control due to its fast infectious cycle and genetic plasticity [2]. Various synthetic fungicides are used in conventional agriculture whereas in some countries, copper-based products are used in organic farming [3]. Such treatments have heavy sanitary and environmental costs, and the products used are progressively becoming banned. Finding new protective strategies against late blight is therefore urgently needed.

Microbial Biological Control Agents (mBCAs), which are microorganisms able to prevent the rise of pests or diseases could be a solution as they are believed to be a healthy and environmentally friendly way of protecting crops [4]. Current attempts to identify new mBCAs require long and costly screenings of isolated bacterial collections, which for practical reasons mostly rely on *in vitro* tests of their antagonistic activities against the pathogen(s) of interest [5–7]. In addition to being time- and cost-intensive, strains selected under these conditions often fail to provide consistent protection in whole plant assays [8], or even in field experiments. This selection of efficient mBCAs could be considerably improved if researchers could be led directly to the most interesting taxa. Recent developments in rhizosphere microbial ecology suggest the existence of an active selection of beneficial microbes by the plants themselves when they are facing challenging conditions [9]. This mechanism, called the cry-for-help hypothesis, might give researchers the much-needed lead to finding more efficient mBCAs.

Indeed, many plants have been shown to respond to various biotic and abiotic stresses by secreting specific root exudates into the rhizosphere. Some of these exudates have a direct effect on their target pathogenic organisms [10, 11], while others rather seem to recruit specific members of rhizosphere bacterial communities. Red clover for example, when facing iron deficiency, has been shown to secrete phenolic compounds, which recruited rhizosphere bacteria able to produce siderophores [12]. *Arabidopsis thaliana* was reported to secrete L-malic acid upon infection with *Pseudomonas syringae*, and this carboxylate was shown to recruit the plant growth promoting rhizobacterium (PGPR) *Bacillus subtilis* F017 [13]. Cucumber was found to secrete tryptophan when infected with *Fusarium oxysporum*, which led to the increased colonisation of the PGPR *Bacillus amyloliquefaciens* SQR9 [14]. When they were infected with the same pathogen, tomatoes secreted a cocktail of chemical compounds, some of which were shown to be antifungal, while others attracted PGPR *Bacillus* spp [15]. Some of these root exudates were also reported to activate specific genes in bacterial members of rhizosphere communities: for example, phenolic compounds secreted by barley upon infection with *Pythium ultimum* activated genes encoding the production of antifungal compounds in *Pseudomonas fluorescens* [16]. Beyond these reported effects of specific root exudates on specific members of plant-associated communities, altered root secretion following a perceived biotic stress (e.g. pathogen attack) has also been observed to lead to community-wide shifts in the relative abundance of different populations. When *Panax notoginseng* was infected with the foliar pathogen *Alternaria panax* [17], changes in relative abundance of different members of the rhizosphere communities were observed, with higher relative abundance of beneficial bacteria and fungi at the expense of pathogenic ones. These community shifts were strong enough to lead to improved germination rates of *Panax* seeds sown in the soil where infected plants had formerly grown, compared to seeds germinating in soils previously harbouring non-infected plants, and even to reduce disease severity of *A. panax* on these new plantlets.

Such evidence for targeted microbiome modulation and recruitment by the plants could be used to point us directly to particularly bioactive members of the communities, thereby reducing the screening time necessary to identify new mBCAs. Since the cry-for-help hypothesis has not yet been investigated in potato, the aim of this study was to test whether late blight infection in potato would also lead to microbiota shifts and to a rhizosphere recruitment of beneficial bacteria, with the ultimate goal to identify new and promising mBCAs against this disease. To this end, we challenged potato plants of two cultivars differing in late blight sensitivity with a controlled infection of *P. infestans*, which was meant as a signal to induce the plant’s reaction, without allowing the infection to proceed further and destroy the plants. We performed this challenge in two consecutive generations of potted plants and analysed the shifts in phyllosphere, rhizosphere and soil bacterial communities occurring as a function of the challenge (infected vs. non-infected), the cultivars (sensitive vs. tolerant) and the generation (first vs. second). In parallel to the amplicon-based community shift studies, we isolated cultivable members of bacterial communities. Those which could be associated to sequences of the amplicon-based community study were selected and assessed for their protective effects against *P. infestans* using bioassays targeting different stages of the oomycete’s life cycle.

## Material and methods

### Exposure of potato plants to P. infestans over multiple generations

The experimental setup used for this transient infection of potato plants is illustrated in Figure 1. Potatoes of susceptible cultivar Bintje and tolerant cultivar Innovator were planted in 2.2 L pots filled with soil. This standard soil was inoculated with a suspension generated by mixing soils collected from copper-free organic potato fields from three different farms in Switzerland (Muensingen 46.884010, 7.559025; Uettligen 46.985864, 7.389619; Niederbipp 47.267207, 7.707649). Bintje and Innovator plants were kept in the greenhouse during 19 and 16 days after emergence respectively and were then transiently infected with *P. infestans*. To this end, the plants were transferred to a growth chamber maintained at 16°C with 85% humidity and were sprayed with distilled water immediately before being sprayed with either a solution of *P. infestans* Rec01 zoospores (50,000 zoospores/mL) or distilled water for the control plants. To keep high humidity conditions, the plants were sprayed with distilled water every hour for 6 hours. After three days, the plants were placed in a second growth chamber at 25°C with 60% humidity for three days to stop the infection, before being transferred back to the greenhouse for 9 days. At least 3 leaflets coming from 3 different mature leaves, 5 mL of rhizospheric soil and 10 mL of bulk soil (respectively referred to as phyllosphere, rhizosphere and soil, 5 replicates each) samples were collected per pot 15 days after the infection and kept at −80°C. The plants (control and infected ones) were removed from the pots 3 days later. The remaining soil as well as the rhizospheric soil were collected and mixed with new potting soil (¼ of previous generation soil with ¾ of new soil). These mixtures were used to grow the second generation of potatoes. In the following sections, the soil mixtures coming from pots which contained infected plants will be referred to as *Phytophthora*-challenged Plant Soil (PPS) and the soil mixture coming from pots which contained non-infected plants will be referred to as Non-challenged Plant Soil (NPS).

**Figure 1:**
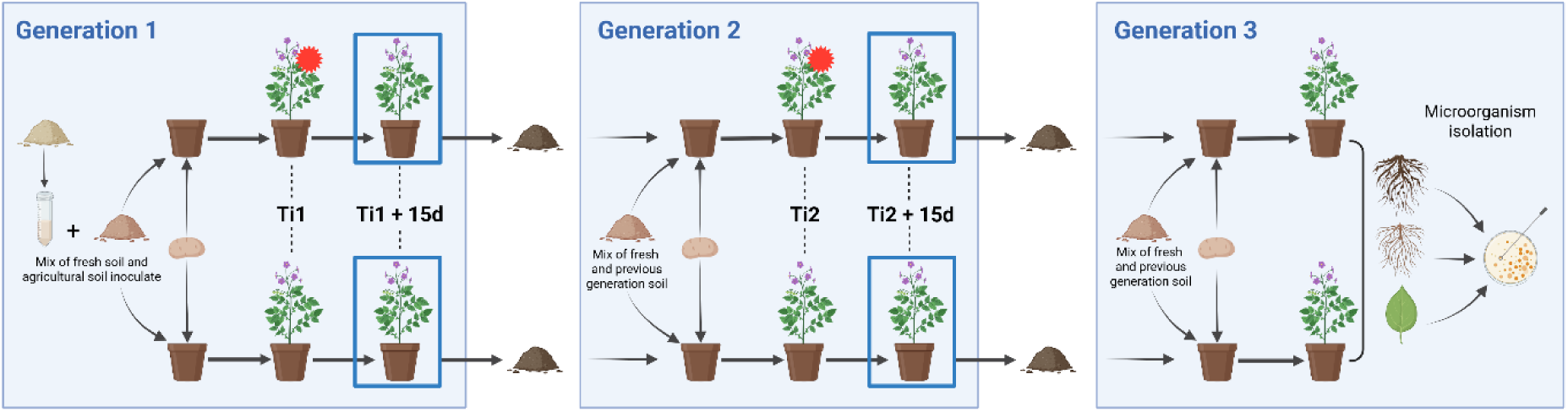
Experimental setup for transient infection of potato plants. Susceptible (Bintje) and tolerant (Innovator) cultivars were planted in pots with soil inoculated with microbial communities from copper-free organic potato fields in Switzerland. After growth in a greenhouse, plants were transferred to a high-humidity growth chamber and infected under control with either Phytophthora infestans zoospores (red star) or distilled water (control). Post-infection, plants were returned to the greenhouse. Successive generations of plants were grown in soil from infected or non-infected plants to examine microbial community shifts. Samples of phyllosphere, rhizosphere, and soil were collected from infected plants 15 days after infection (blue squares) to analyze microbial dynamics. Rhizospheric soil, roots and leaves from the plants of the third generation were used for microorganism isolation. Ti, time of infection.

The second generation of plants was meant to amplify the signal by repeating the same treatments: 15 days after emergence, plants which grew in PPS were transiently infected as described above and the plants which grew in NPS received the same control treatment. Fifteen days after infection, phyllosphere, rhizosphere and soil samples (6 replicates) were harvested and the remaining soil and rhizosphere of the plants were collected, mixed with fresh potting soil (¼ of previous generation soil with ¾ of new soil) and used for the planting of a third generation of potatoes. The third generation’s main purpose was to allow for the isolation of phyllosphere, roots and rhizosphere bacteria from plants growing in PPS.

### Microbiome samples DNA extraction and sequencing

The DNA of microbiome samples was extracted using the NucleoSpin Soil DNA extraction kit (Macherey-Nagel) according to the manufacturer’s instructions. The V5, V6 and V7 hypervariable regions of the 16rRNA were amplified using the 799F (5’-AACMGGATTAGATACCCKG-3’) and 1193R (5’-ACGTCATCCCCACCTTCC- 3’) primers [18] using the Platinum SuperFi II DNA polymerase (Invitrogen, Carlsbad, Californie) for the soil and rhizosphere samples, and the Phusion DNA polymerase (ThermoFisher, Waltham, Massachusetts) for the phyllosphere samples. The ATCC-MSA-3001 (American Type Culture Collection, Manassas, Virginia) mock community standard was used as an internal standard. The bacterial 16S rRNA were separated from the plant mitochondrial 16S by fragment size selection using the PippinHT (Sage Science, Beverly, USA). The library was prepared with the Illumina Nextera XT kit (Illumina, San Diego, USA) at the Bern Next Generation Sequencing Platform, and the sequencing was done using the Illumina MiSeq System. The raw reads are available in the European Nucleotide Archive database, accession PRJEB75732.

### NGS data cleaning and correction with DADA2

The raw reads were cleaned using the fastp 0.20.1 [19] program and their quality was checked using FastQC 0.11.9 [20] and MultiQC 1.8 [21]. The reads were then corrected using the DADA2 package [22] in R. Firstly the reads were trimmed asymmetrically at 261 bp for the forward reads and 200 bp for the reverse reads, to have a long overlap of about 45 bp, with an expected error of 3 bp between the runs. This asymmetrical trim avoided having an overlap in the V5 hypervariable region. The sequences of both runs were merged with a tolerance of up to two mismatches. In the end there were 23420 different Amplicon Sequence Variants (ASVs). The taxonomy was assigned until the Genus rank based on the Silva database, release 138 [23].

### Initial data treatment and alpha diversity analysis

The resulting read table was analysed with R. The samples of soil, rhizosphere and phyllosphere of the plants 15 days after being challenged were selected, and ASVs corresponding to chloroplasts were filtered out based on their phylogeny. At the end of this step, there were 22557 different ASVs. The rarefaction curves were computed with the rarecurve function from the vegan package [24] and plotted with ggplot (Figure S1).

The richness – defined as the number of taxa observed – and Sheldon’s evenness of each sample were calculated with vegan. The effect of the plant generation, compartment (phyllosphere, rhizosphere, soil), treatment (infected, control) and cultivar (Bintje, Innovator) on the alpha diversity were assessed by comparing the samples from the soil and the rhizosphere in a four-way ANOVA with the formula:

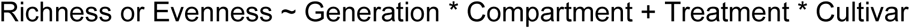

And the samples from the phyllosphere in a Generalised Least Squares (GLS) model with the formula:

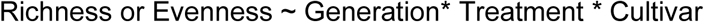

The ANOVA and GLS were followed by Tukey Honest Significant Differences (HSD) post-hoc tests to compare the richness and evenness between infected and control plants within each single compartment, generation and cultivar.

### Low-occurrence ASV removal and beta-diversity analysis

The low-occurring ASVs were filtered out by keeping only the sequences with at least 3 reads in at least 4 samples. There were 6472 ASVs remaining after this step, which represents 29% of the total number of ASVs, but 96% of the total reads from the sequencing. The filtered tables were normalised by Total Sum Scaling (TSS) with the package edgeR [25]. The TSS-normalised table was used for ordinations in order to have an overall idea of the dissimilarity between all the samples. It was firstly plotted based on Bray-Curtis dissimilarity matrix as a PCoA with the phyloseq package [26]. A Distance-Based Redundancy Analysis (dbRDA) based on a Bray-Curtis dissimilarity matrix meant to partial out the effect of the generation on the ordination and to represent more specifically the effect of the compartment was then performed with the vegan package using the formula:

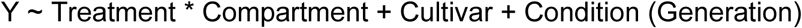

Then another dbRDA aimed at partialling out the effect of the compartment and representing more specifically the effect of the generation was performed with the formula:

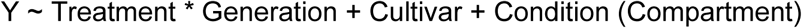

To find which ASVs became more abundant upon infection, the filtered read table was sliced into twelve subtables, each containing the samples of one generation, one compartment and one cultivar. Each subtable was then normalised by Trimmed Mean of the M Component (TMM) before going through a generalised linear model (GLM) fitness test from the edgeR package, using the model:

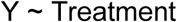

An ASV was considered differentially abundant between infected and uninfected plants if its adjusted p-value (Benjamini-Hochberg method) was below 0.05. Comparing the number of unique differentially abundant (DA) ASVs between each parameter (generation, compartment and cultivar), supported the choice to focus on the community shifts in the rhizosphere of both cultivars.

Descriptive statistics of the differential abundance of every taxonomic level in the rhizosphere of the second generation were calculated based on the TMM-normalised abundance and were all expressed in percentages (Table 1, see after the conclusion). The ratio of the abundance of each group compared to the total abundance was calculated with this formula:

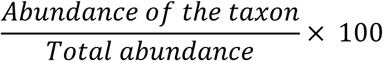

The ratio of taxa which belonged to DA ASVs compared to the total abundance was calculated with this formula:

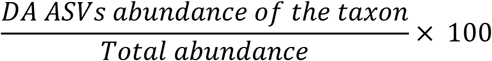

The ratio of taxa which belonged to promoted or inhibited DA ASVs compared to the abundance of their group was calculated with this formula:

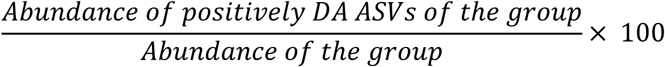

**Table 1:**
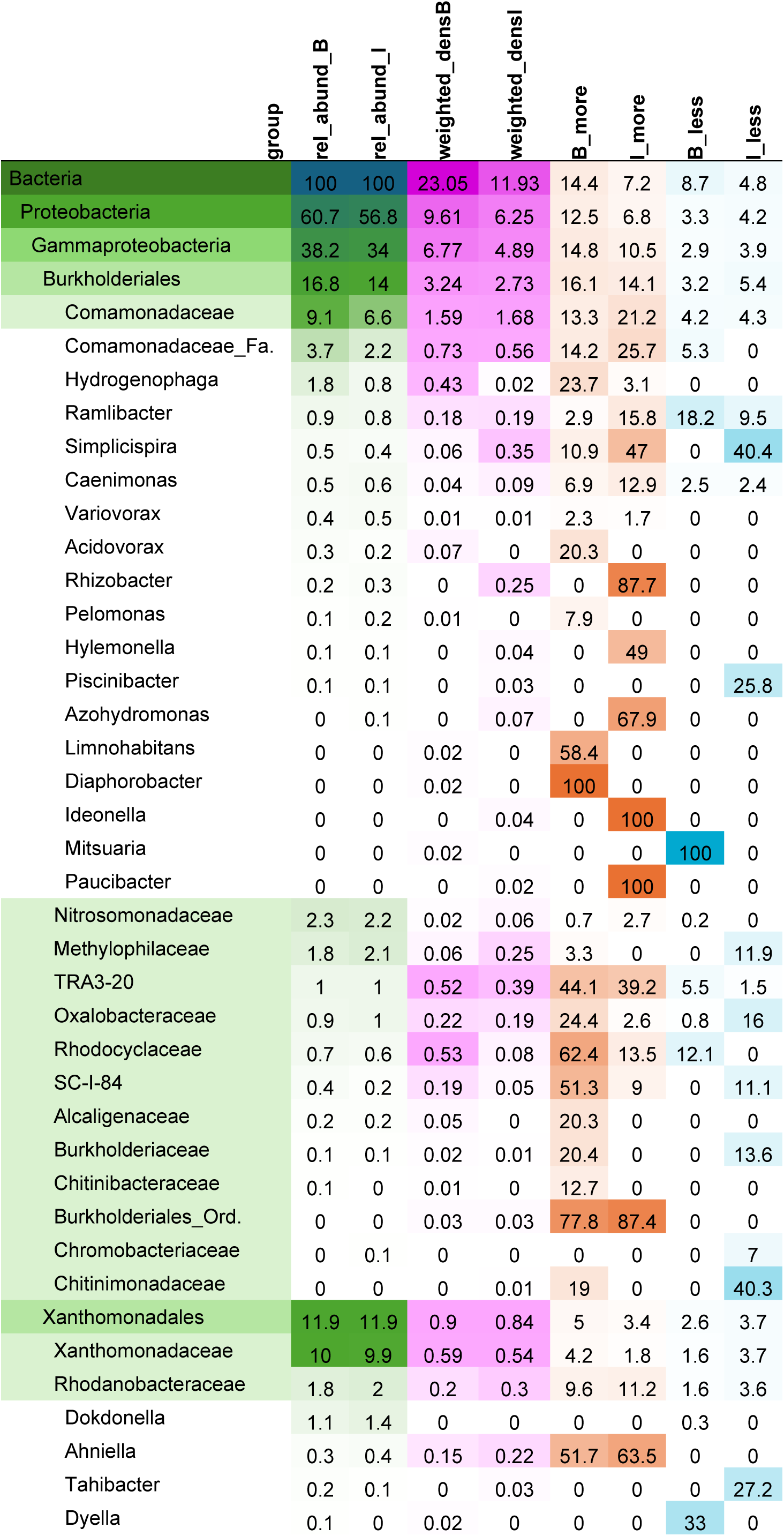
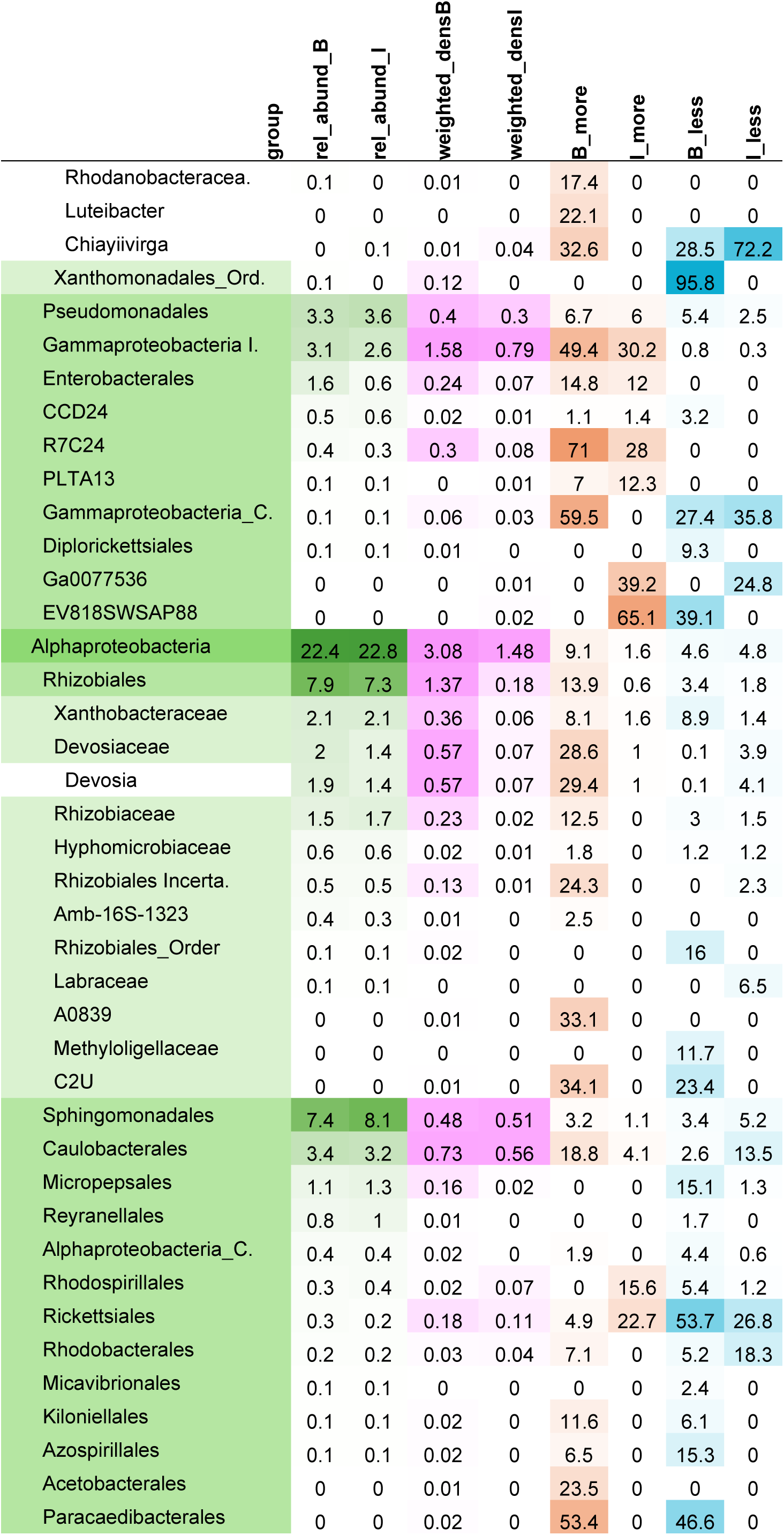
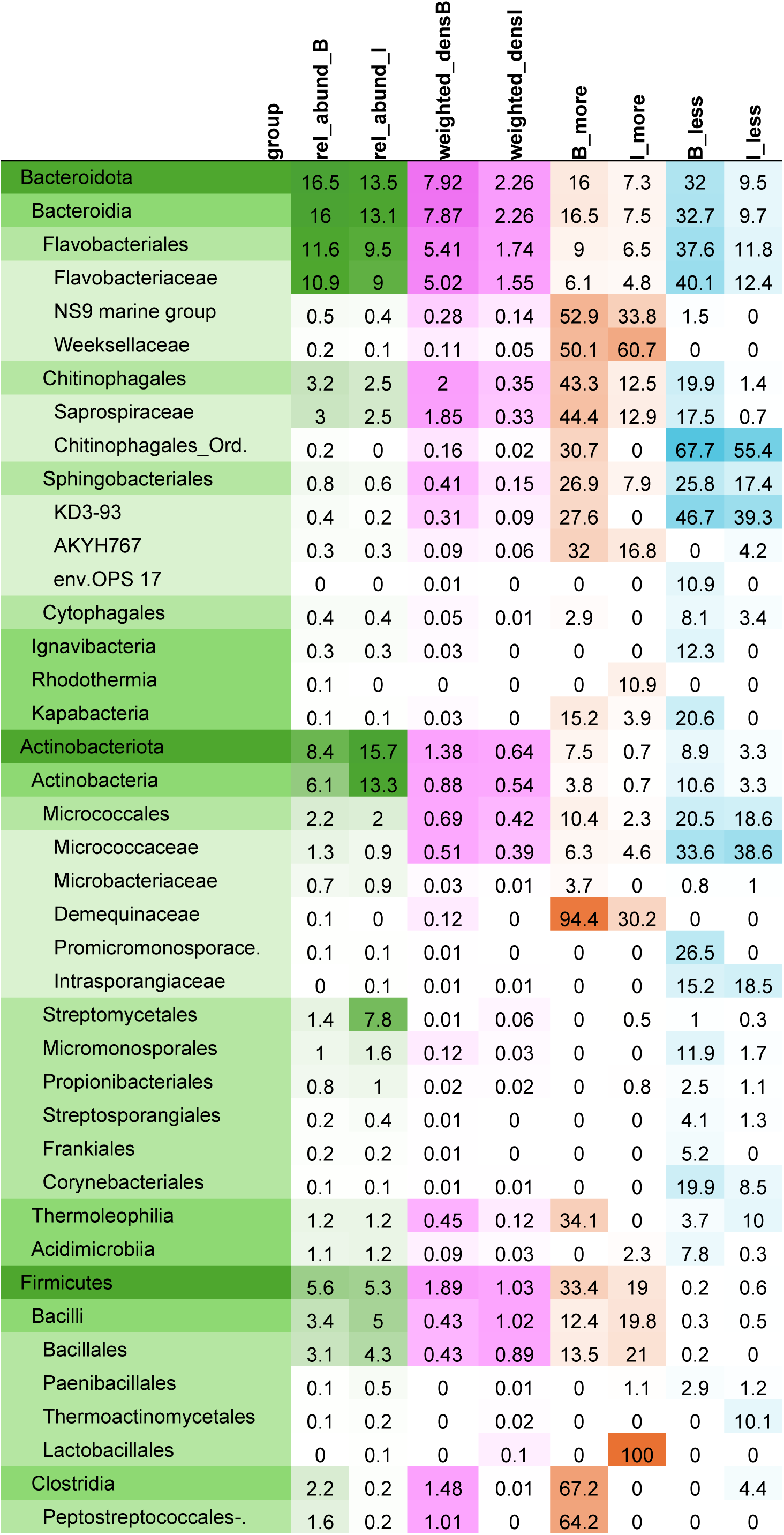
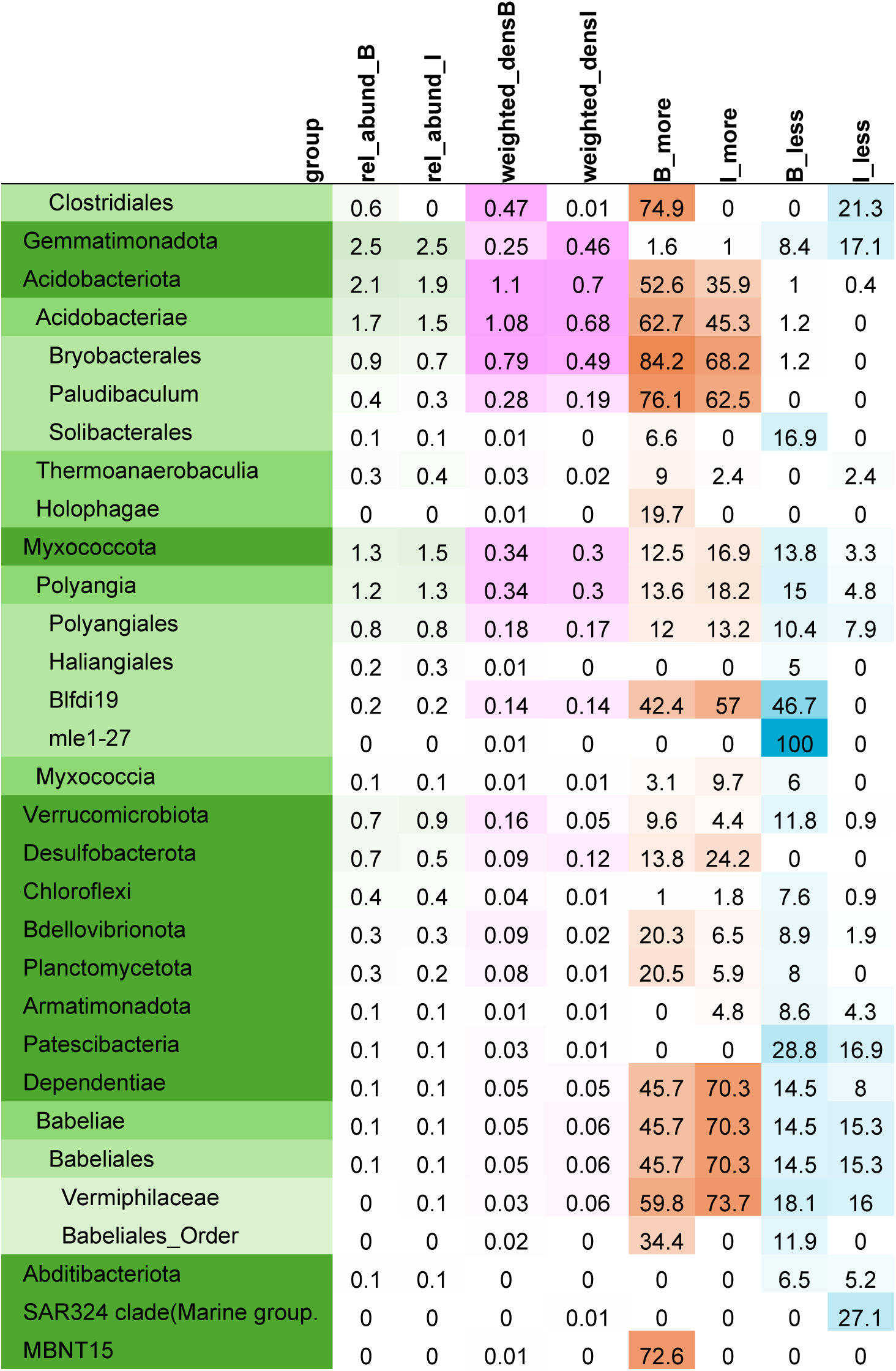
Table of the differential abundance of chosen taxa per cultivar. Descriptive statistics were calculated for every taxa of every taxonomic level present in our study, all were expressed in percentages. The last letter of the column names (B, I) refers to the cultivar. The Rel_abund columns show the abundance of each taxon as a proportion of the total abundance. The Weighted_dens columns show the proportion of the abundance of the DA ASV of each taxon as a proportion of the total abundance. The More columns represent the proportion of the abundance of the ASV which become significantly more abundant upon infection as a proportion of the abundance of the taxon. The Less columns are the same as the More columns, but for significantly less abundant ASV. Taxa shown in this table were kept if their differential abundance characteristics striking enough to be reported in the main text of this article, or if they were closely related to such a taxon. The full table is available in the supplementary materials (Table S3).

### Phylogenetic tree creation and visualisation

The ASVs used for the beta-diversity analysis were aligned with MAFFT 7.490-GCC-10.3.0-with-extensions [27] and the global phylogenetic tree was built using FastTree 2.1.11-GCCcore-10.3.0 [28]. Further work on phylogenetic trees was based on only the data and differential abundance tests of the rhizosphere of the second generation. The ASVs absent from the rhizosphere of the second generation were filtered out using the phyloseq package. Then the ggtree [29] and ggnewscale [30] R packages were used to build 1) a general tree, which summarises the edgeR analysis per order and per cultivar, and 2) several detailed trees showing the differential abundance of every single ASV per cultivar.

The general tree was drawn by agglomerating all the ASVs per order. All the calculations were made by cultivar and based on the TMM-normalised abundance. The abundance was calculated by adding the values of all the ASV of each order. The proportion of DA ASVs within each order was calculated by dividing the sum of the abundances of the DA ASVs by the sum of all the ASVs per order:

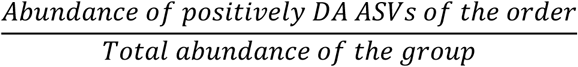

The mean normalised, weighted log-fold change of significantly differentially abundant ASVs of each order, was calculated with this formula:

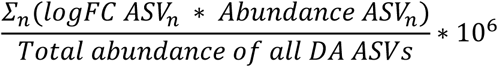

The detailed trees were drawn to represent every single ASV, their abundance and logFC. To keep it readable, the ASVs were subsetted on the basis of their phylogeny in order to have less than 900 ASVs per tree. Only the informative trees were kept (See Supplementary materials).

### Isolation and identification of microbial strains

Four different growth media supplemented with either nystatin (10 μg/mL) or ampicillin + kanamycin (respectively 250 μg/mL and 25 μg/mL), were used for the isolation of the microbial strains: 10% Tryptic Soy Agar (Sigma-Aldrich, Darmstadt, Germany) supplemented with nystatin; 20% Potato Dextrose Agar (Carl Roth, Karlsruhe, Germany) supplemented with ampicillin and kanamycin; Actinomycete Isolation Agar (Sigma-Aldrich) supplemented with nystatin and Artificial Root Exudate (ARE) [31] supplemented with either ampicillin or nystatin and kanamycin.

The isolation was made from three different compartments (leaves, rhizosphere soil and roots) in a 0.1M potassium phosphate isolation buffer at pH 7 with 0.1% Silwet. The leaflets from the leaves 2 and 3 of the healthiest-looking plants of generation 3 were ground in a mortar with 10 mL isolation buffer, then 200 μL of the ground tissue were spread either undiluted or 10-and 100-fold diluted in Petri dishes containing the different growth media. For isolation of root and rhizosphere soil communities, the plants were unrooted and shaken to remove the non-rhizosphere soil. Roots were collected from multiple locations across each plant’s root system and placed loosely into a 50-mL Falcon tube. The tubes were then filled up to 45 mL with isolation buffer, shaken and mildly sonicated (47 kHz) for 5 minutes to separate the soil from the roots. The roots were placed into a new Falcon tube, and the remaining content (soil and root wash corresponding to the rhizosphere soil) from different plants of the same cultivar were mixed together and vortexed. The supernatant was diluted from 10^-1^ to 10^-5^ and 200 μL were plated on each growth medium. The roots were washed two extra times until they appeared white and were then ground in 10 mL isolation buffer in a mortar. The ground tissues from the plants of a same cultivar were mixed together, diluted from 10^-1^ to 10^-7^, and 200 μL were plated on the different growth media.

After 5 days, single colonies were visible. Per cultivar and per isolation medium, all phenotypically distinct colonies were picked using a toothpick and transferred to a new plate for isolation and purification. In total, 636 bacterial isolates were collected. Each bacterial isolate was identified by partial 16S rRNA sequencing using the primers 27F (5’-AGAGTTTGATCCTGGCTCAG-3’) and 1492R (5’-CGGTTACCTTGTTACGACTT-3’). The strains were identified by aligning their 16S rRNA sequence on the SILVA, RDP [32] and NCBI [33] databases. The sequences are available in the European Nucleotide Archive, accession PRJEB75732.

### Identification of strains of interest for biocontrol of P. infestans

As “strain of interest”, we considered the strains for which the abundance of the corresponding ASV differed significantly between transiently infected and control plants. The DA ASVs were identified using the results of the GLM described above. All the ASVs which were significantly differentially abundant between infected and uninfected plants in at least one GLM fitness test were listed. To identify strains corresponding to these ASVs, we proceeded as follows: all 16S rRNA sequences from the isolated strains were assembled as a custom blast database with the BLAST+ (2.15.0) package for Ubuntu [34] and blasted against the list of DA ASVs previously identified using blastn (2.11.0) [35] allowing 1 mismatch and no deletion. If an ASV perfectly matched the 16S rRNA sequence of at least one strain, then all the strains which aligned with one mismatch were removed. As a result 425 strains from the collection were aligned to 74 ASVs. Thirty-five strains aligning with 7 ASVs were removed because they belonged to species listed as potent opportunistic pathogens by the Swiss Federal Office for the Environment [36]. At this stage, there were 390 strains left aligning with 67 ASVs, among which 144 strains perfectly matching with 39 ASVs and 246 strains aligning with 1 mismatch on 28 ASVs. For the biological assays, all the strains matching perfectly any ASV were kept, but only one strain per ASV was kept for the ASVs which aligned with one mismatch, while excluding the filamentous bacteria (e.g. *Streptomyces* sp.) which would not fit the high throughput screening described below. This resulted in the selection of 163 strains aligning with 57 ASVs, which were screened for their biocontrol potential against *P. infestans*.

### Evaluating the biocontrol potential of the strains of interest

#### Preparation of P. infestans zoospore and sporangia suspensions

##### Sporangia suspension

*P. infestans* 208M2 (GFP-tagged) [37] was grown on V8 medium (10% V8 juice supplemented with 1 g/L of CaCO3 and 15 g/L of agar) for 10 to 21 days. Mycelium was then scrapped off the agar and put into a 15 mL Falcon tube, to which sterile distilled water was added until reaching 2 mL. The tube was shaken vigorously to detach the sporangia from the hyphae. The solution was then filtered using a nylon mesh. The sporangia concentration was determined using a Thoma chamber and adjusted to 30,000 sporangia/mL.

##### Zoospore suspension

*P. infestans* 208M2 was grown on V8 medium for 10 to 14 days. The Petri dishes were then wrapped in aluminium foil and 15mL of ice-cold water was poured into them. They were placed at 4°C for 2h and then incubated at room temperature for 30 minutes. After incubation, the liquid was pipetted into a 15 mL Falcon tube coated in aluminium foil. The concentration of zoospores was determined using a Thoma chamber and diluted to 30,000 zoospores/mL.

#### High throughput P. infestans in vitro confrontation assays

##### Bacteria preparation

The selected bacteria were incubated during 24h at 28°C in 200 μL of filtered liquid V8 medium (autoclaved 10% V8 juice supplemented with 1 g/L of CaCO3, filtered using 0.22 μm sterile syringe filters) in 96-well plates. The optical density at 600nm (OD600) of each well was measured using a Cytation5 cell imaging reader (Agilent, Santa Clara, USA). The sporangia and zoospore germination assays were performed at least 3 times per strain.

##### Sporangia germination assays

Fifty μL of the bacterial suspensions were transferred to a new 96-well plate containing 50 μL of sterile 0.9% NaCl solution. Forty μL of 208M2 sporangia suspension were added to the wells. The plates were sealed using parafilm and placed in a box with wet paper towel sheets during 24h at 18 °C until imaging.

##### Zoospore germination assays

Fifty μL of the bacterial suspensions or sterile distilled water was transferred to a new 96-well plate containing 50 μL of sterile NaCl solution (0.9%). Forty μL of 208M2 zoospore suspension were added to the wells. The plates were sealed using parafilm and placed in a box with wet paper towel sheets during 4h at 18 °C until imaging.

##### Imaging, scoring and data treatment

Imaging was performed using a Cytation5 plate reader. For each well, a bright-field image and a GFP fluorescent image (excitation at 395nm) were taken. A germination score from 0 to 4 was given manually for each well. The scores for sporangia germination are 0 for no germination; 1 for low number of small germ tubes; 2 for low number of long germ tubes; 3 for a higher number of middle-sized germ tubes; 4 for full germination. The scores for zoospore germination were 0 for no germination, 1 for few and short germ tubes, 2 for a higher number of middle-sized germ tubes, 3 for full germination with no appressoria, 4 for full germination. The adjusted score was calculated by dividing the germination score of each strain by the average germination score of the distilled water controls of their respective 96-well plate. The adjusted OD600 was calculated by removing the average OD600 of the controls of each 96-well plate from the OD600 of each strain. The adjusted OD600 and adjusted score were used to calculate the germination index using the following calculation:

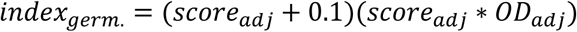

A germination index below 0.25 indicates strong inhibition, while an index between 0.25 to 0.5 signifies moderate inhibition, am index between 0.5 to 0.9 suggests a weak inhibition, and indexes between 0.9 and approximately 1.8 indicate no inhibition. Indexes higher than 1.8 imply that germination was stimulated. The score per strain was calculated by averaging the germination indexes of all the replicates.

The strains were then grouped by phylogenetic group at every taxonomic level (Phylum, Class, Order, Family, Genus). The groups with less than 3 strains were discarded and the rest was compared with a Kruskal-Wallis test followed by Dunn’s post-hoc test (p-value adjusted with the Benjamini-Hochberg method). The strains were then grouped by cultivar and compartment of isolation and were compared by a two-way ANOVA with Type II Sum of Squares, followed by Tukey’s HSD.

The average index for each ASV was determined by averaging the indexes of all associated strains. ASVs were then classified as “MORE” if they showed a significant increase in abundance upon infection in at least one GLM test (see *Low-occurrence ASVs removal and beta-diversity analysis* section above), and “EQUAL” otherwise. The scores of the “MORE” and “EQUAL” groups were compared using a Kruskal-Wallis test. A similar comparison was conducted for strains that decreased in abundance.

## Results

### Transient P. infestans infection did not affect the alpha diversity of bacterial communities

The richness of bacterial communities was affected by both the compartment and the generation when comparing the soil and the rhizosphere. As expected, the soil communities were significantly richer and displayed higher evenness than the rhizosphere communities (Table S1, Figure S2). We also observed that communities were richer in the second generation than in the first one for both compartments. However, neither the treatment (transient infection vs. control) nor the cultivar affected the richness of rhizosphere bacterial communities.

When compared with the soil or the rhizosphere, communities living in the phyllosphere had much lower richness, but relatively high evenness. Richness was not affected by any factor, but evenness was significantly higher in the second generation compared to the first one, and lower in infected plants compared to non-infected ones.

Although no global trends could be observed, transient *P. infestans* infection altered the richness and evenness of subgroups of this experiment in four instances (Figure S2). These four significant changes all occurred in the first generation, and three of them concerned the sensitive cultivar Bintje: in the phyllosphere, which is the site of infection, richness decreased in infected plants compared to control plants, while the reverse tendency was observed in the soil, but not in the rhizosphere. Surprisingly, although richness decreased in leaf communities upon infection, evenness increased, which also occurred in rhizosphere communities. The only impact of transient infection on communities associated with the resistant cultivar Innovator was an increased evenness in the soil communities. Overall, the communities associated with this cultivar reacted less strongly to infection than those associated with the sensitive cultivar (Figure S2).

### Leaf infection had a strong cultivar-dependent impact on the beta diversity of soil and rhizosphere communities

According to the results of the PCoA, all samples grouped into three major clusters (Figure 2a), one for the phyllosphere samples, and one for each generation for the soil and rhizosphere samples. In phyllosphere samples (Figure 2b), communities differed slightly between infected and control plants in the first generation, while such differences could not be observed in the second generation. The soil and rhizosphere samples of the first generation (Figure 2c) clustered according to their compartment, then according to their infection status. As expected, the difference between communities from infected and non-infected plants was greater within the rhizosphere than within the soil communities. The soil and rhizosphere samples of the second generation (Figure 2d) also clustered according to their compartment and infection status but with a remarkably similar clustering pattern between soil and rhizosphere samples. In contrast, much greater differences were observed in the effect of infection on both cultivars than in the first generation: in both soil and rhizosphere communities, those associated with the sensitive cultivar Bintje reacted much more strongly to infection than those associated with the resistant cultivar Innovator (Figure 2d, circles vs. diamonds). This stronger impact of infection on communities from the second generation was further highlighted by the distance-based redundancy analysis partialling out the compartment (Figure S3a), while partialling out the generation (Figure S3b) revealed that shifts in the soil and the rhizosphere communities followed a remarkably similar pattern. In both cases, communities from infected Bintje plants were clearly separated from the other communities.

**Figure 2:**
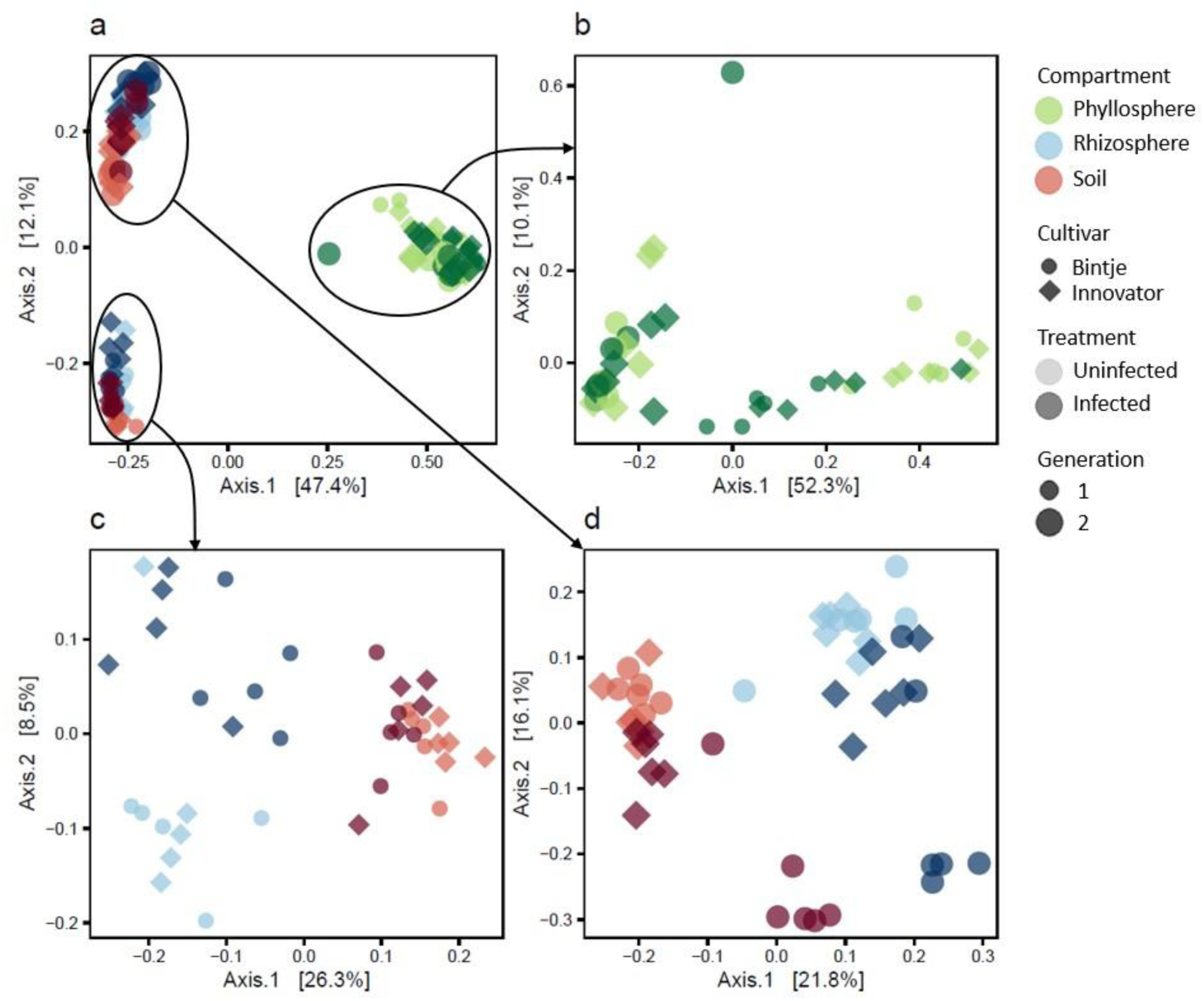
Principal Coordinate Analysis of a) all bacterial communities, b) phyllosphere communities from both generations, c) rhizosphere and soil communities from the first generation, and d) rhizosphere and soil communities from the second generation.

### The phylum composition of phyllosphere and rhizosphere communities changed between generations

The *Proteobacteria* were the most abundant phylum in every compartment (Figures S4 and S5), followed by the *Actinobacteriota*, the *Bacteroidota* and the *Firmicutes*. The *Proteobacteria* were particularly dominant in the phyllosphere of the first generation where they accounted for 87% of the total community (Figures S4 and S5a). This proportion dropped significantly during the second generation where it went down to 68%, mostly due to the proportion of *Firmicutes* booming from 7% to 21%. This drop in *Proteobacteria* in the second generation was not observed in the rhizosphere and soil, where it rose from below 50% to almost 60% (Figure S5 b and c). In the soil, the increase in relative abundance of *Proteobacteria* was compensated mainly by a drop in *Actinobacteriota*, which fell from 23% to 12%, while the *Bacteroidota* and *Firmicutes* remained stable at about 6% and 10%. In the rhizosphere, the increase in *Proteobacteria* was mostly compensated by a drop in the *Firmicutes* population from 15% to 5% in the second generation, while the proportion of *Actinobacteriota*, *Bacteroidota* and other phyla all dropped by 1 to 2% each. The phylum distribution was much more even in the rhizosphere than in the soil in the first generation, but these compartments became remarkably similar in the second generation.

While the infection had a limited effect on the soil phylum composition (Figure S5c), it led to stronger shifts in compartments closely associated with plants. While the phyllosphere phylum composition of Innovator did not change upon infection in the first generation (Figure S5a), the *Actinobacteriota* population in Bintje collapsed from 18% to 4% of the total abundance, which led to an increase in the proportion of *Firmicutes* and *Proteobacteria*. In the second generation, the phyllosphere population in Bintje were stable, whereas in Innovator, *Proteobacteria* increased upon infection at the expense of *Firmicutes* and *Actinobacteriota*. The infection-induced shifts observed in the rhizosphere were common to both cultivars in the first generation (Figure S5b), with an increase in *Proteobacteria* and *Bacteroidota* at the expense of *Firmicutes*, which collapsed. Later, in the second generation, the phyla composition in Innovator hardly changed upon infection, while in Bintje, *Firmicutes* and *Proteobacteria* increased a the expense of the *Bacteroidota* and *Actinobacteriota*.

The number of differentially abundant (DA) Amplicon Sequence Variants (ASVs) was much higher in the second generation (2191) than in the first one (192), with 81 shared DA ASVs. (Figure 3a). More DA ASVs were detected within Bintje-associated communities (1475 specific, 1821 total) than within Innovator-associated ones (481 specific, 827 total) (Figure 3b), with a large majority of them in the soil (1149 specific, 1791 total), less in the rhizosphere (507 specific, 1145 total) and very few in the phyllosphere (4 specific, 8 total) (Figure 3c).

**Figure 3:**
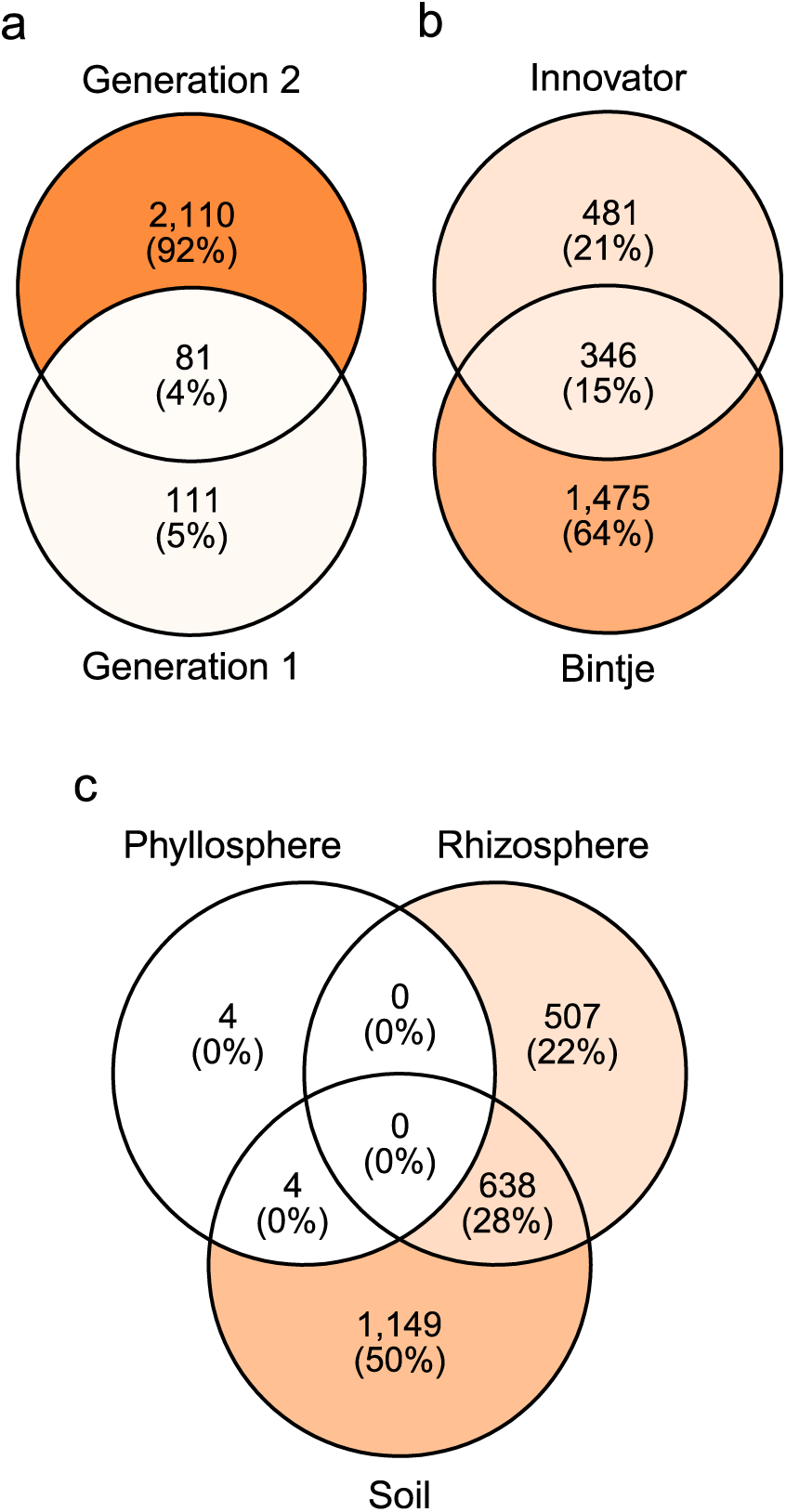
Venn diagrams summarizing the number of differentially abundant ASV detected in the GLMs for microbiome analysis (Table S1a). Each diagram shows the same data split across the different factors: (a) the generation, (b) the cultivar and (c) the compartment. Within a diagram, each set represents the result of several GLM grouped together depending on the samples used in it; each GLM was used in exactly one set. Each set is a list of all the ASV which were differentially abundant in at least one GLM. The numbers in each section of the diagram represent the number and percentage of DA ASV, which were shared between sets or unique to one.

According to these results, shifts in bacterial communities following transient infection with *P. infestans* occurred almost exclusively in the soil and rhizosphere, although infection occurred in leaves. The few observed in the phyllosphere were either driven by significant shifts on very abundant ASVs with a strong impact on community structure, or they were non-significant at the ASV level, leaving them absent from the differential abundance analysis, yet visible at the phylum level. Further analyses were therefore focused on the rhizosphere communities, in view of the higher consistency of their shifts across generations.

### Rhizosphere communities shifts upon infection were stronger in the sensitive cultivar and amplified in the second generation

The bacterial community of the rhizosphere shifted in a cultivar-dependent way in both quantity, which is the percentage of ASVs that belonged to DA ASVs, taking into account their abundance, and in quality, which is whether an ASV was promoted (became more abundant upon infection) or inhibited (became less abundant upon infection).

In the first generation, 12% of the reads were inhibited in Bintje, and 0.1% were promoted, while Innovator had only 1.4% promoted and 0.6% inhibited (Table S2). Most of the inhibited ASVs in Bintje were detected in the *Firmicutes* (12% of the total abundance), with a sharp decline in abundance, while few ASVs shifted significantly in other major phyla. The shifts in Innovator were spread more evenly across the major phyla with 1% in the *Proteobacteria*, 0.6% in the *Firmicutes* and 0.4% in the *Bacteroidota*. In both cultivars, most of the DA ASVs were concentrated in a single order per phylum. Shifts in the *Proteobacteria* were mostly in the *Burkholderiales*, the shifts in the *Firmicutes* were mostly in the *Bacillales*, and the shifts in the *Bacteroidota* were mostly in the *Flavobacteriales*.

Considerably more DA ASVs were detected in the second generation than in the first one. The changes were again much more pronounced in Bintje, where 23% of all ASVs were differentially abundant, than in Innovator where it was 12% (Table 1). However, unlike in the first generation, more DA ASVs were promoted (14% of ASVs in Bintje and 7 % in Innovator) than inhibited (9% in Bintje and 5% in Innovator). The most quantitatively significant shifts happened in the most abundant taxa. In the *Proteobacteria*, all the DA ASVs taken together accounted for 10% of all reads in Bintje and only 6% in Innovator (Table 1), while they accounted respectively for 8% and 2% of ASVs in the *Bacteroidota*. A remarkably small fraction of the *Actinobacteriota* exhibited significant shifts, with their differentially abundant ASVs representing only 1% of the total reads in both cultivars. In contrast, a larger proportion of *Firmicutes* showed shifts, with differentially abundant ASVs constituting 2% of the total reads in Bintje cultivar and 1% in Innovator.

The differential abundance results were summarised into trees, displaying the abundance, the proportion of DA ASVs and the direction and magnitude of the shifts for each order (Figure 4). In addition, subtrees showing the abundance and post infection log-fold change for every ASV of selected taxa per cultivar were created (Figures S6 to S13).

**Figure 4:**
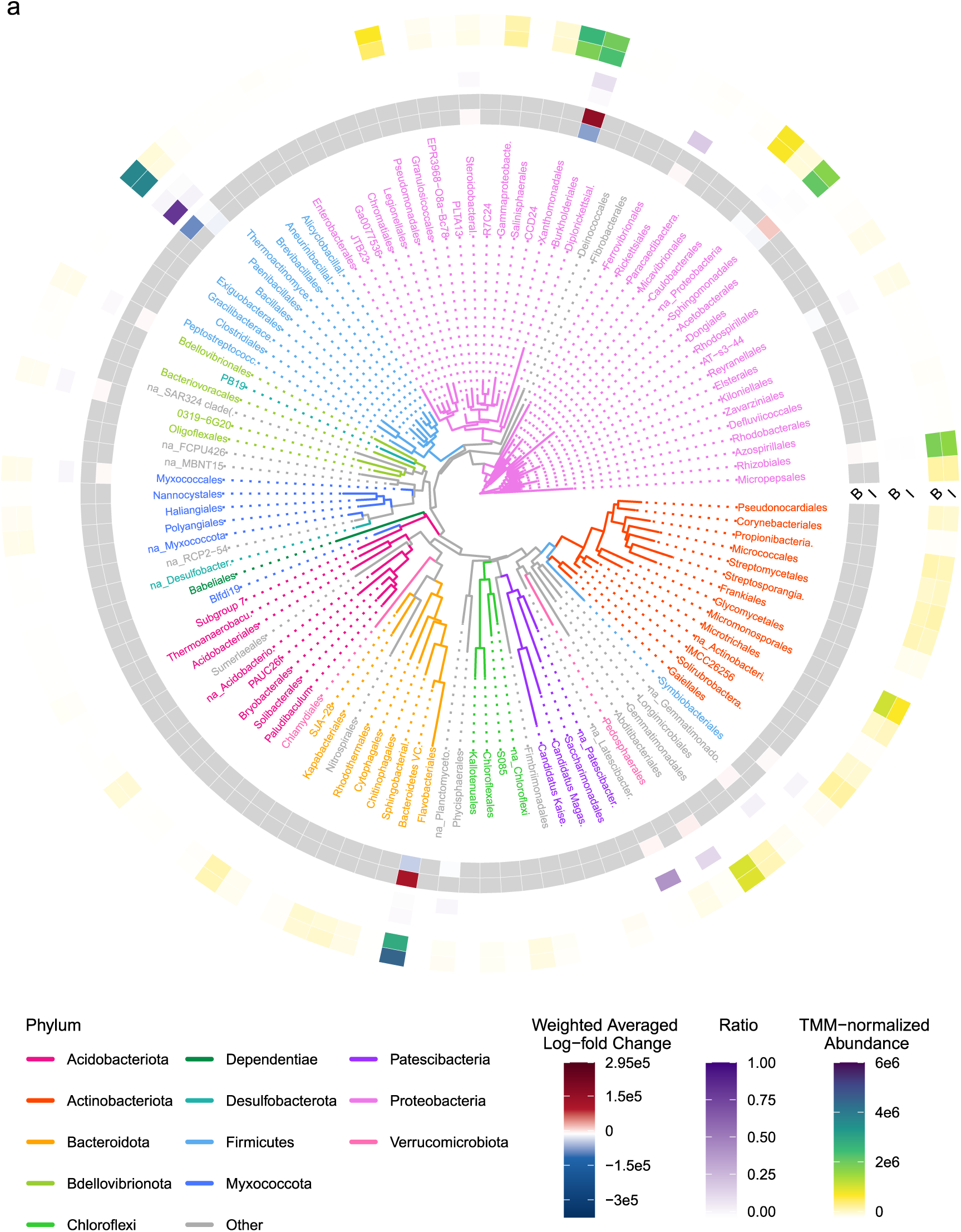
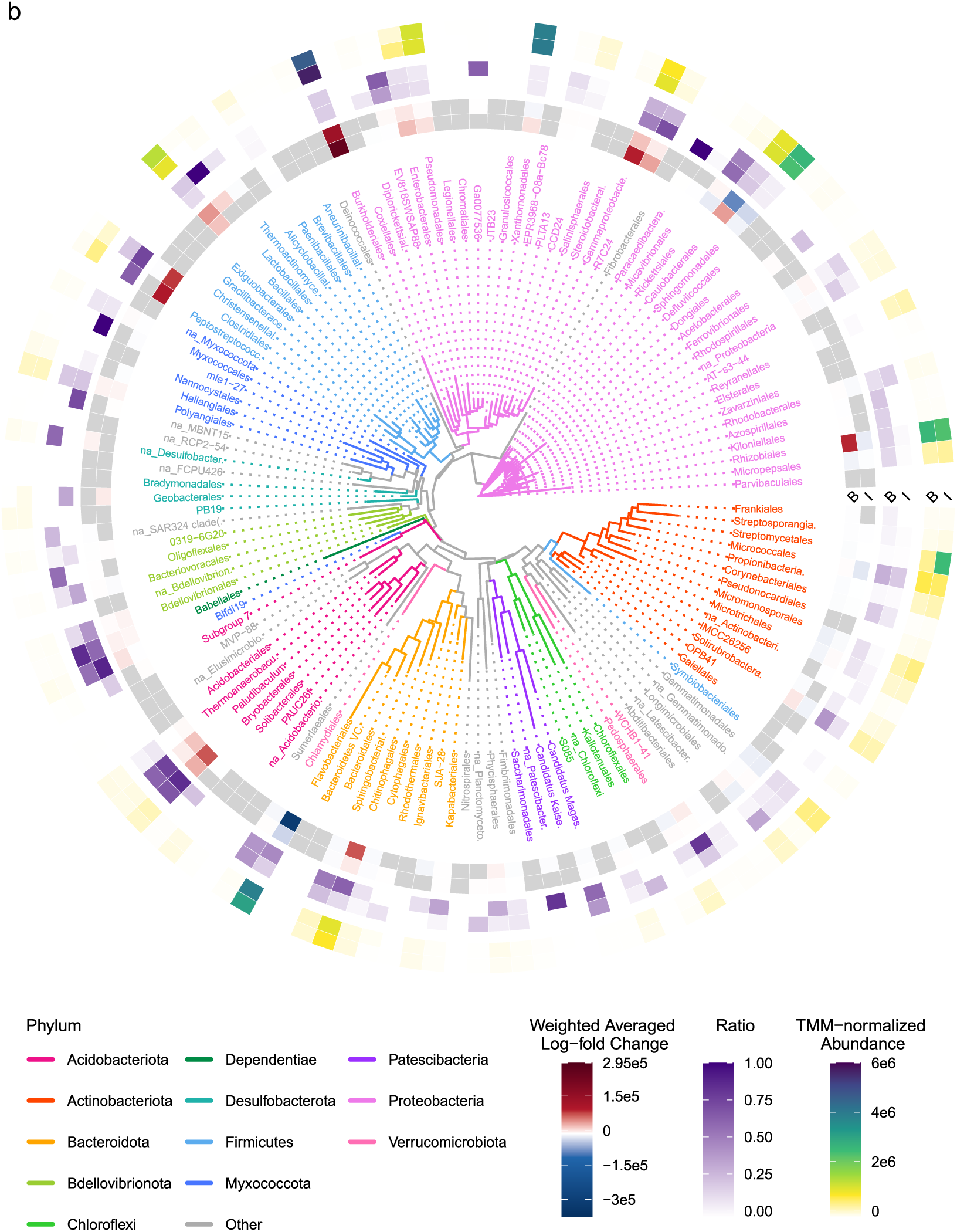
Phylogenetic trees representing the distribution of differentially abundant ASV across the orders and per cultivar in the first (a) and second (b) generation. Each tip represents an order, and their colours represent the phyla they belong to. The tree is surrounded by three annotation circles, the innermost half of which represents the Bintje (B) cultivar, and the outermost half represents Innovator (I). The outer annotation circle represents the TMM-normalized abundance of each order. The middle annotation circle represents the proportion of DA ASV weighted by the abundance of each ASV. The inner annotation circle represents the influence of the infection on the abundance the DA ASV of each order. It was calculated by weighting the mean log-fold change of the DA ASV of each order by the abundance of each ASV and normalizing it by the total abundance of all DA ASV (please see material and methods for more details). For example, at the 3 o’clock position, Rhizobiales were highly abundant (indicated by green squares). While a small fraction of this group exhibited differential abundance in the Bintje cultivar (purple square), the DA ASV showed a substantial increase in abundance on average (red square). In contrast, the Innovator cultivar displayed a minimal proportion of differential abundance (pale purple square), resulting in minimal average logFC difference (pale blue square).

The differences at the order level highlight the cultivar-dependent nature of the microbiome shifts observed. Very few orders displayed DA ASVs in the first generation, and only the *Bacillales* in Bintje had a high proportion of DA ASVs (dark purple, Figure 4a). Most of the orders with a low proportion of DA ASVs had near-neutral weighted averaged logFC values (pale red and pale blue squares in Figure 4a) except for the *Burkholderiales* and *Flavobacteriales*, which were strongly promoted in a few ASV in Innovator (dark red, Figure 4a), and inhibited in Bintje (blue, Figure 4a). The second generation had considerably more DA ASVs (Figure 4b), but most orders which contained DA ASVs had near-neutral weighted averaged logFC. Only two orders had a strong, consistent response between cultivars: *Burkholderiales* (*Proteobacteria*), which was the most abundant order of the study, and *Bryobacterales* (*Acidobacteriota*). In Innovator, only one additional order, *Caulobacterales* (*Proteobacteria*), shifted sharply and became less abundant. In contrast, Bintje exhibited strong shifts in three major orders, such as *Rhizobiales* (*Proteobacteria*) and *Chitinophagales* (*Bacteroidota*), which increased in abundance, and *Flavobacteriales* (*Bacteroidota*), which decreased. Additionally, several minor orders showed significant shifts in Bintje such as *Paludibaculum* (*Acidobacteriota*), *Peptostreptococcales* (*Firmicutes*), *Clostridiales* (*Firmicutes*), *R7C24* (*Proteobacteria*) and unassigned *Gammaproteobacteria*. The cultivar-dependent nature of the shifts appeared even more clearly in the detailed trees where we saw that differing ASVs were differentially abundant even when trends appeared similar at higher taxonomic level, such as in the *Firmicutes* in the first generation (Figure S6) or in the *Gammaproteobacteria* in the second generation (Figures S7 and S8). Bintje and Innovator had no ASV shifting in the same direction in the first generation, and only 4% of their DA ASVs in the second generation.

### Proteobacteria

Like most taxa, the *Proteobacteria* displayed weaker shifts in the first generation than in the second one.

In the first generation, very few ASVs with differential abundance were observed in the *Alphaproteobacteria* despite their prevalence in the community. In the second generation, however, they displayed clear opposite cultivar-dependent trends. While *Alphaproteobacteria* in Bintje were overall promoted with 9% of promoted, and 5% of inhibited taxa, they were mostly inhibited in Innovator with 2% of promoted and 5% of inhibited taxa (Table 1). These differences were mainly due to the *Rhizobiales* and the *Caulobacterales*, which were strongly promoted in Bintje and did not shift or were inhibited in Innovator. Additionally, in Bintje, 29% of *Devosia* ASVs were promoted, with 0.1% showing inhibition. In contrast, Innovator displayed a different pattern, where 4% of ASVs were slightly inhibited and only 1% were promoted. A similar cultivar-dependent trend was observed for *Rickettsiales*: in Bintje, 53% of ASVs were strongly inhibited and only 5% were promoted, while the response of *Rickettsiales* in Innovator was more balanced with 23% of ASVs promoted and 27% inhibited (Table 1).

On the other hand, *Gammaproteobacteria* displayed differential abundance during the first generation, essentially in the *Burkholderiales* where respectively 2% and 12% of the order’s reads were more abundant in Bintje and less abundant in Innovator (Table S2). In the second generation, the *Burkholderiales* shifted similarly in Bintje and in Innovator, with respectively 16% and 14% of the order reads that were promoted, and 3% and 5% that were inhibited (Table 1). Infection affected ASVs across the whole order (Figure S7) with a particularly strong response in the *Comamonadaceae* and *Rhodocyclaceae* families, but without particularly dense differential abundance clusters. Differential abundance clusters were found among other taxa of the *Gammaproteobacteria* despite their lower abundance (Table 1, Figure S8). Although they generally hardly shifted upon infection, a clear and dense cluster was observed in the *Xanthomonadales*, corresponding to the *Ahniella* genus, where 52% and 64% of reads were promoted in Bintje and Innovator respectively (Table 1, Figure S8). More DA ASV clusters were observed to be specific to the Bintje cultivar, such as in the *Enterobacterales*, the R7C24, or in a group of unassigned *Gammaproteobacteria* (*Incertenae sedis*) (Figure S8).

### Bacteroidota

Similarly to what happened in the *Proteobacteria*, the displayed less DA ASVs in Bintje than in Innovator (respectively 1% and 2% of the group abundance) in the first generation, and most of the DA ASVs in Bintje were inhibited while most of the DA ASVs in Innovator were promoted. The differential abundance was detected almost exclusively in the *Flavobacteriales* (Table S2).

By contrast, much more DA ASV was observed in the second generation, with a stronger response to infection in the sensitive cultivar (16% and 32% of respectively promoted and inhibited reads in Bintje) than in the tolerant one (7% and 10% of respectively promoted and inhibited reads in Innovator). Interestingly, each order of the phylum had a different reaction to the infection (Figure S9): Like in the first generation, the *Flavobacteriales* was the order with the strongest inhibition in Bintje (Figure 4) with 38% inhibited reads (Table 1). They were composed of three families with very contrasted shifts upon infection: The *Flavobacteriaceae* were strongly inhibited in Bintje and reacted weakly in Innovator, while the *NS9 marine group* and the *Weeksellaceae* were strongly promoted in both cultivars. The *Chitinophagales* observed were mainly *Saprospiraceae* and showed a strong response in Bintje, with 43% of ASVs promoted and 20% inhibited, while their response was less pronounced in Innovator, where 13% of ASVs were promoted and only 1% inhibited (Table 1). In contrast, the *Sphingobacteriales* were balanced between promoted and inhibited ASVs mainly in Bintje (Table 1, Figure S10). This apparent balance originates from the two most important families of this order having opposite reaction, i.e. the KD3-93 group was inhibited in both cultivars while the *AKYH767* group was promoted in both groups, with an entire branch promoted specifically in Bintje. Lastly the *Cytophagales* showed hardly any feedback to the infection.

### Firmicutes

The *Firmicutes* had by far the strongest DA ratio of the first generation as their population collapsed in Bintje, with 78% of inhibited reads (Table S2), mainly coming from a few clusters of very abundant ASVs among the *Bacillales* (Figure S6). On the other side, only 3.7% of the reads were inhibited in Innovator, despite a strongly reduced abundance (Figure S5). The decrease in the Innovator population was carried by many non-significantly differentially abundant ASVs.

In contrast to the first generation, the *Firmicutes* were mostly promoted in infected plants in the second generation, with stronger changes in Bintje (Figure S11, Table 1). While the *Bacillales* reacted less prominently in Bintje than in Innovator, with respectively 13% and 21% of promoted reads, the *Clostridia* were very strongly promoted in Bintje (67%) and showed an opposite tendency in Innovator, where no ASV was promoted and 4% were inhibited (Figure S11, Table 1).

### Actinobacteriota

The *Actinobacteriota* were remarkably conserved in both generations. In the first generation, there was no differential abundance in Bintje and 3.7% of the *Gaiellales* order belonged to promoted ASVs in Innovator (Table S2).

In the second generation, the phylogenetic tree showed a few scattered DA ASVs, with an unusual majority of inhibited ASVs in Bintje. One cluster was still observable among the *Micrococcales* (Figure S12). Here too, different subgroups showed different, cultivar-dependent responses to infection, e.g. the *Micrococcaceae* were inhibited in both cultivars and the *Demequinaceae* were promoted in Bintje only (Table 1).

### Minor Phyla

No notable shift was observed upon infection in the less abundant phyla of the first generation (Table S2), but several taxa showed a strong reaction in the second generation (Figure S13, Table 1). The *Acidobacteriota* reacted significantly: both of its major orders, *Bryobacterales* and *Paludibaculum*, were highly promoted upon infection in both cultivars (Figure S13). Within the *Myxococcota*, the order *blfdi19* had two peculiarities: first, it was located far away from the other *Myxococcota* on the phylogenetic trees (*Myxococcales*, Figures 4 and S13), and second, it was the only group which responded with strong, balancing shifts in Bintje (42% promoted, 48% inhibited) but with a high promotion in Innovator (57% promoted and no inhibition). The *Dependentiae* only contained ASVs of the *Babeliales* order. These reacted very strongly in both cultivars, with 46% and 70% of promoted reads in Bintje and Innovator respectively, and 15% and 8% of inhibited reads in the respective cultivars (Figure S13, Table 1).

In summary, infection triggered substantial, cultivar-specific shifts in the rhizosphere microbial communities of both generations, although they differed in nature. Despite the number of DA ASVs detected being ten times higher in the second generation than in the first one, both generations had many common points. Firstly, Bintje shifted more strongly than Innovator in both generations, with most of the shifts being concentrated in a few orders: *Burkholderiales*, *Flavobacteriales*, and *Bacillales*, while the *Actinobacteriota* stayed remarkably stable. Each phylum comprised numerous taxa with distinct responses, emphasising the importance of detailed analysis for this type of study. Beyond characterising community shifts following infection, the sequences of DA ASVs were used to identify corresponding bacteria among the strains that were isolated from the third generation of potatoes.

### Anti-Phytophthora activities of strains isolated from leaf-infected plants

From the retrieved isolates, the 16S sequence of 163 strains could be matched to ASVs and corresponded to 57 ASVs (Table S4). These strains comprised 90 *Proteobacteria*, 48 *Firmicutes*, and 25 *Actinobacteriota*. Among these, 14 strains originated from leaves, 74 from roots, and 75 from rhizosphere soil. 88 strains were isolated from Innovator and 75 from Bintje. The ASVs linked to these strains frequently exhibited differential abundance upon infection, with 21 strains becoming more abundant in at least one test, 8 becoming less abundant in at least one test, and 32 remaining unchanged across all tests. The differential abundance of these ASVs varied across compartments: none became differentially abundant in the phyllosphere, 17 did so in the rhizosphere, and 18 in the non-rhizosphere soil. Nine of the selected ASVs exhibited differential abundance in the first generation, while 21 did so in the second generation. Additionally, 21 of these ASVs became differentially abundant in Bintje, vs. 9 did so in Innovator. These strains were tested for their potential inhibition of the germination of the two types of spores produced by *P. infestans,* sporangia and zoospores.

### The ability to inhibit spore germination depended on the cultivar the strains were isolated from and on their phylogenetic identity

Overall, strains were much more efficient in inhibiting sporangia than zoospores, with average germination indexes of 0.46 for sporangia, i.e. a moderate inhibition against 0.8 for zoospores, i.e. a weak inhibition. The most efficient genera against sporangia germination mostly belonged to *Proteobacteria*, e.g. *Phyllobacterium*, *Advenella* and *Pseudomonas*, with respective median indexes of 0.18, 0.21 and 0.31 (Figure 5a). *Nocardioides* from the *Actinobacteriota* was also extremely efficient (0.19), as well as the 14 *Bacillus* strains associated with the ASVs 77 and 68 (0.22 and 0.26 respectively). In contrast, for zoospore germination, only strains belonging to the *Pseudomonas* genus were able to efficiently inhibit this *Phytophthora* developmental stage, with an index of 0.27 (Figure 5b).

**Figure 5:**
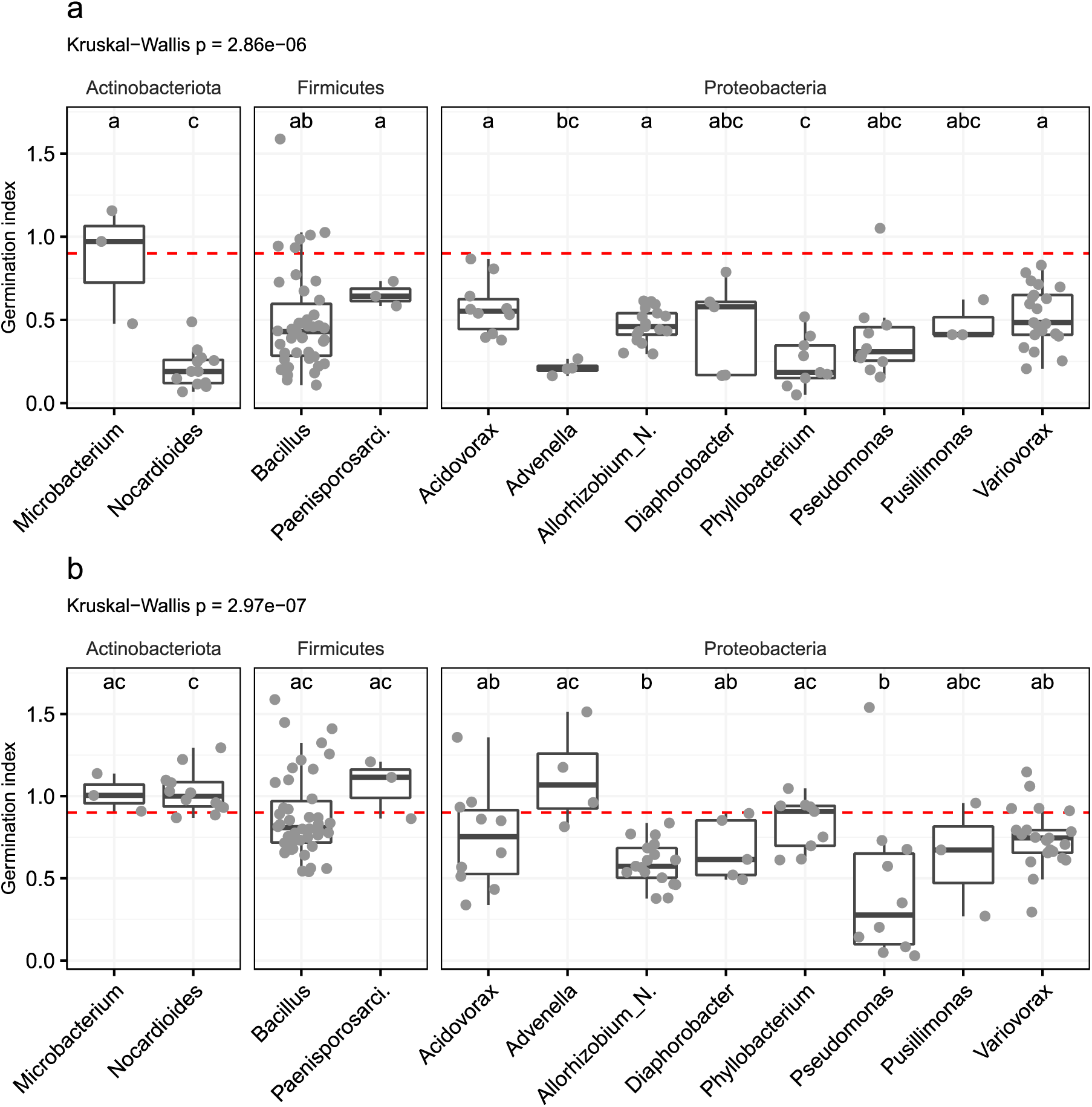
Average final germination indexes per strain for each genus for (a) sporangia germination and (b) zoospore germination. A lower index corresponds to a higher germination inhibition, the red line corresponds to an index of 0.9, below we consider that there is an inhibition. The significance letters are consistent within each sub-figure.

Interestingly, there was an effect of the cultivar of isolation on the strains’ inhibition efficiency of both zoospore and sporangia germination (Figure S14): strains isolated from Innovator exhibited slightly higher inhibitory activity against both zoospore (Dunn’s test p = 0.037) and sporangia (p = 0.044) germination compared to those isolated from Bintje. Additionally, the compartment of isolation significantly influenced zoospore germination inhibition, whereby the strains isolated from the roots were more effective in inhibiting zoospore germination than those isolated from the rhizosphere soil (p = 0.011).

However, we found no overall correlation between the strains’ activity and the differential abundance of the corresponding ASVs in infected vs. non-infected plants, and hence no evidence of a specific plant recruitment mechanism which would have led to enrichment of active strains upon leaf infection (Figure S15). Furthermore, the taxa that most efficiently inhibited spore germination were not associated with ASVs whose abundance strongly differed upon infection in the rhizosphere, except for the *Bacillales* in the first generation.

## Discussion

In this study, we analysed the bacterial community shifts upon late blight infection in two different potato cultivars, Bintje (sensitive) and Innovator (resistant), over two consecutive generations. The overall composition of the microbial communities was similar between the two cultivars, with a few groups slightly more abundant in Bintje (Enterobacterales, *Peptostreptococcales*, *Chitinophagales*, *Burkholderiales* and undefined *Gammaproteobacteria*). This increased abundance was balanced with a higher proportion of *Streptomycetales* in Innovator. These minor differences between the rhizospheres of both cultivars were expected, since plant genotype is considered a secondary driver of the rhizosphere composition behind soil composition, as shown by the comparison of ten common bean cultivars [38]. Another study on seven cultivars of sorghum came to the same conclusion, but found one cultivar with a significantly different rhizosphere composition than the others, and linked this divergence to differences in the strigolactone root secretion [39]. In potato, the comparison of the rhizosphere communities of 51 potato cultivars in one soil revealed that bacterial networks among the cultivars with the highest growth were more developed, but the variations in the rhizosphere composition itself could not be linked to the cultivar’s characteristics [40]. It would have been particularly interesting to characterize the root exudates in our study to determine whether the observed differences were due to variations in the exudation profile between the two genotypes in absence and/or presence of infection.

In our dataset, the main factors influencing the structure of bacterial communities were the plant compartment and the generation, followed by infection status and cultivar. Infection did not affect the alpha diversity in a conclusive way, but significant differences were found in the community structure of the first generation followed by major shifts in the second generation. Other reports about community changes in the rhizosphere infected with various *Phytophthora* species have yielded contrasting results. The diversity of the bacterial rhizosphere of avocado trees did not change if they had symptoms of *P. cinnamomi*-induced root-rot compared to asymptomatic plants, but several phyla including *Actinobacteria* and *Firmicutes* became less abundant while the *Bacteroidetes* became more abundant [41]. In another study, no significant changes were found in the alpha- and beta-diversity of the rhizosphere microbial communities upon root infection of soybean with *P. sojae* [42]. The few studies analysing the impact of the cultivar showed contrasting results: infecting tomato with *P. infestans* led to an increased rhizosphere microbial richness in a susceptible cultivar, but not in the resistant one [43]. In contrast, infecting the roots of common bean with *Fusarium oxysporum* caused increased richness in both sensitive and resistant cultivars [44].

Although we observed that the phyllosphere communities shifted in a cultivar-dependant way upon infection in both generations, only 8 ASVs were differentially abundant in total in this compartment, which is extremely low, especially compared to what has been found after a foliar infection on citrus, where 109 ASVs (20% of the whole community) were promoted in melanose-infected trees [45], or to the extent of the changes observed in rust-infected crabapple, where 17 bacterial orders were differentially abundant compared to healthy trees [46]. A likely explanation for this difference could be that these studies analysed variations in the infected leaves themselves from severely infected plants, while we focused on variations in uninfected leaves from infected plants, as we did not want to compare the bacterial communities of healthy vs. necrotic leaf tissues. Furthermore, the phyllosphere of our greenhouse-grown plants is expected to harbour much lower bacterial richness and abundance compared to plants grown outside due to the importance of environmental factors in the establishment of the phyllosphere communities [47], as reported in lettuce by one study comparing the phyllosphere of greenhouse- and field-grown plants and showing a reduced number of colony forming units by 10- to 100-fold in greenhouse-grown lettuce compared with field-grown ones [48].

Foliar infection caused strong community shifts in the rhizosphere soil in both generations, while causing the bulk soil communities to shift slightly in the first generation and more substantially in the second. These shifts in both soil compartments suggest the existence of a mechanism altering the underground microbiome based on the plant health status, as was already observed in several other plant species infected by various leaf pathogens such as *Arabidopsis thaliana* infected with downy mildew [49], soybean affected by brown spot disease [42], tomatoes infected with late blight [43], and tea trees infected with grey blight [40]. A third of the ASVs of the second generation were differentially abundant in the underground compartments between infected and uninfected plants. These results suggest that the plant triggered rhizosphere shifts upon foliar infection with *P. infestans*. These changes were marginal in the bulk soil of the first generation, but led to the massive shifts observed in the second generation, probably through an amplification of the signal occurring with the repeated infection, keeping in mind that in potted plants, “bulk soil”, although not physically attached to the roots, might still be close enough to the roots to be influenced by the plant. In a setting such as ours, larger differences were observed in community structure in the bulk soil than in the rhizosphere of *A. thaliana* infiltrated over 6 generations with *Pseudomonas syringae*, compared to plants infiltrated with a sterile solution [51]. Meanwhile, the rhizosphere communities stayed under the direct influence of the plant, which led to important changes.

Although both cultivars had relatively similar communities at first, the plant genotype strongly influenced microbial dynamics upon infection, with cultivar-dependent differences in DA ASV counts across both generations. Bintje, the more sensitive cultivar exhibited more than 1700 DA ASV compared to less than 800 in Innovator. The higher responsiveness in Bintje was consistent between generations, especially in the rhizosphere. In the first generation, the DA ASVs in Bintje were almost exclusively inhibited and a large majority of Innovator’s were promoted, while in the second one, most orders had their taxa shifting in the same direction between cultivar, with a majority of promoted ASVs.

The studies that compared rhizosphere shifts in resistant and susceptible cultivars showed different reactions but rarely give details on the nature of the observed shifts. *P. infestans* infection has a stronger impact on the alpha-diversity of the rhizospheric communities of susceptible tomatoes than in the resistant ones [43]. The same study also reported differing beta-diversity shifts between these cultivars, without quantifying them. Another study on common bean infected by the soil-borne pathogen *Fusarium oxysporum* reported rhizosphere shifts in both resistant and susceptible cultivars, with clear family- and genus-level differences in the communities of these cultivars [44].

Interestingly, the rhizosphere of susceptible rapeseed infected with *Plasmodiophora brassicaceae* became a lot more interconnected upon infection, whereas resistant cultivars only showed minor changes [52]. It must be emphasized that unlike in our study, the shifts observed in studies using root infections could be caused by a direct action of the pathogen, not only by a root exudate-mediated reaction of the plant.

The taxa that reacted the most to infection were spread across several phyla: *Firmicutes*, *Proteobacteria*, *Bacteroidota* and *Acidobacteriota*. Most of these DA orders and genera are known to host PGPR.

In the *Firmicutes*, the abundance of the *Bacillales* collapsed upon infection during the first generation in both cultivars, as previously observed in avocado [41], but still increased upon infection. This order is known for the extensively studied *Bacillus* genus, which contains several plant-beneficial species. Some were shown to promote plant growth by secreting phytohormones in the rhizosphere [53] or by increasing nutrient availability by nitrogen fixation [54, 55] and phosphorus solubilization [56, 57]. Furthermore, rhizospheric *Bacillus* were reported to help the plant against pathogens, firstly by triggering ISR, allowing tomatoes to resist viral [58] and fungal [59] infections, secondly by secreting a wide range of powerful pathogen inhibitors, which were able to protect tomatoes from bacterial canker [60] and wheat against take-all [61]. The *Clostridiales* and the *Peptostreptococcales* on the other hand, were much less abundant than the *Bacillales* but were heavily promoted in Bintje following infection in the second generation. While we know that the *Clostridiales* can interact with plants, either as sweet potato pathogens [62] or as a nitrogen-fixing endophytes in grasses [63], the *Peptostreptococcales* were never reported to interact with plants to our knowledge.

The *Burkholderiales* was the most abundant order of the *Proteobacteria*. They were slightly promoted upon infection in the rhizosphere of Innovator during the first generation, and strongly promoted in the rhizosphere of both cultivars during the second generation. Among them, the *Comamonadaceae* had the highest proportion of DA. It is a large family, which contains many cultivable species, including many plant growth promoting strains as well as phytopathogens. Among them, members of the *Variovorax* genus were reported to improve barley and fenugreek plant growth in agricultural soil when inoculated with a nitrogen-fixing *Sinorhizobium* [64]. Members of the *Comamonas* were reported to be great nitrogen fixers [65], and some *Acidovorax* strains improved barley seedlings survival and promoted root growth [66], while others are known cucurbit pathogens [67].

The *Flavobacteriales* were the most prominent and reactive order of the *Bacteroidota*. They were promoted in the resistant cultivar during the first generation, but were strongly inhibited in both cultivars in the second generation. In our study, they were essentially from the *Flavobacterium* genus, which is abundant in the rhizosphere of several species including cucumber [68], lettuce [69], maize [70] and *A. thaliana* [18], and both the rhizosphere and the phyllosphere of wheat [71]. Several isolates could promote plant growth by secreting phytohormones in the rhizosphere [72–74], while a strain is capable of protecting capsicum plants resist *Phytophthora capsicii* root infection [75]. Besides, the *Saprospiraceae* (most of the *Chitinophagales* reads) were heavily promoted in the rhizosphere of Bintje during the second generation. Although they have been observed in the rhizosphere of maize [76] and soils fertilized with manure [77], they have been detected much more frequently in aquatic environments and bioreactors [78].

Lastly in the *Acidobacteriota*, infection led to a considerable promotion of *Bryobacter* genus during the second generation in both cultivars. Several elements point toward a positive influence of this group on plant survival as they were detected in the rhizosphere of plants from environments as inhospitable as the Atacama desert and Antartica [79]. Furthermore their presence was reported to promote duckweed growth [80] and was associated with healthy sesame [81] and *Polygonatum kingianum* [82] plants. In addition, metagenomic studies find many plant growth promoting traits in assembled genomes of members of this genus [83].

When we assessed the biological activity of the isolated strains on *P. infestans* sporangia and zoospore germination, we observed that the genera *Advenella*, *Nocardioides* and *Phyllobacterium* were the best at inhibiting sporangia germination. *Advenella* contains known Plant Growth Promoting Bacteria (PGPB), capable of alleviating salinity stress [84] and reported to be a necessary element of a simple synthetic community capable of reducing root rot incidence by triggering Induced Systemic Resistance [85]. To our knowledge, phytopathogen inhibition activity has only been reported once in this genus [86]. *Nocardioides* are usually found in the soil but are also connected to plants, as some species contain endophytic strains [87, 88]. They have received some attention lately and are extensively studied for their bioremediation potential but are not usually linked to the inhibition of phytopathogenic organisms [89]. Several *Phyllobacterium* species are known for helping plant growth and resilience in several ways. Their presence in the rhizosphere increased resistance to drought stress [90, 91], they increased lateral root growth [92] and solubilized phosphate [91]. Broad-spectrum fungal inhibition was reported once [93], but anti-oomycete activity has never yet been reported to our knowledge.

Inhibiting zoospore germination seemed more challenging than inhibiting sporangia germination, as only *Pseudomonas* strains were able to do so. This genus is very diverse, and several species are extensively studied for their role as plant growth promoters and in plant protection [94]. Interestingly, the strains isolated from the resistant cultivar Innovator were in average more efficient in inhibiting both spores than the strains isolated from the sensitive cultivar Bintje, suggesting that the resistant cultivar’s microbiota might be enriched in protective strains, which might contribute to their host’s higher disease resistance. Despite these findings, we could not link the differential abundance of the ASVs to the direct inhibition of *P. infestans* spores by the ASV-associated strains. This conclusion is constrained by the limited number of DA ASVs that could be associated to cultivable strains from our collection. Several taxa with strong shifts were not represented at all in our collection, such as the *Flavobacterium* and *Saprospiraceae* among the *Chitinophagales*, *Bryobacter* among the *Acidobacteria*, and obligate anaerobic taxa as *Clostridiales* and *Peptostreptococcales*. Furthermore, our method focused on *in vitro* direct inhibition, which may not represent *in planta* protection [95]. Additionally, we cannot exclude that protective mechanisms might rely on interactions between multiple organisms, which we would not see in our experimental setup. For example, *A. thaliana* was protected from downy mildew by a consortium of three rhizosphere-isolated strains that triggered induced systemic resistance [49], and *Fusarium* wilt suppression in a field resulted from the synergistic action of a siderophore-producing *Pseudomonas* and a non-pathogenic *Fusarium* [96, 97]. Deciphering the role of the strains associated to DA ASVs, as single strains or in combinations, will hence require further experiments in ulterior studies integrating the plant as well.

## Conclusion

Foliar infection with *Phytophthora infestans* induced shifts in the potato-associated microbiome, which were first detected in the rhizosphere of the first generation and led to a community divergence with much wider differences in the bulk soil and rhizosphere in the second generation, while the phyllosphere remained stable. Plant genotype also played a major role, as shifts were consistently more important in the susceptible cultivar Bintje that in the resistant cultivar Innovator. While the shifts in the first generation in Bintje were almost exclusively from inhibited ASVs in the *Bacillales*, Innovator’s differentially abundant ASV were primarily promoted in the *Burkholderiales* and *Flavobacteriales*. The shifts observed in the second generation were more similar between cultivars with common non-clustered shifts among the *Burkholderiales* and the *Flavobacteriales*, alongside cultivar-specific clustered shifts in *Chitinophagales*, *Clostridiales*, *Xanthomonadales*, *Bryobacterales*, *Paludibaculum* and *Peptostreptococcales*. Biological assays to assess the spore inhibition potential of strains isolated from potato plants revealed the biocontrol potential of strains otherwise not studied for their antagonism towards phytopathogenic organisms, such as *Advenella*, *Nocardioides* and *Phyllobacterium*. Although we did not see any correlation between differential abundance and biological activity, the overall higher activity of strains isolated from the resistant potato cultivar still suggests that the plant microbiota could play a significant role in its host’s resistance to diseases. It would therefore be of interest to test, in future experiments, whether these active strains associated with the resistant cultivar could be used to improve the protection of the sensitive.

## List of Abbreviations

ASV: Amplicon sequence variant
mBCAs: Microbial Biological Control Agents
DA: Differentially abundant
dbRDA: Distance-based redundancy analysis
GLM: Generalised linear model
GLS: Generalised least squares model
HSD: Tuckey’s honest significance differences
NPS: Non-challenged plant soil
OD600: Optical density at 600nm
PCoA: Principal Coordinate Analysis
PGPR: Plant growth promoting rhizobacterium
PPS: Phytophthora-challenged plant soil
TMM: Trimmed Mean of the M component
TSS: Total sum scaling

## Declarations

### Ethics approval and consent to participate

Not applicable.

### Consent for publication

Not applicable.

### Availability of data and materials

The Miseq sequencing data and the 16S rRNA sequences of the isolated strains are available in the European Nucleotide Archive, accession PRJEB75732 at the following URL: https://www.ebi.ac.uk/ena/browser/view/PRJEB75732.

The scripts used during the current study are available in the PotatoCryForHelp repository on Github, available at this URL: https://github.com/VivienPichon/PotatoCryForHelp.

The intermediate data and figures generated by the scripts are publicly available on Zenodo, at this URL:

https://zenodo.org/records/13384779?preview=1&token=eyJhbGciOiJIUzUxMiJ9.eyJpZCI6IjQ3OWMwMzg1LWE1MDAtNGNkMS04N2RhLTgxZWQxYTg0NDY5MCIsImRhdGEiOnt9LCJyYW5kb20iOiJmMzQyYTAyMjZiMmI2YWI2MDM3Zjg3YzI2NDMxZTQxYSJ9.DLnhZkjhpd7Isb4Dl7inUIFAYP6r0w7jKhCLr13Z3DUr4yU6KcFBgdL0uoO8Y3S_WkjaorkOHjzF3f_Dy_pyMg.

### Author’s contribution

Vivien Pichon did the infection experiment, managed the bacterial strain isolation, the data analysis and the redaction of the article. Mout De Vrieze supervised the lab experiments, and directed the confrontation assays with Rares Cristea. Farès Bellameche participated in isolating microbial strains and reviewing the article. Floriane L’Haridon managed the laboratory and provided technical support for the project. Laurent Falquet supervised the microbiome analysis part of the project. Laure Weisskopf acquired the funding, designed the project and supervised it, except for the microbiome analysis part, and contributed to writing and revising the article. All authors read and approved the final manuscript.

## Acknowledgements

We are grateful for the technical assistance of Aurélie Esseiva, Elissa Feghali, Camila Morales, Eva Trutman, Nicolas Rappo and Tom Lüthi. We would also like to thank the Next Generation Sequencing Platform in Bern for their precious support, Tobias Gelencsér (Research Institute of Organic Agriculture, FiBL) for helping us find the soil inoculum, and Natacha Bodenhausen (FiBL) for her valuable inputs on the project. We are also grateful to the team of the Botanical Garden of Fribourg for their continued help in the greenhouse.

## Supplementary

**Table S1:** Average richness and evenness of bacterial communities in function of compartment, generation, treatment and cultivar. The significance was calculated with a 4-way ANOVA for the soil and rhizosphere samples pooled, a 3-way ANOVA for the soil and rhizosphere samples separated, and with Generalised Least Squares (GLS) for the phyllosphere. The groups written in italics represent the crossed effect of two factors.

**Table S2:** Tables of the differential abundance of all taxa per cultivar in the rhizosphere of the first generation. Descriptive statistics were calculated for every taxa of every taxonomic level present in our study based on the TMM-normalized abundance and were expressed in percentages. The last letter of the column names (B, I) refers to the cultivar. The Rel_abund columns show the abundance of each taxon as a proportion of the whole abundance. The Weighted_dens columns show the proportion of the abundance of the DA ASV of each taxon as a proportion of the total abundance. The More columns represent the proportion of the abundance of the ASV which become significantly more abundant upon infection as a proportion of the abundance of the taxon. The Less columns are the same as the More columns, but for significantly less abundant ASV.

**Table S3:** Tables of the differential abundance of all taxa per cultivar in the rhizosphere of the second generation. Descriptive statistics were calculated for every taxa of every taxonomic level present in our study based on the TMM-normalized abundance and were expressed in percentages. The last letter of the column names (B, I) refers to the cultivar. The Rel_abund columns show the abundance of each taxon as a proportion of the whole abundance. The Weighted_dens columns show the proportion of the abundance of the DA ASV of each taxon as a proportion of the total abundance. The More columns represent the proportion of the abundance of the ASV which become significantly more abundant upon infection as a proportion of the abundance of the taxon. The Less columns are the same as the More columns, but for significantly less abundant ASV.

**Table S4:** Overview of spore inhibition scores and other characteristics of the tested isolates. For each strain, the germination score (SG for the sporangia and ZG for the zoospores), the cultivar and compartment of isolation, and the differential abundance status of the corresponding ASVs observed per experimental factor is given. A germination score below 0.25 indicates strong inhibition, a score of 0.25 - 0.5 indicates moderate inhibition, 0.5 - 0.9 indicates weak inhibition, 0.9 −1.8 indicates no inhibition, and a score higher than 1.8 indicates that germination was stimulated. The differential abundance status (columns I to O) is based on the differential abundance assessment visible on the subtables, defined by a group of samples which share all experimental factors (e.g. all replicates from the rhizosphere of Bintje during the first generation). Differential abundance between uninfected and infected samples within a subtable was assessed with a Generalized Linear Model, and the results were then grouped per experimental factor (for example the phyllosphere group is a summary of the 4 sample subtables from the phyllosphere, encompassing both generations and cultivars). ASV were categorized as follows: ABSENT means that the ASV was observed in no subtable, AMBIG means that it became more abundant in at least one subtable and less abundant in at least one other, EQUAL means that the ASV was observed but did not differ in abundance, MORE means that the ASV became more abundant in at least one subtable, LESS means that it became less abundant in a least one subtable.

**Figure S1:** Rarefaction curves of the samples analysed in this study. The samples were grouped as follows: (a) samples from the soil and rhizosphere of the first generation, (b) samples from the soil and rhizosphere of the second generation, (c) samples from the phyllosphere of both generations. For the soil and rhizosphere plots, the line colour represents the compartment; for the phyllosphere plot the shade represents the generation.

**Figure S2:** Composite boxplot representing the difference of richness and evenness between the samples of infected and uninfected plants for each compartment, generation and cultivar. The colour represents the compartment, the shade represents the plant infection status, the annotation above each boxplot shows the significance. (ns for p > 0.05, * for 0.05 ≥ p > 0.01, ** for 0.01 ≥ p > 0.001).

**Figure S3:** Distance-based redundancy analysis of the soil and rhizosphere communities, partialling out a) the compartments and b) the generations.

**Figure S4:** Comparison of the proportion of the phyla between generations in the soil, the rhizosphere and the phyllosphere.

**Figure S5:** Comparison of the proportion of the phyla between infected and uninfected plants in (a) the phyllosphere, (b) the rhizosphere and (c) the soil for each generation and cultivar.

**Figures S6 to S13:** Detailed phylogenetic trees representing the log-fold change (inner circle) and the abundance (outer circle) per cultivar of every ASV. Each tip represents one ASV, the colours represent the ASV family or order. Each annotation circle is split in two, the innermost half representing the Bintje cultivar and the outermost half representing Innovator. The outer circle represents the abundance, the inner circle represents the log-fold change of the significantly differentially abundant ASV. Grey in the inner circle means no significant differential abundance.

**Figure S14:** Average final germination score of each strain for (a) the sporangia and (b) the zoospores per compartment (top) and cultivar of isolation (bottom).

**Figure S15:** Average final germination score of each ASV categorized by differential abundance for (a) the sporangia and (b) the zoospores.

