## Supplementary figures and images for "Potato foliar infection with *Phytophthora infestans* drives strong, cultivar-specific shifts in rhizosphere communities"

### Supplemental Figure S1

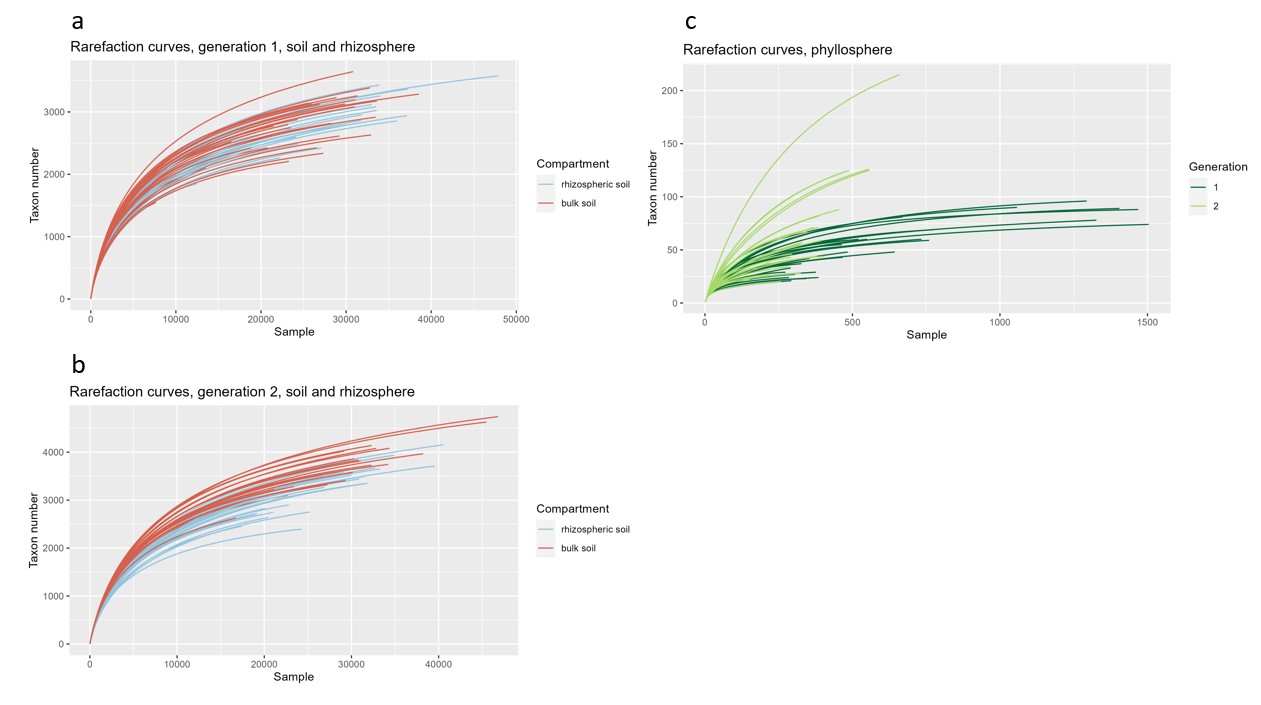

### Supplemental Figure S2

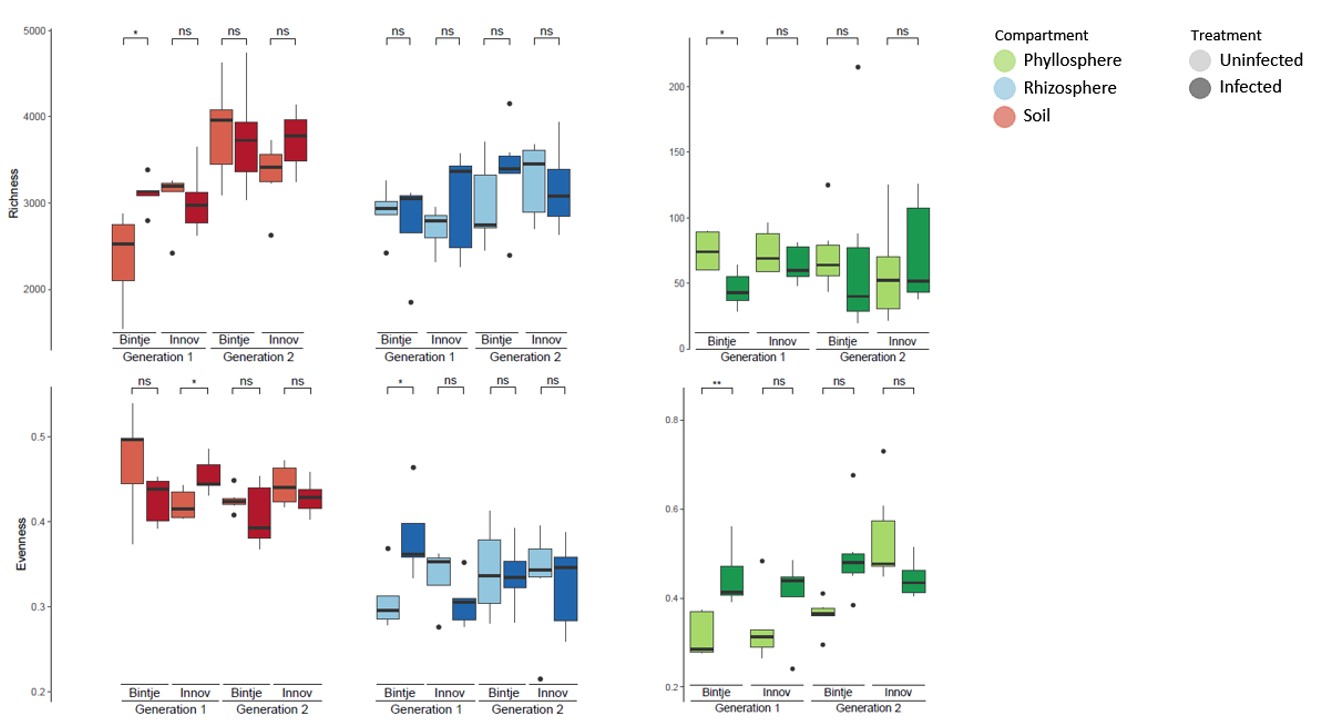

### Supplemental Figure S3

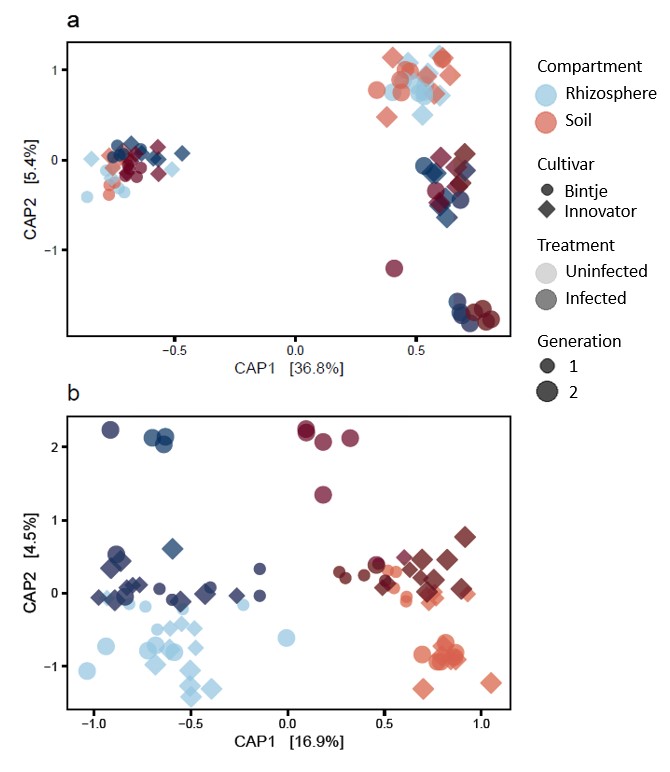

### Supplemental Figure S4

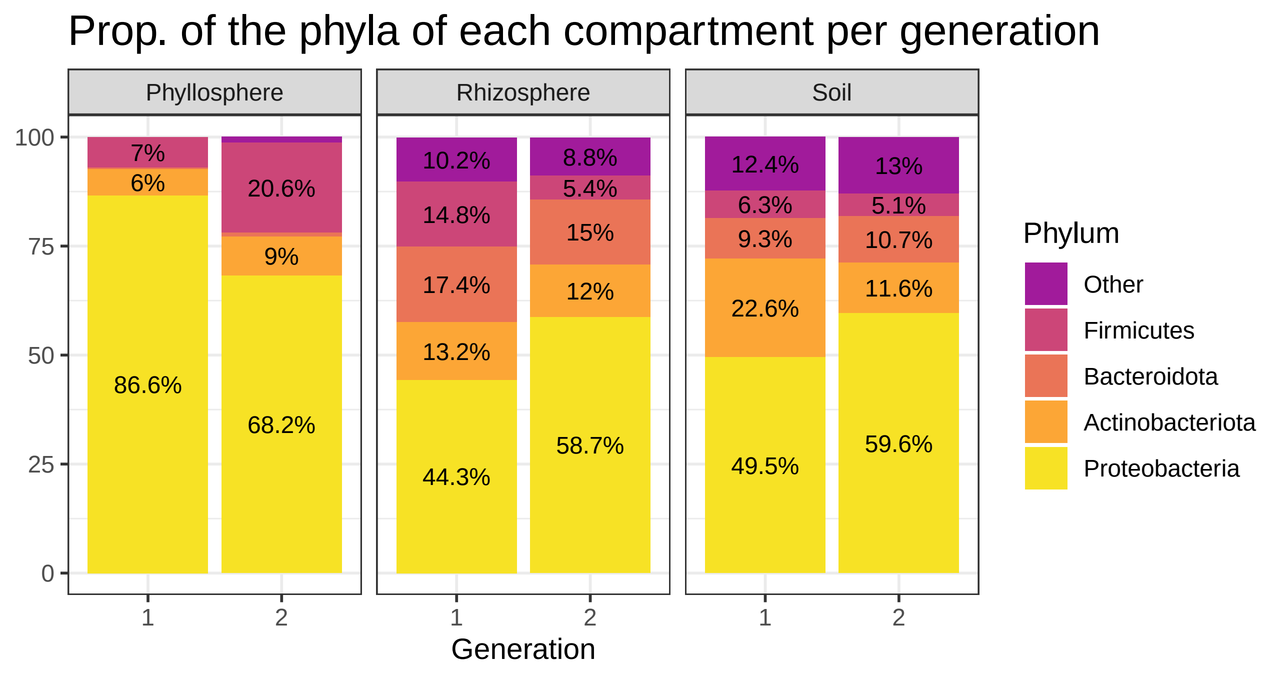

### Supplemental Figure S5

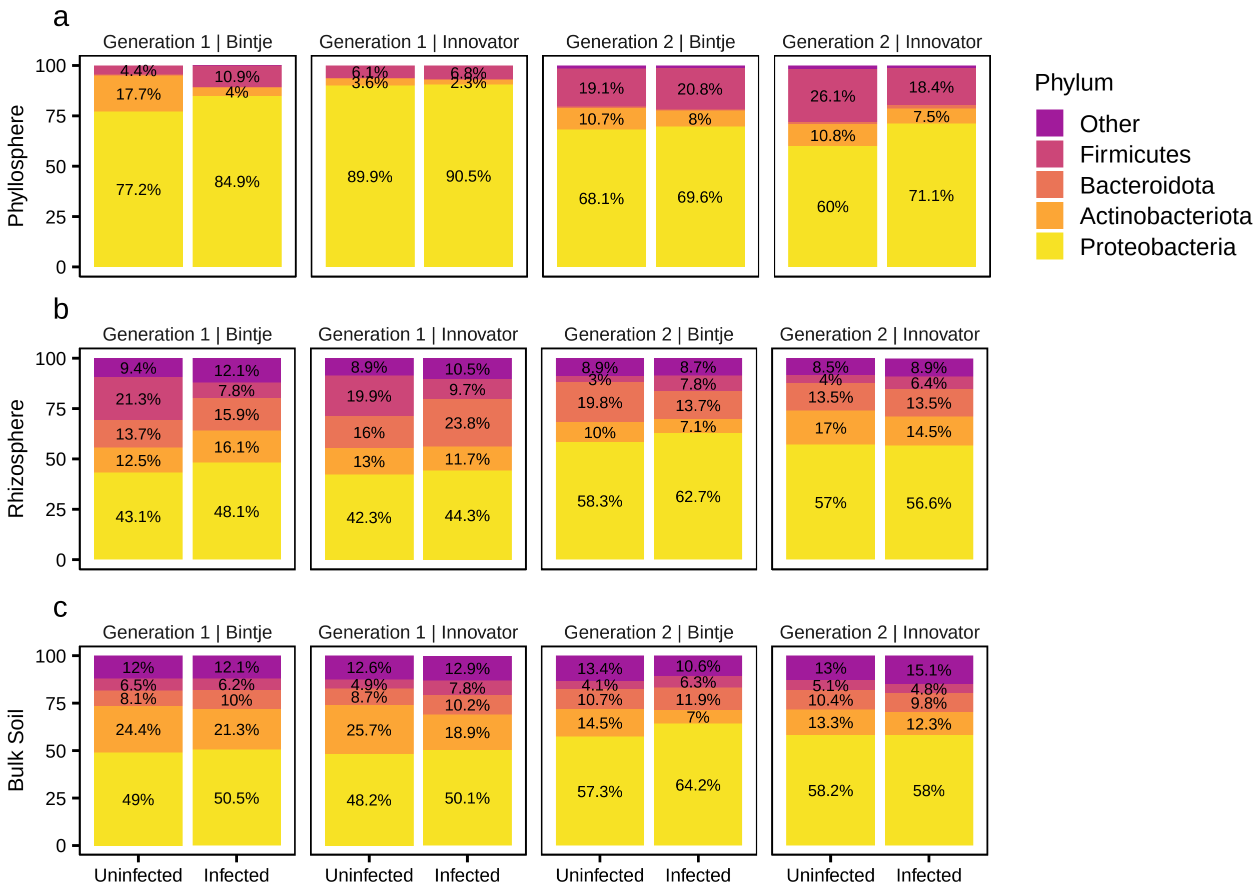

### Supplemental Figure S6

#### Subtree firmicutes, Rhizosphere, p < 0.05

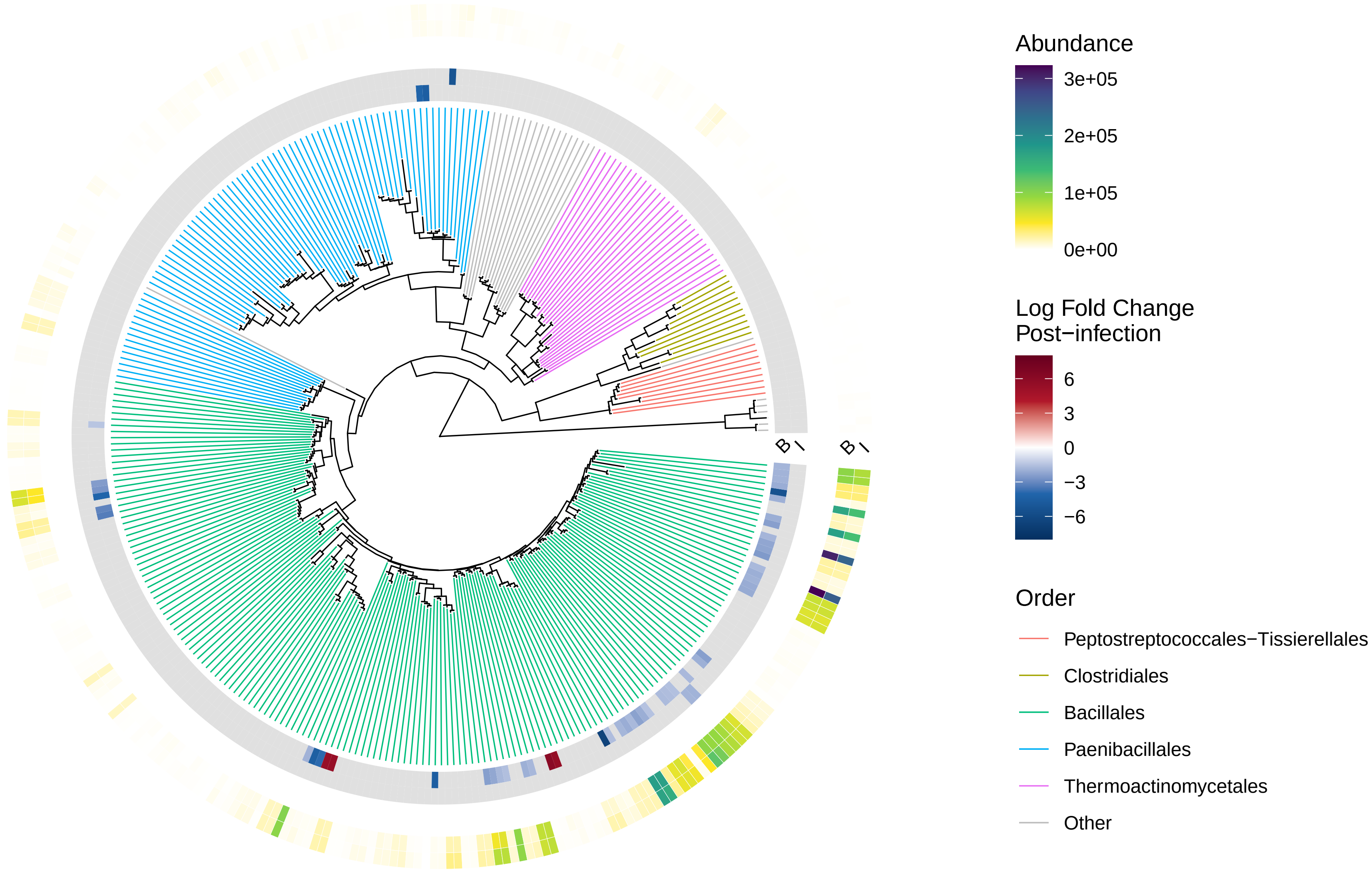

### Supplemental Figure S7

Subtree burkholderiales, Rhizosphere, $p < 0.05$

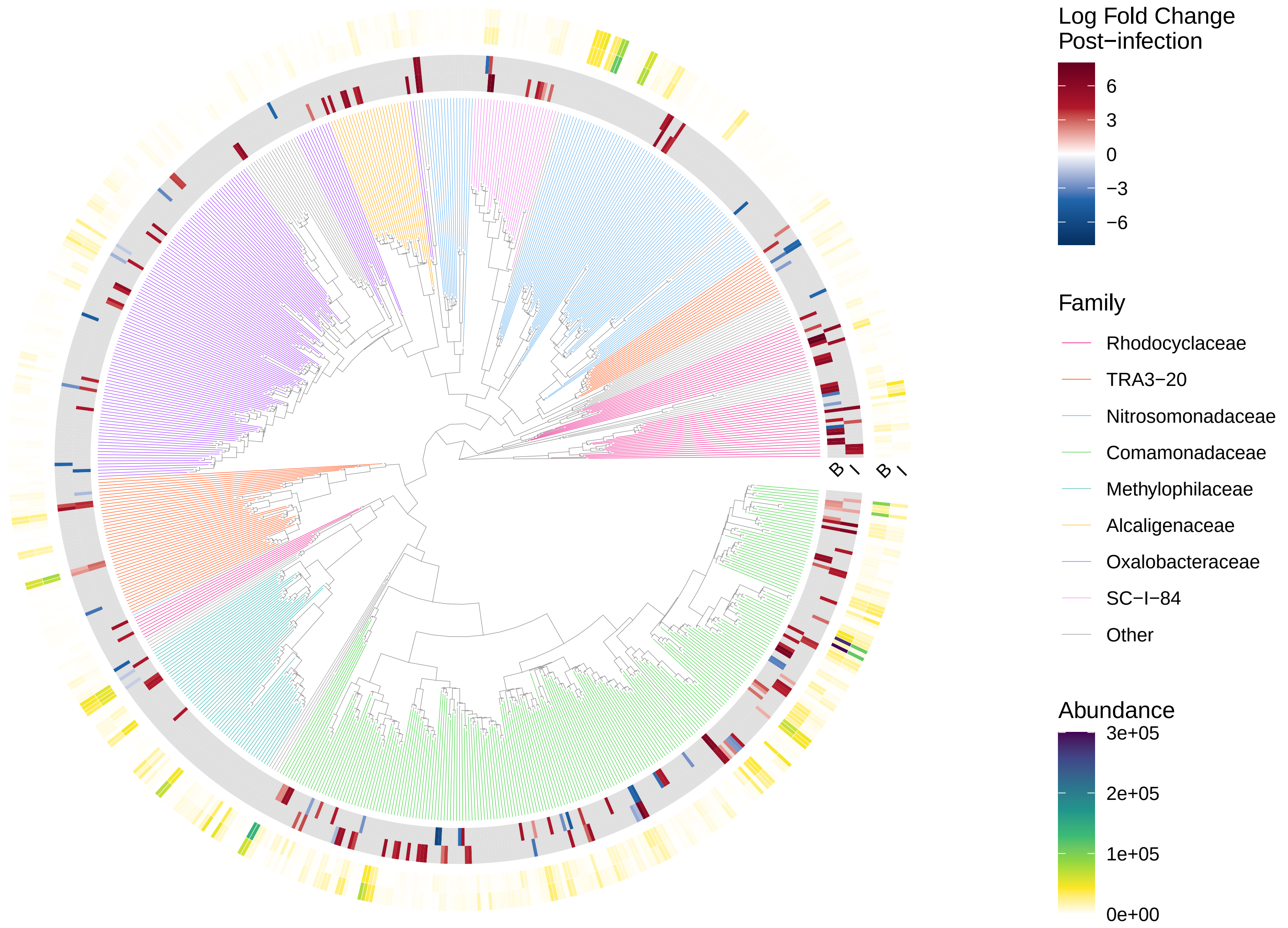

### Supplemental Figure S8

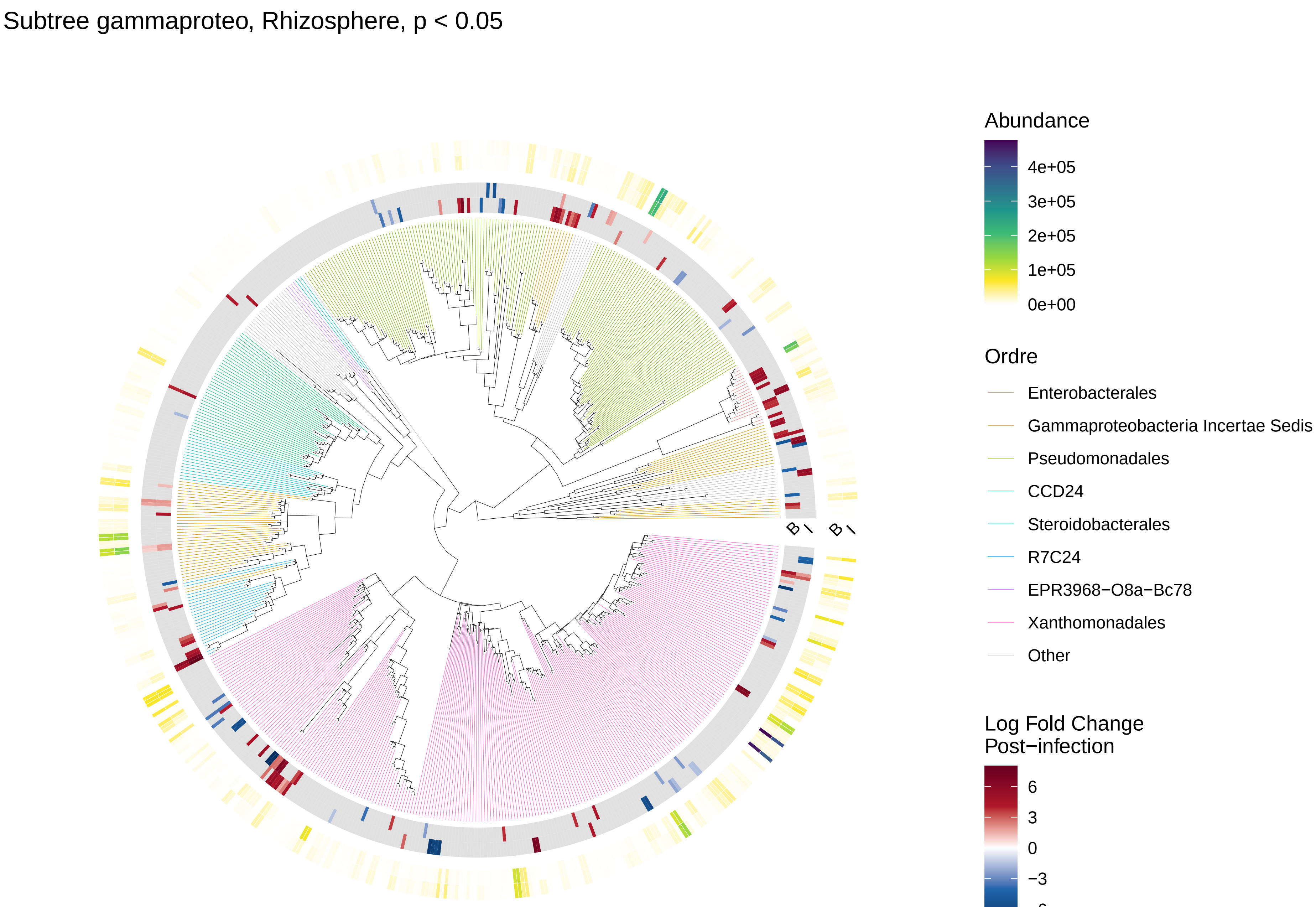

### Supplemental Figure S9

Subtree bacteroidota, Rhizosphere, $p < 0.05$

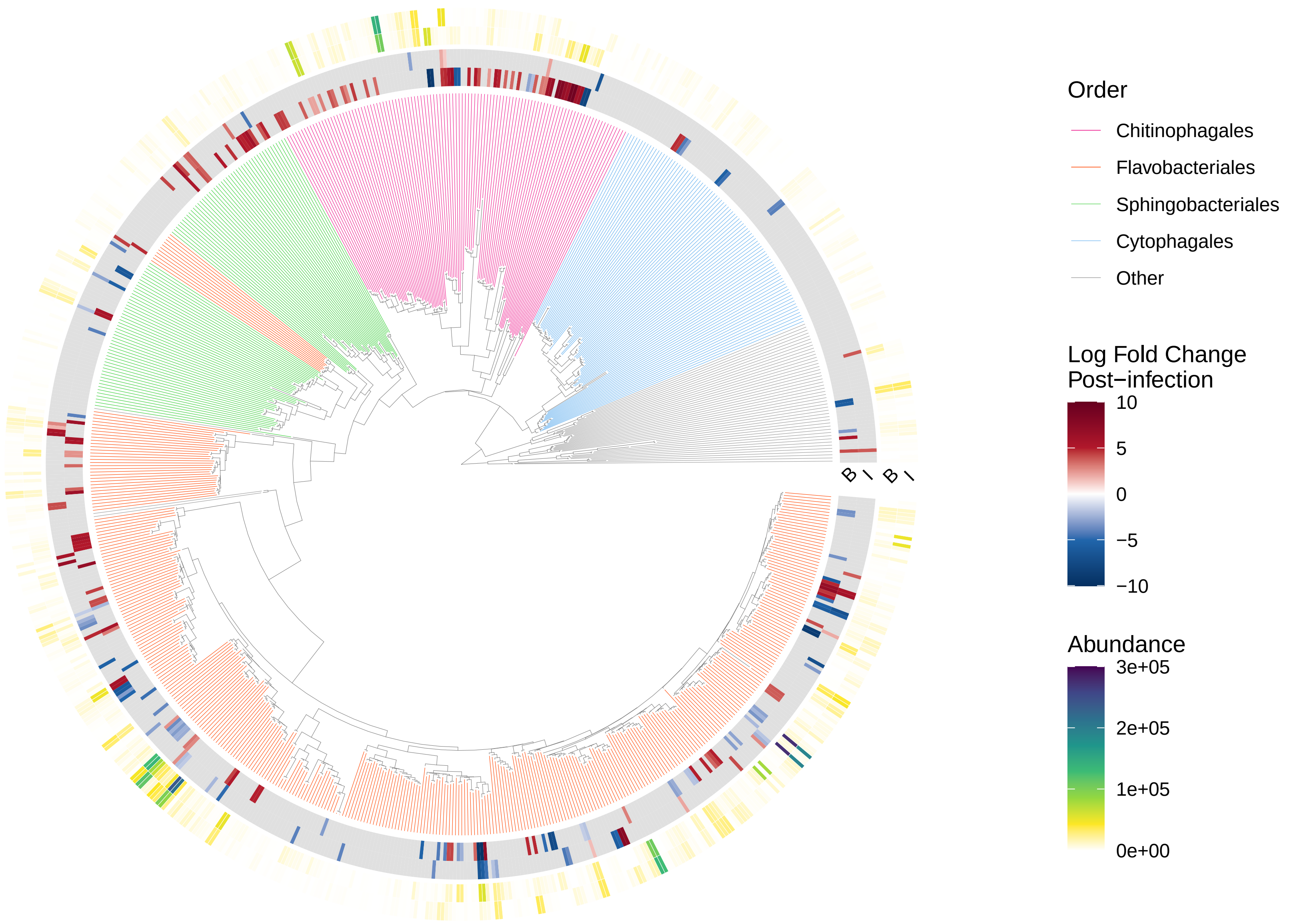

### Supplemental Figure S10

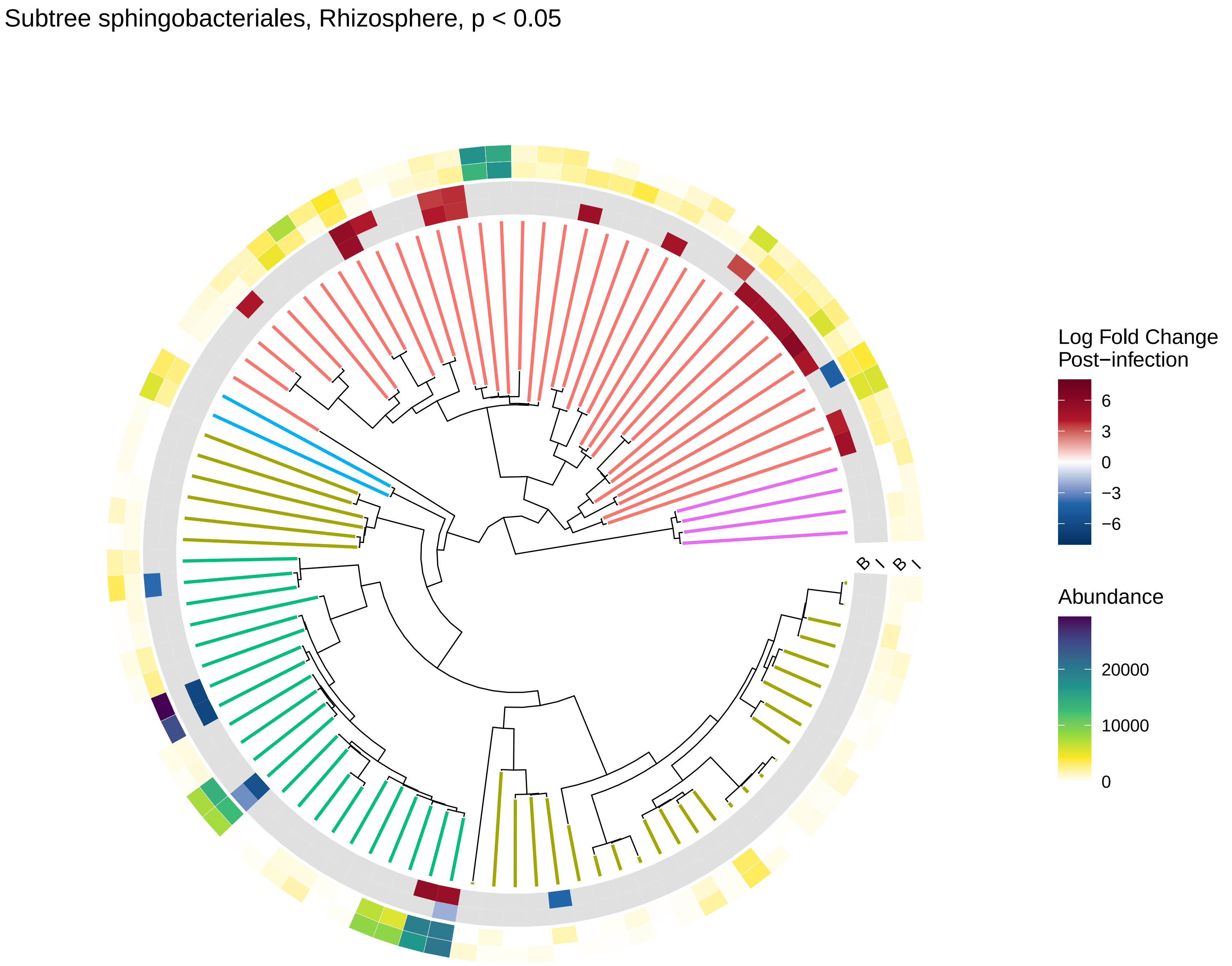

### Supplemental Figure S11

Subtree firmicutes, Rhizosphere, $p < 0.05$

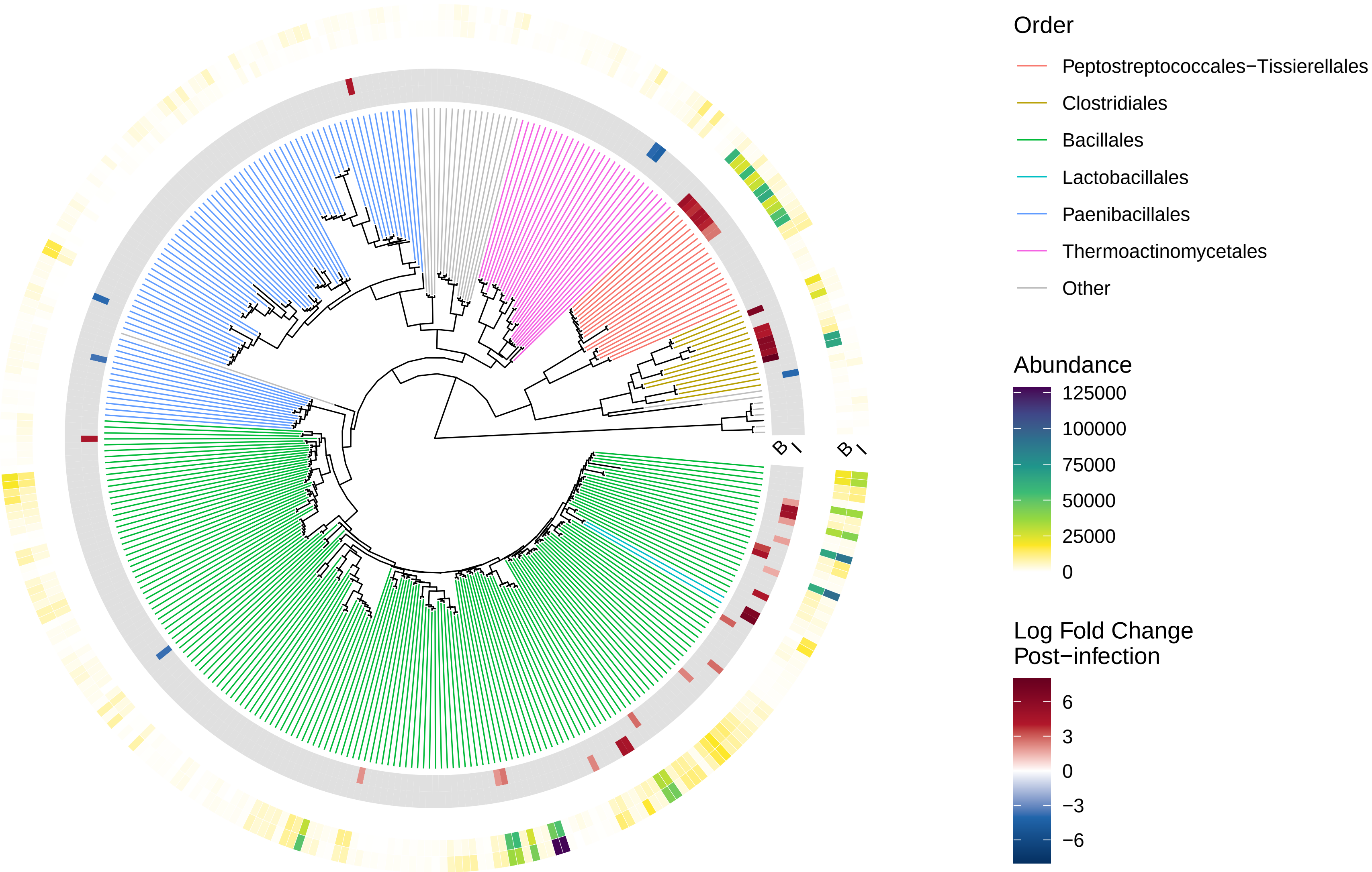

### Supplemental Figure S12

Subtree actinobacteriota, Rhizosphere, $p < 0.05$

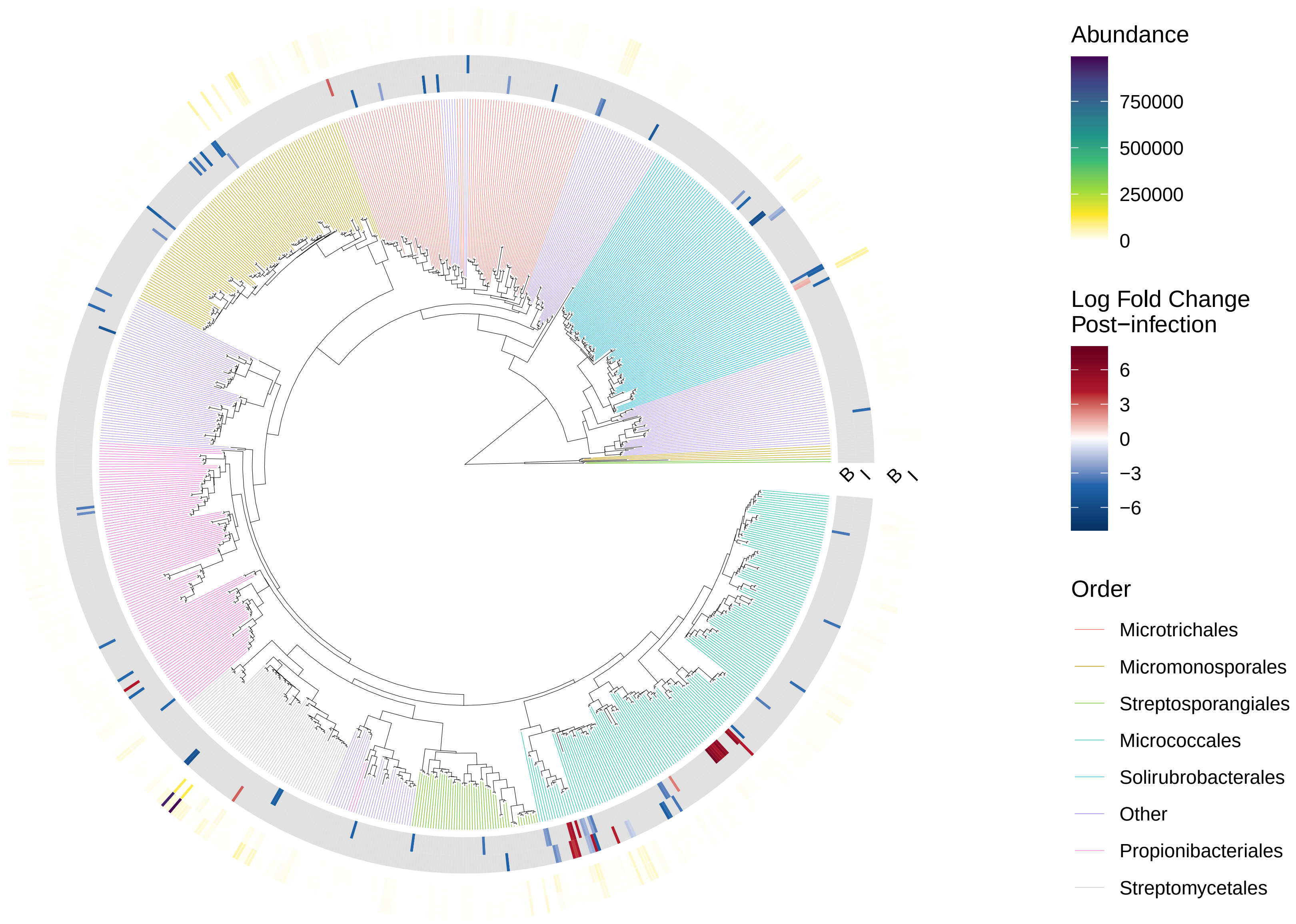

### Supplemental Figure S13

##### Subtree diverse1, Rhizosphere, $p < 0.05$

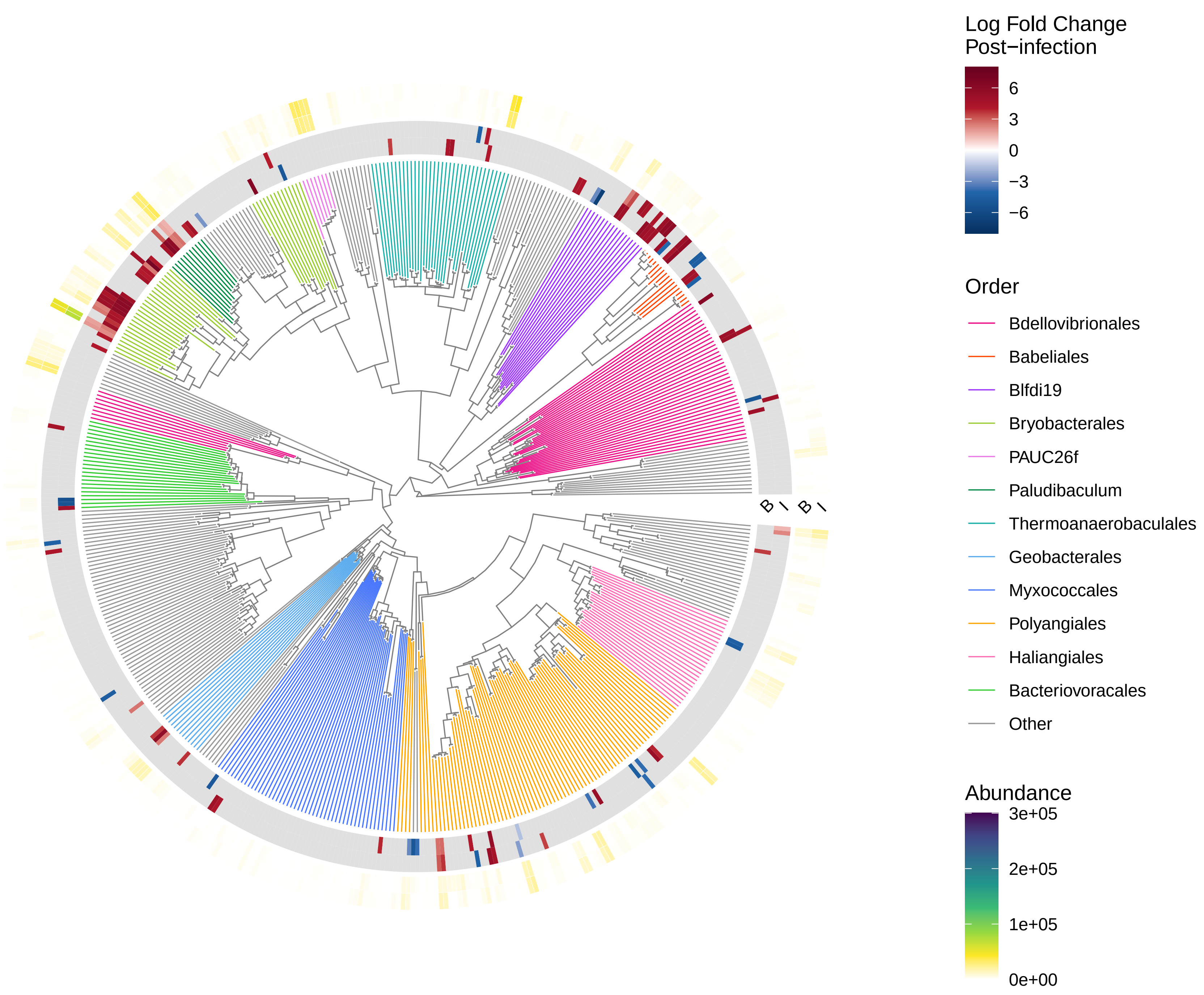

### Supplemental Figure S14

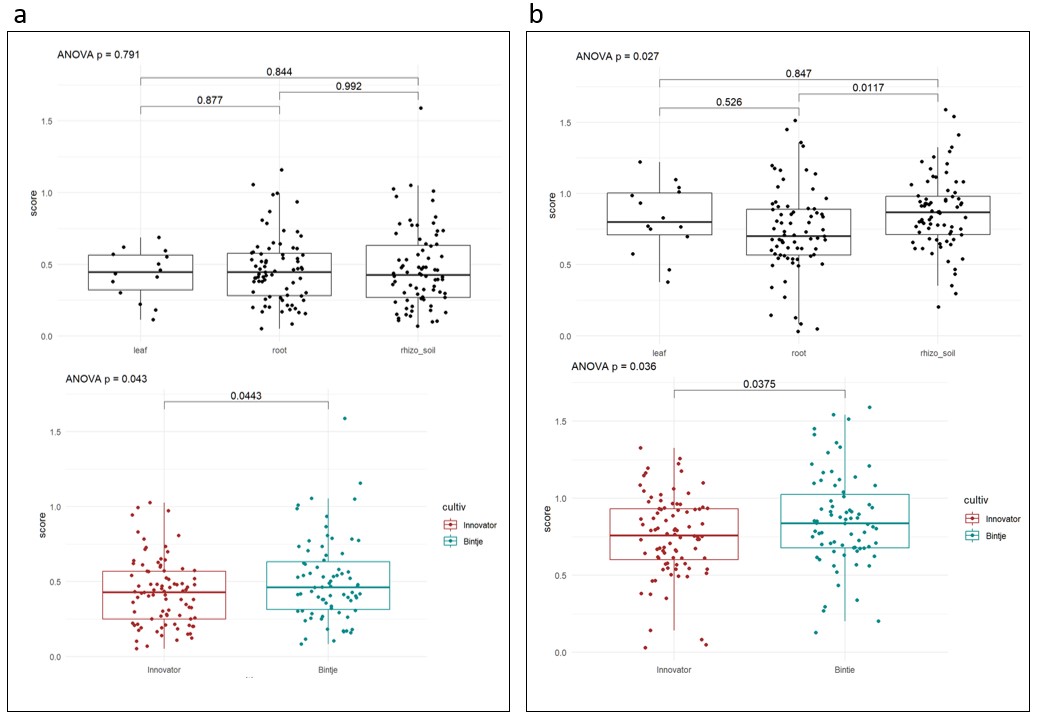

### Supplemental Figure S15

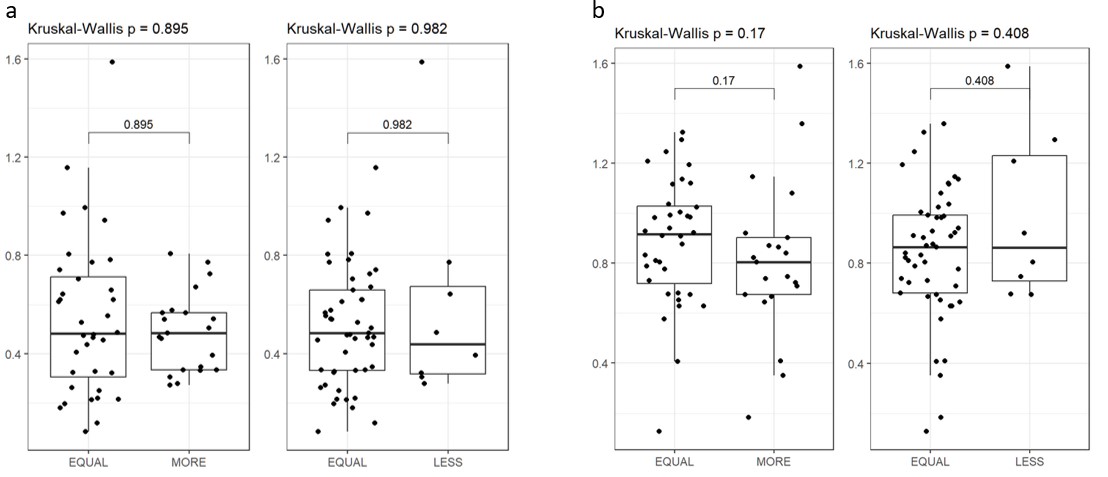
