## Supplemental Table S1 for "Potato foliar infection with *Phytophthora infestans* drives strong, cultivar-specific shifts in rhizosphere communities"

| Compartment | Group | Avg richness | p-value | Avg evenness | p-value |
|---|---|---|---|---|---|
| Soil and rhizosphere pooled | Soil | 3301 | | 0.43 | |
| | Rhizosphere | 3031 | 8.7E-03 | 0.33 | <2E-16 |
| | gen1 | 2860 | | 0.39 | |
| | gen2 | 3420 | 3.3E-07 | 0.38 | 0.42 |
| | Infected | 3259 | | 0.38 | |
| | Non-infected | 3073 | 0.07 | 0.38 | 0.81 |
| | Bintje | 3157 | | 0.39 | |
| | Innovator | 3174 | 0.86 | 0.38 | 0.55 |
| | *Generation x Compartment* | | 0.04 | | 0.30 |
| | *Treatment x Cultivar* | | 0.71 | | 0.78 |
| Soil | gen1 | 2880 | | 0.44 | |
| | gen2 | 3651 | 4.3E-06 | 0.43 | 0.11 |
| | Infected | 3427 | | 0.43 | |
| | Non-infected | 3174 | 0.09 | 0.44 | 0.27 |
| | Bintje | 3302 | | 0.43 | |
| | Innovator | 3299 | 0.98 | 0.44 | 0.59 |
| | *Treatment x Cultivar* | | 0.78 | | 0.06 |
| Rhizosphere | gen1 | 2840 | | 0.33 | |
| | gen2 | 3190 | 1.9E-02 | 0.34 | 0.89 |
| | Infected | 3091 | | 0.34 | |
| | Non-infected | 2971 | 0.4 | 0.33 | 0.63 |
| | Bintje | 3012 | | 0.34 | |
| | Innovator | 3050 | 0.79 | 0.33 | 0.27 |
| | *Treatment x Cultivar* | | 0.82 | | 0.10 |
| Phyllosphere | All | 66.8 | | 0.42 | |
| | gen1 | 64.7 | | 0.38 | |
| | gen2 | 68.6 | 0.28 | 0.46 | 1.2E-03 |
| | Infected | 64.3 | | 0.4 | |
| | Non-infected | 69.4 | 0.15 | 0.39 | 6.7E-03 |
| | Bintje | 66.5 | | 0.41 | |
| | Innovator | 67.1 | 0.72 | 0.44 | 0.36 |
| | *Generation x Treatment* | | 0.62 | | 0.76 |
| | *Treatment x Cultivar* | | 0.41 | | 0.29 |
