## Supplemental Table S2 for "Potato foliar infection with *Phytophthora infestans* drives strong, cultivar-specific shifts in rhizosphere communities"

| group | rank | abund_B | abund_I | rel_abund_ | rel_abund_ | B_more | I_more | B_less | I_less | weighted_d | weighted_densI |
|---|---|---|---|---|---|---|---|---|---|---|---|
| Bacteria | Kingdom | 26612942 | 27479020 | 100 | 100 | 0.1 | 1.4 | 12.1 | 0.6 | 12.22 | 1.97 |
| Proteobac | Phylum | 12078057 | 11909217 | 45.4 | 43.3 | 0.1 | 2.2 | 0.6 | 0.1 | 0.32 | 0.98 |
| Gammap | Class | 6039993 | 6456824 | 22.7 | 23.5 | 0.1 | 3.8 | 1 | 0.1 | 0.24 | 0.91 |
| Burkhold | Order | 2470835 | 2101143 | 9.3 | 7.6 | 0.1 | 11.7 | 2.3 | 0.2 | 0.22 | 0.91 |
| Comam | Family | 978616 | 623754.1 | 3.7 | 2.3 | 0 | 39.3 | 2.3 | 0.8 | 0.08 | 0.91 |
| Comar | Genus | 515320.6 | 330019.9 | 1.9 | 1.2 | 0 | 74.3 | 4.4 | 1.5 | 0.08 | 0.91 |
| Rhizob | Genus | 221884.3 | 28194.4 | 0.8 | 0.1 | 0 | 0 | 0 | 0 | 0 | 0 |
| Variov | Genus | 51010.3 | 38590.8 | 0.2 | 0.1 | 0 | 0 | 0 | 0 | 0 | 0 |
| Hydrog | Genus | 43822.4 | 89965.3 | 0.2 | 0.3 | 0 | 0 | 0 | 0 | 0 | 0 |
| Acidov | Genus | 33993.4 | 20949.3 | 0.1 | 0.1 | 0 | 0 | 0 | 0 | 0 | 0 |
| Ramlib | Genus | 33763.3 | 34799.8 | 0.1 | 0.1 | 0 | 0 | 0 | 0 | 0 | 0 |
| Piscini | Genus | 20070.9 | 20587.9 | 0.1 | 0.1 | 0 | 0 | 0 | 0 | 0 | 0 |
| Pelom | Genus | 12859.7 | 20604.7 | 0 | 0.1 | 0 | 0 | 0 | 0 | 0 | 0 |
| Caenir | Genus | 11770.6 | 9496.4 | 0 | 0 | 0 | 0 | 0 | 0 | 0 | 0 |
| Methyl | Genus | 7992.1 | 2478 | 0 | 0 | 0 | 0 | 0 | 0 | 0 | 0 |
| Hylem | Genus | 4516.2 | 7196.5 | 0 | 0 | 0 | 0 | 0 | 0 | 0 | 0 |
| Limnol | Genus | 4205.8 | 4601.4 | 0 | 0 | 0 | 0 | 0 | 0 | 0 | 0 |
| Azohy | Genus | 3492.3 | 10964.7 | 0 | 0 | 0 | 0 | 0 | 0 | 0 | 0 |
| RS62 r | Genus | 3469.8 | 1713.7 | 0 | 0 | 0 | 0 | 0 | 0 | 0 | 0 |
| Simpli | Genus | 3337.3 | 0 | 0 | 0 | 0 | 0 | 0 | 0 | 0 | 0 |
| Pseud | Genus | 2449.2 | 123.3 | 0 | 0 | 0 | 0 | 0 | 0 | 0 | 0 |
| Pseud | Genus | 1741.8 | 321.3 | 0 | 0 | 0 | 0 | 0 | 0 | 0 | 0 |
| Roseat | Genus | 1001.6 | 381.5 | 0 | 0 | 0 | 0 | 0 | 0 | 0 | 0 |
| Ideone | Genus | 834.9 | 483.8 | 0 | 0 | 0 | 0 | 0 | 0 | 0 | 0 |
| Polaro | Genus | 557.9 | 852.9 | 0 | 0 | 0 | 0 | 0 | 0 | 0 | 0 |
| Inhella | Genus | 247.4 | 731.5 | 0 | 0 | 0 | 0 | 0 | 0 | 0 | 0 |
| Paucib | Genus | 169.7 | 64.9 | 0 | 0 | 0 | 0 | 0 | 0 | 0 | 0 |
| Mitsua | Genus | 104.6 | 127.2 | 0 | 0 | 0 | 0 | 0 | 0 | 0 | 0 |
| Delftia | Genus | 0 | 504.9 | 0 | 0 | 0 | 0 | 0 | 0 | 0 | 0 |
| Nitroso | Family | 534228.3 | 476940.5 | 2 | 1.7 | 0 | 0 | 0 | 0 | 0 | 0 |

| | | | | | | | | | | | |
|---|---|---|---|---|---|---|---|---|---|---|---|
| MND1 | Genus | 323861.1 | 259536.8 | 1.2 | 0.9 | 0 | 0 | 0 | 0 | 0 | 0 |
| Ellin60 | Genus | 130815.9 | 134367.2 | 0.5 | 0.5 | 0 | 0 | 0 | 0 | 0 | 0 |
| IS-44 | Genus | 74531.3 | 78518.2 | 0.3 | 0.3 | 0 | 0 | 0 | 0 | 0 | 0 |
| Nitros | Genus | 2833.7 | 2835.4 | 0 | 0 | 0 | 0 | 0 | 0 | 0 | 0 |
| Nitros | Genus | 1475 | 584.4 | 0 | 0 | 0 | 0 | 0 | 0 | 0 | 0 |
| Nitros | Genus | 711.4 | 738.4 | 0 | 0 | 0 | 0 | 0 | 0 | 0 | 0 |
| P3OB- | Genus | 0 | 360.1 | 0 | 0 | 0 | 0 | 0 | 0 | 0 | 0 |
| TRA3-2 | Family | 366898 | 362548.5 | 1.4 | 1.3 | 0.7 | 0 | 0 | 0 | 0.01 | 0 |
| TRA3-2 | Genus | 366898 | 362548.5 | 1.4 | 1.3 | 0.7 | 0 | 0 | 0 | 0.01 | 0 |
| Methylc | Family | 256296.3 | 346403.1 | 1 | 1.3 | 0 | 0 | 0 | 0 | 0 | 0 |
| Methyl | Genus | 193692.4 | 260486.7 | 0.7 | 0.9 | 0 | 0 | 0 | 0 | 0 | 0 |
| MM1 | Genus | 26608.5 | 30802.7 | 0.1 | 0.1 | 0 | 0 | 0 | 0 | 0 | 0 |
| Methyl | Genus | 24886.1 | 38611.5 | 0.1 | 0.1 | 0 | 0 | 0 | 0 | 0 | 0 |
| Methyl | Genus | 7033.8 | 11592.5 | 0 | 0 | 0 | 0 | 0 | 0 | 0 | 0 |
| Methyl | Genus | 2755.8 | 3576.1 | 0 | 0 | 0 | 0 | 0 | 0 | 0 | 0 |
| MM2 | Genus | 1319.6 | 1333.7 | 0 | 0 | 0 | 0 | 0 | 0 | 0 | 0 |
| Oxalob | Family | 190937.1 | 184852 | 0.7 | 0.7 | 0 | 0 | 18 | 0 | 0.13 | 0 |
| Dugan | Genus | 103644.8 | 59734.3 | 0.4 | 0.2 | 0 | 0 | 31.8 | 0 | 0.12 | 0 |
| Massil | Genus | 51365 | 70362.3 | 0.2 | 0.3 | 0 | 0 | 2.9 | 0 | 0.01 | 0 |
| Novihe | Genus | 17679.2 | 34637.3 | 0.1 | 0.1 | 0 | 0 | 0 | 0 | 0 | 0 |
| Oxalob | Genus | 12318.6 | 8314.9 | 0 | 0 | 0 | 0 | 0 | 0 | 0 | 0 |
| CM1G | Genus | 3352.5 | 5645 | 0 | 0 | 0 | 0 | 0 | 0 | 0 | 0 |
| Pseud | Genus | 2577 | 6051.5 | 0 | 0 | 0 | 0 | 0 | 0 | 0 | 0 |
| Oxalici | Genus | 0 | 106.7 | 0 | 0 | 0 | 0 | 0 | 0 | 0 | 0 |
| Alcalige | Family | 54676.2 | 20827 | 0.2 | 0.1 | 0 | 0 | 0 | 0 | 0 | 0 |
| Adven | Genus | 39818.7 | 6219.9 | 0.1 | 0 | 0 | 0 | 0 | 0 | 0 | 0 |
| Achror | Genus | 7330.3 | 3991.2 | 0 | 0 | 0 | 0 | 0 | 0 | 0 | 0 |
| Alcalig | Genus | 6192.3 | 6754.2 | 0 | 0 | 0 | 0 | 0 | 0 | 0 | 0 |
| Parapı | Genus | 1153.4 | 3666.4 | 0 | 0 | 0 | 0 | 0 | 0 | 0 | 0 |
| Pusillir | Genus | 181.5 | 195.4 | 0 | 0 | 0 | 0 | 0 | 0 | 0 | 0 |
| Burkhol | Family | 39941.1 | 40792.3 | 0.2 | 0.1 | 0 | 0 | 0 | 0 | 0 | 0 |

| | | | | | | | | | | | |
|---|---|---|---|---|---|---|---|---|---|---|---|
| Lautro | Genus | 27837.7 | 24834.3 | 0.1 | 0.1 | 0 | 0 | 0 | 0 | 0 | 0 |
| Burkho | Genus | 11490.2 | 15958 | 0 | 0.1 | 0 | 0 | 0 | 0 | 0 | 0 |
| Limnol | Genus | 613.2 | 0 | 0 | 0 | 0 | 0 | 0 | 0 | 0 | 0 |
| SC-I-84 | Family | 28426.6 | 27003.5 | 0.1 | 0.1 | 0 | 0 | 0 | 0 | 0 | 0 |
| SC-I-8 | Genus | 28426.6 | 27003.5 | 0.1 | 0.1 | 0 | 0 | 0 | 0 | 0 | 0 |
| Neisser | Family | 11602.4 | 8863.1 | 0 | 0 | 0 | 0 | 0 | 0 | 0 | 0 |
| Neisse | Genus | 11602.4 | 8863.1 | 0 | 0 | 0 | 0 | 0 | 0 | 0 | 0 |
| Burkhol | Family | 6293.7 | 6086.4 | 0 | 0 | 0 | 0 | 0 | 0 | 0 | 0 |
| Chitinir | Family | 1469.5 | 1368.8 | 0 | 0 | 0 | 0 | 0 | 0 | 0 | 0 |
| Chitini | Genus | 1469.5 | 1368.8 | 0 | 0 | 0 | 0 | 0 | 0 | 0 | 0 |
| Rhodoc | Family | 1345.5 | 780.7 | 0 | 0 | 0 | 0 | 0 | 0 | 0 | 0 |
| Rhodo | Genus | 1345.5 | 650.8 | 0 | 0 | 0 | 0 | 0 | 0 | 0 | 0 |
| Azoarc | Genus | 0 | 129.8 | 0 | 0 | 0 | 0 | 0 | 0 | 0 | 0 |
| Chromo | Family | 104.6 | 922.4 | 0 | 0 | 0 | 0 | 0 | 0 | 0 | 0 |
| Vogese | Genus | 104.6 | 922.4 | 0 | 0 | 0 | 0 | 0 | 0 | 0 | 0 |
| Xanthom | Order | 1996672 | 2691459 | 7.5 | 9.8 | 0 | 0 | 0 | 0 | 0 | 0 |
| Xanthoi | Family | 1202052 | 1788555 | 4.5 | 6.5 | 0 | 0 | 0 | 0 | 0 | 0 |
| Luteim | Genus | 450147.3 | 492578.3 | 1.7 | 1.8 | 0 | 0 | 0 | 0 | 0 | 0 |
| Pseud | Genus | 344981.2 | 741281 | 1.3 | 2.7 | 0 | 0 | 0 | 0 | 0 | 0 |
| Xantho | Genus | 127058.4 | 139157.9 | 0.5 | 0.5 | 0 | 0 | 0 | 0 | 0 | 0 |
| Arenim | Genus | 109915.5 | 151067.3 | 0.4 | 0.5 | 0 | 0 | 0 | 0 | 0 | 0 |
| Lysoba | Genus | 94589.5 | 103822.2 | 0.4 | 0.4 | 0 | 0 | 0 | 0 | 0 | 0 |
| Therm | Genus | 73502.4 | 146893 | 0.3 | 0.5 | 0 | 0 | 0 | 0 | 0 | 0 |
| Stenot | Genus | 1692.4 | 11623.8 | 0 | 0 | 0 | 0 | 0 | 0 | 0 | 0 |
| Xantho | Genus | 165.3 | 2131.5 | 0 | 0 | 0 | 0 | 0 | 0 | 0 | 0 |
| Rhodan | Family | 773540.8 | 886167.7 | 2.9 | 3.2 | 0 | 0 | 0 | 0 | 0 | 0 |
| Dokdo | Genus | 732694.4 | 806773.7 | 2.8 | 2.9 | 0 | 0 | 0 | 0 | 0 | 0 |
| Rhoda | Genus | 15384.9 | 22004.9 | 0.1 | 0.1 | 0 | 0 | 0 | 0 | 0 | 0 |
| Rudae | Genus | 8600.9 | 4409.5 | 0 | 0 | 0 | 0 | 0 | 0 | 0 | 0 |
| Dyella | Genus | 6618.8 | 13121.7 | 0 | 0 | 0 | 0 | 0 | 0 | 0 | 0 |
| Luteiba | Genus | 5607.5 | 18953 | 0 | 0.1 | 0 | 0 | 0 | 0 | 0 | 0 |

| | | | | | | | | | | | |
|---|---|---|---|---|---|---|---|---|---|---|---|
| Tahiba | Genus | 2270.9 | 2408.6 | 0 | 0 | 0 | 0 | 0 | 0 | 0 | 0 |
| Rhoda | Genus | 1491.8 | 8372.2 | 0 | 0 | 0 | 0 | 0 | 0 | 0 | 0 |
| Ahniell | Genus | 775.1 | 8230.6 | 0 | 0 | 0 | 0 | 0 | 0 | 0 | 0 |
| Chiayii | Genus | 96.7 | 1893.5 | 0 | 0 | 0 | 0 | 0 | 0 | 0 | 0 |
| Xanthoı | Family | 21079.2 | 16736.4 | 0.1 | 0.1 | 0 | 0 | 0 | 0 | 0 | 0 |
| Pseudor | Order | 656258.7 | 866630 | 2.5 | 3.2 | 0 | 0 | 0 | 0 | 0 | 0 |
| Pseudo | Family | 428683.3 | 376244 | 1.6 | 1.4 | 0 | 0 | 0 | 0 | 0 | 0 |
| Blyi10 | Genus | 428400.2 | 375309.2 | 1.6 | 1.4 | 0 | 0 | 0 | 0 | 0 | 0 |
| Pseud | Genus | 283.2 | 934.8 | 0 | 0 | 0 | 0 | 0 | 0 | 0 | 0 |
| Pseudo | Family | 150260.8 | 400215.7 | 0.6 | 1.5 | 0 | 0 | 0 | 0 | 0 | 0 |
| Pseud | Genus | 150061 | 399108 | 0.6 | 1.5 | 0 | 0 | 0 | 0 | 0 | 0 |
| Pseud | Genus | 199.9 | 624 | 0 | 0 | 0 | 0 | 0 | 0 | 0 | 0 |
| Azomo | Genus | 0 | 483.6 | 0 | 0 | 0 | 0 | 0 | 0 | 0 | 0 |
| Moraxel | Family | 32410.6 | 34923.3 | 0.1 | 0.1 | 0 | 0 | 0 | 0 | 0 | 0 |
| Moraxe | Genus | 31536.5 | 34923.3 | 0.1 | 0.1 | 0 | 0 | 0 | 0 | 0 | 0 |
| Acinet | Genus | 874.2 | 0 | 0 | 0 | 0 | 0 | 0 | 0 | 0 | 0 |
| 211ds2 | Family | 25019.4 | 33368.3 | 0.1 | 0.1 | 0 | 0 | 0 | 0 | 0 | 0 |
| 211ds2 | Genus | 25019.4 | 33368.3 | 0.1 | 0.1 | 0 | 0 | 0 | 0 | 0 | 0 |
| Cellvibr | Family | 10972.3 | 15005.7 | 0 | 0.1 | 0 | 0 | 0 | 0 | 0 | 0 |
| Cellvib | Genus | 10972.3 | 15005.7 | 0 | 0.1 | 0 | 0 | 0 | 0 | 0 | 0 |
| Pseudo | Family | 8622 | 6808.1 | 0 | 0 | 0 | 0 | 0 | 0 | 0 | 0 |
| Halieac | Family | 290.1 | 64.9 | 0 | 0 | 0 | 0 | 0 | 0 | 0 | 0 |
| Haliea | Genus | 290.1 | 64.9 | 0 | 0 | 0 | 0 | 0 | 0 | 0 | 0 |
| Gamma | Order | 437626.1 | 367893.4 | 1.6 | 1.3 | 0 | 0 | 0 | 0 | 0 | 0 |
| Unknow | Family | 437626.1 | 367893.4 | 1.6 | 1.3 | 0 | 0 | 0 | 0 | 0 | 0 |
| Acidiba | Genus | 429871.7 | 357767 | 1.6 | 1.3 | 0 | 0 | 0 | 0 | 0 | 0 |
| Unkno | Genus | 3664.5 | 4929.4 | 0 | 0 | 0 | 0 | 0 | 0 | 0 | 0 |
| Candic | Genus | 2055.7 | 2656.4 | 0 | 0 | 0 | 0 | 0 | 0 | 0 | 0 |
| Candic | Genus | 2034.2 | 2540.6 | 0 | 0 | 0 | 0 | 0 | 0 | 0 | 0 |
| CCD24 | Order | 222748.8 | 195642.4 | 0.8 | 0.7 | 0 | 0 | 0 | 0 | 0 | 0 |
| CCD24 | Family | 222748.8 | 195642.4 | 0.8 | 0.7 | 0 | 0 | 0 | 0 | 0 | 0 |

| | | | | | | | | | | | |
|---|---|---|---|---|---|---|---|---|---|---|---|
| Steroido | Order | 102870.6 | 76061.6 | 0.4 | 0.3 | 2.2 | 0 | 0 | 0 | 0.01 | 0 |
| Steroid | Family | 93420.8 | 68352.1 | 0.4 | 0.2 | 0 | 0 | 0 | 0 | 0 | 0 |
| Steroic | Genus | 91482.8 | 67146.7 | 0.3 | 0.2 | 0 | 0 | 0 | 0 | 0 | 0 |
| Steroic | Genus | 1938 | 1205.4 | 0 | 0 | 0 | 0 | 0 | 0 | 0 | 0 |
| Woesei | Family | 9449.8 | 7709.6 | 0 | 0 | 23.6 | 0 | 0 | 0 | 0.01 | 0 |
| Woese | Genus | 5029.5 | 5529.4 | 0 | 0 | 0 | 0 | 0 | 0 | 0 | 0 |
| JTB255 | Genus | 4420.2 | 2180.2 | 0 | 0 | 50.5 | 0 | 0 | 0 | 0.01 | 0 |
| EPR3968 | Order | 85413.9 | 63159.6 | 0.3 | 0.2 | 0 | 0 | 0 | 0 | 0 | 0 |
| EPR396 | Family | 85413.9 | 63159.6 | 0.3 | 0.2 | 0 | 0 | 0 | 0 | 0 | 0 |
| R7C24 | Order | 29903.8 | 25768.8 | 0.1 | 0.1 | 0 | 0 | 0 | 0 | 0 | 0 |
| R7C24_ | Family | 29903.8 | 25768.8 | 0.1 | 0.1 | 0 | 0 | 0 | 0 | 0 | 0 |
| PLTA13 | Order | 12057.3 | 15017.3 | 0 | 0.1 | 0 | 0 | 0 | 0 | 0 | 0 |
| PLTA13 | Family | 12057.3 | 15017.3 | 0 | 0.1 | 0 | 0 | 0 | 0 | 0 | 0 |
| Gamma | Order | 9826.8 | 11438.1 | 0 | 0 | 0 | 0 | 19 | 0 | 0.01 | 0 |
| Diplorick | Order | 5136.1 | 1786.9 | 0 | 0 | 0 | 0 | 0 | 0 | 0 | 0 |
| Diploric | Family | 5136.1 | 1786.9 | 0 | 0 | 0 | 0 | 0 | 0 | 0 | 0 |
| Diplori | Genus | 5136.1 | 1786.9 | 0 | 0 | 0 | 0 | 0 | 0 | 0 | 0 |
| JTB23 | Order | 4855.4 | 4405.2 | 0 | 0 | 0 | 0 | 0 | 0 | 0 | 0 |
| JTB23_( | Family | 4855.4 | 4405.2 | 0 | 0 | 0 | 0 | 0 | 0 | 0 | 0 |
| Enteroba | Order | 3548.3 | 28649.1 | 0 | 0.1 | 0 | 0 | 0 | 0 | 0 | 0 |
| Enterob | Family | 3548.3 | 25816.6 | 0 | 0.1 | 0 | 0 | 0 | 0 | 0 | 0 |
| Cedec | Genus | 2021.3 | 13428.2 | 0 | 0 | 0 | 0 | 0 | 0 | 0 | 0 |
| Entero | Genus | 1213.4 | 7740.4 | 0 | 0 | 0 | 0 | 0 | 0 | 0 | 0 |
| Entero | Genus | 313.7 | 2130.3 | 0 | 0 | 0 | 0 | 0 | 0 | 0 | 0 |
| Lelliott | Genus | 0 | 1432.7 | 0 | 0 | 0 | 0 | 0 | 0 | 0 | 0 |
| Buttiau | Genus | 0 | 1085 | 0 | 0 | 0 | 0 | 0 | 0 | 0 | 0 |
| Aeromo | Family | 0 | 2832.5 | 0 | 0 | 0 | 0 | 0 | 0 | 0 | 0 |
| Aerom | Genus | 0 | 2832.5 | 0 | 0 | 0 | 0 | 0 | 0 | 0 | 0 |
| Salinispl | Order | 1073.8 | 3661.9 | 0 | 0 | 0 | 0 | 0 | 0 | 0 | 0 |
| Solimor | Family | 1073.8 | 3661.9 | 0 | 0 | 0 | 0 | 0 | 0 | 0 | 0 |
| Fontim | Genus | 727.6 | 3493 | 0 | 0 | 0 | 0 | 0 | 0 | 0 | 0 |

| | | | | | | | | | | | |
|---|---|---|---|---|---|---|---|---|---|---|---|
| Panaca | Genus | 346.2 | 168.9 | 0 | 0 | 0 | 0 | 0 | 0 | 0 | 0 |
| Ga0077! | Order | 709.9 | 2785.8 | 0 | 0 | 0 | 0 | 0 | 0 | 0 | 0 |
| Ga0077 | Family | 709.9 | 2785.8 | 0 | 0 | 0 | 0 | 0 | 0 | 0 | 0 |
| Granulos | Order | 369.6 | 648 | 0 | 0 | 0 | 0 | 0 | 0 | 0 | 0 |
| Granulc | Family | 369.6 | 648 | 0 | 0 | 0 | 0 | 0 | 0 | 0 | 0 |
| Chromat | Order | 86.5 | 337.7 | 0 | 0 | 0 | 0 | 0 | 0 | 0 | 0 |
| Sedime | Family | 86.5 | 337.7 | 0 | 0 | 0 | 0 | 0 | 0 | 0 | 0 |
| Sedim | Genus | 86.5 | 337.7 | 0 | 0 | 0 | 0 | 0 | 0 | 0 | 0 |
| Legionel | Order | 0 | 336.2 | 0 | 0 | 0 | 0 | 0 | 0 | 0 | 0 |
| Legione | Family | 0 | 336.2 | 0 | 0 | 0 | 0 | 0 | 0 | 0 | 0 |
| Legion | Genus | 0 | 336.2 | 0 | 0 | 0 | 0 | 0 | 0 | 0 | 0 |
| Alphaprot | Class | 6038064 | 5452393 | 22.7 | 19.8 | 0.2 | 0.3 | 0.2 | 0 | 0.08 | 0.07 |
| Sphingo | Order | 2256883 | 1875646 | 8.5 | 6.8 | 0.1 | 0.9 | 0.4 | 0 | 0.04 | 0.06 |
| Sphingc | Family | 2256883 | 1875646 | 8.5 | 6.8 | 0.1 | 0.9 | 0.4 | 0 | 0.04 | 0.06 |
| Sphing | Genus | 1208587 | 1012636 | 4.5 | 3.7 | 0 | 0 | 0.7 | 0 | 0.03 | 0 |
| Sphing | Genus | 416626.7 | 324541.3 | 1.6 | 1.2 | 0 | 5.1 | 0 | 0 | 0 | 0.06 |
| Alterer | Genus | 373879 | 329390.9 | 1.4 | 1.2 | 0 | 0 | 0 | 0 | 0 | 0 |
| Sphing | Genus | 111163.7 | 92512.2 | 0.4 | 0.3 | 0 | 0 | 0 | 0 | 0 | 0 |
| Ellin60 | Genus | 76758.4 | 68424.8 | 0.3 | 0.2 | 0 | 0 | 0 | 0 | 0 | 0 |
| Sphing | Genus | 53082.4 | 31789 | 0.2 | 0.1 | 6.1 | 0 | 0 | 0 | 0.01 | 0 |
| Novos | Genus | 10766 | 11564 | 0 | 0 | 0 | 0 | 0 | 0 | 0 | 0 |
| Plot4-2 | Genus | 3210.8 | 2818.3 | 0 | 0 | 0 | 0 | 0 | 0 | 0 | 0 |
| Porphy | Genus | 2159.2 | 582.6 | 0 | 0 | 0 | 0 | 0 | 0 | 0 | 0 |
| Qipeng | Genus | 478.2 | 1098.3 | 0 | 0 | 0 | 0 | 0 | 0 | 0 | 0 |
| Parabl | Genus | 171.4 | 159.1 | 0 | 0 | 0 | 0 | 0 | 0 | 0 | 0 |
| Sphing | Genus | 0 | 129.8 | 0 | 0 | 0 | 0 | 0 | 0 | 0 | 0 |
| Rhizobia | Order | 1998670 | 1772690 | 7.5 | 6.5 | 0.2 | 0.1 | 0.1 | 0.1 | 0.02 | 0.01 |
| Devosia | Family | 599206.4 | 528336.9 | 2.3 | 1.9 | 0.6 | 0.4 | 0 | 0 | 0.01 | 0.01 |
| Devosi | Genus | 559784.5 | 498695.6 | 2.1 | 1.8 | 0.7 | 0.4 | 0 | 0 | 0.01 | 0.01 |
| Devosi | Genus | 39421.9 | 29641.3 | 0.1 | 0.1 | 0 | 0 | 0 | 0 | 0 | 0 |
| Hyphon | Family | 394907.6 | 287321.6 | 1.5 | 1 | 0 | 0 | 0 | 0 | 0 | 0 |

| | | | | | | | | | | | |
|---|---|---|---|---|---|---|---|---|---|---|---|
| Pedom | Genus | 232981.3 | 152940.8 | 0.9 | 0.6 | 0 | 0 | 0 | 0 | 0 | 0 |
| Hypho | Genus | 145548.8 | 116586.1 | 0.5 | 0.4 | 0 | 0 | 0 | 0 | 0 | 0 |
| Hypho | Genus | 16377.5 | 17794.7 | 0.1 | 0.1 | 0 | 0 | 0 | 0 | 0 | 0 |
| Xanthol | Family | 343144.3 | 346793.8 | 1.3 | 1.3 | 0 | 0 | 0 | 0 | 0 | 0 |
| Pseud | Genus | 152665.6 | 185071.7 | 0.6 | 0.7 | 0 | 0 | 0 | 0 | 0 | 0 |
| Xanthc | Genus | 78336.9 | 60342.7 | 0.3 | 0.2 | 0 | 0 | 0 | 0 | 0 | 0 |
| Pseud | Genus | 39069.9 | 36716.7 | 0.1 | 0.1 | 0 | 0 | 0 | 0 | 0 | 0 |
| Rhodo | Genus | 36815.1 | 31350.3 | 0.1 | 0.1 | 0 | 0 | 0 | 0 | 0 | 0 |
| Bradyr | Genus | 23754 | 22543.2 | 0.1 | 0.1 | 0 | 0 | 0 | 0 | 0 | 0 |
| Rhodo | Genus | 6941.6 | 6687.6 | 0 | 0 | 0 | 0 | 0 | 0 | 0 | 0 |
| Oligotr | Genus | 3948 | 2983.3 | 0 | 0 | 0 | 0 | 0 | 0 | 0 | 0 |
| Nitrob | Genus | 1113.5 | 1033.5 | 0 | 0 | 0 | 0 | 0 | 0 | 0 | 0 |
| Xanthc | Genus | 499.7 | 64.9 | 0 | 0 | 0 | 0 | 0 | 0 | 0 | 0 |
| Rhizobi | Family | 267969.9 | 250553.8 | 1 | 0.9 | 0 | 0 | 0.6 | 0 | 0.01 | 0 |
| Rhizob | Genus | 47800.8 | 41098.3 | 0.2 | 0.1 | 0 | 0 | 0 | 0 | 0 | 0 |
| Pseud | Genus | 47662.6 | 36720.6 | 0.2 | 0.1 | 0 | 0 | 0 | 0 | 0 | 0 |
| Shinell | Genus | 43391.1 | 71571.1 | 0.2 | 0.3 | 0 | 0 | 0 | 0 | 0 | 0 |
| Mesorl | Genus | 34191.6 | 22279.1 | 0.1 | 0.1 | 0 | 0 | 0 | 0 | 0 | 0 |
| [Rhizol | Genus | 32679.4 | 25059.3 | 0.1 | 0.1 | 0 | 0 | 0 | 0 | 0 | 0 |
| Allorhi | Genus | 24698.5 | 21078.9 | 0.1 | 0.1 | 0 | 0 | 0 | 0 | 0 | 0 |
| Amino | Genus | 11482.2 | 10461.2 | 0 | 0 | 0 | 0 | 0 | 0 | 0 | 0 |
| Ochrol | Genus | 8351.8 | 3026.8 | 0 | 0 | 0 | 0 | 18.8 | 0 | 0.01 | 0 |
| Cicerit | Genus | 6286.9 | 4603.4 | 0 | 0 | 0 | 0 | 0 | 0 | 0 | 0 |
| Phyllol | Genus | 3622.1 | 1687.2 | 0 | 0 | 0 | 0 | 0 | 0 | 0 | 0 |
| Ensifer | Genus | 3425.8 | 10057.1 | 0 | 0 | 0 | 0 | 0 | 0 | 0 | 0 |
| Pseud | Genus | 3087.8 | 129.8 | 0 | 0 | 0 | 0 | 0 | 0 | 0 | 0 |
| Chelat | Genus | 1024 | 265.9 | 0 | 0 | 0 | 0 | 0 | 0 | 0 | 0 |
| Mycop | Genus | 165.3 | 0 | 0 | 0 | 0 | 0 | 0 | 0 | 0 | 0 |
| Aliihoe | Genus | 99.9 | 0 | 0 | 0 | 0 | 0 | 0 | 0 | 0 | 0 |
| Neorhi | Genus | 0 | 2515.1 | 0 | 0 | 0 | 0 | 0 | 0 | 0 | 0 |
| KF-JG3( | Family | 87705 | 87796.8 | 0.3 | 0.3 | 0 | 0 | 0 | 1.8 | 0 | 0.01 |

| | | | | | | | | | | | |
|---|---|---|---|---|---|---|---|---|---|---|---|
| KF-JG3 | Genus | 87705 | 87796.8 | 0.3 | 0.3 | 0 | 0 | 0 | 1.8 | 0 | 0.01 |
| C2U | Family | 77362.5 | 54894.7 | 0.3 | 0.2 | 0 | 0 | 0 | 0 | 0 | 0 |
| C2U_F | Genus | 77362.5 | 54894.7 | 0.3 | 0.2 | 0 | 0 | 0 | 0 | 0 | 0 |
| Amb-16 | Family | 72834.7 | 65911.3 | 0.3 | 0.2 | 0 | 0 | 0 | 0 | 0 | 0 |
| Amb-1 | Genus | 72834.7 | 65911.3 | 0.3 | 0.2 | 0 | 0 | 0 | 0 | 0 | 0 |
| Rhizobi | Family | 52457.7 | 51459.8 | 0.2 | 0.2 | 0 | 0 | 0 | 0 | 0 | 0 |
| Bauldi | Genus | 50624.6 | 49719.8 | 0.2 | 0.2 | 0 | 0 | 0 | 0 | 0 | 0 |
| Rhizob | Genus | 971.3 | 286.2 | 0 | 0 | 0 | 0 | 0 | 0 | 0 | 0 |
| Nordel | Genus | 861.8 | 1453.8 | 0 | 0 | 0 | 0 | 0 | 0 | 0 | 0 |
| Rhizobi | Family | 39762.7 | 38490.2 | 0.1 | 0.1 | 0 | 0 | 0 | 0 | 0 | 0 |
| Beijerin | Family | 24230.3 | 25242.2 | 0.1 | 0.1 | 0 | 0 | 0 | 0 | 0 | 0 |
| Microv | Genus | 12577 | 14735.5 | 0 | 0.1 | 0 | 0 | 0 | 0 | 0 | 0 |
| Bosea | Genus | 10125.4 | 10103.6 | 0 | 0 | 0 | 0 | 0 | 0 | 0 | 0 |
| Neo-b1 | Genus | 797.6 | 0 | 0 | 0 | 0 | 0 | 0 | 0 | 0 | 0 |
| Methyl | Genus | 730.3 | 403.1 | 0 | 0 | 0 | 0 | 0 | 0 | 0 | 0 |
| Labrace | Family | 21385.8 | 19167.9 | 0.1 | 0.1 | 0 | 0 | 0 | 0 | 0 | 0 |
| Labrys | Genus | 21385.8 | 19167.9 | 0.1 | 0.1 | 0 | 0 | 0 | 0 | 0 | 0 |
| D05-2 | Family | 10174.9 | 7014.2 | 0 | 0 | 0 | 0 | 0 | 0 | 0 | 0 |
| D05-2_ | Genus | 10174.9 | 7014.2 | 0 | 0 | 0 | 0 | 0 | 0 | 0 | 0 |
| Methylc | Family | 4831.5 | 7415.4 | 0 | 0 | 0 | 0 | 0 | 0 | 0 | 0 |
| Methyl | Genus | 4831.5 | 7415.4 | 0 | 0 | 0 | 0 | 0 | 0 | 0 | 0 |
| A0839 | Family | 1691.8 | 1544.6 | 0 | 0 | 0 | 0 | 0 | 0 | 0 | 0 |
| A0839_ | Genus | 1691.8 | 1544.6 | 0 | 0 | 0 | 0 | 0 | 0 | 0 | 0 |
| WC3-11 | Family | 920.1 | 408.9 | 0 | 0 | 0 | 0 | 0 | 0 | 0 | 0 |
| WC3-1 | Genus | 920.1 | 408.9 | 0 | 0 | 0 | 0 | 0 | 0 | 0 | 0 |
| Rhodob | Family | 84.8 | 253.4 | 0 | 0 | 0 | 0 | 0 | 0 | 0 | 0 |
| Tepida | Genus | 84.8 | 253.4 | 0 | 0 | 0 | 0 | 0 | 0 | 0 | 0 |
| Kaistiac | Family | 0 | 84.5 | 0 | 0 | 0 | 0 | 0 | 0 | 0 | 0 |
| Kaistia | Genus | 0 | 84.5 | 0 | 0 | 0 | 0 | 0 | 0 | 0 | 0 |
| Cauloba | Order | 901712.2 | 909820.9 | 3.4 | 3.3 | 0 | 0 | 0 | 0 | 0 | 0 |
| Caulob | Family | 627255.2 | 602162.2 | 2.4 | 2.2 | 0 | 0 | 0 | 0 | 0 | 0 |

| | | | | | | | | | | | |
|---|---|---|---|---|---|---|---|---|---|---|---|
| Caulot | Genus | 297202.8 | 279465.9 | 1.1 | 1 | 0 | 0 | 0 | 0 | 0 | 0 |
| Brevur | Genus | 195297.9 | 196059.2 | 0.7 | 0.7 | 0 | 0 | 0 | 0 | 0 | 0 |
| Phenyl | Genus | 88129.8 | 71376.6 | 0.3 | 0.3 | 0 | 0 | 0 | 0 | 0 | 0 |
| Caulot | Genus | 26090.9 | 23601.9 | 0.1 | 0.1 | 0 | 0 | 0 | 0 | 0 | 0 |
| Asticca | Genus | 20533.9 | 31658.7 | 0.1 | 0.1 | 0 | 0 | 0 | 0 | 0 | 0 |
| Hyphon | Family | 274302.9 | 306990 | 1 | 1.1 | 0 | 0 | 0 | 0 | 0 | 0 |
| SWB02 | Genus | 208333.5 | 250526.8 | 0.8 | 0.9 | 0 | 0 | 0 | 0 | 0 | 0 |
| Hirsch | Genus | 65969.4 | 56463.2 | 0.2 | 0.2 | 0 | 0 | 0 | 0 | 0 | 0 |
| Parvula | Family | 154.1 | 415.4 | 0 | 0 | 0 | 0 | 0 | 0 | 0 | 0 |
| Amphi | Genus | 154.1 | 415.4 | 0 | 0 | 0 | 0 | 0 | 0 | 0 | 0 |
| Caulob | Family | 0 | 253.4 | 0 | 0 | 0 | 0 | 0 | 0 | 0 | 0 |
| Micrope | Order | 404848.1 | 400568.7 | 1.5 | 1.5 | 0 | 0 | 0 | 0 | 0 | 0 |
| Micrope | Family | 404848.1 | 400568.7 | 1.5 | 1.5 | 0 | 0 | 0 | 0 | 0 | 0 |
| Microp | Genus | 404848.1 | 400568.7 | 1.5 | 1.5 | 0 | 0 | 0 | 0 | 0 | 0 |
| Alphapro | Order | 163627.9 | 154770.1 | 0.6 | 0.6 | 0 | 0 | 0 | 0 | 0 | 0 |
| Reyranel | Order | 98165.4 | 109972.3 | 0.4 | 0.4 | 0 | 0 | 2.1 | 0 | 0.01 | 0 |
| Reyrane | Family | 98165.4 | 109972.3 | 0.4 | 0.4 | 0 | 0 | 2.1 | 0 | 0.01 | 0 |
| Reyran | Genus | 51000.8 | 54959.1 | 0.2 | 0.2 | 0 | 0 | 0 | 0 | 0 | 0 |
| Reyran | Genus | 47164.7 | 55013.2 | 0.2 | 0.2 | 0 | 0 | 4.5 | 0 | 0.01 | 0 |
| Rhodosp | Order | 97131.6 | 97571.5 | 0.4 | 0.4 | 0 | 0 | 0 | 0 | 0 | 0 |
| Rhodos | Family | 43348.1 | 40625.4 | 0.2 | 0.1 | 0 | 0 | 0 | 0 | 0 | 0 |
| Rhodos | Family | 28087.8 | 28683.9 | 0.1 | 0.1 | 0 | 0 | 0 | 0 | 0 | 0 |
| Rhodo | Genus | 28087.8 | 28683.9 | 0.1 | 0.1 | 0 | 0 | 0 | 0 | 0 | 0 |
| Magnet | Family | 24753.9 | 26199.9 | 0.1 | 0.1 | 0 | 0 | 0 | 0 | 0 | 0 |
| Magne | Genus | 24753.9 | 26199.9 | 0.1 | 0.1 | 0 | 0 | 0 | 0 | 0 | 0 |
| Magnet | Family | 941.9 | 2062.3 | 0 | 0 | 0 | 0 | 0 | 0 | 0 | 0 |
| Magne | Genus | 941.9 | 2062.3 | 0 | 0 | 0 | 0 | 0 | 0 | 0 | 0 |
| Defluviic | Order | 29557.7 | 25168.6 | 0.1 | 0.1 | 0 | 0 | 0 | 0 | 0 | 0 |
| Defluvii | Family | 20214.2 | 15905.4 | 0.1 | 0.1 | 0 | 0 | 0 | 0 | 0 | 0 |
| Defluv | Genus | 20214.2 | 15905.4 | 0.1 | 0.1 | 0 | 0 | 0 | 0 | 0 | 0 |
| Defluvii | Family | 9343.5 | 9263.2 | 0 | 0 | 0 | 0 | 0 | 0 | 0 | 0 |

| | | | | | | | | | | | |
|---|---|---|---|---|---|---|---|---|---|---|---|
| Micavibr | Order | 18291.5 | 15976.3 | 0.1 | 0.1 | 0 | 0 | 0 | 0 | 0 | 0 |
| Micavib | Family | 16555.5 | 14577.8 | 0.1 | 0.1 | 0 | 0 | 0 | 0 | 0 | 0 |
| Micavib | Family | 1736 | 1398.5 | 0 | 0 | 0 | 0 | 0 | 0 | 0 | 0 |
| Micavi | Genus | 1736 | 1398.5 | 0 | 0 | 0 | 0 | 0 | 0 | 0 | 0 |
| Dongiale | Order | 17200.8 | 19032.9 | 0.1 | 0.1 | 0 | 0 | 0 | 0 | 0 | 0 |
| Dongiac | Family | 17200.8 | 19032.9 | 0.1 | 0.1 | 0 | 0 | 0 | 0 | 0 | 0 |
| Dongia | Genus | 17200.8 | 19032.9 | 0.1 | 0.1 | 0 | 0 | 0 | 0 | 0 | 0 |
| Rickettsi | Order | 15317.1 | 20144.9 | 0.1 | 0.1 | 20.2 | 0 | 0 | 0 | 0.01 | 0 |
| SM2D1 | Family | 15317.1 | 20144.9 | 0.1 | 0.1 | 20.2 | 0 | 0 | 0 | 0.01 | 0 |
| SM2D1 | Genus | 15317.1 | 20144.9 | 0.1 | 0.1 | 20.2 | 0 | 0 | 0 | 0.01 | 0 |
| Rhodoba | Order | 10487.7 | 15247.3 | 0 | 0.1 | 0 | 0 | 0 | 0 | 0 | 0 |
| Rhodob | Family | 10487.7 | 15247.3 | 0 | 0.1 | 0 | 0 | 0 | 0 | 0 | 0 |
| Amaric | Genus | 5116 | 6854.7 | 0 | 0 | 0 | 0 | 0 | 0 | 0 | 0 |
| Rhodo | Genus | 3508 | 5094.5 | 0 | 0 | 0 | 0 | 0 | 0 | 0 | 0 |
| Paracc | Genus | 1863.6 | 3298.2 | 0 | 0 | 0 | 0 | 0 | 0 | 0 | 0 |
| Azospiril | Order | 6392.5 | 13564.9 | 0 | 0 | 0 | 0 | 0 | 0 | 0 | 0 |
| Azospir | Family | 4827.9 | 10393.5 | 0 | 0 | 0 | 0 | 0 | 0 | 0 | 0 |
| Azospi | Genus | 4528 | 10140.4 | 0 | 0 | 0 | 0 | 0 | 0 | 0 | 0 |
| Azospi | Genus | 299.8 | 253.1 | 0 | 0 | 0 | 0 | 0 | 0 | 0 | 0 |
| Azospir | Family | 1564.7 | 2340.4 | 0 | 0 | 0 | 0 | 0 | 0 | 0 | 0 |
| Stella | Genus | 1564.7 | 2340.4 | 0 | 0 | 0 | 0 | 0 | 0 | 0 | 0 |
| Inquilin | Family | 0 | 831 | 0 | 0 | 0 | 0 | 0 | 0 | 0 | 0 |
| Inquilii | Genus | 0 | 831 | 0 | 0 | 0 | 0 | 0 | 0 | 0 | 0 |
| Kiloniella | Order | 4040.7 | 3485.8 | 0 | 0 | 0 | 0 | 0 | 0 | 0 | 0 |
| Fodinic | Family | 2130.4 | 2405 | 0 | 0 | 0 | 0 | 0 | 0 | 0 | 0 |
| Fodinic | Genus | 2130.4 | 2405 | 0 | 0 | 0 | 0 | 0 | 0 | 0 | 0 |
| Kiloniel | Family | 1910.2 | 1080.8 | 0 | 0 | 0 | 0 | 0 | 0 | 0 | 0 |
| Kilonie | Genus | 1910.2 | 1080.8 | 0 | 0 | 0 | 0 | 0 | 0 | 0 | 0 |
| Zavarzin | Order | 4039.3 | 3043.5 | 0 | 0 | 0 | 0 | 0 | 0 | 0 | 0 |
| Zavarzii | Family | 4039.3 | 3043.5 | 0 | 0 | 0 | 0 | 0 | 0 | 0 | 0 |
| Elsterale | Order | 3586.4 | 3684.2 | 0 | 0 | 0 | 0 | 0 | 0 | 0 | 0 |

| | | | | | | | | | | | |
|---|---|---|---|---|---|---|---|---|---|---|---|
| Elsteral | Family | 3586.4 | 3684.2 | 0 | 0 | 0 | 0 | 0 | 0 | 0 | 0 |
| Ferrovibı | Order | 3538.4 | 5182.6 | 0 | 0 | 0 | 0 | 0 | 0 | 0 | 0 |
| Ferrovik | Family | 2036.8 | 2927.6 | 0 | 0 | 0 | 0 | 0 | 0 | 0 | 0 |
| Ferrovi | Genus | 2036.8 | 2927.6 | 0 | 0 | 0 | 0 | 0 | 0 | 0 | 0 |
| Taonell | Family | 987.4 | 2125.1 | 0 | 0 | 0 | 0 | 0 | 0 | 0 | 0 |
| Taonel | Genus | 987.4 | 2125.1 | 0 | 0 | 0 | 0 | 0 | 0 | 0 | 0 |
| Ferrovik | Family | 514.2 | 129.8 | 0 | 0 | 0 | 0 | 0 | 0 | 0 | 0 |
| Acetoba | Order | 2235.1 | 3595.7 | 0 | 0 | 0 | 0 | 0 | 0 | 0 | 0 |
| Acetoba | Family | 2235.1 | 3595.7 | 0 | 0 | 0 | 0 | 0 | 0 | 0 | 0 |
| Craurc | Genus | 1779.6 | 1477.2 | 0 | 0 | 0 | 0 | 0 | 0 | 0 | 0 |
| Acetok | Genus | 455.4 | 2034 | 0 | 0 | 0 | 0 | 0 | 0 | 0 | 0 |
| Roseoı | Genus | 0 | 84.5 | 0 | 0 | 0 | 0 | 0 | 0 | 0 | 0 |
| Paracae | Order | 1593.7 | 2317.6 | 0 | 0 | 0 | 0 | 0 | 0 | 0 | 0 |
| Paracae | Family | 1593.7 | 2317.6 | 0 | 0 | 0 | 0 | 0 | 0 | 0 | 0 |
| Candic | Genus | 1039.8 | 1947.8 | 0 | 0 | 0 | 0 | 0 | 0 | 0 | 0 |
| Candic | Genus | 553.9 | 369.8 | 0 | 0 | 0 | 0 | 0 | 0 | 0 | 0 |
| AT-s3-44 | Order | 745.5 | 910.1 | 0 | 0 | 0 | 0 | 0 | 0 | 0 | 0 |
| AT-s3-4 | Family | 745.5 | 910.1 | 0 | 0 | 0 | 0 | 0 | 0 | 0 | 0 |
| Firmicutes | Phylum | 4015643 | 4007149 | 15.1 | 14.6 | 0.3 | 0 | 77.5 | 3.7 | 11.73 | 0.55 |
| Bacilli | Class | 4006686 | 3999868 | 15.1 | 14.6 | 0.3 | 0 | 77.6 | 3.8 | 11.73 | 0.55 |
| Bacillale | Order | 3828384 | 3775369 | 14.4 | 13.7 | 0.3 | 0 | 81.1 | 3.9 | 11.71 | 0.54 |
| Bacillac | Family | 3582330 | 3484343 | 13.5 | 12.7 | 0.3 | 0 | 82.4 | 4.2 | 11.14 | 0.54 |
| Bacillu | Genus | 3451603 | 3342340 | 13 | 12.2 | 0.3 | 0 | 85.4 | 4.4 | 11.12 | 0.54 |
| Bacilla | Genus | 58028.3 | 60297.3 | 0.2 | 0.2 | 0 | 0 | 0 | 0 | 0 | 0 |
| Geoba | Genus | 49391.5 | 55808.5 | 0.2 | 0.2 | 0 | 0 | 0 | 0 | 0 | 0 |
| Virgiba | Genus | 6393.1 | 7197.5 | 0 | 0 | 0 | 0 | 0 | 0 | 0 | 0 |
| Ureiba | Genus | 4979.2 | 4361.7 | 0 | 0 | 0 | 0 | 0 | 0 | 0 | 0 |
| Anoxyk | Genus | 4571.1 | 2881.9 | 0 | 0 | 0 | 0 | 0 | 0 | 0 | 0 |
| Falsiba | Genus | 4163 | 7336.8 | 0 | 0 | 0 | 0 | 100 | 0 | 0.02 | 0 |
| Fictiba | Genus | 3199.8 | 4119.1 | 0 | 0 | 0 | 0 | 0 | 0 | 0 | 0 |
| Planocc | Family | 246054.2 | 291025.5 | 0.9 | 1.1 | 0 | 0 | 61.6 | 0 | 0.57 | 0 |

| | | | | | | | | | | | |
|---|---|---|---|---|---|---|---|---|---|---|---|
| Lysinib | Genus | 65634.8 | 76906.6 | 0.2 | 0.3 | 0 | 0 | 66.1 | 0 | 0.16 | 0 |
| Paenis | Genus | 56793.5 | 55395.9 | 0.2 | 0.2 | 0 | 0 | 28.5 | 0 | 0.06 | 0 |
| Planoc | Genus | 48957.8 | 65900 | 0.2 | 0.2 | 0 | 0 | 94.5 | 0 | 0.17 | 0 |
| Sporos | Genus | 45799.8 | 61600.6 | 0.2 | 0.2 | 0 | 0 | 100 | 0 | 0.17 | 0 |
| Chung | Genus | 28361.3 | 29986.7 | 0.1 | 0.1 | 0 | 0 | 0 | 0 | 0 | 0 |
| Psychr | Genus | 506.9 | 1040.9 | 0 | 0 | 0 | 0 | 0 | 0 | 0 | 0 |
| Domib | Genus | 0 | 194.8 | 0 | 0 | 0 | 0 | 0 | 0 | 0 | 0 |
| Paeniba | Order | 134392.5 | 173701.1 | 0.5 | 0.6 | 0 | 0 | 4.9 | 1.3 | 0.02 | 0.01 |
| Paeniba | Family | 134392.5 | 173701.1 | 0.5 | 0.6 | 0 | 0 | 4.9 | 1.3 | 0.02 | 0.01 |
| Ammo | Genus | 70584 | 77719.5 | 0.3 | 0.3 | 0 | 0 | 0 | 0 | 0 | 0 |
| Paenib | Genus | 63537.2 | 94963.3 | 0.2 | 0.3 | 0 | 0 | 10.4 | 2.3 | 0.02 | 0.01 |
| Cohne | Genus | 271.3 | 1018.4 | 0 | 0 | 0 | 0 | 0 | 0 | 0 | 0 |
| Thermoa | Order | 28412.3 | 35274.2 | 0.1 | 0.1 | 0 | 0 | 0 | 0 | 0 | 0 |
| Thermo | Family | 28412.3 | 35274.2 | 0.1 | 0.1 | 0 | 0 | 0 | 0 | 0 | 0 |
| Shima: | Genus | 15877.9 | 22074.4 | 0.1 | 0.1 | 0 | 0 | 0 | 0 | 0 | 0 |
| Planifil | Genus | 7718.7 | 8848.7 | 0 | 0 | 0 | 0 | 0 | 0 | 0 | 0 |
| Hazen | Genus | 1836.7 | 1588.7 | 0 | 0 | 0 | 0 | 0 | 0 | 0 | 0 |
| Therm | Genus | 1683.4 | 1886.4 | 0 | 0 | 0 | 0 | 0 | 0 | 0 | 0 |
| Therm | Genus | 715.4 | 554.6 | 0 | 0 | 0 | 0 | 0 | 0 | 0 | 0 |
| Laceye | Genus | 580.2 | 321.3 | 0 | 0 | 0 | 0 | 0 | 0 | 0 | 0 |
| Aneurini | Order | 5211.1 | 5656.1 | 0 | 0 | 0 | 0 | 0 | 0 | 0 | 0 |
| Aneurin | Family | 5211.1 | 5656.1 | 0 | 0 | 0 | 0 | 0 | 0 | 0 | 0 |
| Aneuri | Genus | 5211.1 | 5656.1 | 0 | 0 | 0 | 0 | 0 | 0 | 0 | 0 |
| Brevibac | Order | 5071 | 4337.5 | 0 | 0 | 0 | 0 | 0 | 0 | 0 | 0 |
| Breviba | Family | 5071 | 4337.5 | 0 | 0 | 0 | 0 | 0 | 0 | 0 | 0 |
| Brevib | Genus | 5071 | 4337.5 | 0 | 0 | 0 | 0 | 0 | 0 | 0 | 0 |
| Alicyclob | Order | 3689.4 | 4955.8 | 0 | 0 | 0 | 0 | 0 | 0 | 0 | 0 |
| Alicyclo | Family | 3689.4 | 4955.8 | 0 | 0 | 0 | 0 | 0 | 0 | 0 | 0 |
| Tumeb | Genus | 3689.4 | 4955.8 | 0 | 0 | 0 | 0 | 0 | 0 | 0 | 0 |
| Exiguoba | Order | 1526 | 574.3 | 0 | 0 | 0 | 0 | 0 | 0 | 0 | 0 |
| Exiguob | Family | 1526 | 574.3 | 0 | 0 | 0 | 0 | 0 | 0 | 0 | 0 |

| | | | | | | | | | | | |
|---|---|---|---|---|---|---|---|---|---|---|---|
| Exiguo | Genus | 1526 | 574.3 | 0 | 0 | 0 | 0 | 0 | 0 | 0 | 0 |
| Clostridia | Class | 5311.1 | 3562.3 | 0 | 0 | 0 | 0 | 0 | 0 | 0 | 0 |
| Clostridi | Order | 3230.8 | 2220.1 | 0 | 0 | 0 | 0 | 0 | 0 | 0 | 0 |
| Clostric | Family | 3230.8 | 2220.1 | 0 | 0 | 0 | 0 | 0 | 0 | 0 | 0 |
| Clostri | Genus | 1727.3 | 644.7 | 0 | 0 | 0 | 0 | 0 | 0 | 0 | 0 |
| Clostri | Genus | 845.3 | 0 | 0 | 0 | 0 | 0 | 0 | 0 | 0 | 0 |
| Clostri | Genus | 313.7 | 965.7 | 0 | 0 | 0 | 0 | 0 | 0 | 0 | 0 |
| Clostri | Genus | 259.6 | 0 | 0 | 0 | 0 | 0 | 0 | 0 | 0 | 0 |
| Clostri | Genus | 84.8 | 609.6 | 0 | 0 | 0 | 0 | 0 | 0 | 0 | 0 |
| Peptostr | Order | 1501.7 | 1342.2 | 0 | 0 | 0 | 0 | 0 | 0 | 0 | 0 |
| Peptost | Family | 1501.7 | 1342.2 | 0 | 0 | 0 | 0 | 0 | 0 | 0 | 0 |
| Sporac | Genus | 1006.5 | 1095.7 | 0 | 0 | 0 | 0 | 0 | 0 | 0 | 0 |
| Rombc | Genus | 495.2 | 246.5 | 0 | 0 | 0 | 0 | 0 | 0 | 0 | 0 |
| Graciliba | Order | 578.6 | 0 | 0 | 0 | 0 | 0 | 0 | 0 | 0 | 0 |
| Gracilib | Family | 578.6 | 0 | 0 | 0 | 0 | 0 | 0 | 0 | 0 | 0 |
| Symbioba | Class | 3645.9 | 3718.9 | 0 | 0 | 0 | 0 | 0 | 0 | 0 | 0 |
| Symbiob | Order | 3645.9 | 3718.9 | 0 | 0 | 0 | 0 | 0 | 0 | 0 | 0 |
| Symbio | Family | 3645.9 | 3718.9 | 0 | 0 | 0 | 0 | 0 | 0 | 0 | 0 |
| Symbi | Genus | 3645.9 | 3718.9 | 0 | 0 | 0 | 0 | 0 | 0 | 0 | 0 |
| Bacteroid | Phylum | 3920515 | 5513657 | 14.7 | 20.1 | 0.1 | 1.9 | 0.8 | 0.1 | 0.13 | 0.39 |
| Bacteroid | Class | 3470266 | 5084861 | 13 | 18.5 | 0.1 | 2.1 | 0.9 | 0.1 | 0.13 | 0.39 |
| Flavobac | Order | 2964310 | 4543849 | 11.1 | 16.5 | 0.2 | 2.3 | 1 | 0.1 | 0.13 | 0.39 |
| Flavoba | Family | 2809329 | 4390067 | 10.6 | 16 | 0.2 | 2.3 | 0.8 | 0.1 | 0.1 | 0.39 |
| Flavob | Genus | 2481809 | 4053492 | 9.3 | 14.8 | 0 | 2.4 | 0.9 | 0 | 0.09 | 0.36 |
| Aequo | Genus | 163471.5 | 162558.8 | 0.6 | 0.6 | 3 | 1.9 | 0 | 0 | 0.02 | 0.01 |
| Flavob | Genus | 77159.9 | 85576 | 0.3 | 0.3 | 0 | 0 | 0 | 3.9 | 0 | 0.01 |
| Vitellib | Genus | 73606.9 | 85481.9 | 0.3 | 0.3 | 0 | 0 | 0 | 0 | 0 | 0 |
| Subsa | Genus | 7215.5 | 0 | 0 | 0 | 0 | 0 | 0 | 0 | 0 | 0 |
| Gilviba | Genus | 3932.6 | 2552.8 | 0 | 0 | 0 | 0 | 0 | 0 | 0 | 0 |
| Flavivi | Genus | 1553.8 | 0 | 0 | 0 | 0 | 0 | 0 | 0 | 0 | 0 |
| Salinin | Genus | 580.2 | 405.7 | 0 | 0 | 0 | 0 | 0 | 0 | 0 | 0 |

| | | | | | | | | | | | |
|---|---|---|---|---|---|---|---|---|---|---|---|
| Weekse | Family | 105779.3 | 105522.9 | 0.4 | 0.4 | 0 | 0 | 6.5 | 0 | 0.03 | 0 |
| Chryse | Genus | 105779.3 | 105522.9 | 0.4 | 0.4 | 0 | 0 | 6.5 | 0 | 0.03 | 0 |
| NS9 ma | Family | 48644 | 48258.3 | 0.2 | 0.2 | 0 | 4.7 | 0 | 0 | 0 | 0.01 |
| NS9 m | Genus | 48644 | 48258.3 | 0.2 | 0.2 | 0 | 4.7 | 0 | 0 | 0 | 0.01 |
| Crocini | Family | 557.9 | 0 | 0 | 0 | 0 | 0 | 0 | 0 | 0 | 0 |
| Crocin | Genus | 557.9 | 0 | 0 | 0 | 0 | 0 | 0 | 0 | 0 | 0 |
| Cytopha | Order | 210176 | 188302.8 | 0.8 | 0.7 | 0 | 0 | 0 | 0 | 0 | 0 |
| Microsc | Family | 205383.4 | 182630.8 | 0.8 | 0.7 | 0 | 0 | 0 | 0 | 0 | 0 |
| Micros | Genus | 113638.4 | 94322.7 | 0.4 | 0.3 | 0 | 0 | 0 | 0 | 0 | 0 |
| Ohtael | Genus | 73820.9 | 68411.2 | 0.3 | 0.2 | 0 | 0 | 0 | 0 | 0 | 0 |
| Chryse | Genus | 17924.2 | 19642.5 | 0.1 | 0.1 | 0 | 0 | 0 | 0 | 0 | 0 |
| Hassal | Genus | 0 | 254.3 | 0 | 0 | 0 | 0 | 0 | 0 | 0 | 0 |
| Hymen | Family | 4097.6 | 3617.3 | 0 | 0 | 0 | 0 | 0 | 0 | 0 | 0 |
| Adhae | Genus | 4097.6 | 3617.3 | 0 | 0 | 0 | 0 | 0 | 0 | 0 | 0 |
| Cytoph | Family | 695 | 2054.8 | 0 | 0 | 0 | 0 | 0 | 0 | 0 | 0 |
| Cytoph | Genus | 695 | 2054.8 | 0 | 0 | 0 | 0 | 0 | 0 | 0 | 0 |
| Chitinop | Order | 174591 | 229299.9 | 0.7 | 0.8 | 0 | 0 | 0 | 0 | 0 | 0 |
| Saprosp | Family | 169710.4 | 223577.5 | 0.6 | 0.8 | 0 | 0 | 0 | 0 | 0 | 0 |
| Sapros | Genus | 169094.4 | 223259.3 | 0.6 | 0.8 | 0 | 0 | 0 | 0 | 0 | 0 |
| Phaeo | Genus | 616 | 318.1 | 0 | 0 | 0 | 0 | 0 | 0 | 0 | 0 |
| Chitino | Family | 2735.6 | 1881.5 | 0 | 0 | 0 | 0 | 0 | 0 | 0 | 0 |
| 37-13 | Family | 2145.1 | 3841 | 0 | 0 | 0 | 0 | 0 | 0 | 0 | 0 |
| 37-13_ | Genus | 2145.1 | 3841 | 0 | 0 | 0 | 0 | 0 | 0 | 0 | 0 |
| Sphingol | Order | 114994.8 | 118504.5 | 0.4 | 0.4 | 0 | 0 | 0 | 0 | 0 | 0 |
| KD3-93 | Family | 73232.8 | 71623.1 | 0.3 | 0.3 | 0 | 0 | 0 | 0 | 0 | 0 |
| KD3-9 | Genus | 73232.8 | 71623.1 | 0.3 | 0.3 | 0 | 0 | 0 | 0 | 0 | 0 |
| env.OP | Family | 22386.1 | 20954.1 | 0.1 | 0.1 | 0 | 0 | 0 | 0 | 0 | 0 |
| env.OI | Genus | 22386.1 | 20954.1 | 0.1 | 0.1 | 0 | 0 | 0 | 0 | 0 | 0 |
| AKYH76 | Family | 14672.2 | 16305.8 | 0.1 | 0.1 | 0 | 0 | 0 | 0 | 0 | 0 |
| AKYH7 | Genus | 14672.2 | 16305.8 | 0.1 | 0.1 | 0 | 0 | 0 | 0 | 0 | 0 |
| Sphingo | Family | 2409.6 | 6483.3 | 0 | 0 | 0 | 0 | 0 | 0 | 0 | 0 |

| | | | | | | | | | | | |
|---|---|---|---|---|---|---|---|---|---|---|---|
| Sphing | Genus | 2409.6 | 6483.3 | 0 | 0 | 0 | 0 | 0 | 0 | 0 | 0 |
| LiUU-11 | Family | 2070.9 | 2423.8 | 0 | 0 | 0 | 0 | 0 | 0 | 0 | 0 |
| LiUU-1 | Genus | 2070.9 | 2423.8 | 0 | 0 | 0 | 0 | 0 | 0 | 0 | 0 |
| NS11-1 | Family | 223.2 | 714.4 | 0 | 0 | 0 | 0 | 0 | 0 | 0 | 0 |
| NS11-1 | Genus | 223.2 | 714.4 | 0 | 0 | 0 | 0 | 0 | 0 | 0 | 0 |
| Bacteroi | Order | 6193.4 | 4905.3 | 0 | 0 | 0 | 0 | 0 | 0 | 0 | 0 |
| Bactero | Family | 6193.4 | 4905.3 | 0 | 0 | 0 | 0 | 0 | 0 | 0 | 0 |
| Ignavibac | Class | 350517.8 | 328794.1 | 1.3 | 1.2 | 0 | 0 | 0 | 0 | 0 | 0 |
| SJA-28 | Order | 350517.8 | 328794.1 | 1.3 | 1.2 | 0 | 0 | 0 | 0 | 0 | 0 |
| SJA-28_ | Family | 350517.8 | 328794.1 | 1.3 | 1.2 | 0 | 0 | 0 | 0 | 0 | 0 |
| Kapabact | Class | 70085.1 | 69282.8 | 0.3 | 0.3 | 0 | 0 | 0 | 0 | 0 | 0 |
| Kapabac | Order | 70085.1 | 69282.8 | 0.3 | 0.3 | 0 | 0 | 0 | 0 | 0 | 0 |
| Kapaba | Family | 70085.1 | 69282.8 | 0.3 | 0.3 | 0 | 0 | 0 | 0 | 0 | 0 |
| Rhodothe | Class | 29646.2 | 30719.4 | 0.1 | 0.1 | 0 | 0 | 0 | 0 | 0 | 0 |
| Rhodoth | Order | 29646.2 | 30719.4 | 0.1 | 0.1 | 0 | 0 | 0 | 0 | 0 | 0 |
| Rhodotl | Family | 29646.2 | 30719.4 | 0.1 | 0.1 | 0 | 0 | 0 | 0 | 0 | 0 |
| Rhodo | Genus | 29646.2 | 30719.4 | 0.1 | 0.1 | 0 | 0 | 0 | 0 | 0 | 0 |
| Actinobac | Phylum | 3767462 | 3384279 | 14.2 | 12.3 | 0 | 0.1 | 0 | 0 | 0 | 0.01 |
| Actinoba | Class | 2820423 | 2410786 | 10.6 | 8.8 | 0 | 0 | 0 | 0 | 0 | 0 |
| Micromo | Order | 1264055 | 874058.7 | 4.7 | 3.2 | 0 | 0 | 0 | 0 | 0 | 0 |
| Microm | Family | 1264055 | 874058.7 | 4.7 | 3.2 | 0 | 0 | 0 | 0 | 0 | 0 |
| Lueder | Genus | 521435.5 | 364728.6 | 2 | 1.3 | 0 | 0 | 0 | 0 | 0 | 0 |
| Micron | Genus | 285177.1 | 189941.4 | 1.1 | 0.7 | 0 | 0 | 0 | 0 | 0 | 0 |
| Micron | Genus | 257114.5 | 204097.3 | 1 | 0.7 | 0 | 0 | 0 | 0 | 0 | 0 |
| Planta | Genus | 100008.8 | 59869.8 | 0.4 | 0.2 | 0 | 0 | 0 | 0 | 0 | 0 |
| Phytoh | Genus | 40679.5 | 21603.7 | 0.2 | 0.1 | 0 | 0 | 0 | 0 | 0 | 0 |
| Pilimel | Genus | 21945.5 | 18500 | 0.1 | 0.1 | 0 | 0 | 0 | 0 | 0 | 0 |
| Asano | Genus | 15737.8 | 11106.3 | 0.1 | 0 | 0 | 0 | 0 | 0 | 0 | 0 |
| Spirilli | Genus | 15058.8 | 129.8 | 0.1 | 0 | 0 | 0 | 0 | 0 | 0 | 0 |
| Planos | Genus | 2130.7 | 893.7 | 0 | 0 | 0 | 0 | 0 | 0 | 0 | 0 |
| Dactyl | Genus | 2108.2 | 982.7 | 0 | 0 | 0 | 0 | 0 | 0 | 0 | 0 |

| | | | | | | | | | | | |
|---|---|---|---|---|---|---|---|---|---|---|---|
| Phyton | Genus | 1135.1 | 879.7 | 0 | 0 | 0 | 0 | 0 | 0 | 0 | 0 |
| Actino | Genus | 1019.4 | 331.3 | 0 | 0 | 0 | 0 | 0 | 0 | 0 | 0 |
| Stacke | Genus | 339.3 | 909.9 | 0 | 0 | 0 | 0 | 0 | 0 | 0 | 0 |
| Catella | Genus | 164.3 | 84.5 | 0 | 0 | 0 | 0 | 0 | 0 | 0 | 0 |
| Propioni | Order | 349525.3 | 307739.2 | 1.3 | 1.1 | 0 | 0 | 0 | 0 | 0 | 0 |
| Nocard | Family | 315060.8 | 279720 | 1.2 | 1 | 0 | 0 | 0 | 0 | 0 | 0 |
| Kribbe | Genus | 201546 | 166805.1 | 0.8 | 0.6 | 0 | 0 | 0 | 0 | 0 | 0 |
| Nocar | Genus | 89850.3 | 90842 | 0.3 | 0.3 | 0 | 0 | 0 | 0 | 0 | 0 |
| Aerom | Genus | 21303.3 | 19382.7 | 0.1 | 0.1 | 0 | 0 | 0 | 0 | 0 | 0 |
| Nocar | Genus | 2029 | 1506.7 | 0 | 0 | 0 | 0 | 0 | 0 | 0 | 0 |
| Marmo | Genus | 332.2 | 1183.4 | 0 | 0 | 0 | 0 | 0 | 0 | 0 | 0 |
| Propion | Family | 34464.5 | 28019.2 | 0.1 | 0.1 | 0 | 0 | 0 | 0 | 0 | 0 |
| Jiangel | Genus | 24585.1 | 20206.1 | 0.1 | 0.1 | 0 | 0 | 0 | 0 | 0 | 0 |
| Haloac | Genus | 6896.6 | 5809.6 | 0 | 0 | 0 | 0 | 0 | 0 | 0 | 0 |
| Propio | Genus | 2422.4 | 1919.2 | 0 | 0 | 0 | 0 | 0 | 0 | 0 | 0 |
| Microl | Genus | 560.3 | 84.3 | 0 | 0 | 0 | 0 | 0 | 0 | 0 | 0 |
| Streptos | Order | 325258.6 | 276067.8 | 1.2 | 1 | 0 | 0 | 0 | 0 | 0 | 0 |
| Strepto | Family | 206504.7 | 175377.7 | 0.8 | 0.6 | 0 | 0 | 0 | 0 | 0 | 0 |
| Nonon | Genus | 146647.5 | 131392 | 0.6 | 0.5 | 0 | 0 | 0 | 0 | 0 | 0 |
| Sphae | Genus | 29229.7 | 16283.7 | 0.1 | 0.1 | 0 | 0 | 0 | 0 | 0 | 0 |
| Strept | Genus | 25705.4 | 23208.5 | 0.1 | 0.1 | 0 | 0 | 0 | 0 | 0 | 0 |
| Therm | Genus | 2503.9 | 2931 | 0 | 0 | 0 | 0 | 0 | 0 | 0 | 0 |
| Strept | Genus | 2418.2 | 1562.5 | 0 | 0 | 0 | 0 | 0 | 0 | 0 | 0 |
| Thermo | Family | 117237.4 | 98931.2 | 0.4 | 0.4 | 0 | 0 | 0 | 0 | 0 | 0 |
| Actino | Genus | 79204.9 | 66549.5 | 0.3 | 0.2 | 0 | 0 | 0 | 0 | 0 | 0 |
| Therm | Genus | 25859.4 | 22653.9 | 0.1 | 0.1 | 0 | 0 | 0 | 0 | 0 | 0 |
| Therm | Genus | 12061.6 | 9727.8 | 0 | 0 | 0 | 0 | 0 | 0 | 0 | 0 |
| Therm | Genus | 111.6 | 0 | 0 | 0 | 0 | 0 | 0 | 0 | 0 | 0 |
| Nocard | Family | 1516.5 | 1758.9 | 0 | 0 | 0 | 0 | 0 | 0 | 0 | 0 |
| Therm | Genus | 1516.5 | 1758.9 | 0 | 0 | 0 | 0 | 0 | 0 | 0 | 0 |
| Strepton | Order | 309876.7 | 361946.6 | 1.2 | 1.3 | 0 | 0 | 0 | 0 | 0 | 0 |

| | | | | | | | | | | | |
|---|---|---|---|---|---|---|---|---|---|---|---|
| Strepto | Family | 309876.7 | 361946.6 | 1.2 | 1.3 | 0 | 0 | 0 | 0 | 0 | 0 |
| Strepto | Genus | 309876.7 | 361946.6 | 1.2 | 1.3 | 0 | 0 | 0 | 0 | 0 | 0 |
| Pseudon | Order | 237425.1 | 207128.4 | 0.9 | 0.8 | 0 | 0 | 0 | 0 | 0 | 0 |
| Pseudo | Family | 237425.1 | 207128.4 | 0.9 | 0.8 | 0 | 0 | 0 | 0 | 0 | 0 |
| Pseudo | Genus | 220759.6 | 191348.2 | 0.8 | 0.7 | 0 | 0 | 0 | 0 | 0 | 0 |
| Actino | Genus | 12972.9 | 11100.8 | 0 | 0 | 0 | 0 | 0 | 0 | 0 | 0 |
| Crossi | Genus | 3692.6 | 4263.6 | 0 | 0 | 0 | 0 | 0 | 0 | 0 | 0 |
| Saccha | Genus | 0 | 415.7 | 0 | 0 | 0 | 0 | 0 | 0 | 0 | 0 |
| Micrococ | Order | 226098.6 | 274703.6 | 0.8 | 1 | 0 | 0 | 0 | 0 | 0 | 0 |
| Microba | Family | 149039.5 | 183047.5 | 0.6 | 0.7 | 0 | 0 | 0 | 0 | 0 | 0 |
| Lysinir | Genus | 65017.3 | 74016.1 | 0.2 | 0.3 | 0 | 0 | 0 | 0 | 0 | 0 |
| Agrom | Genus | 33030 | 40393.6 | 0.1 | 0.1 | 0 | 0 | 0 | 0 | 0 | 0 |
| Homos | Genus | 17132.4 | 21294.5 | 0.1 | 0.1 | 0 | 0 | 0 | 0 | 0 | 0 |
| Microb | Genus | 9500.9 | 11632.7 | 0 | 0 | 0 | 0 | 0 | 0 | 0 | 0 |
| Parafri | Genus | 6088 | 11716.4 | 0 | 0 | 0 | 0 | 0 | 0 | 0 | 0 |
| Leifsor | Genus | 5619.7 | 8526 | 0 | 0 | 0 | 0 | 0 | 0 | 0 | 0 |
| Microb | Genus | 4430 | 6445.3 | 0 | 0 | 0 | 0 | 0 | 0 | 0 | 0 |
| Galbita | Genus | 4336.6 | 4771.5 | 0 | 0 | 0 | 0 | 0 | 0 | 0 | 0 |
| Salinib | Genus | 2218.9 | 2353.3 | 0 | 0 | 0 | 0 | 0 | 0 | 0 | 0 |
| Conyzi | Genus | 753 | 1331.1 | 0 | 0 | 0 | 0 | 0 | 0 | 0 | 0 |
| Glaciib | Genus | 463.2 | 254.3 | 0 | 0 | 0 | 0 | 0 | 0 | 0 | 0 |
| Schum | Genus | 449.6 | 312.6 | 0 | 0 | 0 | 0 | 0 | 0 | 0 | 0 |
| Micrococ | Family | 53230.7 | 77198.3 | 0.2 | 0.3 | 0 | 0 | 0 | 0 | 0 | 0 |
| Paenai | Genus | 38488.1 | 30316.4 | 0.1 | 0.1 | 0 | 0 | 0 | 0 | 0 | 0 |
| Pseuda | Genus | 6940.2 | 16128.7 | 0 | 0.1 | 0 | 0 | 0 | 0 | 0 | 0 |
| Arthrol | Genus | 4045.7 | 23109.5 | 0 | 0.1 | 0 | 0 | 0 | 0 | 0 | 0 |
| Glutan | Genus | 3756.7 | 5688 | 0 | 0 | 0 | 0 | 0 | 0 | 0 | 0 |
| Microc | Genus | 0 | 1955.7 | 0 | 0 | 0 | 0 | 0 | 0 | 0 | 0 |
| Promici | Family | 19021.8 | 9337.8 | 0.1 | 0 | 0 | 0 | 0 | 0 | 0 | 0 |
| Promic | Genus | 18689 | 8922.1 | 0.1 | 0 | 0 | 0 | 0 | 0 | 0 | 0 |
| Cellulo | Genus | 332.8 | 415.7 | 0 | 0 | 0 | 0 | 0 | 0 | 0 | 0 |

| | | | | | | | | | | | |
|---|---|---|---|---|---|---|---|---|---|---|---|
| Intraspo | Family | 4170.3 | 3900.4 | 0 | 0 | 0 | 0 | 0 | 0 | 0 | 0 |
| Terrab | Genus | 1837.3 | 1001.3 | 0 | 0 | 0 | 0 | 0 | 0 | 0 | 0 |
| Lapillic | Genus | 1389.9 | 1329.4 | 0 | 0 | 0 | 0 | 0 | 0 | 0 | 0 |
| Oryzihu | Genus | 603.8 | 402.8 | 0 | 0 | 0 | 0 | 0 | 0 | 0 | 0 |
| Knoelli | Genus | 339.4 | 829.1 | 0 | 0 | 0 | 0 | 0 | 0 | 0 | 0 |
| Intrasp | Genus | 0 | 337.9 | 0 | 0 | 0 | 0 | 0 | 0 | 0 | 0 |
| Demeq | Family | 636.3 | 1054 | 0 | 0 | 0 | 0 | 0 | 0 | 0 | 0 |
| Demec | Genus | 636.3 | 1054 | 0 | 0 | 0 | 0 | 0 | 0 | 0 | 0 |
| Cellulo | Family | 0 | 165.7 | 0 | 0 | 0 | 0 | 0 | 0 | 0 | 0 |
| Oersko | Genus | 0 | 165.7 | 0 | 0 | 0 | 0 | 0 | 0 | 0 | 0 |
| Frankial | Order | 78618.8 | 76191.8 | 0.3 | 0.3 | 0 | 0 | 0 | 0 | 0 | 0 |
| Geoder | Family | 63861.1 | 52962.6 | 0.2 | 0.2 | 0 | 0 | 0 | 0 | 0 | 0 |
| Blasto | Genus | 60397.1 | 49113.5 | 0.2 | 0.2 | 0 | 0 | 0 | 0 | 0 | 0 |
| Geode | Genus | 3463.9 | 3849 | 0 | 0 | 0 | 0 | 0 | 0 | 0 | 0 |
| Acidoth | Family | 6213.1 | 11351.1 | 0 | 0 | 0 | 0 | 0 | 0 | 0 | 0 |
| Acidot | Genus | 6213.1 | 11351.1 | 0 | 0 | 0 | 0 | 0 | 0 | 0 | 0 |
| Frankia | Family | 4706.6 | 3927.5 | 0 | 0 | 0 | 0 | 0 | 0 | 0 | 0 |
| Jatroph | Genus | 4706.6 | 3927.5 | 0 | 0 | 0 | 0 | 0 | 0 | 0 | 0 |
| Sporich | Family | 3838.1 | 7950.7 | 0 | 0 | 0 | 0 | 0 | 0 | 0 | 0 |
| Sporicl | Genus | 3838.1 | 7950.7 | 0 | 0 | 0 | 0 | 0 | 0 | 0 | 0 |
| Coryneb | Order | 25730.6 | 31081.4 | 0.1 | 0.1 | 0 | 0 | 0 | 0 | 0 | 0 |
| Mycoba | Family | 21823.9 | 26263.4 | 0.1 | 0.1 | 0 | 0 | 0 | 0 | 0 | 0 |
| Mycob | Genus | 21823.9 | 26263.4 | 0.1 | 0.1 | 0 | 0 | 0 | 0 | 0 | 0 |
| Nocard | Family | 3906.8 | 4818 | 0 | 0 | 0 | 0 | 0 | 0 | 0 | 0 |
| Nocard | Genus | 3244.5 | 4490 | 0 | 0 | 0 | 0 | 0 | 0 | 0 | 0 |
| Rhodo | Genus | 662.3 | 328 | 0 | 0 | 0 | 0 | 0 | 0 | 0 | 0 |
| Glycomy | Order | 3834.7 | 1868.9 | 0 | 0 | 0 | 0 | 0 | 0 | 0 | 0 |
| Glycom | Family | 3834.7 | 1868.9 | 0 | 0 | 0 | 0 | 0 | 0 | 0 | 0 |
| Glycor | Genus | 3834.7 | 1868.9 | 0 | 0 | 0 | 0 | 0 | 0 | 0 | 0 |
| Thermole | Class | 478417.8 | 462360.5 | 1.8 | 1.7 | 0 | 0.4 | 0 | 0 | 0 | 0.01 |
| Solirubro | Order | 422224.6 | 408657.6 | 1.6 | 1.5 | 0 | 0 | 0 | 0 | 0 | 0 |

| | | | | | | | | | | | |
|---|---|---|---|---|---|---|---|---|---|---|---|
| 67-14 | Family | 261542.6 | 270740.8 | 1 | 1 | 0 | 0 | 0 | 0 | 0 | 0 |
| 67-14_ | Genus | 261542.6 | 270740.8 | 1 | 1 | 0 | 0 | 0 | 0 | 0 | 0 |
| Solirubı | Family | 160682 | 137916.8 | 0.6 | 0.5 | 0 | 0 | 0 | 0 | 0 | 0 |
| Solirut | Genus | 110108.7 | 99778.9 | 0.4 | 0.4 | 0 | 0 | 0 | 0 | 0 | 0 |
| Conexi | Genus | 27493.7 | 22523.6 | 0.1 | 0.1 | 0 | 0 | 0 | 0 | 0 | 0 |
| JCM 18 | Genus | 15490 | 10004.1 | 0.1 | 0 | 0 | 0 | 0 | 0 | 0 | 0 |
| Solirut | Genus | 7589.6 | 5610.2 | 0 | 0 | 0 | 0 | 0 | 0 | 0 | 0 |
| Gaiellale | Order | 56193.2 | 53702.9 | 0.2 | 0.2 | 0 | 3.7 | 0 | 0 | 0 | 0.01 |
| Gaiellac | Family | 38885.8 | 36628 | 0.1 | 0.1 | 0 | 5.5 | 0 | 0 | 0 | 0.01 |
| Gaiella | Genus | 38885.8 | 36628 | 0.1 | 0.1 | 0 | 5.5 | 0 | 0 | 0 | 0.01 |
| Gaiellal | Family | 17307.4 | 17075 | 0.1 | 0.1 | 0 | 0 | 0 | 0 | 0 | 0 |
| Acidimicr | Class | 465486.6 | 507369.3 | 1.7 | 1.8 | 0 | 0 | 0 | 0 | 0 | 0 |
| Microtric | Order | 245324.8 | 287936.4 | 0.9 | 1 | 0 | 0 | 0 | 0 | 0 | 0 |
| Iamiace | Family | 147550.7 | 188508.1 | 0.6 | 0.7 | 0 | 0 | 0 | 0 | 0 | 0 |
| Iamia | Genus | 147550.7 | 188508.1 | 0.6 | 0.7 | 0 | 0 | 0 | 0 | 0 | 0 |
| Ilumato | Family | 55995.2 | 53079.3 | 0.2 | 0.2 | 0 | 0 | 0 | 0 | 0 | 0 |
| Ilumat | Genus | 30448.8 | 29932.7 | 0.1 | 0.1 | 0 | 0 | 0 | 0 | 0 | 0 |
| Ilumat | Genus | 25042.1 | 22121 | 0.1 | 0.1 | 0 | 0 | 0 | 0 | 0 | 0 |
| CL500 | Genus | 504.3 | 1025.6 | 0 | 0 | 0 | 0 | 0 | 0 | 0 | 0 |
| Microtri | Family | 37144.7 | 38823.3 | 0.1 | 0.1 | 0 | 0 | 0 | 0 | 0 | 0 |
| Microtri | Family | 4634.1 | 7525.7 | 0 | 0 | 0 | 0 | 0 | 0 | 0 | 0 |
| Microti | Genus | 3618.7 | 3639.7 | 0 | 0 | 0 | 0 | 0 | 0 | 0 | 0 |
| IMCC2 | Genus | 1015.4 | 3885.9 | 0 | 0 | 0 | 0 | 0 | 0 | 0 | 0 |
| IMCC26 | Order | 204133.8 | 204904.5 | 0.8 | 0.7 | 0 | 0 | 0 | 0 | 0 | 0 |
| IMCC26 | Family | 204133.8 | 204904.5 | 0.8 | 0.7 | 0 | 0 | 0 | 0 | 0 | 0 |
| Acidimic | Order | 16028.1 | 14528.4 | 0.1 | 0.1 | 0 | 0 | 0 | 0 | 0 | 0 |
| MB-A2-10 | Class | 3134.1 | 3763 | 0 | 0 | 0 | 0 | 0 | 0 | 0 | 0 |
| MB-A2-1 | Order | 3134.1 | 3763 | 0 | 0 | 0 | 0 | 0 | 0 | 0 | 0 |
| Gemmatin | Phylum | 1575450 | 1484515 | 5.9 | 5.4 | 0.2 | 0 | 0 | 0 | 0.01 | 0 |
| Gemmati | Class | 1177394 | 1117853 | 4.4 | 4.1 | 0 | 0 | 0 | 0 | 0 | 0 |
| Gemmat | Order | 1177394 | 1117853 | 4.4 | 4.1 | 0 | 0 | 0 | 0 | 0 | 0 |

| | | | | | | | | | | | |
|---|---|---|---|---|---|---|---|---|---|---|---|
| Gemma | Family | 1177394 | 1117853 | 4.4 | 4.1 | 0 | 0 | 0 | 0 | 0 | 0 |
| Gemm | Genus | 1047073 | 1003420 | 3.9 | 3.7 | 0 | 0 | 0 | 0 | 0 | 0 |
| Gemm | Genus | 130320.8 | 114433.2 | 0.5 | 0.4 | 0 | 0 | 0 | 0 | 0 | 0 |
| Longimic | Class | 334646.1 | 320806 | 1.3 | 1.2 | 0 | 0 | 0 | 0 | 0 | 0 |
| Longimic | Order | 334646.1 | 320806 | 1.3 | 1.2 | 0 | 0 | 0 | 0 | 0 | 0 |
| Longim | Family | 334646.1 | 320806 | 1.3 | 1.2 | 0 | 0 | 0 | 0 | 0 | 0 |
| Longin | Genus | 334646.1 | 320806 | 1.3 | 1.2 | 0 | 0 | 0 | 0 | 0 | 0 |
| S0134 ter | Class | 34140.7 | 20981.3 | 0.1 | 0.1 | 9.9 | 0 | 0 | 0 | 0.01 | 0 |
| S0134 te | Order | 34140.7 | 20981.3 | 0.1 | 0.1 | 9.9 | 0 | 0 | 0 | 0.01 | 0 |
| BD2-11 te | Class | 18851.6 | 15855 | 0.1 | 0.1 | 0 | 0 | 0 | 0 | 0 | 0 |
| BD2-11 t | Order | 18851.6 | 15855 | 0.1 | 0.1 | 0 | 0 | 0 | 0 | 0 | 0 |
| AKAU404 | Class | 7482.4 | 7083.1 | 0 | 0 | 0 | 0 | 0 | 0 | 0 | 0 |
| AKAU40 | Order | 7482.4 | 7083.1 | 0 | 0 | 0 | 0 | 0 | 0 | 0 | 0 |
| Gemmati | Class | 2934.8 | 1937 | 0 | 0 | 0 | 0 | 0 | 0 | 0 | 0 |
| Myxococc | Phylum | 276571.2 | 252868.1 | 1 | 0.9 | 1.1 | 0 | 0 | 0 | 0.01 | 0 |
| Polyangia | Class | 189486.3 | 150998 | 0.7 | 0.5 | 0 | 0 | 0 | 0 | 0 | 0 |
| Polyangi | Order | 109283.3 | 88969 | 0.4 | 0.3 | 0 | 0 | 0 | 0 | 0 | 0 |
| Blrii41 | Family | 80233.5 | 58558.7 | 0.3 | 0.2 | 0 | 0 | 0 | 0 | 0 | 0 |
| Blrii41_ | Genus | 80233.5 | 58558.7 | 0.3 | 0.2 | 0 | 0 | 0 | 0 | 0 | 0 |
| Polyang | Family | 12156.4 | 13328.3 | 0 | 0 | 0 | 0 | 0 | 0 | 0 | 0 |
| Pajaro | Genus | 11991.1 | 13328.3 | 0 | 0 | 0 | 0 | 0 | 0 | 0 | 0 |
| Polyan | Genus | 165.3 | 0 | 0 | 0 | 0 | 0 | 0 | 0 | 0 | 0 |
| Phaseli | Family | 12008.7 | 11417.2 | 0 | 0 | 0 | 0 | 0 | 0 | 0 | 0 |
| Phasel | Genus | 12008.7 | 11417.2 | 0 | 0 | 0 | 0 | 0 | 0 | 0 | 0 |
| Sandara | Family | 3279.2 | 4258.3 | 0 | 0 | 0 | 0 | 0 | 0 | 0 | 0 |
| Sandar | Genus | 2817.8 | 2673.8 | 0 | 0 | 0 | 0 | 0 | 0 | 0 | 0 |
| Sandar | Genus | 461.4 | 1584.6 | 0 | 0 | 0 | 0 | 0 | 0 | 0 | 0 |
| Eel-36e | Family | 1605.4 | 1406.5 | 0 | 0 | 0 | 0 | 0 | 0 | 0 | 0 |
| Eel-36 | Genus | 1605.4 | 1406.5 | 0 | 0 | 0 | 0 | 0 | 0 | 0 | 0 |
| Haliangi | Order | 64883.1 | 51564.5 | 0.2 | 0.2 | 0 | 0 | 0 | 0 | 0 | 0 |
| Haliang | Family | 64883.1 | 51564.5 | 0.2 | 0.2 | 0 | 0 | 0 | 0 | 0 | 0 |

| | | | | | | | | | | | |
|---|---|---|---|---|---|---|---|---|---|---|---|
| Halian | Genus | 64883.1 | 51564.5 | 0.2 | 0.2 | 0 | 0 | 0 | 0 | 0 | 0 |
| Nannocy | Order | 9645.7 | 7779.9 | 0 | 0 | 0 | 0 | 0 | 0 | 0 | 0 |
| Nannoc | Family | 9645.7 | 7779.9 | 0 | 0 | 0 | 0 | 0 | 0 | 0 | 0 |
| Nanno | Genus | 6835.4 | 6611.1 | 0 | 0 | 0 | 0 | 0 | 0 | 0 | 0 |
| Pseud | Genus | 2810.2 | 1168.8 | 0 | 0 | 0 | 0 | 0 | 0 | 0 | 0 |
| Blfdi19 | Order | 5674.3 | 2684.7 | 0 | 0 | 0 | 0 | 0 | 0 | 0 | 0 |
| Blfdi19_ | Family | 5674.3 | 2684.7 | 0 | 0 | 0 | 0 | 0 | 0 | 0 | 0 |
| Myxococ | Class | 86035.7 | 99419.7 | 0.3 | 0.4 | 3.4 | 0 | 0 | 0 | 0.01 | 0 |
| Myxococ | Order | 86035.7 | 99419.7 | 0.3 | 0.4 | 3.4 | 0 | 0 | 0 | 0.01 | 0 |
| Vulgatik | Family | 58357.9 | 61525.7 | 0.2 | 0.2 | 0 | 0 | 0 | 0 | 0 | 0 |
| Vulgati | Genus | 58357.9 | 61525.7 | 0.2 | 0.2 | 0 | 0 | 0 | 0 | 0 | 0 |
| Myxoco | Family | 27677.8 | 37640.6 | 0.1 | 0.1 | 10.6 | 0 | 0 | 0 | 0.01 | 0 |
| Myxoc | Genus | 21633.6 | 19524.4 | 0.1 | 0.1 | 13.5 | 0 | 0 | 0 | 0.01 | 0 |
| Archar | Genus | 4300.4 | 14945.4 | 0 | 0.1 | 0 | 0 | 0 | 0 | 0 | 0 |
| Stigma | Genus | 762 | 1041.6 | 0 | 0 | 0 | 0 | 0 | 0 | 0 | 0 |
| Cystob | Genus | 524 | 1021.1 | 0 | 0 | 0 | 0 | 0 | 0 | 0 | 0 |
| P3OB- | Genus | 457.8 | 1108 | 0 | 0 | 0 | 0 | 0 | 0 | 0 | 0 |
| 27F-14 | Family | 0 | 253.4 | 0 | 0 | 0 | 0 | 0 | 0 | 0 | 0 |
| 27F-14 | Genus | 0 | 253.4 | 0 | 0 | 0 | 0 | 0 | 0 | 0 | 0 |
| bacteriap | Class | 1049.2 | 2450.4 | 0 | 0 | 0 | 0 | 0 | 0 | 0 | 0 |
| bacteria | Order | 1049.2 | 2450.4 | 0 | 0 | 0 | 0 | 0 | 0 | 0 | 0 |
| Acidobact | Phylum | 258542.1 | 246357.5 | 1 | 0.9 | 0 | 0 | 0 | 0 | 0 | 0 |
| Acidobac | Class | 217481.8 | 200035.7 | 0.8 | 0.7 | 0 | 0 | 0 | 0 | 0 | 0 |
| PAUC26 | Order | 189756.2 | 177779.2 | 0.7 | 0.6 | 0 | 0 | 0 | 0 | 0 | 0 |
| PAUC2 | Family | 189756.2 | 177779.2 | 0.7 | 0.6 | 0 | 0 | 0 | 0 | 0 | 0 |
| Solibact | Order | 11408.7 | 8925.2 | 0 | 0 | 0 | 0 | 0 | 0 | 0 | 0 |
| Solibac | Family | 11408.7 | 8925.2 | 0 | 0 | 0 | 0 | 0 | 0 | 0 | 0 |
| Candic | Genus | 11408.7 | 8925.2 | 0 | 0 | 0 | 0 | 0 | 0 | 0 | 0 |
| Bryobact | Order | 11103.1 | 9994.9 | 0 | 0 | 0 | 0 | 0 | 0 | 0 | 0 |
| Bryobac | Family | 11103.1 | 9994.9 | 0 | 0 | 0 | 0 | 0 | 0 | 0 | 0 |
| Bryoba | Genus | 11103.1 | 9994.9 | 0 | 0 | 0 | 0 | 0 | 0 | 0 | 0 |

| | | | | | | | | | | | |
|---|---|---|---|---|---|---|---|---|---|---|---|
| Acidoba | Order | 4093.6 | 2115.2 | 0 | 0 | 0 | 0 | 0 | 0 | 0 | 0 |
| Acidoba | Family | 4093.6 | 2115.2 | 0 | 0 | 0 | 0 | 0 | 0 | 0 | 0 |
| Acidoba | Order | 1120.2 | 422.9 | 0 | 0 | 0 | 0 | 0 | 0 | 0 | 0 |
| Paludiba | Order | 0 | 798.3 | 0 | 0 | 0 | 0 | 0 | 0 | 0 | 0 |
| Paludib | Family | 0 | 798.3 | 0 | 0 | 0 | 0 | 0 | 0 | 0 | 0 |
| Thermoar | Class | 28134 | 36062.1 | 0.1 | 0.1 | 0 | 0 | 0 | 0 | 0 | 0 |
| Thermoa | Order | 28134 | 36062.1 | 0.1 | 0.1 | 0 | 0 | 0 | 0 | 0 | 0 |
| Thermo | Family | 28134 | 36062.1 | 0.1 | 0.1 | 0 | 0 | 0 | 0 | 0 | 0 |
| Subgrc | Genus | 28134 | 36062.1 | 0.1 | 0.1 | 0 | 0 | 0 | 0 | 0 | 0 |
| Subgroup | Class | 10146.5 | 6614.8 | 0 | 0 | 0 | 0 | 0 | 0 | 0 | 0 |
| Subgrou | Order | 10146.5 | 6614.8 | 0 | 0 | 0 | 0 | 0 | 0 | 0 | 0 |
| Holophag | Class | 2779.8 | 3644.9 | 0 | 0 | 0 | 0 | 0 | 0 | 0 | 0 |
| Subgrou | Order | 2779.8 | 3644.9 | 0 | 0 | 0 | 0 | 0 | 0 | 0 | 0 |
| Subgrou | Family | 2779.8 | 3644.9 | 0 | 0 | 0 | 0 | 0 | 0 | 0 | 0 |
| Chloroflex | Phylum | 200335.5 | 183133.4 | 0.8 | 0.7 | 0 | 0 | 0 | 0 | 0 | 0 |
| TK10 | Class | 68855.9 | 65188.9 | 0.3 | 0.2 | 0 | 0 | 0 | 0 | 0 | 0 |
| TK10_Cl | Order | 68855.9 | 65188.9 | 0.3 | 0.2 | 0 | 0 | 0 | 0 | 0 | 0 |
| JG30-KF-( | Class | 66502.8 | 60550.7 | 0.2 | 0.2 | 0 | 0 | 0 | 0 | 0 | 0 |
| JG30-KF- | Order | 66502.8 | 60550.7 | 0.2 | 0.2 | 0 | 0 | 0 | 0 | 0 | 0 |
| Chlorofle | Class | 48696.4 | 47825.7 | 0.2 | 0.2 | 0 | 0 | 0 | 0 | 0 | 0 |
| Chlorofl | Order | 47930.9 | 47244 | 0.2 | 0.2 | 0 | 0 | 0 | 0 | 0 | 0 |
| Herpetc | Family | 43208.9 | 44665.2 | 0.2 | 0.2 | 0 | 0 | 0 | 0 | 0 | 0 |
| Herpet | Genus | 43208.9 | 44665.2 | 0.2 | 0.2 | 0 | 0 | 0 | 0 | 0 | 0 |
| Roseifle | Family | 4722 | 2578.7 | 0 | 0 | 0 | 0 | 0 | 0 | 0 | 0 |
| Roseifl | Genus | 4722 | 2578.7 | 0 | 0 | 0 | 0 | 0 | 0 | 0 | 0 |
| Kalloten | Order | 765.6 | 581.8 | 0 | 0 | 0 | 0 | 0 | 0 | 0 | 0 |
| AKIW78 | Family | 765.6 | 581.8 | 0 | 0 | 0 | 0 | 0 | 0 | 0 | 0 |
| AKIW7 | Genus | 765.6 | 581.8 | 0 | 0 | 0 | 0 | 0 | 0 | 0 | 0 |
| Dehaloco | Class | 15771.4 | 9061.9 | 0.1 | 0 | 0 | 0 | 0 | 0 | 0 | 0 |
| S085 | Order | 15771.4 | 9061.9 | 0.1 | 0 | 0 | 0 | 0 | 0 | 0 | 0 |
| S085_C | Family | 15771.4 | 9061.9 | 0.1 | 0 | 0 | 0 | 0 | 0 | 0 | 0 |

| | | | | | | | | | | | |
|---|---|---|---|---|---|---|---|---|---|---|---|
| Chlorofle | Class | 508.9 | 506.1 | 0 | 0 | 0 | 0 | 0 | 0 | 0 | 0 |
| Bdellovibri | Phylum | 178536.8 | 163309.3 | 0.7 | 0.6 | 1.3 | 1.3 | 0 | 0 | 0.01 | 0.01 |
| Bdellovib | Class | 113577.9 | 96511.2 | 0.4 | 0.4 | 2.1 | 0 | 0 | 0 | 0.01 | 0 |
| Bdellovil | Order | 98215.3 | 86928.6 | 0.4 | 0.3 | 2.4 | 0 | 0 | 0 | 0.01 | 0 |
| Bdellov | Family | 98215.3 | 86928.6 | 0.4 | 0.3 | 2.4 | 0 | 0 | 0 | 0.01 | 0 |
| OM27 | Genus | 51817.4 | 48227.5 | 0.2 | 0.2 | 0 | 0 | 0 | 0 | 0 | 0 |
| Bdello | Genus | 46397.9 | 38701.1 | 0.2 | 0.1 | 5.1 | 0 | 0 | 0 | 0.01 | 0 |
| Bacterio | Order | 15362.5 | 9582.7 | 0.1 | 0 | 0 | 0 | 0 | 0 | 0 | 0 |
| Bacteri | Family | 15362.5 | 9582.7 | 0.1 | 0 | 0 | 0 | 0 | 0 | 0 | 0 |
| Peredi | Genus | 12913.8 | 9582.7 | 0 | 0 | 0 | 0 | 0 | 0 | 0 | 0 |
| Bacter | Genus | 2448.8 | 0 | 0 | 0 | 0 | 0 | 0 | 0 | 0 | 0 |
| Oligoflexi | Class | 64958.9 | 66798.1 | 0.2 | 0.2 | 0 | 3.1 | 0 | 0 | 0 | 0.01 |
| 0319-6G | Order | 63040.6 | 65519.1 | 0.2 | 0.2 | 0 | 3.1 | 0 | 0 | 0 | 0.01 |
| 0319-6 | Family | 63040.6 | 65519.1 | 0.2 | 0.2 | 0 | 3.1 | 0 | 0 | 0 | 0.01 |
| Oligoflex | Order | 1918.3 | 1279 | 0 | 0 | 0 | 0 | 0 | 0 | 0 | 0 |
| Oligofle | Family | 1918.3 | 1279 | 0 | 0 | 0 | 0 | 0 | 0 | 0 | 0 |
| Oligofl | Genus | 1918.3 | 1279 | 0 | 0 | 0 | 0 | 0 | 0 | 0 | 0 |
| Verrucomi | Phylum | 116796.3 | 118756 | 0.4 | 0.4 | 0 | 0 | 0 | 0 | 0 | 0 |
| Verrucom | Class | 99878.8 | 102894.8 | 0.4 | 0.4 | 0 | 0 | 0 | 0 | 0 | 0 |
| Pedosph | Order | 99878.8 | 102894.8 | 0.4 | 0.4 | 0 | 0 | 0 | 0 | 0 | 0 |
| Pedosp | Family | 99878.8 | 102894.8 | 0.4 | 0.4 | 0 | 0 | 0 | 0 | 0 | 0 |
| Pedos | Genus | 95045.7 | 96332.3 | 0.4 | 0.4 | 0 | 0 | 0 | 0 | 0 | 0 |
| Pedos | Genus | 3553.5 | 2376.2 | 0 | 0 | 0 | 0 | 0 | 0 | 0 | 0 |
| Ellin51 | Genus | 835.7 | 902.6 | 0 | 0 | 0 | 0 | 0 | 0 | 0 | 0 |
| Oikopl | Genus | 339.4 | 1100.4 | 0 | 0 | 0 | 0 | 0 | 0 | 0 | 0 |
| Ellin51 | Genus | 104.6 | 1396.4 | 0 | 0 | 0 | 0 | 0 | 0 | 0 | 0 |
| DEV00 | Genus | 0 | 787 | 0 | 0 | 0 | 0 | 0 | 0 | 0 | 0 |
| Chlamydi | Class | 16917.6 | 15861.2 | 0.1 | 0.1 | 0 | 0 | 0 | 0 | 0 | 0 |
| Chlamyc | Order | 16917.6 | 15861.2 | 0.1 | 0.1 | 0 | 0 | 0 | 0 | 0 | 0 |
| cvE6 | Family | 14072.5 | 10317.1 | 0.1 | 0 | 0 | 0 | 0 | 0 | 0 | 0 |
| cvE6_F | Genus | 14072.5 | 10317.1 | 0.1 | 0 | 0 | 0 | 0 | 0 | 0 | 0 |

| | | | | | | | | | | | |
|---|---|---|---|---|---|---|---|---|---|---|---|
| Parachl | Family | 2409 | 4699.3 | 0 | 0 | 0 | 0 | 0 | 0 | 0 | 0 |
| Candic | Genus | 1900 | 2143.3 | 0 | 0 | 0 | 0 | 0 | 0 | 0 | 0 |
| Neoch | Genus | 397.4 | 1012.8 | 0 | 0 | 0 | 0 | 0 | 0 | 0 | 0 |
| Parach | Genus | 111.6 | 1543.2 | 0 | 0 | 0 | 0 | 0 | 0 | 0 | 0 |
| Simkan | Family | 436 | 844.7 | 0 | 0 | 0 | 0 | 0 | 0 | 0 | 0 |
| Simkai | Genus | 436 | 844.7 | 0 | 0 | 0 | 0 | 0 | 0 | 0 | 0 |
| Patesciba | Phylum | 56164.5 | 36527.8 | 0.2 | 0.1 | 0 | 0 | 0 | 0 | 0 | 0 |
| Sacchari | Class | 52973.4 | 32724.4 | 0.2 | 0.1 | 0 | 0 | 0 | 0 | 0 | 0 |
| Sacchari | Order | 52973.4 | 32724.4 | 0.2 | 0.1 | 0 | 0 | 0 | 0 | 0 | 0 |
| YM | Family | 43430.5 | 27335.4 | 0.2 | 0.1 | 0 | 0 | 0 | 0 | 0 | 0 |
| 50 | Genus | 43430.5 | 27335.4 | 0.2 | 0.1 | 0 | 0 | 0 | 0 | 0 | 0 |
| Saccha | Family | 9542.9 | 4823.7 | 0 | 0 | 0 | 0 | 0 | 0 | 0 | 0 |
| Saccha | Family | 0 | 565.3 | 0 | 0 | 0 | 0 | 0 | 0 | 0 | 0 |
| TM7a | Genus | 0 | 565.3 | 0 | 0 | 0 | 0 | 0 | 0 | 0 | 0 |
| Parcubac | Class | 2213.5 | 1790 | 0 | 0 | 0 | 0 | 0 | 0 | 0 | 0 |
| Candida | Order | 2213.5 | 1790 | 0 | 0 | 0 | 0 | 0 | 0 | 0 | 0 |
| Candid | Family | 2213.5 | 1790 | 0 | 0 | 0 | 0 | 0 | 0 | 0 | 0 |
| ABY1 | Class | 580.2 | 1688.9 | 0 | 0 | 0 | 0 | 0 | 0 | 0 | 0 |
| Candida | Order | 580.2 | 1688.9 | 0 | 0 | 0 | 0 | 0 | 0 | 0 | 0 |
| Candid | Family | 580.2 | 1688.9 | 0 | 0 | 0 | 0 | 0 | 0 | 0 | 0 |
| Patesciba | Class | 397.4 | 324.6 | 0 | 0 | 0 | 0 | 0 | 0 | 0 | 0 |
| Planctomy | Phylum | 48233.1 | 53314.4 | 0.2 | 0.2 | 0 | 0 | 3.4 | 0 | 0.01 | 0 |
| Phycisph | Class | 48039.7 | 52189.8 | 0.2 | 0.2 | 0 | 0 | 3.4 | 0 | 0.01 | 0 |
| Phycispl | Order | 48039.7 | 52189.8 | 0.2 | 0.2 | 0 | 0 | 3.4 | 0 | 0.01 | 0 |
| Phycisp | Family | 48039.7 | 52189.8 | 0.2 | 0.2 | 0 | 0 | 3.4 | 0 | 0.01 | 0 |
| SM1AC | Genus | 47765.4 | 51166 | 0.2 | 0.2 | 0 | 0 | 3.5 | 0 | 0.01 | 0 |
| Phycis | Genus | 274.2 | 1023.7 | 0 | 0 | 0 | 0 | 0 | 0 | 0 | 0 |
| OM190 | Class | 193.4 | 744 | 0 | 0 | 0 | 0 | 0 | 0 | 0 | 0 |
| OM190_ | Order | 193.4 | 744 | 0 | 0 | 0 | 0 | 0 | 0 | 0 | 0 |
| vadinHA4 | Class | 0 | 380.6 | 0 | 0 | 0 | 0 | 0 | 0 | 0 | 0 |
| vadinHA | Order | 0 | 380.6 | 0 | 0 | 0 | 0 | 0 | 0 | 0 | 0 |

| | | | | | | | | | | | |
|---|---|---|---|---|---|---|---|---|---|---|---|
| Armatimor | Phylum | 40614.9 | 29741.7 | 0.2 | 0.1 | 0 | 0 | 0 | 0 | 0 | 0 |
| Fimbriimo | Class | 40614.9 | 29741.7 | 0.2 | 0.1 | 0 | 0 | 0 | 0 | 0 | 0 |
| Fimbriim | Order | 40614.9 | 29741.7 | 0.2 | 0.1 | 0 | 0 | 0 | 0 | 0 | 0 |
| Fimbriir | Family | 40614.9 | 29741.7 | 0.2 | 0.1 | 0 | 0 | 0 | 0 | 0 | 0 |
| Fimbrii | Genus | 40614.9 | 29741.7 | 0.2 | 0.1 | 0 | 0 | 0 | 0 | 0 | 0 |
| Abditibact | Phylum | 34817.1 | 33989.7 | 0.1 | 0.1 | 0 | 15.2 | 0 | 0 | 0 | 0.02 |
| Abditibac | Class | 34817.1 | 33989.7 | 0.1 | 0.1 | 0 | 15.2 | 0 | 0 | 0 | 0.02 |
| Abditiba | Order | 34817.1 | 33989.7 | 0.1 | 0.1 | 0 | 15.2 | 0 | 0 | 0 | 0.02 |
| Abditiba | Family | 34817.1 | 33989.7 | 0.1 | 0.1 | 0 | 15.2 | 0 | 0 | 0 | 0.02 |
| Abditib | Genus | 34817.1 | 33989.7 | 0.1 | 0.1 | 0 | 15.2 | 0 | 0 | 0 | 0.02 |
| Desulfoba | Phylum | 12740 | 25176.4 | 0 | 0.1 | 0 | 0 | 0 | 0 | 0 | 0 |
| Desulfurc | Class | 12463.1 | 23688.4 | 0 | 0.1 | 0 | 0 | 0 | 0 | 0 | 0 |
| PB19 | Order | 12463.1 | 23688.4 | 0 | 0.1 | 0 | 0 | 0 | 0 | 0 | 0 |
| PB19_C | Family | 12463.1 | 23688.4 | 0 | 0.1 | 0 | 0 | 0 | 0 | 0 | 0 |
| Desulfoba | Class | 276.9 | 1488 | 0 | 0 | 0 | 0 | 0 | 0 | 0 | 0 |
| RCP2-54 | Phylum | 7205 | 5282.3 | 0 | 0 | 0 | 0 | 0 | 0 | 0 | 0 |
| RCP2-54_ | Class | 7205 | 5282.3 | 0 | 0 | 0 | 0 | 0 | 0 | 0 | 0 |
| Latesciba | Phylum | 6916.7 | 8401.6 | 0 | 0 | 0 | 42.6 | 0 | 0 | 0 | 0.01 |
| Latesciba | Class | 6916.7 | 8401.6 | 0 | 0 | 0 | 42.6 | 0 | 0 | 0 | 0.01 |
| Deinococc | Phylum | 6691 | 6793.6 | 0 | 0 | 0 | 0 | 0 | 0 | 0 | 0 |
| Deinococ | Class | 6691 | 6793.6 | 0 | 0 | 0 | 0 | 0 | 0 | 0 | 0 |
| Deinoco | Order | 6691 | 6793.6 | 0 | 0 | 0 | 0 | 0 | 0 | 0 | 0 |
| Trueper | Family | 4318.2 | 3952 | 0 | 0 | 0 | 0 | 0 | 0 | 0 | 0 |
| Truepe | Genus | 4318.2 | 3952 | 0 | 0 | 0 | 0 | 0 | 0 | 0 | 0 |
| Deinoco | Family | 2372.8 | 2841.6 | 0 | 0 | 0 | 0 | 0 | 0 | 0 | 0 |
| Deinoc | Genus | 2372.8 | 2841.6 | 0 | 0 | 0 | 0 | 0 | 0 | 0 | 0 |
| Dependen | Phylum | 5360.4 | 10558.6 | 0 | 0 | 0 | 0 | 0 | 0 | 0 | 0 |
| Babeliae | Class | 5360.4 | 10558.6 | 0 | 0 | 0 | 0 | 0 | 0 | 0 | 0 |
| Babelial | Order | 5360.4 | 10558.6 | 0 | 0 | 0 | 0 | 0 | 0 | 0 | 0 |
| Babelia | Family | 4595.9 | 9672.4 | 0 | 0 | 0 | 0 | 0 | 0 | 0 | 0 |
| UBA124 | Family | 659.9 | 886.2 | 0 | 0 | 0 | 0 | 0 | 0 | 0 | 0 |

| | | | | | | | | | | | |
|---|---|---|---|---|---|---|---|---|---|---|---|
| UBA12 | Genus | 659.9 | 886.2 | 0 | 0 | 0 | 0 | 0 | 0 | 0 | 0 |
| Vermipl | Family | 104.6 | 0 | 0 | 0 | 0 | 0 | 0 | 0 | 0 | 0 |
| Vermip | Genus | 104.6 | 0 | 0 | 0 | 0 | 0 | 0 | 0 | 0 | 0 |
| Nitrospiro | Phylum | 2114.9 | 1274.8 | 0 | 0 | 0 | 0 | 0 | 0 | 0 | 0 |
| Nitrospiri | Class | 2114.9 | 1274.8 | 0 | 0 | 0 | 0 | 0 | 0 | 0 | 0 |
| Nitrospir | Order | 2114.9 | 1274.8 | 0 | 0 | 0 | 0 | 0 | 0 | 0 | 0 |
| Nitrosp | Family | 2114.9 | 1274.8 | 0 | 0 | 0 | 0 | 0 | 0 | 0 | 0 |
| Nitros | Genus | 2114.9 | 1274.8 | 0 | 0 | 0 | 0 | 0 | 0 | 0 | 0 |
| Sumerlaec | Phylum | 1524.8 | 2306.7 | 0 | 0 | 0 | 0 | 0 | 0 | 0 | 0 |
| Sumerlae | Class | 1524.8 | 2306.7 | 0 | 0 | 0 | 0 | 0 | 0 | 0 | 0 |
| Sumerla | Order | 1524.8 | 2306.7 | 0 | 0 | 0 | 0 | 0 | 0 | 0 | 0 |
| Sumerl | Family | 1524.8 | 2306.7 | 0 | 0 | 0 | 0 | 0 | 0 | 0 | 0 |
| Sumer | Genus | 1524.8 | 2306.7 | 0 | 0 | 0 | 0 | 0 | 0 | 0 | 0 |
| Fibrobacte | Phylum | 1240.8 | 1155.9 | 0 | 0 | 0 | 0 | 0 | 0 | 0 | 0 |
| Fibrobact | Class | 1240.8 | 1155.9 | 0 | 0 | 0 | 0 | 0 | 0 | 0 | 0 |
| Fibrobac | Order | 1240.8 | 1155.9 | 0 | 0 | 0 | 0 | 0 | 0 | 0 | 0 |
| Fibroba | Family | 1240.8 | 1155.9 | 0 | 0 | 0 | 0 | 0 | 0 | 0 | 0 |
| Fibrob | Genus | 1240.8 | 1155.9 | 0 | 0 | 0 | 0 | 0 | 0 | 0 | 0 |
| MBNT15 | Phylum | 893.6 | 717.9 | 0 | 0 | 0 | 0 | 0 | 0 | 0 | 0 |
| MBNT15_ | Class | 893.6 | 717.9 | 0 | 0 | 0 | 0 | 0 | 0 | 0 | 0 |
| SAR324 cl | Phylum | 517.9 | 454.9 | 0 | 0 | 0 | 0 | 0 | 0 | 0 | 0 |
| SAR324 c | Class | 517.9 | 454.9 | 0 | 0 | 0 | 0 | 0 | 0 | 0 | 0 |
| FCPU426 | Phylum | 0 | 81.1 | 0 | 0 | 0 | 0 | 0 | 0 | 0 | 0 |
| FCPU426 | Class | 0 | 81.1 | 0 | 0 | 0 | 0 | 0 | 0 | 0 | 0 |
