## Supplemental Table S3 for "Potato foliar infection with *Phytophthora infestans* drives strong, cultivar-specific shifts in rhizosphere communities"

| group | rank | abund_B | abund_I | rel_abund_ | rel_abund_ | B_more | I_more | B_less | I_less | weighted_d | weighted_densI |
|---|---|---|---|---|---|---|---|---|---|---|---|
| Bacteria | Kingdom | 33843810 | 33357473 | 100 | 100 | 14.4 | 7.2 | 8.7 | 4.8 | 23.05 | 11.93 |
| Proteobac | Phylum | 20527682 | 18948666 | 60.7 | 56.8 | 12.5 | 6.8 | 3.3 | 4.2 | 9.61 | 6.25 |
| Gammap | Class | 12931164 | 11343291 | 38.2 | 34 | 14.8 | 10.5 | 2.9 | 3.9 | 6.77 | 4.89 |
| Burkholc | Order | 5687531 | 4684144 | 16.8 | 14 | 16.1 | 14.1 | 3.2 | 5.4 | 3.24 | 2.73 |
| Comam | Family | 3078487 | 2202077 | 9.1 | 6.6 | 13.3 | 21.2 | 4.2 | 4.3 | 1.59 | 1.68 |
| Comar | Genus | 1267461 | 726605 | 3.7 | 2.2 | 14.2 | 25.7 | 5.3 | 0 | 0.73 | 0.56 |
| Hydrog | Genus | 613979.2 | 270768.5 | 1.8 | 0.8 | 23.7 | 3.1 | 0 | 0 | 0.43 | 0.02 |
| Ramlib | Genus | 291065.5 | 250498.3 | 0.9 | 0.8 | 2.9 | 15.8 | 18.2 | 9.5 | 0.18 | 0.19 |
| Simpli | Genus | 179513.7 | 135286.6 | 0.5 | 0.4 | 10.9 | 47 | 0 | 40.4 | 0.06 | 0.35 |
| Caenir | Genus | 155276.9 | 194439.7 | 0.5 | 0.6 | 6.9 | 12.9 | 2.5 | 2.4 | 0.04 | 0.09 |
| Variov | Genus | 147136.2 | 174751.6 | 0.4 | 0.5 | 2.3 | 1.7 | 0 | 0 | 0.01 | 0.01 |
| Acidov | Genus | 115144 | 59834.8 | 0.3 | 0.2 | 20.3 | 0 | 0 | 0 | 0.07 | 0 |
| Rhizob | Genus | 73065.3 | 93993 | 0.2 | 0.3 | 0 | 87.7 | 0 | 0 | 0 | 0.25 |
| Methyl | Genus | 43188.4 | 61510.2 | 0.1 | 0.2 | 0 | 0 | 0 | 0 | 0 | 0 |
| Pelom | Genus | 30677.2 | 72932.5 | 0.1 | 0.2 | 7.9 | 0 | 0 | 0 | 0.01 | 0 |
| Hylem | Genus | 29566.9 | 27985.5 | 0.1 | 0.1 | 0 | 49 | 0 | 0 | 0 | 0.04 |
| Piscini | Genus | 29138.8 | 44326.1 | 0.1 | 0.1 | 0 | 0 | 0 | 25.8 | 0 | 0.03 |
| Inhella | Genus | 28846.8 | 21666.5 | 0.1 | 0.1 | 0 | 0 | 0 | 0 | 0 | 0 |
| Azohyc | Genus | 16725.6 | 36589.3 | 0 | 0.1 | 0 | 67.9 | 0 | 0 | 0 | 0.07 |
| Limnol | Genus | 14417.6 | 528.6 | 0 | 0 | 58.4 | 0 | 0 | 0 | 0.02 | 0 |
| Diapho | Genus | 6920 | 0 | 0 | 0 | 100 | 0 | 0 | 0 | 0.02 | 0 |
| Ideone | Genus | 6427.7 | 13146.7 | 0 | 0 | 0 | 100 | 0 | 0 | 0 | 0.04 |
| Mitsua | Genus | 6276.8 | 914.3 | 0 | 0 | 0 | 0 | 100 | 0 | 0.02 | 0 |
| Polaro | Genus | 5631.8 | 2148.2 | 0 | 0 | 0 | 0 | 0 | 0 | 0 | 0 |
| Aquab | Genus | 5145.2 | 3525.2 | 0 | 0 | 0 | 0 | 0 | 0 | 0 | 0 |
| Pseud | Genus | 4799.9 | 1049 | 0 | 0 | 0 | 0 | 0 | 0 | 0 | 0 |
| Paucib | Genus | 3753 | 5136.4 | 0 | 0 | 0 | 100 | 0 | 0 | 0 | 0.02 |
| Roseat | Genus | 1857 | 1361.5 | 0 | 0 | 0 | 0 | 0 | 0 | 0 | 0 |
| Pseuda | Genus | 1281.9 | 2006.7 | 0 | 0 | 0 | 0 | 0 | 0 | 0 | 0 |
| RS62 r | Genus | 1091.1 | 1072.2 | 0 | 0 | 0 | 0 | 0 | 0 | 0 | 0 |

| | | | | | | | | | | | |
|---|---|---|---|---|---|---|---|---|---|---|---|
| Delftia | Genus | 99.7 | 0 | 0 | 0 | 0 | 0 | 0 | 0 | 0 | 0 |
| Nitroso | Family | 794597.4 | 746700.7 | 2.3 | 2.2 | 0.7 | 2.7 | 0.2 | 0 | 0.02 | 0.06 |
| Ellin60 | Genus | 532690.5 | 432706.6 | 1.6 | 1.3 | 0 | 0.9 | 0 | 0 | 0 | 0.01 |
| IS-44 | Genus | 159135.9 | 178969.5 | 0.5 | 0.5 | 0 | 0 | 0 | 0 | 0 | 0 |
| MND1 | Genus | 84740.4 | 115404.8 | 0.3 | 0.3 | 3 | 11.1 | 1.9 | 0 | 0.01 | 0.04 |
| Nitros | Genus | 10140.1 | 11398.3 | 0 | 0 | 0 | 0 | 0 | 0 | 0 | 0 |
| Nitros | Genus | 4699.6 | 5154.1 | 0 | 0 | 67.4 | 63.8 | 0 | 0 | 0.01 | 0.01 |
| Nitros | Genus | 3190.8 | 2971.9 | 0 | 0 | 0 | 0 | 0 | 0 | 0 | 0 |
| P3OB- | Genus | 0 | 95.5 | 0 | 0 | 0 | 0 | 0 | 0 | 0 | 0 |
| Methylc | Family | 621245.6 | 686404.6 | 1.8 | 2.1 | 3.3 | 0 | 0 | 11.9 | 0.06 | 0.25 |
| Methyl | Genus | 349069.2 | 258760.2 | 1 | 0.8 | 5.9 | 0 | 0 | 31.6 | 0.06 | 0.25 |
| MM1 | Genus | 95838.3 | 137283.8 | 0.3 | 0.4 | 0 | 0 | 0 | 0 | 0 | 0 |
| Methyl | Genus | 84647.3 | 126927 | 0.3 | 0.4 | 0 | 0 | 0 | 0 | 0 | 0 |
| Methyl | Genus | 65180.4 | 103412.4 | 0.2 | 0.3 | 0 | 0 | 0 | 0 | 0 | 0 |
| MM2 | Genus | 24423.6 | 57057.6 | 0.1 | 0.2 | 0 | 0 | 0 | 0 | 0 | 0 |
| Methyl | Genus | 2086.8 | 2963.5 | 0 | 0 | 0 | 0 | 0 | 0 | 0 | 0 |
| TRA3-2( | Family | 351358.1 | 318743.3 | 1 | 1 | 44.1 | 39.2 | 5.5 | 1.5 | 0.52 | 0.39 |
| TRA3-2 | Genus | 351358.1 | 318743.3 | 1 | 1 | 44.1 | 39.2 | 5.5 | 1.5 | 0.52 | 0.39 |
| Oxaloba | Family | 299398.5 | 337255.4 | 0.9 | 1 | 24.4 | 2.6 | 0.8 | 16 | 0.22 | 0.19 |
| Dugan | Genus | 122631.4 | 115150.9 | 0.4 | 0.3 | 5.5 | 0 | 0 | 35.4 | 0.02 | 0.12 |
| Pseud | Genus | 54726.9 | 28346.1 | 0.2 | 0.1 | 80.2 | 0 | 0 | 7 | 0.13 | 0.01 |
| Massil | Genus | 52264.1 | 111985.2 | 0.2 | 0.3 | 30.2 | 0 | 4.5 | 10.1 | 0.05 | 0.03 |
| Oxalob | Genus | 34534.4 | 23727 | 0.1 | 0.1 | 14.6 | 0 | 0 | 0 | 0.01 | 0 |
| Novihe | Genus | 29095.7 | 54792.7 | 0.1 | 0.2 | 5.6 | 15.7 | 0 | 0 | 0 | 0.03 |
| Hermi | Genus | 2250 | 239.9 | 0 | 0 | 0 | 0 | 0 | 0 | 0 | 0 |
| CM1G | Genus | 2032.6 | 2131.7 | 0 | 0 | 0 | 0 | 0 | 0 | 0 | 0 |
| Oxalici | Genus | 1863.5 | 881.8 | 0 | 0 | 0 | 0 | 0 | 0 | 0 | 0 |
| Rhodoc | Family | 238819.8 | 190519.2 | 0.7 | 0.6 | 62.4 | 13.5 | 12.1 | 0 | 0.53 | 0.08 |
| Azoarc | Genus | 119741.4 | 169394.5 | 0.4 | 0.5 | 63.2 | 10.8 | 24.1 | 0 | 0.31 | 0.05 |
| Dechlo | Genus | 79268.8 | 9177.5 | 0.2 | 0 | 68.2 | 49.3 | 0 | 0 | 0.16 | 0.01 |
| Rhodo | Genus | 18627.1 | 5868.3 | 0.1 | 0 | 59.6 | 0 | 0 | 0 | 0.03 | 0 |

| | | | | | | | | | | | |
|---|---|---|---|---|---|---|---|---|---|---|---|
| Candic | Genus | 8483.2 | 5496.7 | 0 | 0 | 45.6 | 52.5 | 0 | 0 | 0.01 | 0.01 |
| Azospi | Genus | 3896 | 104.8 | 0 | 0 | 61.3 | 0 | 0 | 0 | 0.01 | 0 |
| Uliginc | Genus | 3861.3 | 0 | 0 | 0 | 0 | 0 | 0 | 0 | 0 | 0 |
| Dechlc | Genus | 3004.6 | 0 | 0 | 0 | 63.5 | 0 | 0 | 0 | 0.01 | 0 |
| Propio | Genus | 1497.9 | 0 | 0 | 0 | 0 | 0 | 0 | 0 | 0 | 0 |
| Zooglo | Genus | 439.4 | 477.4 | 0 | 0 | 0 | 0 | 0 | 0 | 0 | 0 |
| SC-I-84 | Family | 123262.9 | 75426.8 | 0.4 | 0.2 | 51.3 | 9 | 0 | 11.1 | 0.19 | 0.05 |
| SC-I-8 | Genus | 123262.9 | 75426.8 | 0.4 | 0.2 | 51.3 | 9 | 0 | 11.1 | 0.19 | 0.05 |
| Alcalige | Family | 78094 | 59392.4 | 0.2 | 0.2 | 20.3 | 0 | 0 | 0 | 0.05 | 0 |
| Achror | Genus | 20513.8 | 4824 | 0.1 | 0 | 77.2 | 0 | 0 | 0 | 0.05 | 0 |
| Advene | Genus | 18463.4 | 13310.1 | 0.1 | 0 | 0 | 0 | 0 | 0 | 0 | 0 |
| Alcalig | Genus | 18357.1 | 22250.8 | 0.1 | 0.1 | 0 | 0 | 0 | 0 | 0 | 0 |
| Parapu | Genus | 11684.6 | 8109.7 | 0 | 0 | 0 | 0 | 0 | 0 | 0 | 0 |
| Candic | Genus | 7046.9 | 6029.6 | 0 | 0 | 0 | 0 | 0 | 0 | 0 | 0 |
| Pusillir | Genus | 2028.2 | 4868.2 | 0 | 0 | 0 | 0 | 0 | 0 | 0 | 0 |
| Burkhol | Family | 40085.9 | 20966.1 | 0.1 | 0.1 | 20.4 | 0 | 0 | 13.6 | 0.02 | 0.01 |
| Lautro | Genus | 30566.8 | 18379.9 | 0.1 | 0.1 | 26.8 | 0 | 0 | 15.5 | 0.02 | 0.01 |
| Burkhc | Genus | 5438.8 | 2185.5 | 0 | 0 | 0 | 0 | 0 | 0 | 0 | 0 |
| Limnol | Genus | 4080.3 | 400.7 | 0 | 0 | 0 | 0 | 0 | 0 | 0 | 0 |
| Chitinib | Family | 19127.4 | 712.6 | 0.1 | 0 | 12.7 | 0 | 0 | 0 | 0.01 | 0 |
| Formiv | Genus | 19127.4 | 712.6 | 0.1 | 0 | 12.7 | 0 | 0 | 0 | 0.01 | 0 |
| Burkhol | Family | 14693.9 | 9540.2 | 0 | 0 | 77.8 | 87.4 | 0 | 0 | 0.03 | 0.03 |
| Chromc | Family | 11559.2 | 21705.8 | 0 | 0.1 | 0 | 0 | 0 | 7 | 0 | 0 |
| Vogese | Genus | 11559.2 | 21705.8 | 0 | 0.1 | 0 | 0 | 0 | 7 | 0 | 0 |
| Chitinir | Family | 8011.7 | 6669.3 | 0 | 0 | 19 | 0 | 0 | 40.3 | 0 | 0.01 |
| Chitini | Genus | 8011.7 | 6669.3 | 0 | 0 | 19 | 0 | 0 | 40.3 | 0 | 0.01 |
| B1-7BS | Family | 5329.5 | 3564.9 | 0 | 0 | 0 | 0 | 0 | 0 | 0 | 0 |
| B1-7BS | Genus | 5329.5 | 3564.9 | 0 | 0 | 0 | 0 | 0 | 0 | 0 | 0 |
| Neisser | Family | 1824.8 | 2685.5 | 0 | 0 | 0 | 0 | 0 | 0 | 0 | 0 |
| Neisse | Genus | 1824.8 | 2685.5 | 0 | 0 | 0 | 0 | 0 | 0 | 0 | 0 |
| A21b | Family | 1635.2 | 1781.2 | 0 | 0 | 0 | 0 | 0 | 0 | 0 | 0 |

| | | | | | | | | | | | |
|---|---|---|---|---|---|---|---|---|---|---|---|
| A21b_I | Genus | 1635.2 | 1781.2 | 0 | 0 | 0 | 0 | 0 | 0 | 0 | 0 |
| Xanthom | Order | 4042029 | 3968303 | 11.9 | 11.9 | 5 | 3.4 | 2.6 | 3.7 | 0.9 | 0.84 |
| Xanthoi | Family | 3393851 | 3298401 | 10 | 9.9 | 4.2 | 1.8 | 1.6 | 3.7 | 0.59 | 0.54 |
| Pseudo | Genus | 1132776 | 957924.2 | 3.3 | 2.9 | 0 | 0 | 0 | 0 | 0 | 0 |
| Luteim | Genus | 583171.2 | 700687.9 | 1.7 | 2.1 | 0 | 0 | 1.5 | 7.1 | 0.03 | 0.15 |
| Arenim | Genus | 455247.8 | 583359.4 | 1.3 | 1.7 | 1.4 | 4.9 | 0.5 | 7.3 | 0.03 | 0.21 |
| Xantho | Genus | 434971.2 | 399000.5 | 1.3 | 1.2 | 6.7 | 3.8 | 1.8 | 0 | 0.11 | 0.05 |
| Lysoba | Genus | 408542.3 | 225508.2 | 1.2 | 0.7 | 10.5 | 0 | 5.3 | 9.2 | 0.19 | 0.06 |
| Therm | Genus | 354712.2 | 391422.9 | 1 | 1.2 | 17.7 | 0 | 4.4 | 2.4 | 0.23 | 0.03 |
| Stenot | Genus | 21133.2 | 35674.4 | 0.1 | 0.1 | 4.4 | 42.9 | 0 | 0 | 0 | 0.05 |
| Xantho | Genus | 3297.8 | 4824 | 0 | 0 | 0 | 0 | 0 | 0 | 0 | 0 |
| Rhodan | Family | 606562.1 | 667600.1 | 1.8 | 2 | 9.6 | 11.2 | 1.6 | 3.6 | 0.2 | 0.3 |
| Dokdo | Genus | 389032.1 | 452793 | 1.1 | 1.4 | 0 | 0 | 0.3 | 0 | 0 | 0 |
| Ahniell | Genus | 98811.2 | 117984.5 | 0.3 | 0.4 | 51.7 | 63.5 | 0 | 0 | 0.15 | 0.22 |
| Tahiba | Genus | 58307.5 | 34950 | 0.2 | 0.1 | 0 | 0 | 0 | 27.2 | 0 | 0.03 |
| Dyella | Genus | 19494.7 | 11365.4 | 0.1 | 0 | 0 | 0 | 33 | 0 | 0.02 | 0 |
| Rhoda | Genus | 17968.6 | 11089.2 | 0.1 | 0 | 17.4 | 0 | 0 | 0 | 0.01 | 0 |
| Luteiba | Genus | 7572.4 | 3406.2 | 0 | 0 | 22.1 | 0 | 0 | 0 | 0 | 0 |
| Chiayii | Genus | 6961.9 | 19952.2 | 0 | 0.1 | 32.6 | 0 | 28.5 | 72.2 | 0.01 | 0.04 |
| Rhoda | Genus | 6763 | 14041.6 | 0 | 0 | 0 | 0 | 0 | 0 | 0 | 0 |
| Rudae | Genus | 1650.8 | 2017.9 | 0 | 0 | 0 | 0 | 0 | 0 | 0 | 0 |
| Xanthoi | Family | 41615.8 | 2301.9 | 0.1 | 0 | 0 | 0 | 95.8 | 0 | 0.12 | 0 |
| Pseudon | Order | 1118095 | 1207133 | 3.3 | 3.6 | 6.7 | 6 | 5.4 | 2.5 | 0.4 | 0.3 |
| Pseudo | Family | 863069.4 | 959586.2 | 2.6 | 2.9 | 1.4 | 7.5 | 4.4 | 1.9 | 0.15 | 0.27 |
| Pseudo | Genus | 859652.4 | 956788.8 | 2.5 | 2.9 | 1.4 | 7.5 | 4.4 | 1.9 | 0.15 | 0.27 |
| Pseudo | Genus | 1782.3 | 2187.8 | 0 | 0 | 0 | 0 | 0 | 0 | 0 | 0 |
| Azomo | Genus | 1634.7 | 609.6 | 0 | 0 | 0 | 0 | 0 | 0 | 0 | 0 |
| Moraxel | Family | 86016.9 | 94321.8 | 0.3 | 0.3 | 2 | 0 | 0 | 0 | 0 | 0 |
| Moraxe | Genus | 81746.4 | 90837.2 | 0.2 | 0.3 | 0 | 0 | 0 | 0 | 0 | 0 |
| Acinet | Genus | 4270.5 | 3484.7 | 0 | 0 | 39.4 | 0 | 0 | 0 | 0 | 0 |
| Pseudo | Family | 51488.4 | 52854.3 | 0.2 | 0.2 | 0 | 0 | 31.2 | 8.1 | 0.05 | 0.01 |

| | | | | | | | | | | | |
|---|---|---|---|---|---|---|---|---|---|---|---|
| Blyi10 | Genus | 51488.4 | 52609.6 | 0.2 | 0.2 | 0 | 0 | 31.2 | 8.1 | 0.05 | 0.01 |
| Pseud | Genus | 0 | 244.7 | 0 | 0 | 0 | 0 | 0 | 0 | 0 | 0 |
| Cellvibr | Family | 45989.1 | 31146.7 | 0.1 | 0.1 | 41.4 | 0 | 0 | 0 | 0.06 | 0 |
| Cellvib | Genus | 45989.1 | 31146.7 | 0.1 | 0.1 | 41.4 | 0 | 0 | 0 | 0.06 | 0 |
| Halieac | Family | 43011.7 | 15788.4 | 0.1 | 0 | 100 | 0 | 0 | 0 | 0.13 | 0 |
| Haliea | Genus | 43011.7 | 15788.4 | 0.1 | 0 | 100 | 0 | 0 | 0 | 0.13 | 0 |
| 211ds2 | Family | 17585.7 | 25085.9 | 0.1 | 0.1 | 0 | 0 | 0 | 0 | 0 | 0 |
| 211ds: | Genus | 17585.7 | 25085.9 | 0.1 | 0.1 | 0 | 0 | 0 | 0 | 0 | 0 |
| Pseudo | Family | 10933.8 | 28349.9 | 0 | 0.1 | 0 | 0 | 60 | 25.2 | 0.02 | 0.02 |
| Gammaı | Order | 1064637 | 867232.6 | 3.1 | 2.6 | 49.4 | 30.2 | 0.8 | 0.3 | 1.58 | 0.79 |
| Unknov | Family | 1064637 | 867232.6 | 3.1 | 2.6 | 49.4 | 30.2 | 0.8 | 0.3 | 1.58 | 0.79 |
| Acidiba | Genus | 852905.8 | 691013.5 | 2.5 | 2.1 | 45.1 | 33.4 | 0.2 | 0 | 1.14 | 0.69 |
| Unkno | Genus | 126565.2 | 88396.3 | 0.4 | 0.3 | 98.4 | 20.1 | 0 | 0 | 0.37 | 0.05 |
| Candic | Genus | 56221.3 | 67318 | 0.2 | 0.2 | 10.4 | 0 | 0 | 0 | 0.02 | 0 |
| Candic | Genus | 28944.2 | 20504.8 | 0.1 | 0.1 | 36.7 | 65.6 | 22.5 | 12.2 | 0.05 | 0.05 |
| Enteroba | Order | 552259.1 | 189301.8 | 1.6 | 0.6 | 14.8 | 12 | 0 | 0 | 0.24 | 0.07 |
| Enterob | Family | 525725.5 | 154117.9 | 1.6 | 0.5 | 14.8 | 14.7 | 0 | 0 | 0.23 | 0.07 |
| Lelliott | Genus | 344910.8 | 39645.9 | 1 | 0.1 | 0 | 0 | 0 | 0 | 0 | 0 |
| Entero | Genus | 118812.9 | 31678.2 | 0.4 | 0.1 | 22.8 | 0 | 0 | 0 | 0.08 | 0 |
| Cedec | Genus | 45694.5 | 36040 | 0.1 | 0.1 | 91 | 0 | 0 | 0 | 0.12 | 0 |
| Entero | Genus | 13806.1 | 36666.1 | 0 | 0.1 | 64.7 | 34.2 | 0 | 0 | 0.03 | 0.04 |
| Buttiaı | Genus | 2501.2 | 10087.8 | 0 | 0 | 0 | 100 | 0 | 0 | 0 | 0.03 |
| Aeromc | Family | 26533.7 | 35183.9 | 0.1 | 0.1 | 16.6 | 0 | 0 | 0 | 0.01 | 0 |
| Aerom | Genus | 26533.7 | 35183.9 | 0.1 | 0.1 | 16.6 | 0 | 0 | 0 | 0.01 | 0 |
| CCD24 | Order | 184436.8 | 193512 | 0.5 | 0.6 | 1.1 | 1.4 | 3.2 | 0 | 0.02 | 0.01 |
| CCD24_ | Family | 184436.8 | 193512 | 0.5 | 0.6 | 1.1 | 1.4 | 3.2 | 0 | 0.02 | 0.01 |
| R7C24 | Order | 145393 | 91110.9 | 0.4 | 0.3 | 71 | 28 | 0 | 0 | 0.3 | 0.08 |
| R7C24_ | Family | 145393 | 91110.9 | 0.4 | 0.3 | 71 | 28 | 0 | 0 | 0.3 | 0.08 |
| Steroido | Order | 50711.7 | 34358.9 | 0.1 | 0.1 | 0 | 0 | 0 | 0 | 0 | 0 |
| Steroid | Family | 50421.4 | 34165.8 | 0.1 | 0.1 | 0 | 0 | 0 | 0 | 0 | 0 |
| Steroic | Genus | 37035.5 | 23840.9 | 0.1 | 0.1 | 0 | 0 | 0 | 0 | 0 | 0 |

| | | | | | | | | | | | |
|---|---|---|---|---|---|---|---|---|---|---|---|
| Steroic | Genus | 13385.9 | 10324.9 | 0 | 0 | 0 | 0 | 0 | 0 | 0 | 0 |
| Woesei | Family | 290.3 | 193.1 | 0 | 0 | 0 | 0 | 0 | 0 | 0 | 0 |
| JTB255 | Genus | 290.3 | 0 | 0 | 0 | 0 | 0 | 0 | 0 | 0 | 0 |
| Woese | Genus | 0 | 193.1 | 0 | 0 | 0 | 0 | 0 | 0 | 0 | 0 |
| PLTA13 | Order | 23199.6 | 26815.5 | 0.1 | 0.1 | 7 | 12.3 | 0 | 0 | 0 | 0.01 |
| PLTA13 | Family | 23199.6 | 26815.5 | 0.1 | 0.1 | 7 | 12.3 | 0 | 0 | 0 | 0.01 |
| Gamma | Order | 22775.3 | 24478.8 | 0.1 | 0.1 | 59.5 | 0 | 27.4 | 35.8 | 0.06 | 0.03 |
| Diplorick | Order | 22518.4 | 29676.5 | 0.1 | 0.1 | 0 | 0 | 9.3 | 0 | 0.01 | 0 |
| Diploric | Family | 22518.4 | 29676.5 | 0.1 | 0.1 | 0 | 0 | 9.3 | 0 | 0.01 | 0 |
| Diplori | Genus | 22518.4 | 29676.5 | 0.1 | 0.1 | 0 | 0 | 9.3 | 0 | 0.01 | 0 |
| Coxiellal | Order | 6365.7 | 3050.7 | 0 | 0 | 0 | 0 | 0 | 0 | 0 | 0 |
| Coxiella | Family | 6365.7 | 3050.7 | 0 | 0 | 0 | 0 | 0 | 0 | 0 | 0 |
| Coxiell | Genus | 6365.7 | 3050.7 | 0 | 0 | 0 | 0 | 0 | 0 | 0 | 0 |
| Ga0077 | Order | 4444.7 | 5881.1 | 0 | 0 | 0 | 39.2 | 0 | 24.8 | 0 | 0.01 |
| Ga0077 | Family | 4444.7 | 5881.1 | 0 | 0 | 0 | 39.2 | 0 | 24.8 | 0 | 0.01 |
| EV818S\ | Order | 3112 | 10245.2 | 0 | 0 | 0 | 65.1 | 39.1 | 0 | 0 | 0.02 |
| EV818S | Family | 3112 | 10245.2 | 0 | 0 | 0 | 65.1 | 39.1 | 0 | 0 | 0.02 |
| EPR396 | Order | 1204.2 | 1627.6 | 0 | 0 | 0 | 0 | 0 | 0 | 0 | 0 |
| EPR396 | Family | 1204.2 | 1627.6 | 0 | 0 | 0 | 0 | 0 | 0 | 0 | 0 |
| Chromat | Order | 1094.6 | 3230.4 | 0 | 0 | 0 | 0 | 0 | 0 | 0 | 0 |
| Sedime | Family | 1094.6 | 3230.4 | 0 | 0 | 0 | 0 | 0 | 0 | 0 | 0 |
| Sedim | Genus | 1094.6 | 3230.4 | 0 | 0 | 0 | 0 | 0 | 0 | 0 | 0 |
| JTB23 | Order | 564.7 | 81.6 | 0 | 0 | 0 | 0 | 0 | 0 | 0 | 0 |
| JTB23_( | Family | 564.7 | 81.6 | 0 | 0 | 0 | 0 | 0 | 0 | 0 | 0 |
| Legionel | Order | 528.8 | 1418 | 0 | 0 | 0 | 0 | 0 | 0 | 0 | 0 |
| Legione | Family | 528.8 | 1418 | 0 | 0 | 0 | 0 | 0 | 0 | 0 | 0 |
| Legion | Genus | 528.8 | 1418 | 0 | 0 | 0 | 0 | 0 | 0 | 0 | 0 |
| Salinispl | Order | 164.3 | 1576 | 0 | 0 | 0 | 0 | 0 | 0 | 0 | 0 |
| Solimor | Family | 164.3 | 1576 | 0 | 0 | 0 | 0 | 0 | 0 | 0 | 0 |
| Fontim | Genus | 164.3 | 118.8 | 0 | 0 | 0 | 0 | 0 | 0 | 0 | 0 |
| Panaca | Genus | 0 | 1457.3 | 0 | 0 | 0 | 0 | 0 | 0 | 0 | 0 |

| | | | | | | | | | | | |
|---|---|---|---|---|---|---|---|---|---|---|---|
| Granulos | Order | 99.7 | 112.6 | 0 | 0 | 0 | 0 | 0 | 0 | 0 | 0 |
| Granulo | Family | 99.7 | 112.6 | 0 | 0 | 0 | 0 | 0 | 0 | 0 | 0 |
| Alphaprot | Class | 7596518 | 7605375 | 22.4 | 22.8 | 9.1 | 1.6 | 4.6 | 4.8 | 3.08 | 1.48 |
| Rhizobia | Order | 2671774 | 2441710 | 7.9 | 7.3 | 13.9 | 0.6 | 3.4 | 1.8 | 1.37 | 0.18 |
| Xanthol | Family | 721154.2 | 685582.8 | 2.1 | 2.1 | 8.1 | 1.6 | 8.9 | 1.4 | 0.36 | 0.06 |
| Pseud | Genus | 233122 | 234772.6 | 0.7 | 0.7 | 0 | 1.4 | 1.4 | 2.8 | 0.01 | 0.03 |
| Xantho | Genus | 204915.1 | 162806.4 | 0.6 | 0.5 | 19.8 | 0 | 4.6 | 0 | 0.15 | 0 |
| Rhodo | Genus | 175588.3 | 169173.7 | 0.5 | 0.5 | 2.5 | 0 | 29.2 | 0 | 0.16 | 0 |
| Pseud | Genus | 55613.5 | 53249.5 | 0.2 | 0.2 | 20.4 | 13.6 | 0 | 5.5 | 0.03 | 0.03 |
| Bradyr | Genus | 31613.9 | 37690.3 | 0.1 | 0.1 | 0 | 0 | 0 | 0 | 0 | 0 |
| Rhodo | Genus | 8145 | 15421.9 | 0 | 0 | 0 | 0 | 0 | 0 | 0 | 0 |
| Variiba | Genus | 7009.4 | 6809.1 | 0 | 0 | 32.9 | 0 | 0 | 0 | 0.01 | 0 |
| Oligotr | Genus | 2666.3 | 2124.8 | 0 | 0 | 0 | 0 | 0 | 0 | 0 | 0 |
| Nitroba | Genus | 2480.6 | 3432.7 | 0 | 0 | 0 | 0 | 0 | 0 | 0 | 0 |
| Xantho | Genus | 0 | 101.8 | 0 | 0 | 0 | 0 | 0 | 0 | 0 | 0 |
| Devosia | Family | 675888.3 | 480257.6 | 2 | 1.4 | 28.6 | 1 | 0.1 | 3.9 | 0.57 | 0.07 |
| Devosi | Genus | 656489.5 | 460718.1 | 1.9 | 1.4 | 29.4 | 1 | 0.1 | 4.1 | 0.57 | 0.07 |
| Devosi | Genus | 19398.9 | 19539.5 | 0.1 | 0.1 | 0 | 0 | 0 | 0 | 0 | 0 |
| Rhizobi | Family | 500890 | 556767 | 1.5 | 1.7 | 12.5 | 0 | 3 | 1.5 | 0.23 | 0.02 |
| Shinell | Genus | 116715.3 | 184955.4 | 0.3 | 0.6 | 0 | 0 | 0 | 0 | 0 | 0 |
| Rhizob | Genus | 86243.4 | 86862.6 | 0.3 | 0.3 | 24.8 | 0 | 3.4 | 0 | 0.07 | 0 |
| Mesorl | Genus | 84268.3 | 79662.8 | 0.2 | 0.2 | 16.3 | 0 | 0 | 7.8 | 0.04 | 0.02 |
| Allorhi | Genus | 50315.7 | 49126.3 | 0.1 | 0.1 | 3.7 | 0 | 7.6 | 0 | 0.02 | 0 |
| [Rhizol | Genus | 43837.4 | 32334.9 | 0.1 | 0.1 | 27.6 | 0 | 0 | 0 | 0.04 | 0 |
| Ensifer | Genus | 34127.4 | 35454.8 | 0.1 | 0.1 | 20.5 | 0 | 0 | 0 | 0.02 | 0 |
| Pseuda | Genus | 30380.6 | 32191.5 | 0.1 | 0.1 | 0 | 0 | 0 | 6.2 | 0 | 0.01 |
| Amino | Genus | 21912.3 | 28327.7 | 0.1 | 0.1 | 0 | 0 | 0 | 0 | 0 | 0 |
| Ciceril | Genus | 15806.2 | 18605 | 0 | 0.1 | 28.1 | 0 | 51.5 | 0 | 0.04 | 0 |
| Aliihoe | Genus | 5163.7 | 305.4 | 0 | 0 | 0 | 0 | 0 | 0 | 0 | 0 |
| Neorhi | Genus | 2954.2 | 1981 | 0 | 0 | 0 | 0 | 0 | 0 | 0 | 0 |
| Pseud | Genus | 2786.6 | 987 | 0 | 0 | 0 | 0 | 0 | 0 | 0 | 0 |

| | | | | | | | | | | | |
|---|---|---|---|---|---|---|---|---|---|---|---|
| Ochrol | Genus | 2359.2 | 2861.4 | 0 | 0 | 0 | 0 | 0 | 0 | 0 | 0 |
| Mycop | Genus | 2247.4 | 688.6 | 0 | 0 | 100 | 0 | 0 | 0 | 0.01 | 0 |
| Phyllol | Genus | 1027.3 | 538.7 | 0 | 0 | 0 | 0 | 0 | 0 | 0 | 0 |
| Chelat | Genus | 744.9 | 1883.9 | 0 | 0 | 0 | 0 | 0 | 0 | 0 | 0 |
| Hyphon | Family | 204682.2 | 188918.6 | 0.6 | 0.6 | 1.8 | 0 | 1.2 | 1.2 | 0.02 | 0.01 |
| Hypho | Genus | 97412.8 | 90732.2 | 0.3 | 0.3 | 1.6 | 0 | 0 | 0 | 0 | 0 |
| Hypho | Genus | 65921.1 | 54483.3 | 0.2 | 0.2 | 3.2 | 0 | 3.9 | 0 | 0.01 | 0 |
| Pedom | Genus | 41348.3 | 43703.1 | 0.1 | 0.1 | 0 | 0 | 0 | 5.4 | 0 | 0.01 |
| Rhizobi | Family | 177899.9 | 170661.1 | 0.5 | 0.5 | 24.3 | 0 | 0 | 2.3 | 0.13 | 0.01 |
| Bauldi | Genus | 154573.2 | 151148.8 | 0.5 | 0.5 | 24.6 | 0 | 0 | 2.6 | 0.11 | 0.01 |
| Rhizob | Genus | 14291.6 | 13133.7 | 0 | 0 | 21.5 | 0 | 0 | 0 | 0.01 | 0 |
| Nordel | Genus | 9035.1 | 6378.6 | 0 | 0 | 24.4 | 0 | 0 | 0 | 0.01 | 0 |
| Amb-16 | Family | 133769.4 | 107661.8 | 0.4 | 0.3 | 2.5 | 0 | 0 | 0 | 0.01 | 0 |
| Amb-1 | Genus | 133769.4 | 107661.8 | 0.4 | 0.3 | 2.5 | 0 | 0 | 0 | 0.01 | 0 |
| Beijerin | Family | 87128.7 | 63661.9 | 0.3 | 0.2 | 0 | 0 | 0 | 0 | 0 | 0 |
| Bosea | Genus | 51061.3 | 29886.7 | 0.2 | 0.1 | 0 | 0 | 0 | 0 | 0 | 0 |
| Microv | Genus | 35970.9 | 33775.2 | 0.1 | 0.1 | 0 | 0 | 0 | 0 | 0 | 0 |
| Methyl | Genus | 96.5 | 0 | 0 | 0 | 0 | 0 | 0 | 0 | 0 | 0 |
| KF-JG30 | Family | 56195.6 | 68058.4 | 0.2 | 0.2 | 0 | 0 | 0 | 0 | 0 | 0 |
| KF-JG3 | Genus | 56195.6 | 68058.4 | 0.2 | 0.2 | 0 | 0 | 0 | 0 | 0 | 0 |
| Rhizobi | Family | 33938.4 | 34848.7 | 0.1 | 0.1 | 0 | 0 | 16 | 0 | 0.02 | 0 |
| Labrace | Family | 19381.4 | 20264.6 | 0.1 | 0.1 | 0 | 0 | 0 | 6.5 | 0 | 0 |
| Labrys | Genus | 19381.4 | 20264.6 | 0.1 | 0.1 | 0 | 0 | 0 | 6.5 | 0 | 0 |
| WC3-1 | Family | 17646.9 | 15156.9 | 0.1 | 0 | 0 | 0 | 0 | 0 | 0 | 0 |
| WC3-1 | Genus | 17646.9 | 15156.9 | 0.1 | 0 | 0 | 0 | 0 | 0 | 0 | 0 |
| A0839 | Family | 13782.6 | 11616.1 | 0 | 0 | 33.1 | 0 | 0 | 0 | 0.01 | 0 |
| A0839 | Genus | 13782.6 | 11616.1 | 0 | 0 | 33.1 | 0 | 0 | 0 | 0.01 | 0 |
| Methylc | Family | 10329.6 | 9748 | 0 | 0 | 0 | 0 | 11.7 | 0 | 0 | 0 |
| Methyl | Genus | 10329.6 | 9748 | 0 | 0 | 0 | 0 | 11.7 | 0 | 0 | 0 |
| C2U | Family | 8244.1 | 12740.6 | 0 | 0 | 34.1 | 0 | 23.4 | 0 | 0.01 | 0 |
| C2U_F | Genus | 8244.1 | 12740.6 | 0 | 0 | 34.1 | 0 | 23.4 | 0 | 0.01 | 0 |

| | | | | | | | | | | | |
|---|---|---|---|---|---|---|---|---|---|---|---|
| D05-2 | Family | 6465.3 | 6896.5 | 0 | 0 | 0 | 0 | 0 | 0 | 0 | 0 |
| D05-2_ | Genus | 6465.3 | 6896.5 | 0 | 0 | 0 | 0 | 0 | 0 | 0 | 0 |
| Kaistiac | Family | 2766.6 | 8138.7 | 0 | 0 | 0 | 0 | 0 | 0 | 0 | 0 |
| Kaistia | Genus | 2766.6 | 8138.7 | 0 | 0 | 0 | 0 | 0 | 0 | 0 | 0 |
| Rhodob | Family | 1610.6 | 730.8 | 0 | 0 | 0 | 0 | 0 | 0 | 0 | 0 |
| Tepida | Genus | 1610.6 | 730.8 | 0 | 0 | 0 | 0 | 0 | 0 | 0 | 0 |
| Sphingo | Order | 2491318 | 2714972 | 7.4 | 8.1 | 3.2 | 1.1 | 3.4 | 5.2 | 0.48 | 0.51 |
| Sphingo | Family | 2491318 | 2714972 | 7.4 | 8.1 | 3.2 | 1.1 | 3.4 | 5.2 | 0.48 | 0.51 |
| Sphing | Genus | 1456793 | 1686461 | 4.3 | 5.1 | 1.1 | 0 | 0 | 0 | 0.05 | 0 |
| Sphing | Genus | 532866.5 | 645860.4 | 1.6 | 1.9 | 2.4 | 4.5 | 9.4 | 15.1 | 0.19 | 0.38 |
| Alterer | Genus | 225795.8 | 183710.4 | 0.7 | 0.6 | 17.7 | 0 | 4 | 9 | 0.14 | 0.05 |
| Sphing | Genus | 90870 | 96219.1 | 0.3 | 0.3 | 0 | 0 | 0 | 24.6 | 0 | 0.07 |
| Novos | Genus | 62203.4 | 31563.2 | 0.2 | 0.1 | 15.4 | 0 | 29.4 | 0 | 0.08 | 0 |
| Sphing | Genus | 61263.3 | 31140.7 | 0.2 | 0.1 | 0 | 0 | 12.3 | 0 | 0.02 | 0 |
| Ellin60 | Genus | 41141.7 | 30617.4 | 0.1 | 0.1 | 0 | 0 | 0 | 3.5 | 0 | 0 |
| Sphing | Genus | 5782.6 | 910.1 | 0 | 0 | 0 | 0 | 0 | 0 | 0 | 0 |
| Qipeng | Genus | 4656.3 | 1947.2 | 0 | 0 | 0 | 0 | 0 | 67.4 | 0 | 0 |
| Porphy | Genus | 4363.6 | 2679.6 | 0 | 0 | 0 | 0 | 0 | 0 | 0 | 0 |
| Plot4-2 | Genus | 3941.2 | 3170.4 | 0 | 0 | 0 | 0 | 0 | 0 | 0 | 0 |
| Rhizorl | Genus | 996.2 | 0 | 0 | 0 | 0 | 0 | 0 | 0 | 0 | 0 |
| Parabl | Genus | 644.6 | 692.3 | 0 | 0 | 0 | 0 | 0 | 0 | 0 | 0 |
| Cauloba | Order | 1149671 | 1058088 | 3.4 | 3.2 | 18.8 | 4.1 | 2.6 | 13.5 | 0.73 | 0.56 |
| Caulob | Family | 598497.3 | 530986.5 | 1.8 | 1.6 | 5.5 | 1.1 | 5.1 | 25.5 | 0.19 | 0.42 |
| Brevur | Genus | 288997.5 | 209694.1 | 0.9 | 0.6 | 6.8 | 2.9 | 1.3 | 1.8 | 0.07 | 0.03 |
| Caulot | Genus | 120421.2 | 101590.4 | 0.4 | 0.3 | 3 | 0 | 7.9 | 33.2 | 0.04 | 0.1 |
| Caulot | Genus | 80210.6 | 138514.7 | 0.2 | 0.4 | 0 | 0 | 14.9 | 70.8 | 0.04 | 0.29 |
| Phenyl | Genus | 63782.6 | 40860.6 | 0.2 | 0.1 | 1.6 | 0 | 4.2 | 0 | 0.01 | 0 |
| Asticca | Genus | 45085.4 | 40326.7 | 0.1 | 0.1 | 19.1 | 0 | 5.5 | 0 | 0.03 | 0 |
| Hyphon | Family | 547844.2 | 525768.9 | 1.6 | 1.6 | 33 | 7.1 | 0 | 1.4 | 0.53 | 0.13 |
| SWB02 | Genus | 352193.8 | 324026.4 | 1 | 1 | 44 | 0.8 | 0 | 2.3 | 0.46 | 0.03 |
| Hirsch | Genus | 195650.5 | 201742.5 | 0.6 | 0.6 | 13.1 | 17.1 | 0 | 0 | 0.08 | 0.1 |

| | | | | | | | | | | | |
|---|---|---|---|---|---|---|---|---|---|---|---|
| Parvula | Family | 3112.3 | 1146.5 | 0 | 0 | 64.9 | 0 | 0 | 0 | 0.01 | 0 |
| Amphi | Genus | 3112.3 | 1146.5 | 0 | 0 | 64.9 | 0 | 0 | 0 | 0.01 | 0 |
| Caulob | Family | 217 | 186.3 | 0 | 0 | 0 | 0 | 0 | 0 | 0 | 0 |
| Micrope | Order | 369782.7 | 438629.9 | 1.1 | 1.3 | 0 | 0 | 15.1 | 1.3 | 0.16 | 0.02 |
| Micrope | Family | 369782.7 | 438629.9 | 1.1 | 1.3 | 0 | 0 | 15.1 | 1.3 | 0.16 | 0.02 |
| Microp | Genus | 369782.7 | 438629.9 | 1.1 | 1.3 | 0 | 0 | 15.1 | 1.3 | 0.16 | 0.02 |
| Reyranel | Order | 276695.5 | 343451 | 0.8 | 1 | 0 | 0 | 1.7 | 0 | 0.01 | 0 |
| Reyrane | Family | 276695.5 | 343451 | 0.8 | 1 | 0 | 0 | 1.7 | 0 | 0.01 | 0 |
| Reyran | Genus | 200243.8 | 249581 | 0.6 | 0.7 | 0 | 0 | 2.4 | 0 | 0.01 | 0 |
| Reyran | Genus | 76451.7 | 93870 | 0.2 | 0.3 | 0 | 0 | 0 | 0 | 0 | 0 |
| Alphapr | Order | 120815.5 | 136806.4 | 0.4 | 0.4 | 1.9 | 0 | 4.4 | 0.6 | 0.02 | 0 |
| Rhodosp | Order | 112384.4 | 130058.3 | 0.3 | 0.4 | 0 | 15.6 | 5.4 | 1.2 | 0.02 | 0.07 |
| Rhodos | Family | 73261.4 | 85882.4 | 0.2 | 0.3 | 0 | 23.6 | 6.7 | 1.8 | 0.01 | 0.07 |
| Magnet | Family | 22367.8 | 28242.1 | 0.1 | 0.1 | 0 | 0 | 0 | 0 | 0 | 0 |
| Magne | Genus | 22367.8 | 28242.1 | 0.1 | 0.1 | 0 | 0 | 0 | 0 | 0 | 0 |
| Rhodos | Family | 16755.3 | 15933.8 | 0 | 0 | 0 | 0 | 7.1 | 0 | 0 | 0 |
| Rhodo | Genus | 16755.3 | 15933.8 | 0 | 0 | 0 | 0 | 7.1 | 0 | 0 | 0 |
| Rickettsi | Order | 104371.4 | 77271.4 | 0.3 | 0.2 | 4.9 | 22.7 | 53.7 | 26.8 | 0.18 | 0.11 |
| SM2D1 | Family | 104177.6 | 76376.3 | 0.3 | 0.2 | 5 | 23 | 53.8 | 27.1 | 0.18 | 0.11 |
| SM2D1 | Genus | 104177.6 | 76376.3 | 0.3 | 0.2 | 5 | 23 | 53.8 | 27.1 | 0.18 | 0.11 |
| Ricketts | Family | 193.9 | 895.1 | 0 | 0 | 0 | 0 | 0 | 0 | 0 | 0 |
| Rhodoba | Order | 72827.4 | 68553.6 | 0.2 | 0.2 | 7.1 | 0 | 5.2 | 18.3 | 0.03 | 0.04 |
| Rhodob | Family | 72827.4 | 68553.6 | 0.2 | 0.2 | 7.1 | 0 | 5.2 | 18.3 | 0.03 | 0.04 |
| Rhodo | Genus | 29826.8 | 26811.2 | 0.1 | 0.1 | 0 | 0 | 0 | 0 | 0 | 0 |
| Paracc | Genus | 20635.1 | 28728.4 | 0.1 | 0.1 | 11.6 | 0 | 0 | 43.7 | 0.01 | 0.04 |
| Amaric | Genus | 11774.1 | 12890.7 | 0 | 0 | 0 | 0 | 32.1 | 0 | 0.01 | 0 |
| Cereib | Genus | 7126.1 | 0 | 0 | 0 | 0 | 0 | 0 | 0 | 0 | 0 |
| Tabrizi | Genus | 2780.5 | 123.4 | 0 | 0 | 100 | 0 | 0 | 0 | 0.01 | 0 |
| Pseud | Genus | 684.9 | 0 | 0 | 0 | 0 | 0 | 0 | 0 | 0 | 0 |
| Dongiale | Order | 52407 | 49983.1 | 0.2 | 0.1 | 0 | 0 | 0 | 0 | 0 | 0 |
| Dongiac | Family | 52407 | 49983.1 | 0.2 | 0.1 | 0 | 0 | 0 | 0 | 0 | 0 |

| | | | | | | | | | | | |
|---|---|---|---|---|---|---|---|---|---|---|---|
| Dongia | Genus | 52407 | 49983.1 | 0.2 | 0.1 | 0 | 0 | 0 | 0 | 0 | 0 |
| Micavibr | Order | 46288.1 | 25605.7 | 0.1 | 0.1 | 0 | 0 | 2.4 | 0 | 0 | 0 |
| Micavib | Family | 34479.2 | 15882 | 0.1 | 0 | 0 | 0 | 3.3 | 0 | 0 | 0 |
| Micavib | Family | 11808.9 | 9723.6 | 0 | 0 | 0 | 0 | 0 | 0 | 0 | 0 |
| Micavi | Genus | 11808.9 | 9723.6 | 0 | 0 | 0 | 0 | 0 | 0 | 0 | 0 |
| Kiloniella | Order | 31725.3 | 27280.5 | 0.1 | 0.1 | 11.6 | 0 | 6.1 | 0 | 0.02 | 0 |
| Fodinic | Family | 31479.9 | 26853.6 | 0.1 | 0.1 | 11.7 | 0 | 6.1 | 0 | 0.02 | 0 |
| Fodinic | Genus | 31479.9 | 26853.6 | 0.1 | 0.1 | 11.7 | 0 | 6.1 | 0 | 0.02 | 0 |
| Kiloniel | Family | 245.4 | 426.9 | 0 | 0 | 0 | 0 | 0 | 0 | 0 | 0 |
| Kilonie | Genus | 245.4 | 426.9 | 0 | 0 | 0 | 0 | 0 | 0 | 0 | 0 |
| Azospiril | Order | 26228.3 | 17680.6 | 0.1 | 0.1 | 6.5 | 0 | 15.3 | 0 | 0.02 | 0 |
| Azospir | Family | 15390.3 | 3962.4 | 0 | 0 | 0 | 0 | 26.1 | 0 | 0.01 | 0 |
| Azospi | Genus | 13832.2 | 3096.1 | 0 | 0 | 0 | 0 | 29.1 | 0 | 0.01 | 0 |
| Azospi | Genus | 1558.1 | 866.3 | 0 | 0 | 0 | 0 | 0 | 0 | 0 | 0 |
| Azospir | Family | 6658.9 | 10047.3 | 0 | 0 | 0 | 0 | 0 | 0 | 0 | 0 |
| Stella | Genus | 6658.9 | 10047.3 | 0 | 0 | 0 | 0 | 0 | 0 | 0 | 0 |
| Inquilin | Family | 4179 | 3670.9 | 0 | 0 | 40.7 | 0 | 0 | 0 | 0.01 | 0 |
| Inquili | Genus | 4179 | 3670.9 | 0 | 0 | 40.7 | 0 | 0 | 0 | 0.01 | 0 |
| Defluviic | Order | 23381.3 | 28262.5 | 0.1 | 0.1 | 0 | 0 | 0 | 0 | 0 | 0 |
| Defluvii | Family | 14456.7 | 15900 | 0 | 0 | 0 | 0 | 0 | 0 | 0 | 0 |
| Defluvii | Family | 8924.6 | 12362.6 | 0 | 0 | 0 | 0 | 0 | 0 | 0 | 0 |
| Defluv | Genus | 8924.6 | 12362.6 | 0 | 0 | 0 | 0 | 0 | 0 | 0 | 0 |
| Acetoba | Order | 13996.3 | 13775.9 | 0 | 0 | 23.5 | 0 | 0 | 0 | 0.01 | 0 |
| Acetoba | Family | 13996.3 | 13775.9 | 0 | 0 | 23.5 | 0 | 0 | 0 | 0.01 | 0 |
| Acetob | Genus | 7617.6 | 11412.6 | 0 | 0 | 0 | 0 | 0 | 0 | 0 | 0 |
| Roseo | Genus | 6164.9 | 2161.7 | 0 | 0 | 53.4 | 0 | 0 | 0 | 0.01 | 0 |
| Craurc | Genus | 213.7 | 201.5 | 0 | 0 | 0 | 0 | 0 | 0 | 0 | 0 |
| Elsterale | Order | 10963.8 | 18046.8 | 0 | 0.1 | 0 | 0 | 0 | 0 | 0 | 0 |
| Elsteral | Family | 10963.8 | 18046.8 | 0 | 0.1 | 0 | 0 | 0 | 0 | 0 | 0 |
| Ferrovibi | Order | 8435.9 | 3226 | 0 | 0 | 0 | 0 | 0 | 0 | 0 | 0 |
| Ferrovit | Family | 7348.5 | 1661.7 | 0 | 0 | 0 | 0 | 0 | 0 | 0 | 0 |

| | | | | | | | | | | | |
|---|---|---|---|---|---|---|---|---|---|---|---|
| Ferrovi | Genus | 7348.5 | 1661.7 | 0 | 0 | 0 | 0 | 0 | 0 | 0 | 0 |
| Taonell | Family | 1087.5 | 1163.8 | 0 | 0 | 0 | 0 | 0 | 0 | 0 | 0 |
| Taonel | Genus | 1087.5 | 1163.8 | 0 | 0 | 0 | 0 | 0 | 0 | 0 | 0 |
| Ferrovit | Family | 0 | 400.5 | 0 | 0 | 0 | 0 | 0 | 0 | 0 | 0 |
| Paracae | Order | 7613 | 5184.9 | 0 | 0 | 53.4 | 0 | 46.6 | 0 | 0.02 | 0 |
| Paracae | Family | 7613 | 5184.9 | 0 | 0 | 53.4 | 0 | 46.6 | 0 | 0.02 | 0 |
| Paraca | Genus | 4062 | 549 | 0 | 0 | 100 | 0 | 0 | 0 | 0.01 | 0 |
| Candic | Genus | 3551 | 3814.2 | 0 | 0 | 0 | 0 | 100 | 0 | 0.01 | 0 |
| Candic | Genus | 0 | 821.7 | 0 | 0 | 0 | 0 | 0 | 0 | 0 | 0 |
| Zavarzin | Order | 3397.5 | 4721.4 | 0 | 0 | 0 | 0 | 0 | 0 | 0 | 0 |
| Zavarzir | Family | 3397.5 | 4721.4 | 0 | 0 | 0 | 0 | 0 | 0 | 0 | 0 |
| Parvibac | Order | 1979 | 1320.7 | 0 | 0 | 0 | 0 | 0 | 0 | 0 | 0 |
| Parviba | Family | 1979 | 1320.7 | 0 | 0 | 0 | 0 | 0 | 0 | 0 | 0 |
| Parviba | Genus | 1979 | 1320.7 | 0 | 0 | 0 | 0 | 0 | 0 | 0 | 0 |
| AT-s3-44 | Order | 462.5 | 746.2 | 0 | 0 | 0 | 0 | 0 | 0 | 0 | 0 |
| AT-s3-4 | Family | 462.5 | 746.2 | 0 | 0 | 0 | 0 | 0 | 0 | 0 | 0 |
| Bacteroidc | Phylum | 5588157 | 4504678 | 16.5 | 13.5 | 16 | 7.3 | 32 | 9.5 | 7.92 | 2.26 |
| Bacteroid | Class | 5415949 | 4371355 | 16 | 13.1 | 16.5 | 7.5 | 32.7 | 9.7 | 7.87 | 2.26 |
| Flavobac | Order | 3934554 | 3167965 | 11.6 | 9.5 | 9 | 6.5 | 37.6 | 11.8 | 5.41 | 1.74 |
| Flavoba | Family | 3685804 | 3002701 | 10.9 | 9 | 6.1 | 4.8 | 40.1 | 12.4 | 5.02 | 1.55 |
| Flavob | Genus | 3592884 | 2970369 | 10.6 | 8.9 | 6.1 | 4.9 | 40.9 | 12.5 | 4.99 | 1.55 |
| Vitellib | Genus | 50463.6 | 6975.9 | 0.1 | 0 | 0 | 0 | 14.2 | 18 | 0.02 | 0 |
| Flavob | Genus | 19130.7 | 7862.5 | 0.1 | 0 | 0 | 0 | 0 | 0 | 0 | 0 |
| Salinin | Genus | 7030 | 4039.9 | 0 | 0 | 0 | 0 | 0 | 0 | 0 | 0 |
| Aequo | Genus | 6382.7 | 8357.2 | 0 | 0 | 0 | 0 | 0 | 0 | 0 | 0 |
| Subsa | Genus | 4734.2 | 2554.2 | 0 | 0 | 100 | 0 | 0 | 0 | 0.01 | 0 |
| Gilviba | Genus | 3828.1 | 1667.3 | 0 | 0 | 0 | 0 | 0 | 0 | 0 | 0 |
| Flavivi | Genus | 1349.9 | 874.8 | 0 | 0 | 0 | 0 | 0 | 0 | 0 | 0 |
| NS9 ma | Family | 174254.9 | 139130.3 | 0.5 | 0.4 | 52.9 | 33.8 | 1.5 | 0 | 0.28 | 0.14 |
| NS9 m | Genus | 174254.9 | 139130.3 | 0.5 | 0.4 | 52.9 | 33.8 | 1.5 | 0 | 0.28 | 0.14 |
| Weekse | Family | 73206.9 | 26134.1 | 0.2 | 0.1 | 50.1 | 60.7 | 0 | 0 | 0.11 | 0.05 |

| | | | | | | | | | | | |
|---|---|---|---|---|---|---|---|---|---|---|---|
| Chryse | Genus | 73206.9 | 26134.1 | 0.2 | 0.1 | 50.1 | 60.7 | 0 | 0 | 0.11 | 0.05 |
| Flavoba | Family | 1288.4 | 0 | 0 | 0 | 0 | 0 | 0 | 0 | 0 | 0 |
| Chitinop | Order | 1071648 | 848897.5 | 3.2 | 2.5 | 43.3 | 12.5 | 19.9 | 1.4 | 2 | 0.35 |
| Sapros | Family | 1009082 | 824276.8 | 3 | 2.5 | 44.4 | 12.9 | 17.5 | 0.7 | 1.85 | 0.33 |
| Sapros | Genus | 850636.9 | 726035.3 | 2.5 | 2.2 | 50.6 | 1.5 | 5.6 | 0.8 | 1.41 | 0.05 |
| Phaeo | Genus | 158444.6 | 98241.5 | 0.5 | 0.3 | 10.8 | 96.9 | 81.4 | 0 | 0.43 | 0.29 |
| Chitino | Family | 53718.7 | 11761.9 | 0.2 | 0 | 30.7 | 0 | 67.7 | 55.4 | 0.16 | 0.02 |
| 37-13 | Family | 8847.7 | 12858.8 | 0 | 0 | 0 | 0 | 0 | 0 | 0 | 0 |
| 37-13_ | Genus | 8847.7 | 12858.8 | 0 | 0 | 0 | 0 | 0 | 0 | 0 | 0 |
| Sphingol | Order | 265521.8 | 204405.5 | 0.8 | 0.6 | 26.9 | 7.9 | 25.8 | 17.4 | 0.41 | 0.15 |
| KD3-93 | Family | 142905.6 | 80161.4 | 0.4 | 0.2 | 27.6 | 0 | 46.7 | 39.3 | 0.31 | 0.09 |
| KD3-9 | Genus | 142905.6 | 80161.4 | 0.4 | 0.2 | 27.6 | 0 | 46.7 | 39.3 | 0.31 | 0.09 |
| AKYH76 | Family | 100289.5 | 95556 | 0.3 | 0.3 | 32 | 16.8 | 0 | 4.2 | 0.09 | 0.06 |
| AKYH7 | Genus | 100289.5 | 95556 | 0.3 | 0.3 | 32 | 16.8 | 0 | 4.2 | 0.09 | 0.06 |
| env.OP | Family | 15666.6 | 15897.2 | 0 | 0 | 0 | 0 | 10.9 | 0 | 0.01 | 0 |
| env.OF | Genus | 15666.6 | 15897.2 | 0 | 0 | 0 | 0 | 10.9 | 0 | 0.01 | 0 |
| LiUU-11 | Family | 4899.9 | 8855.3 | 0 | 0 | 0 | 0 | 0 | 0 | 0 | 0 |
| LiUU-1 | Genus | 4899.9 | 8855.3 | 0 | 0 | 0 | 0 | 0 | 0 | 0 | 0 |
| Sphing | Family | 1760.1 | 3935.5 | 0 | 0 | 0 | 0 | 0 | 0 | 0 | 0 |
| Sphing | Genus | 1760.1 | 3935.5 | 0 | 0 | 0 | 0 | 0 | 0 | 0 | 0 |
| Cytopha | Order | 143028.6 | 148941.8 | 0.4 | 0.4 | 2.9 | 0 | 8.1 | 3.4 | 0.05 | 0.01 |
| Microsc | Family | 113773.1 | 120833.5 | 0.3 | 0.4 | 3.6 | 0 | 10.1 | 4.1 | 0.05 | 0.01 |
| Micros | Genus | 63817.5 | 73920.2 | 0.2 | 0.2 | 6.4 | 0 | 6.9 | 6.8 | 0.03 | 0.01 |
| Ohtael | Genus | 33057.4 | 31294.1 | 0.1 | 0.1 | 0 | 0 | 21.6 | 0 | 0.02 | 0 |
| Chryse | Genus | 16530 | 15298.1 | 0 | 0 | 0 | 0 | 0 | 0 | 0 | 0 |
| Hassal | Genus | 368.2 | 321.1 | 0 | 0 | 0 | 0 | 0 | 0 | 0 | 0 |
| Hymen | Family | 28953.2 | 28108.3 | 0.1 | 0.1 | 0 | 0 | 0 | 0 | 0 | 0 |
| Adhae | Genus | 28953.2 | 28108.3 | 0.1 | 0.1 | 0 | 0 | 0 | 0 | 0 | 0 |
| Cytopha | Family | 302.4 | 0 | 0 | 0 | 0 | 0 | 0 | 0 | 0 | 0 |
| Cytoph | Genus | 302.4 | 0 | 0 | 0 | 0 | 0 | 0 | 0 | 0 | 0 |
| Bacteroi | Order | 1197.2 | 0 | 0 | 0 | 0 | 0 | 0 | 0 | 0 | 0 |

| | | | | | | | | | | | |
|---|---|---|---|---|---|---|---|---|---|---|---|
| Prolixib | Family | 1197.2 | 0 | 0 | 0 | 0 | 0 | 0 | 0 | 0 | 0 |
| WCHB | Genus | 1197.2 | 0 | 0 | 0 | 0 | 0 | 0 | 0 | 0 | 0 |
| Bacteroi | Order | 0 | 1145 | 0 | 0 | 0 | 0 | 0 | 0 | 0 | 0 |
| Bactero | Family | 0 | 1145 | 0 | 0 | 0 | 0 | 0 | 0 | 0 | 0 |
| Ignavibac | Class | 91408.3 | 91233.9 | 0.3 | 0.3 | 0 | 0 | 12.3 | 0 | 0.03 | 0 |
| SJA-28 | Order | 90407.7 | 88795.1 | 0.3 | 0.3 | 0 | 0 | 12.4 | 0 | 0.03 | 0 |
| SJA-28_ | Family | 90407.7 | 88795.1 | 0.3 | 0.3 | 0 | 0 | 12.4 | 0 | 0.03 | 0 |
| Ignaviba | Order | 1000.6 | 2438.7 | 0 | 0 | 0 | 0 | 0 | 0 | 0 | 0 |
| Ignavib | Family | 1000.6 | 2438.7 | 0 | 0 | 0 | 0 | 0 | 0 | 0 | 0 |
| Rhodothe | Class | 48557.6 | 10412.2 | 0.1 | 0 | 0 | 10.9 | 0 | 0 | 0 | 0 |
| Rhodoth | Order | 48557.6 | 10412.2 | 0.1 | 0 | 0 | 10.9 | 0 | 0 | 0 | 0 |
| Rhodotl | Family | 48557.6 | 10412.2 | 0.1 | 0 | 0 | 10.9 | 0 | 0 | 0 | 0 |
| Rhodo | Genus | 48557.6 | 10412.2 | 0.1 | 0 | 0 | 10.9 | 0 | 0 | 0 | 0 |
| Kapabact | Class | 32241.6 | 31676.6 | 0.1 | 0.1 | 15.2 | 3.9 | 20.6 | 0 | 0.03 | 0 |
| Kapabac | Order | 32241.6 | 31676.6 | 0.1 | 0.1 | 15.2 | 3.9 | 20.6 | 0 | 0.03 | 0 |
| Kapaba | Family | 32241.6 | 31676.6 | 0.1 | 0.1 | 15.2 | 3.9 | 20.6 | 0 | 0.03 | 0 |
| Actinobac | Phylum | 2848155 | 5241963 | 8.4 | 15.7 | 7.5 | 0.7 | 8.9 | 3.3 | 1.38 | 0.64 |
| Actinobac | Class | 2062870 | 4445174 | 6.1 | 13.3 | 3.8 | 0.7 | 10.6 | 3.3 | 0.88 | 0.54 |
| Micrococ | Order | 757559 | 669434.9 | 2.2 | 2 | 10.4 | 2.3 | 20.5 | 18.6 | 0.69 | 0.42 |
| Micrococ | Family | 434824.5 | 304772 | 1.3 | 0.9 | 6.3 | 4.6 | 33.6 | 38.6 | 0.51 | 0.39 |
| Pseuda | Genus | 185171.7 | 101464.4 | 0.5 | 0.3 | 0 | 2.2 | 79 | 49 | 0.43 | 0.16 |
| Paenai | Genus | 175101.8 | 168758.1 | 0.5 | 0.5 | 0 | 0 | 0 | 34.7 | 0 | 0.18 |
| Glutan | Genus | 47667.3 | 11211.3 | 0.1 | 0 | 54.9 | 36.8 | 0 | 0 | 0.08 | 0.01 |
| Arthrol | Genus | 15395.9 | 15276.6 | 0 | 0 | 7.8 | 16 | 0 | 61.3 | 0 | 0.04 |
| Microc | Genus | 10301.3 | 7638.4 | 0 | 0 | 0 | 66 | 0 | 0 | 0 | 0.02 |
| Paenig | Genus | 1186.4 | 423.2 | 0 | 0 | 0 | 0 | 0 | 0 | 0 | 0 |
| Microba | Family | 244932.5 | 297561.7 | 0.7 | 0.9 | 3.7 | 0 | 0.8 | 1 | 0.03 | 0.01 |
| Lysinir | Genus | 82040.1 | 59272.8 | 0.2 | 0.2 | 0 | 0 | 2.5 | 0 | 0.01 | 0 |
| Agrom | Genus | 69800 | 88641.1 | 0.2 | 0.3 | 0 | 0 | 0 | 0 | 0 | 0 |
| Microb | Genus | 19355.8 | 18398.2 | 0.1 | 0.1 | 0 | 0 | 0 | 5 | 0 | 0 |
| Homos | Genus | 18824.9 | 55028.9 | 0.1 | 0.2 | 0 | 0 | 0 | 3.7 | 0 | 0.01 |

| | | | | | | | | | | | |
|---|---|---|---|---|---|---|---|---|---|---|---|
| Galbita | Genus | 16052 | 8044.4 | 0 | 0 | 55.7 | 0 | 0 | 0 | 0.03 | 0 |
| Microb | Genus | 11043.6 | 22780.1 | 0 | 0.1 | 0 | 0 | 0 | 0 | 0 | 0 |
| Salinib | Genus | 10366.1 | 21424.9 | 0 | 0.1 | 0 | 0 | 0 | 0 | 0 | 0 |
| Parafri | Genus | 9830.4 | 10799.9 | 0 | 0 | 0 | 0 | 0 | 0 | 0 | 0 |
| Schum | Genus | 3895.6 | 8773.9 | 0 | 0 | 0 | 0 | 0 | 0 | 0 | 0 |
| Conyzi | Genus | 1713.2 | 1869.6 | 0 | 0 | 0 | 0 | 0 | 0 | 0 | 0 |
| Glaciib | Genus | 1169.2 | 1547.5 | 0 | 0 | 0 | 0 | 0 | 0 | 0 | 0 |
| Leifsor | Genus | 841.5 | 980.1 | 0 | 0 | 0 | 0 | 0 | 0 | 0 | 0 |
| Demeq | Family | 44731.9 | 4302.8 | 0.1 | 0 | 94.4 | 30.2 | 0 | 0 | 0.12 | 0 |
| Demec | Genus | 44731.9 | 4302.8 | 0.1 | 0 | 94.4 | 30.2 | 0 | 0 | 0.12 | 0 |
| Promici | Family | 17454.6 | 41274.1 | 0.1 | 0.1 | 0 | 0 | 26.5 | 0 | 0.01 | 0 |
| Promic | Genus | 13718.9 | 32758.5 | 0 | 0.1 | 0 | 0 | 25.3 | 0 | 0.01 | 0 |
| Cellulc | Genus | 3735.7 | 8515.6 | 0 | 0 | 0 | 0 | 30.6 | 0 | 0 | 0 |
| Intraspo | Family | 14413.7 | 20515.5 | 0 | 0.1 | 0 | 0 | 15.2 | 18.5 | 0.01 | 0.01 |
| Knoelli | Genus | 8415.1 | 13244.6 | 0 | 0 | 0 | 0 | 0 | 0 | 0 | 0 |
| Intrasp | Genus | 2231.3 | 1985.5 | 0 | 0 | 0 | 0 | 65.2 | 46.1 | 0 | 0 |
| Lapillic | Genus | 1738.6 | 1354.4 | 0 | 0 | 0 | 0 | 42.2 | 0 | 0 | 0 |
| Terrab. | Genus | 1737.9 | 2888.7 | 0 | 0 | 0 | 0 | 0 | 100 | 0 | 0.01 |
| Oryzihi | Genus | 290.8 | 1042.3 | 0 | 0 | 0 | 0 | 0 | 0 | 0 | 0 |
| Celluloi | Family | 1201.8 | 1008.9 | 0 | 0 | 0 | 0 | 0 | 0 | 0 | 0 |
| Oerskc | Genus | 1201.8 | 1008.9 | 0 | 0 | 0 | 0 | 0 | 0 | 0 | 0 |
| Strepton | Order | 479933.3 | 2596730 | 1.4 | 7.8 | 0 | 0.5 | 1 | 0.3 | 0.01 | 0.06 |
| Strepto | Family | 479933.3 | 2596730 | 1.4 | 7.8 | 0 | 0.5 | 1 | 0.3 | 0.01 | 0.06 |
| Strepto | Genus | 479933.3 | 2596730 | 1.4 | 7.8 | 0 | 0.5 | 1 | 0.3 | 0.01 | 0.06 |
| Micromc | Order | 346058.3 | 549126.2 | 1 | 1.6 | 0 | 0 | 11.9 | 1.7 | 0.12 | 0.03 |
| Microm | Family | 346058.3 | 549126.2 | 1 | 1.6 | 0 | 0 | 11.9 | 1.7 | 0.12 | 0.03 |
| Micron | Genus | 197354.8 | 375989.8 | 0.6 | 1.1 | 0 | 0 | 16.4 | 0.6 | 0.1 | 0.01 |
| Luedei | Genus | 47317.5 | 49611.8 | 0.1 | 0.1 | 0 | 0 | 0 | 0 | 0 | 0 |
| Pilimel | Genus | 42309.4 | 64043.9 | 0.1 | 0.2 | 0 | 0 | 0 | 0 | 0 | 0 |
| Micron | Genus | 21822.1 | 31377.4 | 0.1 | 0.1 | 0 | 0 | 0 | 0 | 0 | 0 |
| Phytoh | Genus | 13957 | 11405.7 | 0 | 0 | 0 | 0 | 28 | 15.9 | 0.01 | 0.01 |

| | | | | | | | | | | | |
|---|---|---|---|---|---|---|---|---|---|---|---|
| Spirilli | Genus | 10121 | 982.2 | 0 | 0 | 0 | 0 | 46.7 | 0 | 0.01 | 0 |
| Planta | Genus | 8109.2 | 11663.5 | 0 | 0 | 0 | 0 | 0 | 43.8 | 0 | 0.02 |
| Longis | Genus | 2114.2 | 163.1 | 0 | 0 | 0 | 0 | 0 | 0 | 0 | 0 |
| Asano | Genus | 1525.1 | 2710.8 | 0 | 0 | 0 | 0 | 0 | 0 | 0 | 0 |
| Planos | Genus | 811.3 | 81.6 | 0 | 0 | 0 | 0 | 0 | 0 | 0 | 0 |
| Dactyl | Genus | 293.5 | 1096.4 | 0 | 0 | 0 | 0 | 0 | 0 | 0 | 0 |
| Catella | Genus | 226.9 | 0 | 0 | 0 | 0 | 0 | 0 | 0 | 0 | 0 |
| Actino | Genus | 96.5 | 0 | 0 | 0 | 0 | 0 | 0 | 0 | 0 | 0 |
| Propioni | Order | 265297.4 | 340636 | 0.8 | 1 | 0 | 0.8 | 2.5 | 1.1 | 0.02 | 0.02 |
| Nocard | Family | 259786 | 332195 | 0.8 | 1 | 0 | 0.8 | 2.6 | 1.1 | 0.02 | 0.02 |
| Nocar | Genus | 173286.9 | 219379.4 | 0.5 | 0.7 | 0 | 0 | 3.2 | 0 | 0.02 | 0 |
| Kribbe | Genus | 55839.9 | 75727.1 | 0.2 | 0.2 | 0 | 0 | 1.9 | 0 | 0 | 0 |
| Aerom | Genus | 19562.6 | 23498.5 | 0.1 | 0.1 | 0 | 11.6 | 0 | 15.5 | 0 | 0.02 |
| Marmo | Genus | 10610.2 | 12982.5 | 0 | 0 | 0 | 0 | 0 | 0 | 0 | 0 |
| Nocar | Genus | 486.4 | 607.6 | 0 | 0 | 0 | 0 | 0 | 0 | 0 | 0 |
| Propion | Family | 5511.3 | 8440.9 | 0 | 0 | 0 | 0 | 0 | 0 | 0 | 0 |
| Jiangel | Genus | 3842.4 | 5220.5 | 0 | 0 | 0 | 0 | 0 | 0 | 0 | 0 |
| Haloa | Genus | 1273 | 2444.9 | 0 | 0 | 0 | 0 | 0 | 0 | 0 | 0 |
| Propio | Genus | 299 | 664.1 | 0 | 0 | 0 | 0 | 0 | 0 | 0 | 0 |
| Microl | Genus | 96.9 | 111.5 | 0 | 0 | 0 | 0 | 0 | 0 | 0 | 0 |
| Streptos | Order | 79515.3 | 118028 | 0.2 | 0.4 | 0 | 0 | 4.1 | 1.3 | 0.01 | 0 |
| Strepto | Family | 61341.2 | 90793 | 0.2 | 0.3 | 0 | 0 | 0 | 1.7 | 0 | 0 |
| Nonon | Genus | 50095.9 | 74577.3 | 0.1 | 0.2 | 0 | 0 | 0 | 0 | 0 | 0 |
| Therm | Genus | 5837.1 | 9394.1 | 0 | 0 | 0 | 0 | 0 | 0 | 0 | 0 |
| Sphae | Genus | 2921.6 | 2925.3 | 0 | 0 | 0 | 0 | 0 | 0 | 0 | 0 |
| Strept | Genus | 2292.8 | 3814.7 | 0 | 0 | 0 | 0 | 0 | 40.4 | 0 | 0 |
| Strept | Genus | 193.9 | 81.6 | 0 | 0 | 0 | 0 | 0 | 0 | 0 | 0 |
| Thermo | Family | 16178.2 | 25890.3 | 0 | 0.1 | 0 | 0 | 12.6 | 0 | 0.01 | 0 |
| Actino | Genus | 9528.4 | 12388 | 0 | 0 | 0 | 0 | 0 | 0 | 0 | 0 |
| Therm | Genus | 3867.7 | 6557 | 0 | 0 | 0 | 0 | 52.6 | 0 | 0.01 | 0 |
| Therm | Genus | 2350.5 | 5760.4 | 0 | 0 | 0 | 0 | 0 | 0 | 0 | 0 |

| | | | | | | | | | | | |
|---|---|---|---|---|---|---|---|---|---|---|---|
| Therm | Genus | 431.6 | 1184.7 | 0 | 0 | 0 | 0 | 0 | 0 | 0 | 0 |
| Nocard | Family | 1995.9 | 1344.7 | 0 | 0 | 0 | 0 | 61.7 | 0 | 0 | 0 |
| Therm | Genus | 1995.9 | 1344.7 | 0 | 0 | 0 | 0 | 61.7 | 0 | 0 | 0 |
| Pseudor | Order | 61349 | 84426.5 | 0.2 | 0.3 | 0 | 0 | 0 | 0 | 0 | 0 |
| Pseudo | Family | 61349 | 84426.5 | 0.2 | 0.3 | 0 | 0 | 0 | 0 | 0 | 0 |
| Pseud | Genus | 58714.8 | 81366.1 | 0.2 | 0.2 | 0 | 0 | 0 | 0 | 0 | 0 |
| Saccha | Genus | 1413.5 | 672.4 | 0 | 0 | 0 | 0 | 0 | 0 | 0 | 0 |
| Actino | Genus | 1220.7 | 2387.9 | 0 | 0 | 0 | 0 | 0 | 0 | 0 | 0 |
| Frankial | Order | 50870.7 | 52225.1 | 0.2 | 0.2 | 0 | 0 | 5.2 | 0 | 0.01 | 0 |
| Geoder | Family | 40135.5 | 43083.7 | 0.1 | 0.1 | 0 | 0 | 0 | 0 | 0 | 0 |
| Blasto | Genus | 39356.2 | 41810.7 | 0.1 | 0.1 | 0 | 0 | 0 | 0 | 0 | 0 |
| Geode | Genus | 779.2 | 1273 | 0 | 0 | 0 | 0 | 0 | 0 | 0 | 0 |
| Sporich | Family | 5118 | 7724.4 | 0 | 0 | 0 | 0 | 0 | 0 | 0 | 0 |
| Sporic | Genus | 5118 | 7724.4 | 0 | 0 | 0 | 0 | 0 | 0 | 0 | 0 |
| Acidoth | Family | 4821.8 | 1038.4 | 0 | 0 | 0 | 0 | 54.7 | 0 | 0.01 | 0 |
| Acidot | Genus | 4821.8 | 1038.4 | 0 | 0 | 0 | 0 | 54.7 | 0 | 0.01 | 0 |
| Frankia | Family | 795.5 | 378.5 | 0 | 0 | 0 | 0 | 0 | 0 | 0 | 0 |
| Jatropl | Genus | 795.5 | 378.5 | 0 | 0 | 0 | 0 | 0 | 0 | 0 | 0 |
| Coryneb | Order | 22287.4 | 34567.1 | 0.1 | 0.1 | 0 | 0 | 19.9 | 8.5 | 0.01 | 0.01 |
| Mycoba | Family | 21668 | 33904.4 | 0.1 | 0.1 | 0 | 0 | 20.4 | 8.6 | 0.01 | 0.01 |
| Mycob | Genus | 21668 | 33904.4 | 0.1 | 0.1 | 0 | 0 | 20.4 | 8.6 | 0.01 | 0.01 |
| Nocard | Family | 619.3 | 662.7 | 0 | 0 | 0 | 0 | 0 | 0 | 0 | 0 |
| Nocard | Genus | 328.6 | 119.9 | 0 | 0 | 0 | 0 | 0 | 0 | 0 | 0 |
| Rhodo | Genus | 290.7 | 542.8 | 0 | 0 | 0 | 0 | 0 | 0 | 0 | 0 |
| Thermole | Class | 402342.3 | 388700 | 1.2 | 1.2 | 34.1 | 0 | 3.7 | 10 | 0.45 | 0.12 |
| Solirubro | Order | 368588.9 | 344319.2 | 1.1 | 1 | 37.3 | 0 | 4.1 | 11 | 0.45 | 0.11 |
| 67-14 | Family | 184930.8 | 171851.6 | 0.5 | 0.5 | 0 | 0 | 7.6 | 19.3 | 0.04 | 0.1 |
| 67-14_ | Genus | 184930.8 | 171851.6 | 0.5 | 0.5 | 0 | 0 | 7.6 | 19.3 | 0.04 | 0.1 |
| Solirubr | Family | 183658.2 | 172467.6 | 0.5 | 0.5 | 74.8 | 0 | 0.5 | 2.7 | 0.41 | 0.01 |
| Solirub | Genus | 172514.7 | 162358.9 | 0.5 | 0.5 | 79.6 | 0 | 0.6 | 2.9 | 0.41 | 0.01 |
| JCM 18 | Genus | 4311.4 | 5044.6 | 0 | 0 | 0 | 0 | 0 | 0 | 0 | 0 |

| | | | | | | | | | | | |
|---|---|---|---|---|---|---|---|---|---|---|---|
| Solirut | Genus | 3491.3 | 1559.4 | 0 | 0 | 0 | 0 | 0 | 0 | 0 | 0 |
| Conexi | Genus | 3340.7 | 3504.8 | 0 | 0 | 0 | 0 | 0 | 0 | 0 | 0 |
| Gaiellale | Order | 33753.4 | 44380.8 | 0.1 | 0.1 | 0 | 0 | 0 | 2.5 | 0 | 0 |
| Gaiellac | Family | 21378.8 | 24380.2 | 0.1 | 0.1 | 0 | 0 | 0 | 4.5 | 0 | 0 |
| Gaiella | Genus | 21378.8 | 24380.2 | 0.1 | 0.1 | 0 | 0 | 0 | 4.5 | 0 | 0 |
| Gaiellal | Family | 12374.5 | 20000.6 | 0 | 0.1 | 0 | 0 | 0 | 0 | 0 | 0 |
| Acidimicr | Class | 380680.3 | 406889.9 | 1.1 | 1.2 | 0 | 2.3 | 7.8 | 0.3 | 0.09 | 0.03 |
| Microtric | Order | 216768.5 | 228695.6 | 0.6 | 0.7 | 0 | 4 | 9.7 | 0 | 0.06 | 0.03 |
| Iamiace | Family | 86730.4 | 88974.2 | 0.3 | 0.3 | 0 | 0 | 8 | 0 | 0.02 | 0 |
| Iamia | Genus | 86730.4 | 88974.2 | 0.3 | 0.3 | 0 | 0 | 8 | 0 | 0.02 | 0 |
| Ilumato | Family | 85904.3 | 83772.2 | 0.3 | 0.3 | 0 | 11 | 9.7 | 0 | 0.02 | 0.03 |
| Ilumat | Genus | 65086.8 | 65286.2 | 0.2 | 0.2 | 0 | 14.1 | 2.1 | 0 | 0 | 0.03 |
| Ilumat | Genus | 10559.4 | 12225.4 | 0 | 0 | 0 | 0 | 65.7 | 0 | 0.02 | 0 |
| CL500 | Genus | 10258 | 6260.6 | 0 | 0 | 0 | 0 | 0 | 0 | 0 | 0 |
| Microtri | Family | 33229.2 | 36090.6 | 0.1 | 0.1 | 0 | 0 | 17.3 | 0 | 0.02 | 0 |
| Microtri | Family | 10904.6 | 19858.7 | 0 | 0.1 | 0 | 0 | 0 | 0 | 0 | 0 |
| Microt | Genus | 8882.6 | 13567.4 | 0 | 0 | 0 | 0 | 0 | 0 | 0 | 0 |
| IMCC2 | Genus | 2022 | 6291.3 | 0 | 0 | 0 | 0 | 0 | 0 | 0 | 0 |
| IMCC26 | Order | 123865.4 | 155738.5 | 0.4 | 0.5 | 0 | 0 | 7 | 0 | 0.03 | 0 |
| IMCC26 | Family | 123865.4 | 155738.5 | 0.4 | 0.5 | 0 | 0 | 7 | 0 | 0.03 | 0 |
| Acidimic | Order | 40046.4 | 22455.8 | 0.1 | 0.1 | 0 | 0 | 0 | 5.1 | 0 | 0 |
| Coriobac | Class | 1777.9 | 0 | 0 | 0 | 0 | 0 | 0 | 0 | 0 | 0 |
| OPB41 | Order | 1777.9 | 0 | 0 | 0 | 0 | 0 | 0 | 0 | 0 | 0 |
| OPB41_ | Family | 1777.9 | 0 | 0 | 0 | 0 | 0 | 0 | 0 | 0 | 0 |
| MB-A2-1( | Class | 483.7 | 1199.1 | 0 | 0 | 0 | 0 | 0 | 0 | 0 | 0 |
| MB-A2-1 | Order | 483.7 | 1199.1 | 0 | 0 | 0 | 0 | 0 | 0 | 0 | 0 |
| Firmicutes | Phylum | 1900728 | 1752401 | 5.6 | 5.3 | 33.4 | 19 | 0.2 | 0.6 | 1.89 | 1.03 |
| Bacilli | Class | 1152124 | 1680112 | 3.4 | 5 | 12.4 | 19.8 | 0.3 | 0.5 | 0.43 | 1.02 |
| Bacillale | Order | 1051809 | 1421607 | 3.1 | 4.3 | 13.5 | 21 | 0.2 | 0 | 0.43 | 0.89 |
| Bacillac | Family | 946183.9 | 1271541 | 2.8 | 3.8 | 14.9 | 23.4 | 0.3 | 0 | 0.42 | 0.89 |
| Bacillu | Genus | 859130.2 | 1170708 | 2.5 | 3.5 | 16.4 | 25 | 0 | 0 | 0.42 | 0.88 |

| | | | | | | | | | | | |
|---|---|---|---|---|---|---|---|---|---|---|---|
| Geoba | Genus | 41876.6 | 43008.9 | 0.1 | 0.1 | 0 | 0 | 0 | 0 | 0 | 0 |
| Bacilla | Genus | 18363.8 | 26372.5 | 0.1 | 0.1 | 0 | 22 | 0 | 0 | 0 | 0.02 |
| Fictiba | Genus | 12050.8 | 11311.4 | 0 | 0 | 0 | 0 | 20.1 | 0 | 0.01 | 0 |
| Ureiba | Genus | 9502.6 | 14807.5 | 0 | 0 | 0 | 0 | 0 | 0 | 0 | 0 |
| Anoxyb | Genus | 3305.7 | 4018.2 | 0 | 0 | 0 | 0 | 0 | 0 | 0 | 0 |
| Falsiba | Genus | 1254 | 356.3 | 0 | 0 | 0 | 0 | 0 | 0 | 0 | 0 |
| Virgiba | Genus | 700.1 | 957.4 | 0 | 0 | 0 | 0 | 0 | 0 | 0 | 0 |
| Planocc | Family | 105625.1 | 150066.2 | 0.3 | 0.4 | 1.8 | 0 | 0 | 0 | 0.01 | 0 |
| Paenis | Genus | 53432.6 | 79905.7 | 0.2 | 0.2 | 3.6 | 0 | 0 | 0 | 0.01 | 0 |
| Lysinib | Genus | 25989.4 | 37291.4 | 0.1 | 0.1 | 0 | 0 | 0 | 0 | 0 | 0 |
| Chung | Genus | 7690.4 | 16799.3 | 0 | 0.1 | 0 | 0 | 0 | 0 | 0 | 0 |
| Planor | Genus | 6002.6 | 0 | 0 | 0 | 0 | 0 | 0 | 0 | 0 | 0 |
| Planoc | Genus | 3648.4 | 7978.4 | 0 | 0 | 0 | 0 | 0 | 0 | 0 | 0 |
| Sporos | Genus | 3341.4 | 6847.5 | 0 | 0 | 0 | 0 | 0 | 0 | 0 | 0 |
| Domib | Genus | 3089.2 | 922.4 | 0 | 0 | 0 | 0 | 0 | 0 | 0 | 0 |
| Psychr | Genus | 2431.2 | 321.4 | 0 | 0 | 0 | 0 | 0 | 0 | 0 | 0 |
| Paeniba | Order | 50699.2 | 150580.3 | 0.1 | 0.5 | 0 | 1.1 | 2.9 | 1.2 | 0 | 0.01 |
| Paeniba | Family | 50699.2 | 150580.3 | 0.1 | 0.5 | 0 | 1.1 | 2.9 | 1.2 | 0 | 0.01 |
| Paenib | Genus | 38992.8 | 131779.3 | 0.1 | 0.4 | 0 | 1.2 | 0 | 1.4 | 0 | 0.01 |
| Ammo | Genus | 11074.4 | 15780.5 | 0 | 0 | 0 | 0 | 13 | 0 | 0 | 0 |
| Cohne | Genus | 631.9 | 3020.4 | 0 | 0 | 0 | 0 | 0 | 0 | 0 | 0 |
| Thermoa | Order | 35492.6 | 57035.2 | 0.1 | 0.2 | 0 | 0 | 0 | 10.1 | 0 | 0.02 |
| Thermo | Family | 35492.6 | 57035.2 | 0.1 | 0.2 | 0 | 0 | 0 | 10.1 | 0 | 0.02 |
| Hazen | Genus | 16525.1 | 29443.8 | 0 | 0.1 | 0 | 0 | 0 | 19.6 | 0 | 0.02 |
| Laceye | Genus | 5916.7 | 11297.8 | 0 | 0 | 0 | 0 | 0 | 0 | 0 | 0 |
| Planifil | Genus | 4687.8 | 5004.8 | 0 | 0 | 0 | 0 | 0 | 0 | 0 | 0 |
| Shima | Genus | 4600.8 | 5918.8 | 0 | 0 | 0 | 0 | 0 | 0 | 0 | 0 |
| Therm | Genus | 3762.2 | 4718 | 0 | 0 | 0 | 0 | 0 | 0 | 0 | 0 |
| Therm | Genus | 0 | 652 | 0 | 0 | 0 | 0 | 0 | 0 | 0 | 0 |
| Aneurini | Order | 6547.6 | 7654.4 | 0 | 0 | 0 | 0 | 0 | 0 | 0 | 0 |
| Aneurin | Family | 6547.6 | 7654.4 | 0 | 0 | 0 | 0 | 0 | 0 | 0 | 0 |

| | | | | | | | | | | | |
|---|---|---|---|---|---|---|---|---|---|---|---|
| Aneuri | Genus | 6547.6 | 7654.4 | 0 | 0 | 0 | 0 | 0 | 0 | 0 | 0 |
| Brevibac | Order | 3280.2 | 8313.8 | 0 | 0 | 0 | 0 | 0 | 0 | 0 | 0 |
| Breviba | Family | 3280.2 | 8313.8 | 0 | 0 | 0 | 0 | 0 | 0 | 0 | 0 |
| Brevib | Genus | 3280.2 | 8313.8 | 0 | 0 | 0 | 0 | 0 | 0 | 0 | 0 |
| Exiguoba | Order | 2033.4 | 1425.3 | 0 | 0 | 0 | 0 | 0 | 0 | 0 | 0 |
| Exiguob | Family | 2033.4 | 1425.3 | 0 | 0 | 0 | 0 | 0 | 0 | 0 | 0 |
| Exiguo | Genus | 2033.4 | 1425.3 | 0 | 0 | 0 | 0 | 0 | 0 | 0 | 0 |
| Lactoba | Order | 1169 | 33376.4 | 0 | 0.1 | 0 | 100 | 0 | 0 | 0 | 0.1 |
| Carnob | Family | 1169 | 33376.4 | 0 | 0.1 | 0 | 100 | 0 | 0 | 0 | 0.1 |
| Tricho | Genus | 1169 | 33376.4 | 0 | 0.1 | 0 | 100 | 0 | 0 | 0 | 0.1 |
| Alicyclob | Order | 1092.7 | 119.9 | 0 | 0 | 0 | 0 | 0 | 0 | 0 | 0 |
| Alicyclo | Family | 1092.7 | 119.9 | 0 | 0 | 0 | 0 | 0 | 0 | 0 | 0 |
| Tumeb | Genus | 1092.7 | 119.9 | 0 | 0 | 0 | 0 | 0 | 0 | 0 | 0 |
| Clostridia | Class | 744767.4 | 66453.3 | 2.2 | 0.2 | 67.2 | 0 | 0 | 4.4 | 1.48 | 0.01 |
| Peptostr | Order | 529789.4 | 52879 | 1.6 | 0.2 | 64.2 | 0 | 0 | 0 | 1.01 | 0 |
| Peptost | Family | 521420.4 | 52755.5 | 1.5 | 0.2 | 65.2 | 0 | 0 | 0 | 1.01 | 0 |
| Sporac | Genus | 411957.4 | 44763.7 | 1.2 | 0.1 | 82.6 | 0 | 0 | 0 | 1.01 | 0 |
| Rombo | Genus | 109463 | 7991.8 | 0.3 | 0 | 0 | 0 | 0 | 0 | 0 | 0 |
| Anaerov | Family | 8368.9 | 123.4 | 0 | 0 | 0 | 0 | 0 | 0 | 0 | 0 |
| Anaero | Genus | 8368.9 | 123.4 | 0 | 0 | 0 | 0 | 0 | 0 | 0 | 0 |
| Clostridi | Order | 213552.8 | 13574.4 | 0.6 | 0 | 74.9 | 0 | 0 | 21.3 | 0.47 | 0.01 |
| Clostric | Family | 213552.8 | 13574.4 | 0.6 | 0 | 74.9 | 0 | 0 | 21.3 | 0.47 | 0.01 |
| Clostri | Genus | 184241.6 | 6249.6 | 0.5 | 0 | 75.6 | 0 | 0 | 0 | 0.41 | 0 |
| Clostri | Genus | 16184.4 | 611.4 | 0 | 0 | 100 | 0 | 0 | 0 | 0.05 | 0 |
| Clostri | Genus | 6844.2 | 1989.7 | 0 | 0 | 66.6 | 0 | 0 | 0 | 0.01 | 0 |
| Clostri | Genus | 5509.2 | 4628.2 | 0 | 0 | 0 | 0 | 0 | 62.5 | 0 | 0.01 |
| Clostri | Genus | 773.5 | 95.5 | 0 | 0 | 0 | 0 | 0 | 0 | 0 | 0 |
| Graciliba | Order | 1057 | 0 | 0 | 0 | 0 | 0 | 0 | 0 | 0 | 0 |
| Lutispo | Family | 1057 | 0 | 0 | 0 | 0 | 0 | 0 | 0 | 0 | 0 |
| Lutisp | Genus | 1057 | 0 | 0 | 0 | 0 | 0 | 0 | 0 | 0 | 0 |
| Christen | Order | 368.2 | 0 | 0 | 0 | 0 | 0 | 0 | 0 | 0 | 0 |

| | | | | | | | | | | | |
|---|---|---|---|---|---|---|---|---|---|---|---|
| Christe | Family | 368.2 | 0 | 0 | 0 | 0 | 0 | 0 | 0 | 0 | 0 |
| Christe | Genus | 368.2 | 0 | 0 | 0 | 0 | 0 | 0 | 0 | 0 | 0 |
| Symbioba | Class | 3837.1 | 5836 | 0 | 0 | 0 | 0 | 0 | 0 | 0 | 0 |
| Symbiob | Order | 3837.1 | 5836 | 0 | 0 | 0 | 0 | 0 | 0 | 0 | 0 |
| Symbio | Family | 3837.1 | 5836 | 0 | 0 | 0 | 0 | 0 | 0 | 0 | 0 |
| Symbi | Genus | 3837.1 | 5836 | 0 | 0 | 0 | 0 | 0 | 0 | 0 | 0 |
| Gemmatin | Phylum | 846818.6 | 849987.7 | 2.5 | 2.5 | 1.6 | 1 | 8.4 | 17.1 | 0.25 | 0.46 |
| Gemmati | Class | 545552.1 | 566926.4 | 1.6 | 1.7 | 0.9 | 0.9 | 12.6 | 11.8 | 0.22 | 0.22 |
| Gemmat | Order | 545552.1 | 566926.4 | 1.6 | 1.7 | 0.9 | 0.9 | 12.6 | 11.8 | 0.22 | 0.22 |
| Gemma | Family | 545552.1 | 566926.4 | 1.6 | 1.7 | 0.9 | 0.9 | 12.6 | 11.8 | 0.22 | 0.22 |
| Gemm | Genus | 482223.9 | 494897.4 | 1.4 | 1.5 | 0.6 | 0.6 | 14.3 | 11.5 | 0.21 | 0.18 |
| Gemm | Genus | 63328.2 | 72029 | 0.2 | 0.2 | 3.6 | 2.9 | 0 | 14.2 | 0.01 | 0.04 |
| Longimic | Class | 255859.6 | 238049.8 | 0.8 | 0.7 | 3.9 | 1.1 | 0 | 33.7 | 0.03 | 0.25 |
| Longimi | Order | 255859.6 | 238049.8 | 0.8 | 0.7 | 3.9 | 1.1 | 0 | 33.7 | 0.03 | 0.25 |
| Longim | Family | 255859.6 | 238049.8 | 0.8 | 0.7 | 3.9 | 1.1 | 0 | 33.7 | 0.03 | 0.25 |
| Longin | Genus | 248187.8 | 236087.8 | 0.7 | 0.7 | 1.9 | 0.5 | 0 | 34 | 0.01 | 0.24 |
| YC-ZS | Genus | 7671.9 | 1962 | 0 | 0 | 67 | 66.1 | 0 | 0 | 0.02 | 0 |
| S0134 ter | Class | 29031.6 | 23613.1 | 0.1 | 0.1 | 0 | 0 | 5.4 | 0 | 0 | 0 |
| S0134 te | Order | 29031.6 | 23613.1 | 0.1 | 0.1 | 0 | 0 | 5.4 | 0 | 0 | 0 |
| BD2-11 te | Class | 12422.5 | 16299.9 | 0 | 0 | 0 | 24.1 | 13.8 | 0 | 0.01 | 0.01 |
| BD2-11 t | Order | 12422.5 | 16299.9 | 0 | 0 | 0 | 24.1 | 13.8 | 0 | 0.01 | 0.01 |
| AKAU404 | Class | 3758.9 | 5003.1 | 0 | 0 | 0 | 0 | 0 | 0 | 0 | 0 |
| AKAU40 | Order | 3758.9 | 5003.1 | 0 | 0 | 0 | 0 | 0 | 0 | 0 | 0 |
| Gemmati | Class | 193.9 | 95.5 | 0 | 0 | 0 | 0 | 0 | 0 | 0 | 0 |
| Acidobact | Phylum | 694115.8 | 639451 | 2.1 | 1.9 | 52.6 | 35.9 | 1 | 0.4 | 1.1 | 0.7 |
| Acidobac | Class | 574163.1 | 500139.2 | 1.7 | 1.5 | 62.7 | 45.3 | 1.2 | 0 | 1.08 | 0.68 |
| Bryobact | Order | 312956.1 | 239090.4 | 0.9 | 0.7 | 84.2 | 68.2 | 1.2 | 0 | 0.79 | 0.49 |
| Bryobac | Family | 312956.1 | 239090.4 | 0.9 | 0.7 | 84.2 | 68.2 | 1.2 | 0 | 0.79 | 0.49 |
| Bryoba | Genus | 312956.1 | 239090.4 | 0.9 | 0.7 | 84.2 | 68.2 | 1.2 | 0 | 0.79 | 0.49 |
| Paludiba | Order | 124916.3 | 101775 | 0.4 | 0.3 | 76.1 | 62.5 | 0 | 0 | 0.28 | 0.19 |
| Paludib | Family | 124916.3 | 101775 | 0.4 | 0.3 | 76.1 | 62.5 | 0 | 0 | 0.28 | 0.19 |

| | | | | | | | | | | | |
|---|---|---|---|---|---|---|---|---|---|---|---|
| PAUC26 | Order | 115272 | 133966.1 | 0.3 | 0.4 | 0 | 0 | 0 | 0 | 0 | 0 |
| PAUC2( | Family | 115272 | 133966.1 | 0.3 | 0.4 | 0 | 0 | 0 | 0 | 0 | 0 |
| Solibact | Order | 19457.3 | 19868.2 | 0.1 | 0.1 | 6.6 | 0 | 16.9 | 0 | 0.01 | 0 |
| Solibac | Family | 19457.3 | 19868.2 | 0.1 | 0.1 | 6.6 | 0 | 16.9 | 0 | 0.01 | 0 |
| Candic | Genus | 19457.3 | 19868.2 | 0.1 | 0.1 | 6.6 | 0 | 16.9 | 0 | 0.01 | 0 |
| Acidoba | Order | 1461.8 | 5032.3 | 0 | 0 | 0 | 0 | 0 | 0 | 0 | 0 |
| Acidoba | Family | 1461.8 | 5032.3 | 0 | 0 | 0 | 0 | 0 | 0 | 0 | 0 |
| Acidoba | Order | 99.7 | 407.3 | 0 | 0 | 0 | 0 | 0 | 0 | 0 | 0 |
| Thermoar | Class | 94450.1 | 118371.5 | 0.3 | 0.4 | 9 | 2.4 | 0 | 2.4 | 0.03 | 0.02 |
| Thermoa | Order | 94450.1 | 118371.5 | 0.3 | 0.4 | 9 | 2.4 | 0 | 2.4 | 0.03 | 0.02 |
| Thermo | Family | 94450.1 | 118371.5 | 0.3 | 0.4 | 9 | 2.4 | 0 | 2.4 | 0.03 | 0.02 |
| Subgrc | Genus | 94450.1 | 118371.5 | 0.3 | 0.4 | 9 | 2.4 | 0 | 2.4 | 0.03 | 0.02 |
| Holophag | Class | 15731.8 | 8158.8 | 0 | 0 | 19.7 | 0 | 0 | 0 | 0.01 | 0 |
| Subgrou | Order | 15731.8 | 8158.8 | 0 | 0 | 19.7 | 0 | 0 | 0 | 0.01 | 0 |
| Subgrou | Family | 15731.8 | 8158.8 | 0 | 0 | 19.7 | 0 | 0 | 0 | 0.01 | 0 |
| Subgroup | Class | 9770.8 | 12781.4 | 0 | 0 | 0 | 0 | 0 | 0 | 0 | 0 |
| Subgrou | Order | 9770.8 | 12781.4 | 0 | 0 | 0 | 0 | 0 | 0 | 0 | 0 |
| Myxococc | Phylum | 440834 | 489657.6 | 1.3 | 1.5 | 12.5 | 16.9 | 13.8 | 3.3 | 0.34 | 0.3 |
| Polyangia | Class | 405212.7 | 442552 | 1.2 | 1.3 | 13.6 | 18.2 | 15 | 4.8 | 0.34 | 0.3 |
| Polyangi | Order | 276487.4 | 266047.2 | 0.8 | 0.8 | 12 | 13.2 | 10.4 | 7.9 | 0.18 | 0.17 |
| BIrii41 | Family | 174563 | 171834 | 0.5 | 0.5 | 15.9 | 20.5 | 10.3 | 11.6 | 0.14 | 0.17 |
| BIrii41_ | Genus | 174563 | 171834 | 0.5 | 0.5 | 15.9 | 20.5 | 10.3 | 11.6 | 0.14 | 0.17 |
| Polyang | Family | 56247.1 | 45936 | 0.2 | 0.1 | 9.4 | 0 | 0 | 0 | 0.02 | 0 |
| Pajaro | Genus | 46376.8 | 41177.1 | 0.1 | 0.1 | 0 | 0 | 0 | 0 | 0 | 0 |
| Polyan | Genus | 5561.1 | 2370.7 | 0 | 0 | 59.7 | 0 | 0 | 0 | 0.01 | 0 |
| Polyan | Genus | 4309.2 | 2388.2 | 0 | 0 | 46.2 | 0 | 0 | 0 | 0.01 | 0 |
| Phaseli | Family | 30541.8 | 25511.6 | 0.1 | 0.1 | 0 | 0 | 17.2 | 4.4 | 0.02 | 0 |
| Phasel | Genus | 30541.8 | 25511.6 | 0.1 | 0.1 | 0 | 0 | 17.2 | 4.4 | 0.02 | 0 |
| Sandara | Family | 15135.5 | 22765.6 | 0 | 0.1 | 0 | 0 | 37.5 | 0 | 0.02 | 0 |
| Sandar | Genus | 13687.2 | 21486.7 | 0 | 0.1 | 0 | 0 | 41.5 | 0 | 0.02 | 0 |
| Sandar | Genus | 1448.3 | 1278.9 | 0 | 0 | 0 | 0 | 0 | 0 | 0 | 0 |

| | | | | | | | | | | | |
|---|---|---|---|---|---|---|---|---|---|---|---|
| Haliangi | Order | 68840.7 | 85569 | 0.2 | 0.3 | 0 | 0 | 5 | 0 | 0.01 | 0 |
| Haliang | Family | 68840.7 | 85569 | 0.2 | 0.3 | 0 | 0 | 5 | 0 | 0.01 | 0 |
| Halian | Genus | 68840.7 | 85569 | 0.2 | 0.3 | 0 | 0 | 5 | 0 | 0.01 | 0 |
| Blfdi19 | Order | 52119.2 | 79203.1 | 0.2 | 0.2 | 42.4 | 57 | 46.7 | 0 | 0.14 | 0.14 |
| Blfdi19_ | Family | 52119.2 | 79203.1 | 0.2 | 0.2 | 42.4 | 57 | 46.7 | 0 | 0.14 | 0.14 |
| mle1-27 | Order | 3916.1 | 439.2 | 0 | 0 | 0 | 0 | 100 | 0 | 0.01 | 0 |
| mle1-2 | Family | 3916.1 | 439.2 | 0 | 0 | 0 | 0 | 100 | 0 | 0.01 | 0 |
| Nannocy | Order | 3849.3 | 11293.4 | 0 | 0 | 0 | 0 | 0 | 0 | 0 | 0 |
| Nannoc | Family | 3849.3 | 11293.4 | 0 | 0 | 0 | 0 | 0 | 0 | 0 | 0 |
| Nanno | Genus | 3849.3 | 11293.4 | 0 | 0 | 0 | 0 | 0 | 0 | 0 | 0 |
| Myxococ | Class | 34735.5 | 45221.8 | 0.1 | 0.1 | 3.1 | 9.7 | 6 | 0 | 0.01 | 0.01 |
| Myxococ | Order | 34735.5 | 45221.8 | 0.1 | 0.1 | 3.1 | 9.7 | 6 | 0 | 0.01 | 0.01 |
| Myxoco | Family | 21093.9 | 28248.5 | 0.1 | 0.1 | 5.1 | 15.5 | 9.9 | 0 | 0.01 | 0.01 |
| P3OB- | Genus | 11726.4 | 17482.2 | 0 | 0.1 | 0 | 25 | 17.8 | 0 | 0.01 | 0.01 |
| Archar | Genus | 4268 | 2082.2 | 0 | 0 | 0 | 0 | 0 | 0 | 0 | 0 |
| Myxoc | Genus | 3684.6 | 7637.1 | 0 | 0 | 29.5 | 0 | 0 | 0 | 0 | 0 |
| Stigma | Genus | 1414.9 | 1047 | 0 | 0 | 0 | 0 | 0 | 0 | 0 | 0 |
| Vulgatil | Family | 13641.6 | 16973.2 | 0 | 0.1 | 0 | 0 | 0 | 0 | 0 | 0 |
| Vulgati | Genus | 13641.6 | 16973.2 | 0 | 0.1 | 0 | 0 | 0 | 0 | 0 | 0 |
| Myxococ | Class | 885.8 | 1720.7 | 0 | 0 | 0 | 0 | 0 | 0 | 0 | 0 |
| bacteriap | Class | 0 | 163.1 | 0 | 0 | 0 | 0 | 0 | 0 | 0 | 0 |
| bacteria | Order | 0 | 163.1 | 0 | 0 | 0 | 0 | 0 | 0 | 0 | 0 |
| Verrucomi | Phylum | 250414.9 | 290389.1 | 0.7 | 0.9 | 9.6 | 4.4 | 11.8 | 0.9 | 0.16 | 0.05 |
| Verrucom | Class | 187065.7 | 248955.4 | 0.6 | 0.7 | 1.8 | 1.7 | 10.6 | 0 | 0.07 | 0.01 |
| Pedosph | Order | 187065.7 | 248955.4 | 0.6 | 0.7 | 1.8 | 1.7 | 10.6 | 0 | 0.07 | 0.01 |
| Pedosp | Family | 187065.7 | 248955.4 | 0.6 | 0.7 | 1.8 | 1.7 | 10.6 | 0 | 0.07 | 0.01 |
| Pedos | Genus | 168199.5 | 224520.3 | 0.5 | 0.7 | 0 | 1.9 | 11.8 | 0 | 0.06 | 0.01 |
| Ellin51 | Genus | 11438.1 | 18483 | 0 | 0.1 | 0 | 0 | 0 | 0 | 0 | 0 |
| Oikopl | Genus | 3596 | 2058.1 | 0 | 0 | 92.2 | 0 | 0 | 0 | 0.01 | 0 |
| Pedos | Genus | 1957 | 1657.3 | 0 | 0 | 0 | 0 | 0 | 0 | 0 | 0 |
| DEV00 | Genus | 970.9 | 818.8 | 0 | 0 | 0 | 0 | 0 | 0 | 0 | 0 |

| | | | | | | | | | | | |
|---|---|---|---|---|---|---|---|---|---|---|---|
| Ellin51 | Genus | 904.2 | 1417.9 | 0 | 0 | 0 | 0 | 0 | 0 | 0 | 0 |
| Chlamydi | Class | 39205.4 | 27007.6 | 0.1 | 0.1 | 6.8 | 25.6 | 34.7 | 16.2 | 0.05 | 0.03 |
| Chlamyc | Order | 39205.4 | 27007.6 | 0.1 | 0.1 | 6.8 | 25.6 | 34.7 | 16.2 | 0.05 | 0.03 |
| Parachl | Family | 18287.8 | 3769.5 | 0.1 | 0 | 14.6 | 32.4 | 33.1 | 0 | 0.03 | 0 |
| Neoch | Genus | 11824.7 | 1221.6 | 0 | 0 | 0 | 100 | 51.1 | 0 | 0.02 | 0 |
| Paracl | Genus | 5721.1 | 2303.2 | 0 | 0 | 46.7 | 0 | 0 | 0 | 0.01 | 0 |
| Candic | Genus | 741.9 | 244.7 | 0 | 0 | 0 | 0 | 0 | 0 | 0 | 0 |
| cvE6 | Family | 17103.8 | 20729.2 | 0.1 | 0.1 | 0 | 27.5 | 44.1 | 13 | 0.02 | 0.03 |
| cvE6_F | Genus | 17103.8 | 20729.2 | 0.1 | 0.1 | 0 | 27.5 | 44.1 | 13 | 0.02 | 0.03 |
| Simkan | Family | 3813.8 | 2509 | 0 | 0 | 0 | 0 | 0 | 66.9 | 0 | 0.01 |
| Simkai | Genus | 3813.8 | 2509 | 0 | 0 | 0 | 0 | 0 | 66.9 | 0 | 0.01 |
| Kiritimati | Class | 24143.8 | 14426.1 | 0.1 | 0 | 81.3 | 20.1 | 0 | 0 | 0.06 | 0.01 |
| WCHB1- | Order | 24143.8 | 14426.1 | 0.1 | 0 | 81.3 | 20.1 | 0 | 0 | 0.06 | 0.01 |
| WCHB1 | Family | 24143.8 | 14426.1 | 0.1 | 0 | 81.3 | 20.1 | 0 | 0 | 0.06 | 0.01 |
| Desulfoba | Phylum | 227998.6 | 162402.4 | 0.7 | 0.5 | 13.8 | 24.2 | 0 | 0 | 0.09 | 0.12 |
| Desulfurc | Class | 186305.7 | 97515.2 | 0.6 | 0.3 | 16.8 | 0 | 0 | 0 | 0.09 | 0 |
| PB19 | Order | 95018.9 | 93248.1 | 0.3 | 0.3 | 0 | 0 | 0 | 0 | 0 | 0 |
| PB19_C | Family | 95018.9 | 93248.1 | 0.3 | 0.3 | 0 | 0 | 0 | 0 | 0 | 0 |
| Geobact | Order | 90987.8 | 4267.1 | 0.3 | 0 | 34.5 | 0 | 0 | 0 | 0.09 | 0 |
| Geobac | Family | 90987.8 | 4267.1 | 0.3 | 0 | 34.5 | 0 | 0 | 0 | 0.09 | 0 |
| Citrifer | Genus | 89058.2 | 4267.1 | 0.3 | 0 | 35.2 | 0 | 0 | 0 | 0.09 | 0 |
| Geoba | Genus | 1929.5 | 0 | 0 | 0 | 0 | 0 | 0 | 0 | 0 | 0 |
| Bradymc | Order | 299 | 0 | 0 | 0 | 0 | 0 | 0 | 0 | 0 | 0 |
| Bradym | Family | 299 | 0 | 0 | 0 | 0 | 0 | 0 | 0 | 0 | 0 |
| Desulfob | Class | 41692.9 | 64887.2 | 0.1 | 0.2 | 0 | 60.5 | 0 | 0 | 0 | 0.12 |
| Chloroflex | Phylum | 140438.4 | 138580.4 | 0.4 | 0.4 | 1 | 1.8 | 7.6 | 0.9 | 0.04 | 0.01 |
| JG30-KF-( | Class | 59395.9 | 47855.1 | 0.2 | 0.1 | 2.4 | 5.3 | 0 | 2.7 | 0 | 0.01 |
| JG30-KF- | Order | 59395.9 | 47855.1 | 0.2 | 0.1 | 2.4 | 5.3 | 0 | 2.7 | 0 | 0.01 |
| TK10 | Class | 46766.5 | 37566.5 | 0.1 | 0.1 | 0 | 2.5 | 22.8 | 0 | 0.03 | 0 |
| TK10_Cl | Order | 46766.5 | 37566.5 | 0.1 | 0.1 | 0 | 2.5 | 22.8 | 0 | 0.03 | 0 |
| Chlorofle | Class | 24563 | 41540.2 | 0.1 | 0.1 | 0 | 0 | 0 | 2.8 | 0 | 0 |

| | | | | | | | | | | | |
|---|---|---|---|---|---|---|---|---|---|---|---|
| Chloroflo | Order | 20674.1 | 37322.5 | 0.1 | 0.1 | 0 | 0 | 0 | 0 | 0 | 0 |
| Roseifle | Family | 19466.9 | 31376.9 | 0.1 | 0.1 | 0 | 0 | 0 | 0 | 0 | 0 |
| Roseifl | Genus | 19466.9 | 31376.9 | 0.1 | 0.1 | 0 | 0 | 0 | 0 | 0 | 0 |
| Herpeto | Family | 1207.2 | 5945.6 | 0 | 0 | 0 | 0 | 0 | 0 | 0 | 0 |
| Herpet | Genus | 1207.2 | 5945.6 | 0 | 0 | 0 | 0 | 0 | 0 | 0 | 0 |
| Kalloten | Order | 3888.9 | 4217.7 | 0 | 0 | 0 | 0 | 0 | 27.3 | 0 | 0 |
| AKIW78 | Family | 3888.9 | 4217.7 | 0 | 0 | 0 | 0 | 0 | 27.3 | 0 | 0 |
| AKIW7 | Genus | 3888.9 | 4217.7 | 0 | 0 | 0 | 0 | 0 | 27.3 | 0 | 0 |
| Dehaloco | Class | 9317 | 11135.2 | 0 | 0 | 0 | 0 | 0 | 0 | 0 | 0 |
| S085 | Order | 9317 | 11135.2 | 0 | 0 | 0 | 0 | 0 | 0 | 0 | 0 |
| S085_C | Family | 9317 | 11135.2 | 0 | 0 | 0 | 0 | 0 | 0 | 0 | 0 |
| Chlorofle | Class | 396.1 | 483.4 | 0 | 0 | 0 | 0 | 0 | 0 | 0 | 0 |
| Bdellovibri | Phylum | 100763.8 | 98589.2 | 0.3 | 0.3 | 20.3 | 6.5 | 8.9 | 1.9 | 0.09 | 0.02 |
| Bdellovib | Class | 65335.3 | 51323.6 | 0.2 | 0.2 | 25.1 | 12.4 | 13.7 | 0 | 0.07 | 0.02 |
| Bdellovil | Order | 40258.4 | 31639.7 | 0.1 | 0.1 | 35.5 | 13.6 | 6 | 0 | 0.05 | 0.01 |
| Bdellov | Family | 40258.4 | 31639.7 | 0.1 | 0.1 | 35.5 | 13.6 | 6 | 0 | 0.05 | 0.01 |
| Bdello | Genus | 37174.1 | 23105.2 | 0.1 | 0.1 | 38.5 | 18.7 | 6.5 | 0 | 0.05 | 0.01 |
| OM27 | Genus | 3084.4 | 8534.5 | 0 | 0 | 0 | 0 | 0 | 0 | 0 | 0 |
| Bacterio | Order | 25076.8 | 19683.9 | 0.1 | 0.1 | 8.4 | 10.5 | 26.1 | 0 | 0.03 | 0.01 |
| Bacteri | Family | 25076.8 | 19683.9 | 0.1 | 0.1 | 8.4 | 10.5 | 26.1 | 0 | 0.03 | 0.01 |
| Peredi | Genus | 22977.3 | 18859 | 0.1 | 0.1 | 0 | 11 | 28.5 | 0 | 0.02 | 0.01 |
| Bacter | Genus | 2099.5 | 824.9 | 0 | 0 | 100 | 0 | 0 | 0 | 0.01 | 0 |
| Oligoflexi | Class | 35332.1 | 46773 | 0.1 | 0.1 | 11.5 | 3.3 | 0 | 4 | 0.01 | 0.01 |
| 0319-6G | Order | 33474.6 | 44064.6 | 0.1 | 0.1 | 12.1 | 0 | 0 | 4.3 | 0.01 | 0.01 |
| 0319-6 | Family | 33474.6 | 44064.6 | 0.1 | 0.1 | 12.1 | 0 | 0 | 4.3 | 0.01 | 0.01 |
| Oligofle | Order | 1857.5 | 2708.4 | 0 | 0 | 0 | 57.8 | 0 | 0 | 0 | 0 |
| Oligofle | Family | 1857.5 | 2708.4 | 0 | 0 | 0 | 57.8 | 0 | 0 | 0 | 0 |
| Oligofl | Genus | 1857.5 | 2708.4 | 0 | 0 | 0 | 57.8 | 0 | 0 | 0 | 0 |
| Bdellovib | Class | 96.5 | 492.6 | 0 | 0 | 0 | 0 | 0 | 0 | 0 | 0 |
| Planctomy | Phylum | 94743.3 | 61355.6 | 0.3 | 0.2 | 20.5 | 5.9 | 8 | 0 | 0.08 | 0.01 |
| Phycisph | Class | 85192.3 | 53242.4 | 0.3 | 0.2 | 23.9 | 7.1 | 8.8 | 0 | 0.08 | 0.01 |

| | | | | | | | | | | | |
|---|---|---|---|---|---|---|---|---|---|---|---|
| Phycisph | Order | 85192.3 | 53242.4 | 0.3 | 0.2 | 23.9 | 7.1 | 8.8 | 0 | 0.08 | 0.01 |
| Phycisp | Family | 85192.3 | 53242.4 | 0.3 | 0.2 | 23.9 | 7.1 | 8.8 | 0 | 0.08 | 0.01 |
| SM1A0 | Genus | 81967.2 | 49337.4 | 0.2 | 0.1 | 22.7 | 7.7 | 9.2 | 0 | 0.08 | 0.01 |
| Phycis | Genus | 3225.1 | 3905 | 0 | 0 | 56.2 | 0 | 0 | 0 | 0.01 | 0 |
| OM190 | Class | 6349.7 | 2996.4 | 0 | 0 | 0 | 0 | 0 | 0 | 0 | 0 |
| OM190_ | Order | 6349.7 | 2996.4 | 0 | 0 | 0 | 0 | 0 | 0 | 0 | 0 |
| vadinHA4 | Class | 3201.3 | 5116.8 | 0 | 0 | 0 | 37.3 | 0 | 0 | 0 | 0.01 |
| vadinHA | Order | 3201.3 | 5116.8 | 0 | 0 | 0 | 37.3 | 0 | 0 | 0 | 0.01 |
| Armatimo | Phylum | 38458.8 | 33883.8 | 0.1 | 0.1 | 0 | 4.8 | 8.6 | 4.3 | 0.01 | 0.01 |
| Fimbriim | Class | 38458.8 | 33883.8 | 0.1 | 0.1 | 0 | 4.8 | 8.6 | 4.3 | 0.01 | 0.01 |
| Fimbriim | Order | 38458.8 | 33883.8 | 0.1 | 0.1 | 0 | 4.8 | 8.6 | 4.3 | 0.01 | 0.01 |
| Fimbriir | Family | 38458.8 | 33883.8 | 0.1 | 0.1 | 0 | 4.8 | 8.6 | 4.3 | 0.01 | 0.01 |
| Fimbrii | Genus | 38458.8 | 33883.8 | 0.1 | 0.1 | 0 | 4.8 | 8.6 | 4.3 | 0.01 | 0.01 |
| Patesciba | Phylum | 33519.6 | 24053.1 | 0.1 | 0.1 | 0 | 0 | 28.8 | 16.9 | 0.03 | 0.01 |
| Saccharir | Class | 18853.2 | 12428.7 | 0.1 | 0 | 0 | 0 | 0 | 0 | 0 | 0 |
| Sacchari | Order | 18853.2 | 12428.7 | 0.1 | 0 | 0 | 0 | 0 | 0 | 0 | 0 |
| Saccha | Family | 14600.4 | 6682.9 | 0 | 0 | 0 | 0 | 0 | 0 | 0 | 0 |
| Saccha | Family | 2272.8 | 4254.6 | 0 | 0 | 0 | 0 | 0 | 0 | 0 | 0 |
| TM7a | Genus | 2272.8 | 4254.6 | 0 | 0 | 0 | 0 | 0 | 0 | 0 | 0 |
| YM | Family | 1980.1 | 1491.3 | 0 | 0 | 0 | 0 | 0 | 0 | 0 | 0 |
| 50 | Genus | 1980.1 | 1491.3 | 0 | 0 | 0 | 0 | 0 | 0 | 0 | 0 |
| ABY1 | Class | 9858.2 | 9953.5 | 0 | 0 | 0 | 0 | 60 | 40.9 | 0.02 | 0.01 |
| Candida | Order | 9858.2 | 9953.5 | 0 | 0 | 0 | 0 | 60 | 40.9 | 0.02 | 0.01 |
| Candid | Family | 9858.2 | 9953.5 | 0 | 0 | 0 | 0 | 60 | 40.9 | 0.02 | 0.01 |
| Graciliba | Class | 3750.1 | 0 | 0 | 0 | 0 | 0 | 100 | 0 | 0.01 | 0 |
| Graciliba | Order | 3750.1 | 0 | 0 | 0 | 0 | 0 | 100 | 0 | 0.01 | 0 |
| Patesciba | Class | 827 | 1074.8 | 0 | 0 | 0 | 0 | 0 | 0 | 0 | 0 |
| Parcubac | Class | 231.2 | 596.1 | 0 | 0 | 0 | 0 | 0 | 0 | 0 | 0 |
| Candida | Order | 231.2 | 596.1 | 0 | 0 | 0 | 0 | 0 | 0 | 0 | 0 |
| Candid | Family | 231.2 | 596.1 | 0 | 0 | 0 | 0 | 0 | 0 | 0 | 0 |
| Dependen | Phylum | 29566.2 | 23088.3 | 0.1 | 0.1 | 45.7 | 70.3 | 14.5 | 8 | 0.05 | 0.05 |

| | | | | | | | | | | | |
|---|---|---|---|---|---|---|---|---|---|---|---|
| Babeliae | Class | 29566.2 | 23088.3 | 0.1 | 0.1 | 45.7 | 70.3 | 14.5 | 15.3 | 0.05 | 0.06 |
| Babelial | Order | 29566.2 | 23088.3 | 0.1 | 0.1 | 45.7 | 70.3 | 14.5 | 15.3 | 0.05 | 0.06 |
| Vermipl | Family | 15164.6 | 22038 | 0 | 0.1 | 59.8 | 73.7 | 18.1 | 16 | 0.03 | 0.06 |
| Vermip | Genus | 15164.6 | 22038 | 0 | 0.1 | 59.8 | 73.7 | 18.1 | 16 | 0.03 | 0.06 |
| Babelia | Family | 12909.7 | 237.5 | 0 | 0 | 34.4 | 0 | 11.9 | 0 | 0.02 | 0 |
| UBA124 | Family | 1491.9 | 812.8 | 0 | 0 | 0 | 0 | 0 | 0 | 0 | 0 |
| UBA12 | Genus | 1491.9 | 812.8 | 0 | 0 | 0 | 0 | 0 | 0 | 0 | 0 |
| Abditibact | Phylum | 22430.6 | 23474.2 | 0.1 | 0.1 | 0 | 0 | 6.5 | 5.2 | 0 | 0 |
| Abditibac | Class | 22430.6 | 23474.2 | 0.1 | 0.1 | 0 | 0 | 6.5 | 5.2 | 0 | 0 |
| Abditiba | Order | 22430.6 | 23474.2 | 0.1 | 0.1 | 0 | 0 | 6.5 | 5.2 | 0 | 0 |
| Abditiba | Family | 22430.6 | 23474.2 | 0.1 | 0.1 | 0 | 0 | 6.5 | 5.2 | 0 | 0 |
| Abditib | Genus | 22430.6 | 23474.2 | 0.1 | 0.1 | 0 | 0 | 6.5 | 5.2 | 0 | 0 |
| Deinococc | Phylum | 16019 | 32354.5 | 0 | 0.1 | 0 | 0 | 0 | 0 | 0 | 0 |
| Deinococ | Class | 16019 | 32354.5 | 0 | 0.1 | 0 | 0 | 0 | 0 | 0 | 0 |
| Deinoco | Order | 16019 | 32354.5 | 0 | 0.1 | 0 | 0 | 0 | 0 | 0 | 0 |
| Trueper | Family | 15910.5 | 15439.3 | 0 | 0 | 0 | 0 | 0 | 0 | 0 | 0 |
| Truepe | Genus | 15910.5 | 15439.3 | 0 | 0 | 0 | 0 | 0 | 0 | 0 | 0 |
| Deinoc | Family | 108.5 | 16915.3 | 0 | 0.1 | 0 | 0 | 0 | 0 | 0 | 0 |
| Deinoc | Genus | 108.5 | 16915.3 | 0 | 0.1 | 0 | 0 | 0 | 0 | 0 | 0 |
| SAR324 cl | Phylum | 14019.7 | 12474.4 | 0 | 0 | 0 | 0 | 0 | 27.1 | 0 | 0.01 |
| SAR324 c | Class | 14019.7 | 12474.4 | 0 | 0 | 0 | 0 | 0 | 27.1 | 0 | 0.01 |
| Latesciba | Phylum | 12883.9 | 18018.4 | 0 | 0.1 | 0 | 0 | 0 | 0 | 0 | 0 |
| Latesciba | Class | 12883.9 | 18018.4 | 0 | 0.1 | 0 | 0 | 0 | 0 | 0 | 0 |
| Fibrobacte | Phylum | 5189.8 | 2199.5 | 0 | 0 | 0 | 0 | 0 | 0 | 0 | 0 |
| Fibrobact | Class | 5189.8 | 2199.5 | 0 | 0 | 0 | 0 | 0 | 0 | 0 | 0 |
| Fibrobac | Order | 5189.8 | 2199.5 | 0 | 0 | 0 | 0 | 0 | 0 | 0 | 0 |
| Fibroba | Family | 5189.8 | 2199.5 | 0 | 0 | 0 | 0 | 0 | 0 | 0 | 0 |
| possib | Genus | 3233.8 | 1421.4 | 0 | 0 | 0 | 0 | 0 | 0 | 0 | 0 |
| Fibrob | Genus | 1956 | 778.1 | 0 | 0 | 0 | 0 | 0 | 0 | 0 | 0 |
| Elusimicro | Phylum | 3166.4 | 4688.6 | 0 | 0 | 0 | 0 | 0 | 0 | 0 | 0 |
| Lineage II | Class | 2453.7 | 3457.1 | 0 | 0 | 0 | 0 | 0 | 0 | 0 | 0 |

| | | | | | | | | | | | |
|---|---|---|---|---|---|---|---|---|---|---|---|
| Lineage | Order | 2453.7 | 3457.1 | 0 | 0 | 0 | 0 | 0 | 0 | 0 | 0 |
| Elusimicr | Class | 712.7 | 1231.4 | 0 | 0 | 0 | 0 | 0 | 0 | 0 | 0 |
| MVP-88 | Order | 712.7 | 1231.4 | 0 | 0 | 0 | 0 | 0 | 0 | 0 | 0 |
| MVP-88 | Family | 712.7 | 1231.4 | 0 | 0 | 0 | 0 | 0 | 0 | 0 | 0 |
| MBNT15 | Phylum | 2995.1 | 1482.5 | 0 | 0 | 72.6 | 0 | 0 | 0 | 0.01 | 0 |
| MBNT15_ | Class | 2995.1 | 1482.5 | 0 | 0 | 72.6 | 0 | 0 | 0 | 0.01 | 0 |
| Sumerlaec | Phylum | 1739.2 | 1014.1 | 0 | 0 | 0 | 0 | 0 | 0 | 0 | 0 |
| Sumerlae | Class | 1739.2 | 1014.1 | 0 | 0 | 0 | 0 | 0 | 0 | 0 | 0 |
| Sumerla | Order | 1739.2 | 1014.1 | 0 | 0 | 0 | 0 | 0 | 0 | 0 | 0 |
| Sumerla | Family | 1739.2 | 1014.1 | 0 | 0 | 0 | 0 | 0 | 0 | 0 | 0 |
| Sumer | Genus | 1739.2 | 1014.1 | 0 | 0 | 0 | 0 | 0 | 0 | 0 | 0 |
| RCP2-54 | Phylum | 1519.6 | 2539 | 0 | 0 | 0 | 0 | 0 | 0 | 0 | 0 |
| RCP2-54_ | Class | 1519.6 | 2539 | 0 | 0 | 0 | 0 | 0 | 0 | 0 | 0 |
| FCPU426 | Phylum | 1330 | 0 | 0 | 0 | 0 | 0 | 0 | 0 | 0 | 0 |
| FCPU426 | Class | 1330 | 0 | 0 | 0 | 0 | 0 | 0 | 0 | 0 | 0 |
| Nitrospirot | Phylum | 122.7 | 81.6 | 0 | 0 | 0 | 0 | 0 | 0 | 0 | 0 |
| Nitrospiri | Class | 122.7 | 81.6 | 0 | 0 | 0 | 0 | 0 | 0 | 0 | 0 |
| Nitrospir | Order | 122.7 | 81.6 | 0 | 0 | 0 | 0 | 0 | 0 | 0 | 0 |
| Nitrosp | Family | 122.7 | 81.6 | 0 | 0 | 0 | 0 | 0 | 0 | 0 | 0 |
| Nitrosp | Genus | 122.7 | 81.6 | 0 | 0 | 0 | 0 | 0 | 0 | 0 | 0 |
