## Supplemental Table S4 for "Potato foliar infection with *Phytophthora infestans* drives strong, cultivar-specific shifts in rhizosphere communities"

| | Genus | strain | score_SG | score_ZG | Cultivar | Compartment | ASV | Phyllosphere | Rhizosphere | Soil | Gen 1 | Gen 2 | Bintje | Innovator |
|---|---|---|---|---|---|---|---|---|---|---|---|---|---|---|
| Actinobacteriota | Rhodococcus | AS109 | 0.25 | 0.83 | Innovator | rhizo_soil | asv003126 | EQUAL | EQUAL | EQUAL | EQUAL | EQUAL | EQUAL | EQUAL |
| | Agromyces | AR021 | 0.43 | 1.16 | Innovator | root | asv000372 | ABSENT | EQUAL | EQUAL | EQUAL | EQUAL | EQUAL | EQUAL |
| | | BR121 | 1.05 | 1.33 | Bintje | root | | | | | | | | |
| | Microbacterium | AS086 | 0.97 | 1.00 | Innovator | rhizo_soil | asv002271 | EQUAL | EQUAL | EQUAL | EQUAL | EQUAL | EQUAL | EQUAL |
| | | BR064 | 1.16 | 1.14 | Bintje | root | asv002346 | EQUAL | EQUAL | EQUAL | EQUAL | EQUAL | EQUAL | EQUAL |
| | | AR022 | 0.48 | 0.91 | Innovator | root | asv006178 | EQUAL | EQUAL | EQUAL | EQUAL | EQUAL | EQUAL | EQUAL |
| | Arthrobacter | BS170 | 0.77 | 0.91 | Bintje | rhizo_soil | asv003790 | EQUAL | EQUAL | EQUAL | EQUAL | EQUAL | EQUAL | EQUAL |
| | | BS230 | 0.55 | 0.68 | Bintje | rhizo_soil | asv004743 | EQUAL | EQUAL | EQUAL | EQUAL | EQUAL | EQUAL | EQUAL |
| | Paenarthrobacter | AR070 | 0.49 | 0.68 | Innovator | root | asv000575 | EQUAL | EQUAL | LESS | EQUAL | LESS | LESS | EQUAL |
| | | AS049 | 0.32 | 0.58 | Innovator | rhizo_soil | asv003625 | ABSENT | EQUAL | EQUAL | EQUAL | EQUAL | EQUAL | EQUAL |
| | Promicromonospo. | BS107 | 0.18 | 0.92 | Bintje | rhizo_soil | asv000134 | ABSENT | EQUAL | EQUAL | EQUAL | EQUAL | EQUAL | EQUAL |
| | Micromonospora | BL047b | 0.55 | 1.01 | Bintje | leaf | asv003269 | ABSENT | EQUAL | EQUAL | EQUAL | EQUAL | EQUAL | EQUAL |
| | | BL047c | 0.69 | 1.04 | Bintje | leaf | | | | | | | | |
| | Nocardioides | AR010 | 0.26 | 0.87 | Innovator | root | asv000395 | ABSENT | EQUAL | EQUAL | EQUAL | EQUAL | EQUAL | EQUAL |
| | | AR046 | 0.19 | 1.03 | Innovator | root | | | | | | | | |
| | | AR051 | 0.27 | 0.89 | Innovator | root | | | | | | | | |
| | | AS010 | 0.10 | 1.02 | Innovator | rhizo_soil | | | | | | | | |
| | | AS106 | 0.07 | 1.22 | Innovator | rhizo_soil | | | | | | | | |
| | | AS108 | 0.15 | 0.96 | Innovator | rhizo_soil | | | | | | | | |
| | | AS121b | 0.19 | 0.93 | Innovator | rhizo_soil | | | | | | | | |
| | | BR081 | 0.25 | 0.94 | Bintje | root | | | | | | | | |
| | | BS088 | 0.49 | 1.08 | Bintje | rhizo_soil | | | | | | | | |
| | Nocardioides | AS133 | 0.12 | 0.98 | Innovator | rhizo_soil | asv001872 | ABSENT | EQUAL | EQUAL | EQUAL | EQUAL | EQUAL | EQUAL |
| | | BL047a | 0.11 | 1.10 | Bintje | leaf | | | | | | | | |
| | | BS157 | 0.32 | 1.29 | Bintje | rhizo_soil | asv002615 | EQUAL | EQUAL | LESS | EQUAL | LESS | LESS | EQUAL |
| | | AR039 | 0.38 | 0.79 | Innovator | root | | | | | | | | |
| | | AS001 | 0.48 | 0.75 | Innovator | rhizo_soil | | | | | | | | |
| | | AS013 | 0.43 | 0.73 | Innovator | rhizo_soil | | | | | | | | |
| | | AS035 | 0.48 | 0.76 | Innovator | rhizo_soil | | | | | | | | |
| | | BL074 | 0.30 | 0.75 | Bintje | leaf | | | | | | | | |
| | | BL096 | 0.50 | 0.69 | Bintje | leaf | | | | | | | | |
| | | BR014a | 0.93 | 1.45 | Bintje | root | | | | | | | | |

| | | | | | | | | | | | | | | |
|---|---|---|---|---|---|---|---|---|---|---|---|---|---|---|
| Firmicutes | Bacillus | BR066 | 0.37 | 0.89 | Bintje | root | asv000047 | EQUAL | MORE | EQUAL | MORE | EQUAL | MORE | EQUAL |
| | | BS139 | 0.53 | 0.65 | Bintje | rhizo_soil | | | | | | | | |
| | | BS150 | 0.31 | 0.87 | Bintje | rhizo_soil | | | | | | | | |
| | | BS177 | 0.43 | 0.87 | Bintje | rhizo_soil | | | | | | | | |
| | | BS199 | 0.30 | 0.73 | Bintje | rhizo_soil | | | | | | | | |
| | | BS200 | 0.46 | 0.78 | Bintje | rhizo_soil | | | | | | | | |
| | | BS222 | 1.01 | 0.83 | Bintje | rhizo_soil | | | | | | | | |
| | | BS232 | 0.35 | 0.80 | Bintje | rhizo_soil | | | | | | | | |
| | | BS236 | 0.47 | 0.81 | Bintje | rhizo_soil | | | | | | | | |
| | | AR001 | 0.45 | 0.68 | Innovator | root | asv000068 | EQUAL | LESS | MORE | LESS | MORE | LESS | MORE |
| | | AR002 | 0.73 | 1.10 | Innovator | root | | | | | | | | |
| | | AR005 | 0.20 | 0.54 | Innovator | root | | | | | | | | |
| | | AR042 | 0.21 | 0.54 | Innovator | root | | | | | | | | |
| | | AR057 | 0.28 | 0.66 | Innovator | root | | | | | | | | |
| | | AS022 | 0.11 | 0.80 | Innovator | rhizo_soil | | | | | | | | |
| | | AS029 | 0.14 | 0.93 | Innovator | rhizo_soil | | | | | | | | |
| | | AS040 | 0.27 | 0.64 | Innovator | rhizo_soil | | | | | | | | |
| | | BL049 | 0.18 | 1.22 | Bintje | leaf | | | | | | | | |
| | | BR107 | 0.39 | 0.56 | Bintje | root | | | | | | | | |
| | | BS196 | 0.39 | 1.17 | Bintje | rhizo_soil | | | | | | | | |
| | | BS132 | 0.77 | 1.59 | Bintje | rhizo_soil | asv000070 | EQUAL | LESS | MORE | LESS | MORE | AMBIG. | EQUAL |
| | | AL004 | 0.22 | 0.83 | Innovator | leaf | asv000077 | EQUAL | LESS | MORE | LESS | MORE | AMBIG. | EQUAL |
| | | AR037 | 0.44 | 0.85 | Innovator | root | | | | | | | | |
| | | BR134 | 0.17 | 0.56 | Bintje | root | | | | | | | | |
| | | AS146 | 0.39 | 0.92 | Innovator | rhizo_soil | asv000244 | EQUAL | LESS | MORE | LESS | MORE | AMBIG. | EQUAL |
| | | BS102 | 1.59 | 0.68 | Bintje | rhizo_soil | asv000285 | EQUAL | LESS | EQUAL | LESS | EQUAL | LESS | EQUAL |
| | | AS123 | 0.94 | 1.32 | Innovator | rhizo_soil | asv000843 | EQUAL | EQUAL | EQUAL | EQUAL | EQUAL | EQUAL | EQUAL |
| | | AS021 | 0.73 | 1.26 | Innovator | rhizo_soil | asv001219 | EQUAL | MORE | EQUAL | EQUAL | MORE | EQUAL | MORE |
| | | AS144 | 1.03 | 1.08 | Innovator | rhizo_soil | | | | | | | | |
| | | BR088 | 0.46 | 0.84 | Bintje | root | | | | | | | | |
| | | BS131 | 0.67 | 1.41 | Bintje | rhizo_soil | | | | | | | | |
| | | BR136 | 0.99 | 1.16 | Bintje | root | asv002388 | ABSENT | EQUAL | EQUAL | EQUAL | EQUAL | EQUAL | EQUAL |
| | | BS113 | 0.24 | 0.72 | Bintje | rhizo_soil | | | | | | | | |
| | | AL028 | 0.62 | 0.98 | Innovator | leaf | asv002748 | EQUAL | EQUAL | EQUAL | EQUAL | EQUAL | EQUAL | EQUAL |
| | | BS135 | 0.26 | 0.78 | Bintje | rhizo_soil | asv004823 | ABSENT | EQUAL | EQUAL | EQUAL | EQUAL | EQUAL | EQUAL |
| | Lysinibacillus | BR127 | 0.40 | 0.63 | Bintje | root | asv007669 | ABSENT | EQUAL | EQUAL | EQUAL | EQUAL | EQUAL | EQUAL |

| | | | | | | | | | | | | | |
|---|---|---|---|---|---|---|---|---|---|---|---|---|---|
| Paenisporosarci. | AR095 | 0.58 | 0.86 | Innovator | root | asv000216 | EQUAL | EQUAL | EQUAL | EQUAL | EQUAL | EQUAL | EQUAL |
| | BS179 | 0.73 | 1.11 | Bintje | rhizo_soil | | | | | | | | |
| | BS106 | 0.64 | 1.21 | Bintje | rhizo_soil | asv000945 | EQUAL | EQUAL | LESS | EQUAL | LESS | LESS | EQUAL |
| Psychrobacillus | BS080 | 0.70 | 1.12 | Bintje | rhizo_soil | asv003647 | ABSENT | EQUAL | EQUAL | EQUAL | EQUAL | EQUAL | EQUAL |
| Brevibacillus | AS028 | 0.81 | 0.79 | Innovator | rhizo_soil | asv004156 | ABSENT | EQUAL | EQUAL | EQUAL | EQUAL | EQUAL | EQUAL |
| Kaistia | BS237 | 0.78 | 0.98 | Bintje | rhizo_soil | asv002208 | ABSENT | EQUAL | EQUAL | ABSENT | EQUAL | EQUAL | EQUAL |
| Allorhizobium_N. | AL034 | 0.38 | 0.38 | Innovator | leaf | asv003204 | EQUAL | EQUAL | EQUAL | EQUAL | EQUAL | EQUAL | EQUAL |
| | AL039 | 0.41 | 0.46 | Innovator | leaf | | | | | | | | |
| | AR061 | 0.52 | 0.38 | Innovator | root | | | | | | | | |
| | AL064 | 0.46 | 0.57 | Innovator | leaf | asv005149 | EQUAL | EQUAL | EQUAL | EQUAL | EQUAL | EQUAL | EQUAL |
| | AR077 | 0.61 | 0.61 | Innovator | root | | | | | | | | |
| | AR081 | 0.57 | 0.50 | Innovator | root | | | | | | | | |
| | AR083 | 0.30 | 0.57 | Innovator | root | | | | | | | | |
| | AR116 | 0.43 | 0.54 | Innovator | root | | | | | | | | |
| | AS054 | 0.30 | 0.64 | Innovator | rhizo_soil | | | | | | | | |
| | AS058 | 0.44 | 0.61 | Innovator | rhizo_soil | | | | | | | | |
| | AS064 | 0.48 | 0.46 | Innovator | rhizo_soil | | | | | | | | |
| | AS070 | 0.36 | 0.54 | Innovator | rhizo_soil | | | | | | | | |
| | BL067 | 0.59 | 0.77 | Bintje | leaf | | | | | | | | |
| | BL071 | 0.43 | 0.76 | Bintje | leaf | | | | | | | | |
| | BR126 | 0.52 | 0.71 | Bintje | root | | | | | | | | |
| | BR150 | 0.61 | 0.68 | Bintje | root | | | | | | | | |
| | BR153 | 0.54 | 0.84 | Bintje | root | | | | | | | | |
| Ensifer | BS138 | 0.54 | 1.08 | Bintje | rhizo_soil | asv004665 | ABSENT | MORE | MORE | EQUAL | MORE | MORE | EQUAL |
| Fa_Rhizobiaceae | BS141 | 0.53 | 0.88 | Bintje | rhizo_soil | asv003045 | ABSENT | EQUAL | EQUAL | EQUAL | EQUAL | EQUAL | EQUAL |
| Ochrobactrum | BR046 | 0.08 | 0.13 | Bintje | root | asv002231 | ABSENT | EQUAL | EQUAL | EQUAL | EQUAL | EQUAL | EQUAL |
| | AR056 | 0.99 | 1.19 | Innovator | root | asv004651 | ABSENT | EQUAL | EQUAL | EQUAL | EQUAL | EQUAL | EQUAL |
| Phyllobacterium | AR008 | 0.52 | 1.05 | Innovator | root | asv000475 | ABSENT | EQUAL | MORE | MORE | EQUAL | EQUAL | MORE |
| | AS006 | 0.17 | 0.94 | Innovator | rhizo_soil | | | | | | | | |
| | AS056 | 0.35 | 0.61 | Innovator | rhizo_soil | | | | | | | | |
| | AR177 | 0.05 | 0.75 | Innovator | root | asv002046 | ABSENT | EQUAL | EQUAL | EQUAL | EQUAL | EQUAL | EQUAL |
| | AR179 | 0.18 | 0.62 | Innovator | root | | | | | | | | |
| | AS178 | 0.15 | 0.94 | Innovator | rhizo_soil | | | | | | | | |
| | BR015 | 0.40 | 0.91 | Bintje | root | | | | | | | | |
| | BR099 | 0.28 | 0.70 | Bintje | root | | | | | | | | |
| | BS071 | 0.10 | 0.95 | Bintje | rhizo_soil | | | | | | | | |

| | | | | | | | | | | | | | | |
|---|---|---|---|---|---|---|---|---|---|---|---|---|---|---|
| Proteobacteria | Achromobacter | AS060 | 0.57 | 0.90 | Innovator | rhizo_soil | asv005242 | EQUAL | MORE | EQUAL | EQUAL | MORE | MORE | EQUAL |
| | Advenella | AR097 | 0.21 | 1.18 | Innovator | root | asv000086 | ABSENT | EQUAL | EQUAL | EQUAL | EQUAL | EQUAL | EQUAL |
| | | AS050 | 0.21 | 0.96 | Innovator | rhizo_soil | | | | | | | | |
| | | AS085 | 0.16 | 0.81 | Innovator | rhizo_soil | | | | | | | | |
| | | BR157 | 0.27 | 1.51 | Bintje | root | | | | | | | | |
| | Fa_Alcaligenaceae | BS103 | 0.38 | 0.71 | Bintje | rhizo_soil | asv000375 | ABSENT | EQUAL | EQUAL | EQUAL | EQUAL | EQUAL | EQUAL |
| | | AR063 | 0.47 | 0.67 | Innovator | root | asv002448 | EQUAL | MORE | EQUAL | EQUAL | MORE | MORE | EQUAL |
| | Pusillimonas | BR094 | 0.62 | 0.67 | Bintje | root | asv000375 | ABSENT | EQUAL | EQUAL | EQUAL | EQUAL | EQUAL | EQUAL |
| | | BR144 | 0.41 | 0.27 | Bintje | root | | | | | | | | |
| | | BS231 | 0.41 | 0.96 | Bintje | rhizo_soil | | | | | | | | |
| | Acidovorax | AR150 | 0.64 | 0.51 | Innovator | root | asv000049 | ABSENT | MORE | MORE | MORE | EQUAL | EQUAL | MORE |
| | | BR010 | 0.87 | 0.57 | Bintje | root | | | | | | | | |
| | | BR101 | 0.56 | 0.85 | Bintje | root | | | | | | | | |
| | | BS181 | 0.39 | 0.86 | Bintje | rhizo_soil | | | | | | | | |
| | | BS192 | 0.42 | 0.43 | Bintje | rhizo_soil | | | | | | | | |
| | | AL005 | 0.57 | 0.93 | Innovator | leaf | asv000438 | ABSENT | MORE | EQUAL | MORE | MORE | EQUAL | MORE |
| | | AR009 | 0.54 | 0.96 | Innovator | root | | | | | | | | |
| | | BR112 | 0.53 | 0.66 | Bintje | root | | | | | | | | |
| | | BR133 | 0.38 | 0.34 | Bintje | root | | | | | | | | |
| | | BR013 | 0.81 | 1.36 | Bintje | root | asv001884 | EQUAL | MORE | MORE | EQUAL | MORE | MORE | MORE |
| | Diaphorobacter | AR118 | 0.61 | 0.49 | Innovator | root | asv003333 | ABSENT | MORE | MORE | ABSENT | MORE | MORE | EQUAL |
| | | BR049 | 0.58 | 0.89 | Bintje | root | | | | | | | | |
| | | BR050 | 0.17 | 0.85 | Bintje | root | | | | | | | | |
| | | BR156 | 0.16 | 0.61 | Bintje | root | | | | | | | | |
| | | BS123 | 0.79 | 0.52 | Bintje | rhizo_soil | | | | | | | | |
| | Fa_Comamonadaceae | AR132 | 0.38 | 0.54 | Innovator | root | asv001155 | ABSENT | EQUAL | MORE | EQUAL | MORE | MORE | EQUAL |
| | Hydrogenophaga | BR109 | 0.47 | 0.93 | Bintje | root | asv000670 | ABSENT | EQUAL | EQUAL | EQUAL | EQUAL | EQUAL | EQUAL |
| | | AR076 | 0.40 | 0.66 | Innovator | root | asv000668 | ABSENT | EQUAL | MORE | EQUAL | MORE | MORE | EQUAL |
| | | AR143 | 0.70 | 0.61 | Innovator | root | | | | | | | | |
| | | AR144 | 0.48 | 0.73 | Innovator | root | | | | | | | | |
| | | AR176 | 0.78 | 0.79 | Innovator | root | | | | | | | | |
| | | AR178 | 0.65 | 0.79 | Innovator | root | | | | | | | | |
| | | AR190 | 0.31 | 0.49 | Innovator | root | | | | | | | | |
| | | AS024 | 0.46 | 0.66 | Innovator | rhizo_soil | | | | | | | | |
| | | AS026 | 0.83 | 0.77 | Innovator | rhizo_soil | | | | | | | | |
| | | AS181 | 0.60 | 0.75 | Innovator | rhizo_soil | | | | | | | | |

| | | | | | | | | | | | | | |
|---|---|---|---|---|---|---|---|---|---|---|---|---|---|
| | AS182 | 0.71 | 1.15 | Innovator | rhizo_soil | | | | | | | | |
| Variovorax | BR034 | 0.73 | 0.60 | Bintje | root | | | | | | | | |
| | BS037 | 0.48 | 0.91 | Bintje | rhizo_soil | | | | | | | | |
| | BS038 | 0.25 | 0.29 | Bintje | rhizo_soil | | | | | | | | |
| | BS061 | 0.33 | 0.78 | Bintje | rhizo_soil | | | | | | | | |
| | BS158 | 0.42 | 0.62 | Bintje | rhizo_soil | | | | | | | | |
| | AR201 | 0.41 | 0.75 | Innovator | root | asv001155 | ABSENT | EQUAL | MORE | EQUAL | MORE | MORE | EQUAL |
| | AS093 | 0.21 | 0.93 | Innovator | rhizo_soil | | | | | | | | |
| | AR032 | 0.57 | 0.71 | Innovator | root | | | | | | | | |
| | AS027 | 0.44 | 1.06 | Innovator | rhizo_soil | asv002573 | ABSENT | MORE | MORE | EQUAL | MORE | MORE | MORE |
| | AS080 | 0.64 | 0.67 | Innovator | rhizo_soil | | | | | | | | |
| | AS180 | 0.63 | 0.93 | Innovator | rhizo_soil | | | | | | | | |
| Fa_Enterobacteriac. | AR093 | 0.21 | 0.80 | Innovator | root | asv000512 | ABSENT | EQUAL | EQUAL | EQUAL | EQUAL | EQUAL | EQUAL |
| | BS119 | 1.05 | 1.54 | Bintje | rhizo_soil | asv000051 | EQUAL | EQUAL | MORE | EQUAL | MORE | MORE | EQUAL |
| | BS130 | 0.29 | 0.20 | Bintje | rhizo_soil | | | | | | | | |
| | AS044 | 0.27 | 0.35 | Innovator | rhizo_soil | asv000300 | EQUAL | EQUAL | MORE | EQUAL | MORE | MORE | EQUAL |
| | AR071 | 0.42 | 0.57 | Innovator | root | | | | | | | | |
| Pseudomonas | AR088 | 0.25 | 0.03 | Innovator | root | asv000701 | ABSENT | MORE | MORE | EQUAL | MORE | MORE | MORE |
| | AR125 | 0.20 | 0.08 | Innovator | root | | | | | | | | |
| | AR168 | 0.47 | 0.05 | Innovator | root | | | | | | | | |
| | AR060 | 0.51 | 0.14 | Innovator | root | asv001047 | ABSENT | MORE | MORE | EQUAL | MORE | MORE | MORE |
| | BR058 | 0.16 | 0.68 | Bintje | root | | | | | | | | |
| | AR075 | 0.33 | 0.73 | Innovator | root | asv004400 | EQUAL | EQUAL | EQUAL | EQUAL | EQUAL | EQUAL | EQUAL |
